# The ERGtools2 package: A Toolset for Processing and Analysing Visual Electrophysiology Data

**DOI:** 10.1101/2024.08.27.609856

**Authors:** Moritz Lindner

## Abstract

**Introduction:** Visual electrophysiology techniques play a key role in assessing the functional integrity of the visual pathway. Recent advancements and applications demand flexible analysis tools. Heren, ERGtools2, an open-source R package designed for processing and analysing visual electrophysiology data is introduced.

**Methods:** A dataset comprising Electroretinogram (ERG) recordings from both eyes of C57Bl/6J mice, subjected to standard ISCEV stimuli, was used to present the functionality of ERGtools2. ERGtools2 stores and organizes all recordings, metadata, and measurement information from individual examination in a single object, maintaining raw data throughout the analysis process.

**Results:** A standard workflow is presented exemplifying how ERGtools2 can be used to efficiently import, pre-process and analyse ERG data. Following this workflow, basic ERG measurements like *a* and *B*-wave amplitudes and visualisation of a single exam as well as group statistics are obtained. Moreover, special use cases are described, including for the handling of noisy data and the storage of data in the HDF5 format to ensure long-term preservation and accessibility. ERGtools2 also provides a graphical interface.

**Discussion:** ERGtools2 provides a comprehensive, flexible, and device-independent solution for visual electrophysiology data analysis. Its emphasis on maintaining raw data integrity, combined with advanced processing and analysis capabilities, makes it a useful tool for preclinical and clinical research applications. The open-source nature and the use of open data formats promote reproducibility and data sharing in visual neurosciences.

## Introduction

Visual electrophysiology techniques, including electroretinography (ERG) and visually evoked potentials (VEP) enable a direct functional assessment of the visual pathway, from photoreceptors to the visual cortex [9; 16; 10; 15; 7].

Over recent years the clinical use of electrophysiology techniques has evolved. Refined retinal imaging techniques and genetic testing [3; 4] now offer alternative or complementary approaches for differential diagnosis in retinal disease and optic neuropathy [4; 6] and current psychophysical techniques offer space resolved functional assessment [11; 1]. Visual electrophysiology, however, can dissect the functional integrity of the visual pathway at specific levels thus yielding information that cannot be obtained by any other techniques [9]. Thereby, electrophysiological findings may be pathognomonic for several conditions [4; 14; 5]. Functional changes as objectified by retinal electrophysiology may precede structural changes by years and even occur in conditions without any noticeable fundus changes [22]. Since vision prosthetic approaches are emerging visual electrophysiology has additionally gained attention as a functional objective endpoint [2] and in preclinical research visual electrophysiology is of particular relevance as laboratory mice, despite a sophisticated retinal circuity, make relatively little use of their visual function on behavioural level [8].

Especially more novel areas of application for visual electrophysiology now require for flexible analysis strategies. For instance, electrophysiological responses rescued or restored by a novel therapeutic approach might be too weak to be reliably measure using standard ERG/VEP readouts while still containing important evidence of the efficacy of that approach. In such cases, alternative analysis strategies like performing discrete Fourier transformations on Flicker ERGs may provide higher sensitivity [2]. Such analyses are however usually not possible using the software provided with the ERG/VEP device.

Moreover, shareability, reusability and long-term preservation of research data has moved into public focus. Taking measures to ensuring their reusability and long-term preservation is increasingly demanded by cost payers in research as well as clinical regulators and animal welfare authorities. These considerations are reflected in the FAIR Data Principles (FAIR: “Findable, Accessible, Interoperable and Reusable”, [21]). Despite recent efforts [17], in visual electrophysiology, both in research and clinics, a standard data format in line with the FAIR principles is yet not widely established. Anyone wishing to analyse visual electrophysiology data does not only need the access to the data but also to a (usually proprietary and) file format specific analysis software.

Here we present ERGtools2, an open-source framework for processing and analysing retinal electrophysiology data in R [13]. It comes with a command-line interface as well as a visual interface for efficient data processing and offers data storing in the open HDF5 format [20]. In this paper, technical aspects of the ERGtools2 package are briefly described in Methods and examples of its functionality are presented in Results. ERGtools2, in its version 0.8, is available as supplemental material (Supplemental Material 1) to this paper and the latest version can be downloaded from GitHub (https://github.com/moritzlindner/ERGtools2). Example code is provided in two supplementary files (Supplemental Material 2 and 3).

## Results

### A standard workflow

#### The test dataset

To demonstrate the functionality of ERGtools2, we utilized electroretinogram data recorded from the left eyes of five C57Bl/6J mice (median age 7.4 weeks, inter-quartile range: 7.1 – 9 weeks) in response to a stimulus protocol following ISCEV recommendations. The stimulus protocol consisted of a sequence of three dark-adapted and three light-adapted steps, with each step consisting of a single flash stimulus. Intensity ranges for dark-adapted stimuli were 0.001 cd*s/m^2^ to 3 cd*s/m^2^ and 1 cd*s/m^2^ to 10 cd*s/m^2^ for light adapted stimuli. Each step was repeated least three times.

A couple of actions are necessary in order to import such a dataset into ERGtools2 and perform measurements on it or visualize the data. These steps are summarized in **Figure 1 A** and described in detail below. The nomenclature of visual electrophysiology recordings as used herein is provided in the methods section.

**Figure 1:**
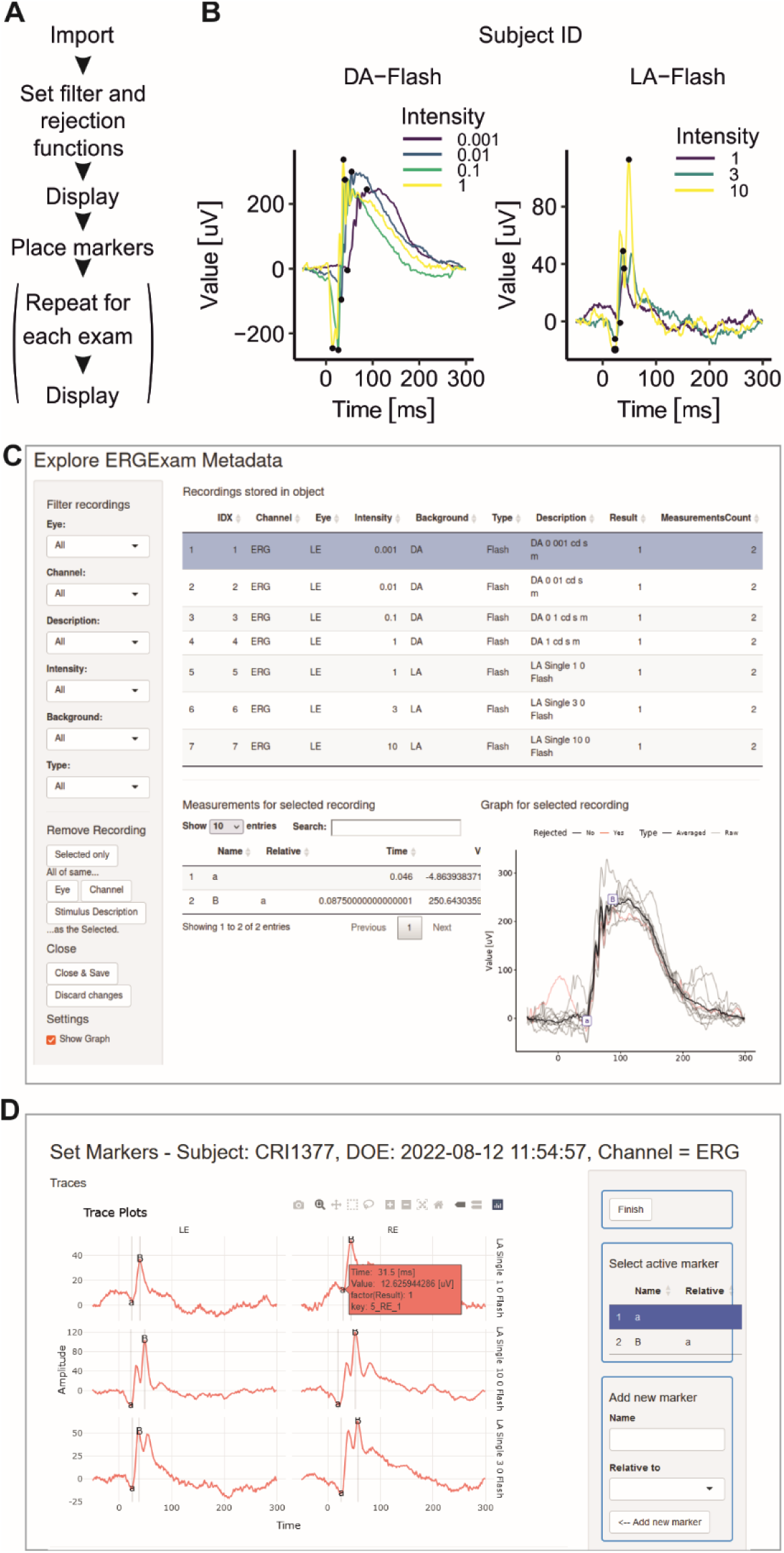
Visualization and manipulation of an ERGExam object using ERGtools2. **(A)** Standard workflow to import and process visual electrophysiology data using ERGtools2. **(B)** Visualization of an ERG exam from a left eye of a mouse as obtained by running *ggERGExam()*. Recordings are automatically colour coded by stimulus intensity and panelled out by adaptation state (“DA” and “LA”). **(C)** An interactive interface for viewing the content of an ERGExam object can be called typing *exploreERGExam().* Besides viewing, it also allows subsetting an individual ERGExam object. **(D)** An interactive interface for placing trace markers can be called via *interactiveMeasurements()*.

#### Importing data

ERGtools2 can import visual electrophysiology data from any file format that can be read into R. The first step in the process involves creating an EPhysData object, which stores the recording data along with the associated time trace. Data recorded from one single channel of a single eye will be stored into one such object. A new object will be created for the data from the next channel and eye, and so forth. This can be done using the *newEPhysData()* function. Once the individual recordings are imported as EPhysData objects, they are assembled into an ERGExam object using the *newERGExam()* function. In the ERGExam object the individual recordings are stored together with their descriptive metadata such as the recording laterality (e.g., left or right eye) and the channel type (e.g., ERG or VEP). Defining information for the stimuli used (e.g. adaptation state and intensity) as well as information on the subject, experimental cohort, and exam date is also stored in that object.

In practical terms, users first create a list of all their EPhysData objects, this list would then be passed to the *newERGExam()* method as the parameter *“Data”*. Metadata and stimulus information is passed on as the parameters “*Metadata*” and “*Stimulus*”, respectively. Each entry in the *Metadata* must be aligned with the corresponding entry in *Data*, so it’s crucial to ensure both are in the correct order. The stimulus information is matched to the *Data* by the Step column, which is an essential part of the *Metadata* table. Additionally, the user provides exam-related information (parameter: *ExamInfo*) in the form of a list containing at least the acquisition protocol name (*ProtocolName*) and the exam date (*ExamDate*). Subject details are provided through the *SubjectInfo* parameter, which is also a list that includes at least the subject’s name (*Subject*) and date of birth (*DOB*). A visual summary of the structure of the ERGExam object is shown in **Figure 2** and in the Methods section. Although setting up the import routine for the first time may require some work, once established, it can be reused for all subsequent exams, provided they are stored in the same format. R code illustrating two scenarios of such an import procedure is provided as supplemental material (Supplemental Material 2).

**Figure 2:**
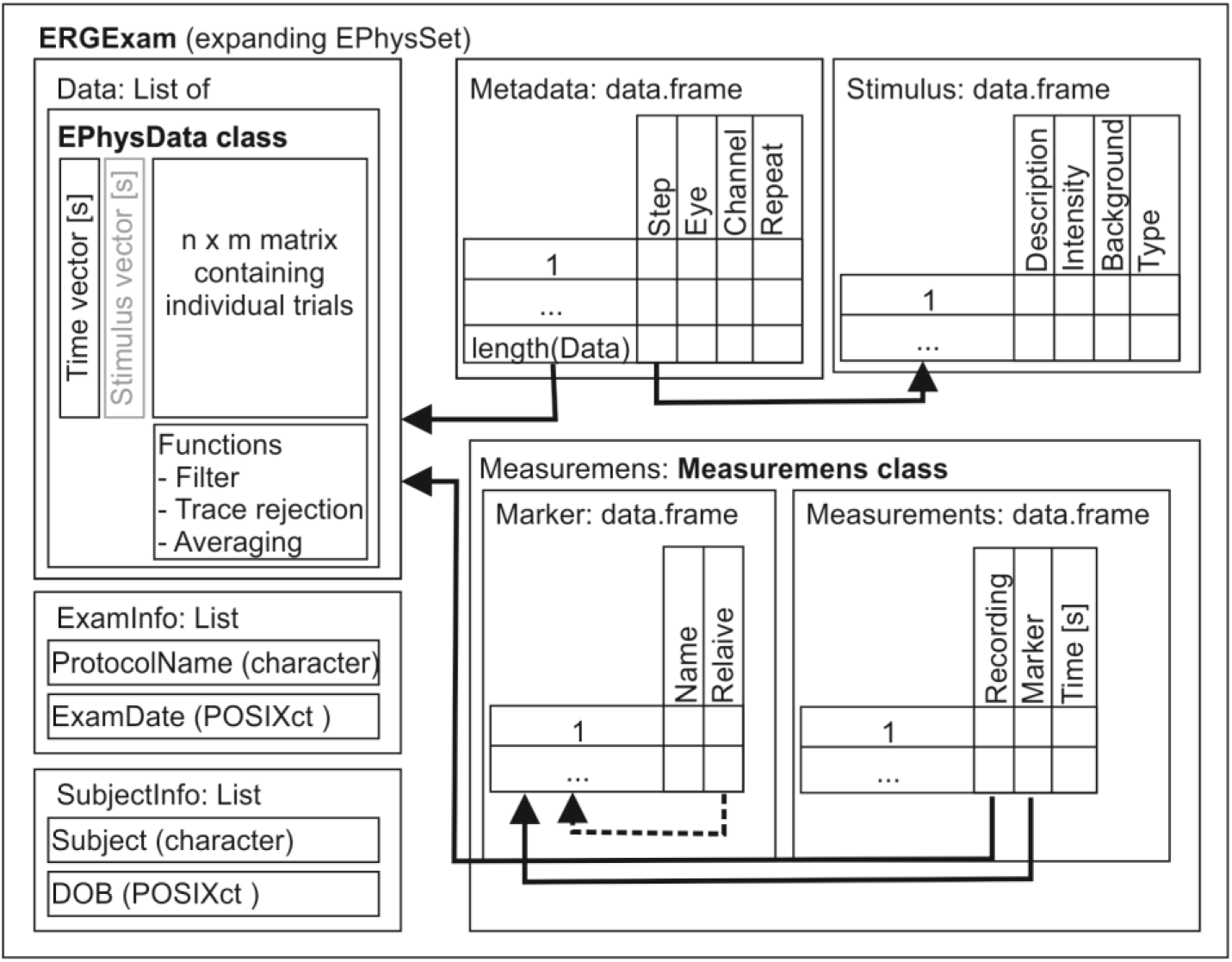
The structure of an ERGExam object, the container of visual electrophysiology data in ERGtools2. Details can be found in the main text as well as in R typing *“?ERGtools2::‘ERGtools2-package’”*.

Importantly, the raw data remains unaltered throughout the analysis process, ensuring data integrity. ERGtools2 also contains a convenience function for importing ERG exams from Diagnosys’s Espion software. Thus, with *ImportEspion()*,exams can be imported with hardly any coding effort.

#### Configuring the data object for analysis

After importing the recordings, the next step is to prepare the data for analysis. ERGtools2 includes functions for averaging repeated trials and filtering and outlier removal to improve the signal-to-noise ratio. The function *SetStandardFunctions()* is designed to streamline this process: For each recording in the object it sets a linear detrending filter. In case of repeated recordings, it then looks for outliers among those and marks them for exclusion. Finally, the averaging function is defined as *mean()*. These default settings work well for most standard applications. On special occasions, e.g. when specific bandpass filtering is desired to extract oscillatory potentials, this can be achieved using *FilterFunction()*. Additionally, ERGtools2 offers several alternative algorithms for identification and exclusion of outlier trials and user defined functions are equally supported.

Importantly, when such functions are set for an ERGExam object, validity checks are automatically performed to ensure their in-exam validity. For example, if a filter function is changed for a recording from the left eye, the same change must also be applied to the corresponding recording from the right eye as well.

#### Visualizing and subsetting the data

Once those preprocessing functions are set, the exam can be displayed using the *ggERGExam()* command, which panels out the recordings in the exam: Eyes in columns and channels and/or adaptation conditions in rows. The different steps, usually representing increasing stimulus intensities, are colour coded and plotted to the same panel (**Figure 1 B**). The user might now find that not all parts of the exam are of relevance for further analysis. Unnecessary recordings can be removed either at the command-line level using *Subset()* or *DropRecordings()* or interactively using *exploreERGExam()* (**Figure 1 C**).

#### Placing Markers for Measurement

Subsequently, one needs to position the markers for performing measurements on the recordings, e.g. timing and amplitude of characteristic waveforms. *AutoPlaceMarkers()* tries to automatically identify a and B wave positions on flash ERG traces as well as P1, N1, and P2 waves for VEP traces. Also, peak detection on flicker ERG traces is supported. *AutoPlaceMarkers()* is optimized for the waveform patterns expected in wild-type (C57Bl/6J) mice. Once the marker positions are set, these are also displayed when calling *ggERGExam()* and a table containing timing and amplitudes for all markers can be retrieved calling *Measurements()*. If the automatic marker placement did not provide satisfactory results, interactive marker placement is possible using *interactiveMeasurements()* (**Figure 1 D**).

#### Summarizing and analysing a set of exams

In a research setting, an investigator may want to compare between exams acquired from subjects belonging to different experimental cohorts. To achieve this, ERGtools2 contains a set of functions that can handle lists of ERGExam objects e.g. to plot the exams side-by-side (*ggPlotRecordings()*), draw intensity response curves (*ggIntensitySequence(),* **Figure 3**), or simply obtain measurements for specific markers for all recordings in the list (*CollectMeasurements()*). The latter could then be used for any downstream analysis, e.g. to for descriptive statistics.

**Figure 3:**
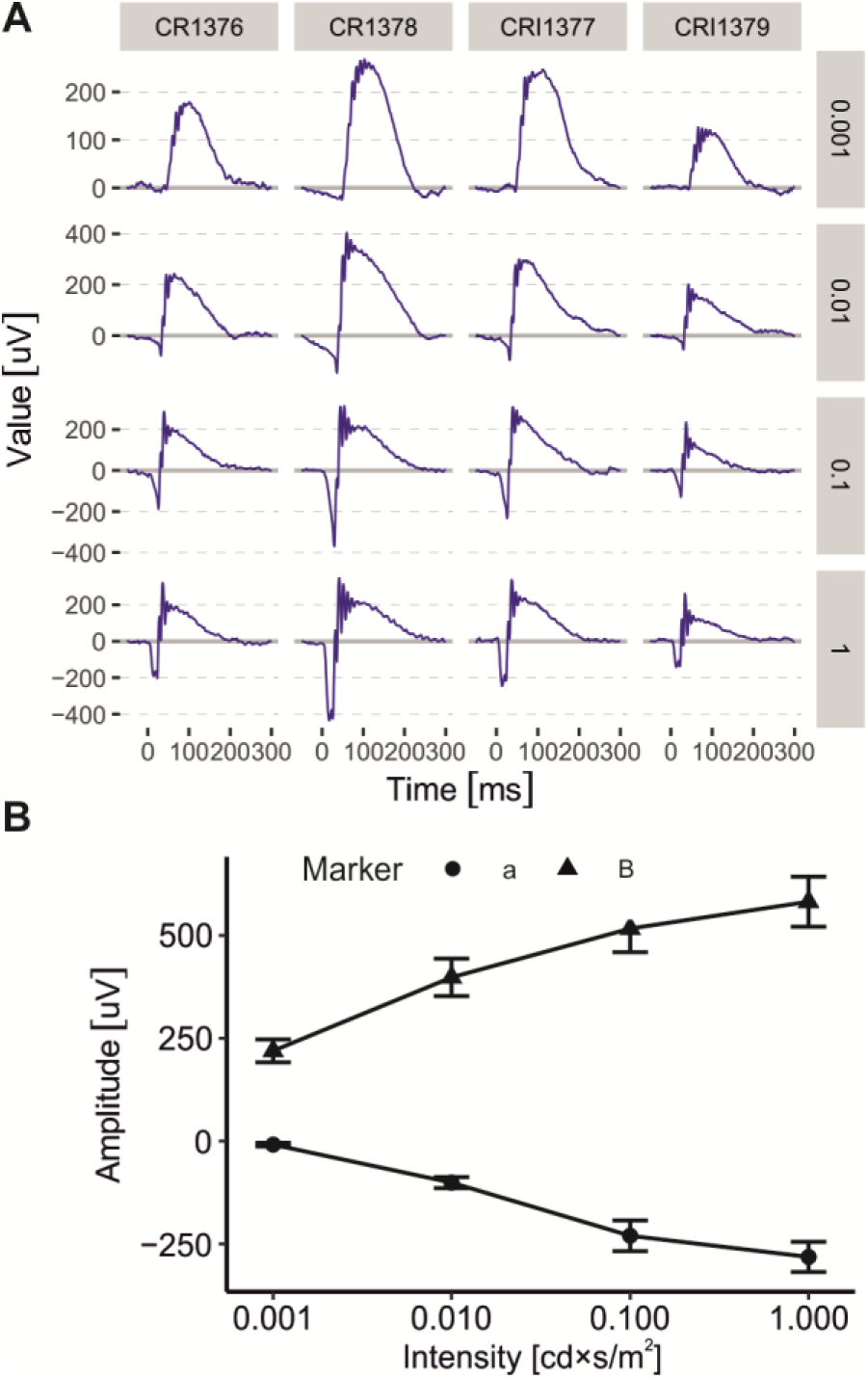
Visualization of multiple ERG exams. **(A)** Automatic visualization of a set of visual electrophysiology exams using *ggPlotRecordings()*. Individual traces are panelled out by subject (columns) and stimulus conditions (rows). Only recordings for dark-adapted stimuli are shown. **(B)** Summary intensity-response curves for a and B-wave amplitudes from the exams shown in (A), visualized using *ggIntensitySequence()*.

### Special use cases

#### Automatic rejection of outlier recordings

One particularly challenge about processing and analysing electroretinogram data is the sometimes-poor signal-to-noise ratio that needs to be dealt with. This usually includes filtering as well as averaging of repeated measurements. Additionally, manual or threshold-based removal of trials that are considered to be outliers or particularly noisy can be performed before averaging. Manual rejection is labour intensive and prone to human bias. Also, threshold-based approaches can be problematic in cases where the noise on the trace has a dominating component with a specific polarity. This would be the case for breathing or heartbeat artefacts on mouse visual electrophysiology recordings. ERGtools2 therefore offers functions that identify outlier trials based on their similarity to the other trials. The function *autoreject.by.distance()*, for instance, calculates a distance matrix between the trials and rejects those that deviate from all others by more than a threshold value (default: one standard deviation). Applying this function does not require any prior information on the expected signal amplitude or other recording-specific parameters, and thus works without manual user input. Still, a function for threshold-based trial rejection is also available. The performance of *autoreject.by.distance()* for unbiased trial rejection is exemplified in **Figure 4 A**.

**Figure 4:**
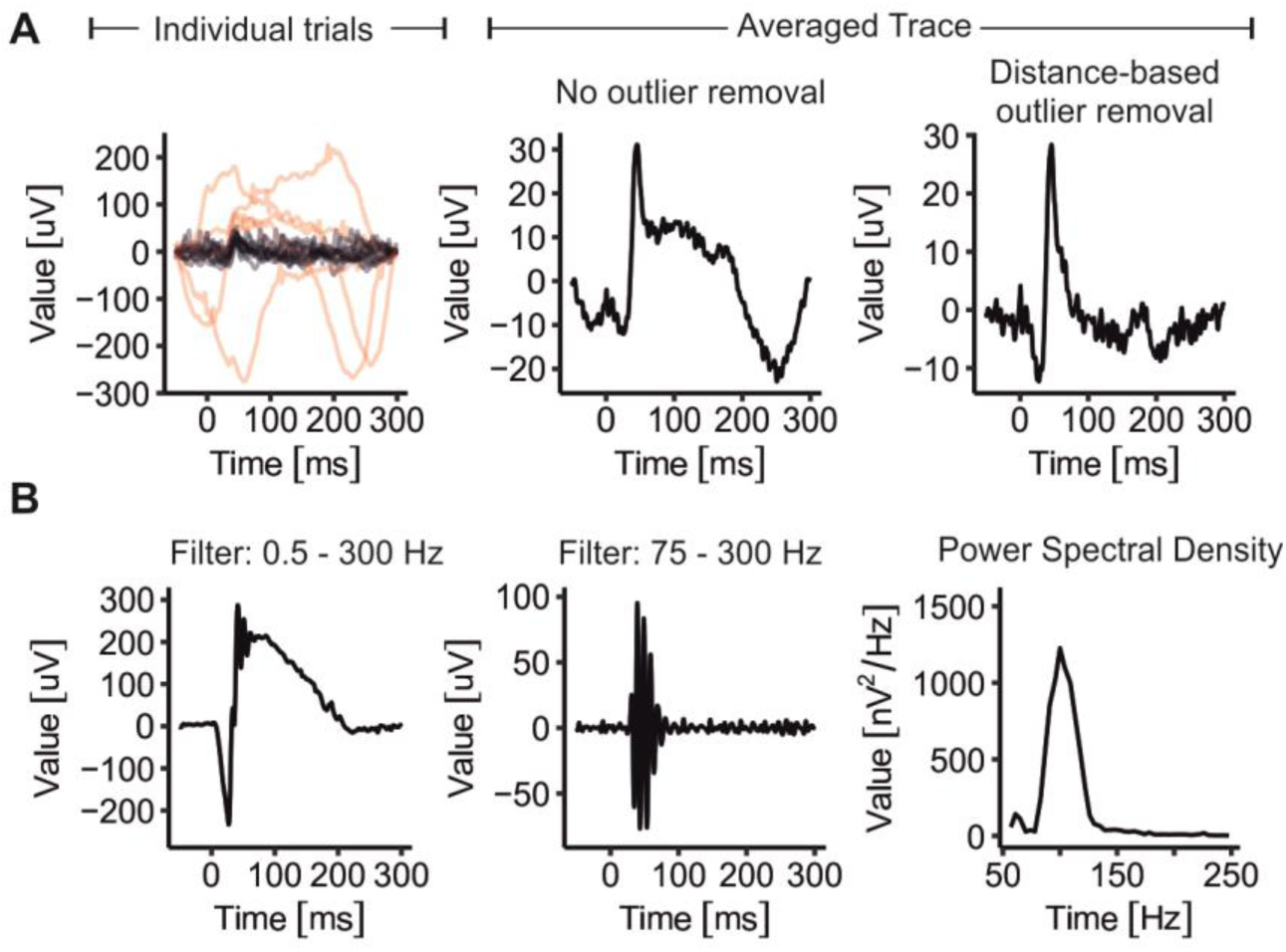
Special use cases: **(A)** Automatic rejection of outlier recordings using the *autoreject.by.distance()* function. Left panel: Raw trials of an ERG signal obtained in response to a light-adapted 1 cd*s/cm^2^ flash stimulus. Trials identified as outliers by *autoreject.by.distance()* are highlighted in red. Middle and right panel: Average traces from the raw trials shown on the left without (middle) and with (right) outlier trials removed. **(B)** Signal obtained in response to a dark-adapted 0.1 cd*s/cm^2^ flash stimulus. Left panel shows the recording after setting a 0.5-300 75-300

#### Extracting and analysing oscillatory potentials

Inner retinal network activity is reflected by the oscillatory potentials (OPs) which can be observed overlaying the B-wave in electroretinogram recordings [12]. Bandpass filter windows have been established to extract OPs from the remaining signal and these are commonly characterized describing their amplitude and frequency. Extraction of such information from ERG recordings would usually require a relatively high amount of manual coding. Using ERGtools2 the desired bandpass-filter window (usually 75 – 300 Hz) can be relatively easy set for an individual recording using *FilterFunction()<-* in conjunction with the *filter.bandpass()* function (**Figure 4 B**, left and middle panel). In analogy, data can be Fourier-transformed to extract peak frequency and spectral power using the *PSD* (= “power spectral density”) method (**Figure 4 B**, right panel).

#### Storing data for long-term preservation and shareability

To facilitate shareability and reusability of visual electrophysiology data ERGtools2 offers the option to store ERGExam objects in the Hierarchical Data Format version 5 [HDF5, 20] format using the *Save()* method. Its open, non-proprietary nature ensures long-term preservation and accessibility of the data – independent of any particular software – and its hierarchical structure enables grouping of data into logical self-documenting categories. The HDF5 file created by *Save()* mirrors the structure of the ERGExam object. Exams once saved into an HDF5 file can read into ERGtools2 again using *LoadERGExam()*.

## Discussion

ERGtools2 allows for efficient processing and analysis or visual electrophysiology data. It is written in the free open-source programming language R [13] and its code is itself free to use. To the best of our knowledge, ERGtools2 is thus the first comprehensive free and public domain toolbox for analysing visual electrophysiology data. ERGtools2 has an extensive command library allowing for complex and reproducible programming-based data processing but also offers an interactive interface which makes it possible to use the toolbox with minimal prior experience in programming. It supports storing data in the HDF5 format to support shareability, reusability and long-term preservation of visual electrophysiology data.

In the results section we have exemplified how ERGtools2 can be employed for routine/standard workup of ERG data as well as advanced data analysis like performing Fourier transformations for assessing the spectral power of an ERG signal and oscillatory potentials within it. All code used in this paper is available as supplemental material (Supplemental Material 3) and can serve as a starting point for own analyses.

The concept of ERGtools2 is to store unfiltered raw data alongside with filter functions, as well as rejection and averaging functions. Those filters are then applied when viewing a recording or extracting measurements. There might only be moderate advantage of this approach in a (clinical) routine setting where filter bandwidths for the different types of examinations are well established. In certain research settings, however, it can be particularly helpful: This could be the case when studying prosthetic visual restorative approaches like subretinal implants, optogenetic therapies or even stem cell-based approaches [8; 18; 2]. Herein electrical light responses may differ substantially from their physiological shapes and patterns, sometimes in not entirely predictable ways. If filtering is done right upon acquisition, potentially valuable components of the signal are irreversibly lost, while with raw data being stored and kept, changing and adjusting filters remains an option.

ERGtools2 is fully independent of the device and software used to acquire the data. Under the precondition that the system used for data acquisition supports raw data export, ERGtools2 can be used to process and analyse the data. Consequently, researchers can standardize their ERG data processing workflow, ensuring consistency and reliability regardless of the initial acquisition platform. Assuming stimulus standardization is not an issue, this facilitates collaboration and data sharing across different laboratories or clinical sites. In the context of laboratory research this is becoming increasingly relevant, not only in view of growing efforts to reduce the number of animals used in research, but also in view of justified claims for easier and practicable data sharing [21]. In the context of clinical research, it may facilitate data exchange between clinical study sites and reading centres, and, particularly by giving the reading centres access to the raw data, which might help them to better evaluate the quality of the submitted data.

In this regard it is also of importance that, beyond supporting convenient but still flexible analysis of visual electrophysiology data, ERGtools2 tools puts a special focus on data integrity. While the user can subset an exam to only keep its relevant parts for further analysis (e.g. only the data from one eye), filtering or averaging functions are only stored in the object but not applied to the data until visualization or measurement methods are called. This ensures that the raw data remains unaltered and readily available for re-analysis, while preventing any inadvertent loss of information or introduction of biases e.g. by accidental repeated application of filters to the data.

In line with the FAIR Data Principles [21], data once imported into ERGtools2 can be stored into HDF5 files [20], which has been implemented with a particular focus on long-term accessibility of data. The HDF5 file format is widely used and broadly accessible using open-source libraries. The structure in which ERGtools2 saves the data is self-documenting and resembles the structure of the ERGExam object. Thus, accessibility of the data is ensured also beyond the lifetime of any particular software, including ERGtools2 itself. Of note, there are ongoing efforts to implement data standards that can contain a broad variety of neuroscientific data with the aim to improve shareability and findability of the data (https://www.nwb.org/nwb-software/). This NWB format is also based on HDF5. Nevertheless, in the current release ERGtools2 uses is own file structure as described above. Main reason for this is that the data structure is substantially more simplistic (and therefore self-explanatory) than the versatile NBW format. Another recent effort puts a focus on visual electrophysiology data in particular: The ELVisML format is based on XML (eXtensible Markup Language), and capable of comprehensively storing ERG/VEP traces together with relevant metadata including stimulus and subject information [17]. The ELVisML format provides extensive capabilities for standardized storage of, e.g., the stimulus conditions, including calibration, as well as encrypted storage of patient-identifying data. The HDF5 format presented herein is extensible to also store these types of information. However, a key advantage of the binary HDF5 format lies in its ability to efficiently handle larger volumes of data, such as raw traces, rather than just averaged responses, which is essential for more detailed analyses.

Future releases of ERGtools2 should enable interaction with the ELVisML as well as the NBW format, while the current standard will always remain supported. Similarly, it is planned that future releases should provide more flexible versions of the auto-processing commands like *SetStandardFunctions()* and *AutoPlaceMarkers()* to ensure that these functions can be easily adopted for use with any species as well as any non-standard stimulus protocol.

### Conclusion

In conclusion, ERGtools2 is an open-source data analysis platform with a particular focus on data integrity, shareability and long-term preservation. It enables both, efficient routine data analysis, achievable without prior experience in programming as well as sophisticated customized data analysis and processing.

## MATERIAL AND METHODS

### Animal studies

All procedures were performed with the approval of the Giessen Regional Animal Health Authority (File No: G61/2021) and in accordance with the ARVO Statement for the Use of Animals in Ophthalmic and Vision Research. C57BL/6J were purchased from Charles River (Sulzfeld, Germany) and were kept under a 12-h light light/dark cycle, with no restriction on food and water. Electroretinogram recordings were performed at the age of 9-12 weeks usually at Zeitgeber time 3-6h under general anaesthesia. General anaesthesia was induced and maintained using isoflurane (Baxter, Deerfield, IL, USA). The maintenance dose was approximately 1% isoflurane in 0.2-0.5 L/min O_2_. Anaesthesia was administered using a Univentor 410-Q vaporizer (UNO Roestvaststaal BV, Zevenaar, The Netherlands). Pupil dilation was achieved by local application of Tropicamide (1%) and Phenylephrine (2.5%) eye drops. Electroretinograms were recorded using Celeris Rodent ERG system (Diagnosys LLC, Lowell, MA, USA) employing an integrated light guide stimulator and electrode. Oxybuprocaine hydrochloride (4 mg/ml) was installed to the eye before placement of the stimulator/electrodes.

### Electrophysiological Recordings

Electroretinograms were recorded using Espion software (Diagnosys) and digitized with a sampling rate of 2000 Hz. Mice were dark-adapted for at least 20 min before the beginning of the recording and thereafter only handled under dim red light. Before the beginning of the light adapted recordings, mice were exposed to 30 cd/m^2^ for at least 8 minutes. Flash stimulus duration was 4 ms maximum.

Recordings were exported from the Espion database into CSV files using the software’s inbuilt function. Export settings were set as follows: Export: Table of Content, Header Table, Marker Table, Stimulus Table and Data Table, Separator: Tab; Options: Titles, Vertical; Include: Steps, Channels, Results; Data Columns: Contents, Results, Sweeps.

### Structure and nomenclature of (visual) electrophysiological data

In electrophysiological tests, the fundamental unit of data is the time series, where data traces are recorded over a specified time period in response to a particular stimulus. These recordings are often repeated multiple times to enhance the signal-to-noise ratio through subsequent averaging. These repetitions are referred to as *trials,* while a complete set of trials would be considered as a *recording*. Recordings may be gathered from different electrodes, often simultaneously, and they are commonly referred to as *Channels*. In the context of visual electrophysiology there could be an ERG and a VEP channel acquired simultaneously, for instance. When dealing with paired organs, such as the eye, the stimulus or electrode may also have a laterality. In ERGtools2, each recording is therefore additionally defined by the *Eye* the signal corresponds to. Sequential recordings conducted under varying stimulus conditions, are referred to as *Steps*. Each step is characterized by the specifics of the stimulus, such as intensity, duration, and adaptation state of the eye, along with other parameters like the stimulus periodicity (e.g., flash or flicker).

### The ERGtools2 Package

The core of the ERGtools2 package is the ERGExam object, which stores all recordings from an individual examination. The structure of the ERGExam object is summarized in **Figure 2**. A single recording (including all its trials) is stored in an EPhysData object alongside with a time trace. The data slot of ERGExam represents a list of EPhysData objects thus containing all original data from an individual examination. This list of data is associated with the metadata table that contains all relevant contextual information, like the Eye that has been recorded from, the Channel, and the Step ID. Each step is further defined by the Stimulus Table, where a humane readable description of the stimulus may be stored along with the stimulus intensity, adaptation state or background light intensity and any other information defining the type of the stimulus. Additional information can be stored in both, the metadata and the stimulus table in user defined columns. To this end, the object contains the data together with all relevant information on the recording conditions.

ERG and VEP data can be analysed by measuring and comparing the amplitudes of the traces at defined positions, such as the a-wave representing the strong initial negative deflection of a recording. These measurements are stored in the Measurements slot. It contains the definition of the individual measurement point, termed *markers*, including their name, and whether amplitudes for that marker should be measured relative to any other marker or with reference to the baseline. It also stores the information on the recording an individual marker has been placed on and at what point on the time axis. The amplitudes at the position of the marker are not stored in the object but are computed each time the measurements are called to ensure data consistency.

The data in an ERGExam is normally stored as raw data and functions for filtering and averaging the data or rejecting poor quality/outlier trials can be stored within the object. Technically, these functions ae stored within each EPhysData object, but the ERGExam object ensures that these are consistent across the entire object. For instance, all recordings belonging to an individual channel and step must be filtered uniformly.

The EPhysData object additionally contains a slot termed “*StimulusTrace*”. This slot can be used (optionally) to store a vector representing the underlying stimulus trace, or any other trace that has the same timing as the actual data, e.g. the readings of a feedback-photodiode. ERGtools2 however, currently does not contain any options to visualize this trace alongside with the data.

A detailed description of the ERGExam object and all methods to access or modify this object can be found in the package’s help pages that can be accessed by typing *“?ERGtools2::‘ERGtools2-packageߣ”* into the R console after installation of the package. Key commands are additionally listed in **Table 1**. The ERGtools2 package expands on the functionality provided by the EPhysData (https://github.com/moritzlindner/EPhysData/) and EPhysMethods (https://github.com/moritzlindner/EPhysMethods/) packages.

**Table 1:** Important functions and methods implemented in the ERGtools2 package.

| Command | Description |
| --- | --- |
| `newERGExam()` | Create a new ERGExam object |
| `Save()`, `Load()` | Saves and loads the ERGExam object into and hdf5 file |
| `SetStandardFunctions()` | Sets standard functions for processing ERGExam data, defining default functions for averaging, filtering, and signal rejection based on the stimulus type. |
| `AutoPlaceMarkers()` | Automatically sets markers depending on the channel (e.g., ERG, VEP, OP) and stimulus type (Flash, Flicker). |
| `Where()` | Returns the indices of recordings matching the given criteria |
| `Subset()` | Subsets an ERGExam object into a new object of the same class |
| `Measurements()` | Returns the Measurements table |
| `interactiveMeasurements()` | Allows for interactive visual placement of markers. |
| `exploreERGExam()` | Explores the ERGExam object interactively. |
| `ggERGExam()` | Uses ggplot2 to plot the content of an ERGExam object |
| `ggIntensitySequence()` | Uses ggplot2 to plot intensity sequence for a list of ERG exams |

### Software requirements

ERGtools2 was developed in R, version 4.3.1, a free language for statistical computing and RStudio 2023.06.1 IDE on a Linux system (Ubuntu 22.04 LTS). The package was additionally tested Windows 10. ERGtools2 is available on GitHub (https://github.com/moritzlindner/ERGtools2) and submission to the Comprehensive R Archive Network (CRAN) is thought.

The package can be installed using the following lines of code:

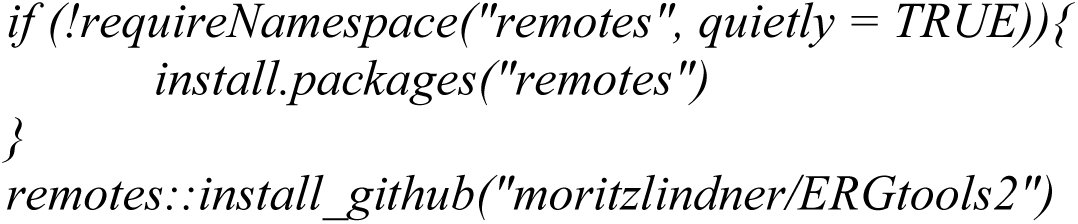

All package dependencies are available via CRAN except for the EPhysData (https://github.com/moritzlindner/EPhysData/) and EPhysMethods (https://github.com/moritzlindner/EPhysMethods/) packages, which can be obtained from GitHub.

## Declarations

### Author Contributions

*Participated in research design: ML*

*Conducted experiments: ML*

*Performed data analysis: ML*

*Wrote or contributed to the writing of the manuscript: ML*

### Data Availability

The datasets generated during and/or analysed during the current study are available from the corresponding author on reasonable request.

### Ethics approval

This study did not involve human participants. Animal work was performed with approval of the relevant authorities and in accordance with the institutional Ethics Guidelines of Animal Care. Further details are provided in the Methods section.

### Consent to publish

Not applicable.

