## Supplemental Material 1 for "The ERGtools2 package: A Toolset for Processing and Analysing Visual Electrophysiology Data": 404.html

Page not found (404) • ERGtools2
 


  

 


Toggle navigation


ERGtools2
0.7.0

- Reference

### Page not found (404)

Content not found. Please use links in the navbar.

#### Contents

Developed by Moritz Lindner.

Site built with pkgdown 2.0.7.
