## Supplemental Material 1 for "The ERGtools2 package: A Toolset for Processing and Analysing Visual Electrophysiology Data": authors.html

Authors and Citation • ERGtools2       

Toggle navigation


ERGtools2
0.7.0

- Reference

### Authors

- **Moritz Lindner**. Maintainer.

### Citation

person) (2024).
*ERGtools2: Importing and analysing Electroretinogram data in R*.
R package version 0.7.0.

```
@Manual{,
  title = {ERGtools2: Importing and analysing Electroretinogram data in R},
  author = {{person)}},
  year = {2024},
  note = {R package version 0.7.0},
}
```

Developed by Moritz Lindner.

Site built with pkgdown 2.0.7.
