## Supplemental Material 1 for "The ERGtools2 package: A Toolset for Processing and Analysing Visual Electrophysiology Data": index.html

Importing and analysing Electroretinogram data in R • ERGtools2
 


  


 


Toggle navigation


ERGtools2
0.7.0

- Reference

This package contains an environment for working with electroretinogram data. It contains an import method for Diagnosys Espion data, but allows reading in of data also from other manufacturers with limited coding effort. Standard procedures like averaging, subsetting an visualization of individual exams are supported.

### License

- GPL (>= 3)

### Citation

- Citing ERGtools2

### Developers

- Moritz Lindner   
   Maintainer

Developed by Moritz Lindner.

Site built with pkgdown 2.0.7.
