## Supplemental Material 1 for "The ERGtools2 package: A Toolset for Processing and Analysing Visual Electrophysiology Data": as.data.frame.html

as.data.frame for ERGProtocol, ERGStep, ERGChannel and ERGMarker — as.data.frame • ERGtools2       

Toggle navigation


ERGtools2
0.7.0

- Reference

### as.data.frame for ERGProtocol, ERGStep, ERGChannel and ERGMarker

`as.data.frame.Rd`

Converts an ERGExam, ERGMeasurements-class, ERGProtocol, ERGStep, ERGChannel or ERGMarker object to a data frame format.
Note that the `as.data.frame` method for ERGExam objects is inherited from the EPhysData-package package. See: EPhysData::as.data.frame-method.

```
# S4 method for ERGMeasurements
as.data.frame(x, row.names = NULL, optional = FALSE, ...)

# S4 method for ERGMarker
as.data.frame(x, row.names = NULL, optional = FALSE, ...)

# S4 method for ERGChannel
as.data.frame(x, row.names = NULL, optional = FALSE, ...)

# S4 method for ERGStep
as.data.frame(x, row.names = NULL, optional = FALSE, ...)

as.data.frame
```

#### Format

An object of class `character` of length 1.

#### Arguments

x
:   An ERGProtocol, ERGStep, ERGChannel or ERGMarker object.

...
:   currently unused.

#### Value

A data frame representing the ERGExam, ERGMeasurements, ERGProtocol, ERGStep, ERGChannel or ERGMarker object in long format.

#### Details

as.data.frame convert various ERG-related objects (ERGExam, ERGMeasurements, ERGProtocol, ERGMarker, ERGChannel, and ERGStep) to data frame formats, facilitating data manipulation and analysis. Each method extracts relevant information from the respective object and organizes it into a structured data frame.

- For ERGExam it is inherited from the EPhysData-package.
- For ERGMeasurements-class it is an alias for Measurements() (see: Measurements-Methods).

The ERGProtocol-related methods are experimental. These include:

- For ERGMarker objects, the resulting data frame contains two columns: Marker.Name and Marker.Relative.to, representing the marker's name and its relative position.
- For ERGChannel objects, the resulting data frame includes channel properties such as name, eye, frequency cutoffs, and inversion status, along with information about associated markers.
- For ERGStep objects, the resulting data frame includes step properties such as description, adaptation, and recording parameters, along with information about associated channels.
- For ERGProtocol objects, the resulting data frame includes protocol properties such as name and export date, along with information about associated steps.

These data frames provide comprehensive representations of the respective ERG-related objects, allowing for further analysis, visualization, or integration with other data.

#### Functions

- `as.data.frame(ERGMeasurements)`: Method for ERGMeasurements
- `as.data.frame(ERGMarker)`: Method for ERGMarker
- `as.data.frame(ERGChannel)`: Method for ERGChannel
- `as.data.frame(ERGStep)`: Method for ERGStep
- `as.data.frame`: Method for ERGProtocol

#### See also

ERGExam, ERGMeasurements-class, ERGProtocol, EPhysData-package, Measurements-Methods

#### Examples

```
data(ERG)
ERG <- SetStandardFunctions(ERG)
ERG <- Subset(ERG,where=list(Step=as.integer(1),Eye="RE")) # converting the whole object would return a huhge data.frame
head(as.data.frame(ERG))
#>   Step Channel Result Eye Channel_Name Recording Repeat       Time
#> 1    1     ERG      1  RE          ERG         1      1 -50.0 [ms]
#> 2    1     ERG      1  RE          ERG         1      1 -49.5 [ms]
#> 3    1     ERG      1  RE          ERG         1      1 -49.0 [ms]
#> 4    1     ERG      1  RE          ERG         1      1 -48.5 [ms]
#> 5    1     ERG      1  RE          ERG         1      1 -48.0 [ms]
#> 6    1     ERG      1  RE          ERG         1      1 -47.5 [ms]
#>            Value
#> 1 -9290.523 [nV]
#> 2 -7466.938 [nV]
#> 3 -6383.305 [nV]
#> 4 -5494.484 [nV]
#> 5 -4521.727 [nV]
#> 6 -3468.445 [nV]
```

#### Contents

Developed by Moritz Lindner.

Site built with pkgdown 2.0.7.
