## Supplemental Material 1 for "The ERGtools2 package: A Toolset for Processing and Analysing Visual Electrophysiology Data": as.std.channelname.html

Convert non-standard channel names to standard channel names — as.std.channelname • ERGtools2       

Toggle navigation


ERGtools2
0.7.0

- Reference

### Convert non-standard channel names to standard channel names

`as.std.channelname.Rd`

Convert non-standard channel names to standard channel names

```
as.std.channelname(channel_str, clear.unmatched = F)

is.std.channelname(channel_str)

erg_str()

op_str()

vep_str()
```

#### Arguments

channel\_str
:   Character vector of channel name strings to be converted.

clear.unmatched
:   Logical indicating whether to clear unmatched strings.

#### Value

A character vector with standardized channel names.

For is.std.channelname: Logical vector indicating whether each element in `channel_str` contains a word that describes a standard channel name.

#### Functions

- `is.std.channelname()`: Check if channel name strings are standard channel names
- `erg_str()`: Get standard ERG channel name strings
- `op_str()`: Get standard oscillatory potentials channel name strings
- `vep_str()`: Get standard VEP channel name strings

#### Examples

```
if (FALSE) {
as.std.channelname(c("ERG_auto", "C-wave", "OPs", "Nonstandard"), clear.unmatched = TRUE)
}
if (FALSE) {
is.std.channelname(c("ERG_auto", "C-wave", "OPs", "Nonstandard"))
}
if (FALSE) {
erg_str()
}
if (FALSE) {
op_str()
}

if (FALSE) {
vep_str()
}
```

#### Contents

Developed by Moritz Lindner.

Site built with pkgdown 2.0.7.
