## Supplemental Material 1 for "The ERGtools2 package: A Toolset for Processing and Analysing Visual Electrophysiology Data": as.std.eyename.html

Convert eye identifier strings to standard notation — as.std.eyename • ERGtools2       

Toggle navigation


ERGtools2
0.7.0

- Reference

### Convert eye identifier strings to standard notation

`as.std.eyename.Rd`

Convert eye identifier strings to standard notation

```
as.std.eyename(eye_str, exact = T, warn.only = F)

eye.haystack()

od_str()

os_str()
```

#### Arguments

eye\_str
:   Character vector of eye identifier strings.

exact
:   If `TRUE`: Require exact match. If `FALSE`: Whole word match is sufficient.

warn.only
:   Invalid eye identifier strings will only cause a warning, not an error.

#### Value

A character vector with standardized eye identifiers ('RE' for right eye, 'LE' for left eye).

#### Functions

- `eye.haystack()`: Get standard eye identifier strings
- `od_str()`: Get standard right eye identifier strings
- `os_str()`: Get standard left eye identifier strings

#### Examples

```
as.std.eyename(c("RE", "OD", "OS", "Right"))
#> [1] "RE" "RE" "LE" "RE"
if (FALSE) {
eye.haystack()
}#'
if (FALSE) {
od_str()
}
if (FALSE) {
os_str()
}
```

#### Contents

Developed by Moritz Lindner.

Site built with pkgdown 2.0.7.
