## Supplemental Material 1 for "The ERGtools2 package: A Toolset for Processing and Analysing Visual Electrophysiology Data": AutoPlaceMarkers.html

AutoPlaceMarkers for ERG/VEP Recordings — AutoPlaceMarkers • ERGtools2       

Toggle navigation


ERGtools2
0.7.0

- Reference

### AutoPlaceMarkers for ERG/VEP Recordings

`AutoPlaceMarkers.Rd`

These methods automatically place markers for ERG/VEP recordings in an ERGExam object.

```
AutoPlaceMarkers(
  X,
  Channel.names = pairlist(ERG = "ERG", VEP = "VEP"),
  Stimulus.type.names = pairlist(Flash = "Flash", Flicker = "Flicker")
)

AutoPlaceAB(
  X,
  robust.peak.filter.bands = c(5, 75),
  true.peak.tolerance = as_units(c(70, 30), "ms")
)

AutoPlaceFlicker(
  X,
  robust.peak.filter.bands = as_units(c(0.5, 300), "Hz"),
  true.peak.tolerance = as_units(c(12, 15), "ms")
)

AutoPlaceVEP(
  X,
  robust.peak.filter.bands = c(3, 75),
  true.peak.tolerance = as_units(c(80, 20), "ms")
)
```

#### Arguments

X
:   An ERGExam object for `AutoPlaceMarkers()` or an EPhysData::EPhysData-class object for the lower level methods AutoPlaceAB, AutoPlaceFlicker or AutoPlaceVEP.

Channel.names
:   A `pairlist` specifying channel names.

Stimulus.type.names
:   A base::pairlist specifying the names identifying the different stimulus types, e.g., `Flash="Flash"` or `Flash="Blitz"`.

robust.peak.filter.bands
:   A numeric vector of length 2 specifying the lower and upper bounds of the frequency band used for initial peak idetification n the lowe-level methods

true.peak.tolerance
:   A vector of class units and length 2 specifying the tolerance range around true peaks. Must be time values (i.e. a unit convertibel into 'seconds').

#### Value

An updated `ERGExam` object with markers placed.

#### Details

These methods are used to automatically place markers for ERGs/VEPs.  
  
`AutoPlaceMarkers()` sets markers depending on the channel (E.g. ERG, VEP, OP,...) and stimulus type (Flash, Flicker), defined via the `Channel.names` and `Stimulus.type.names` arguments. Markers are placed using the lower level methods AutoPlaceAB, AutoPlaceFlicker or AutoPlaceVEP function depending on the stimulus type.  
  
AutoPlaceAB, AutoPlaceFlicker and AutoPlaceVEP are the lower level functions which perform the actual marker placement on the EPhysData::EPhysData-class objects contained in the ERGExam object. These methods are usually not called directly by a user, unless she/he wants to perform or re-run marker placement only on certain recordings while leaving previously set markers unchanged for the others.
There working principle is that they apply robust peak filtering within defined frequency bands (low frequency band by default) to locate the gross position of the most prominent peaks peaks and then look for the peak in data using the preset filter function (FilterFunction) to accurately identify the actual peak position.   
  
Currently, supported are:

- a and B waves for Flash ERG
- N1, P1 (and Frequency) for Flicker ERGs
- P1, N1, and P2 for Flash ERGs

#### Functions

- `AutoPlaceMarkers()`: Automatically sets markers depending on the channel (E.g. ERG, VEP, OP,...) and stimulus type (Flash, FLicker).
- `AutoPlaceAB()`: places the a and B waves on Flash ERG data stored in an an EPhysData::EPhysData-class object.
- `AutoPlaceFlicker()`: places the N1 and P1 markers and determines 1/frequency (period) for Flicker ERG data stored in an an EPhysData::EPhysData-class object
- `AutoPlaceVEP()`: places the P1, N1 and P2 markers for Flash VEP data stored in an an EPhysData::EPhysData-class object

#### Examples

```
data(ERG)
ERG<-SetStandardFunctions(ERG)
imported_Markers<-Measurements(ERG)
#> Retrieving record values for the given time points.
#> ================================================================================
head(imported_Markers)
#>   Recording Step     Description Channel Result Eye Name Relative       Time
#> 1         1    1 DA 0 01 cd s m      ERG      1  RE    a     <NA> 0.0315 [s]
#> 2         1    1 DA 0 01 cd s m      ERG      1  RE    B        a 0.0515 [s]
#> 3         2    1 DA 0 01 cd s m      ERG      1  LE    a     <NA> 0.0315 [s]
#> 4         2    1 DA 0 01 cd s m      ERG      1  LE    B        a 0.0505 [s]
#> 5         5    2    DA 1 cd s m      ERG      1  RE    a     <NA> 0.0130 [s]
#> 6         5    2    DA 1 cd s m      ERG      1  RE    B        a 0.0355 [s]
#>           Voltage Channel_Name Recording.y
#> 1  -53.17071 [uV]          ERG           1
#> 2  346.34079 [uV]          ERG           1
#> 3  -79.03807 [uV]          ERG           2
#> 4  303.53579 [uV]          ERG           2
#> 5 -202.88360 [uV]          ERG           5
#> 6  544.37543 [uV]          ERG           5
ERG<-ClearMeasurements(ERG)
imported_Markers_cleared<-Measurements(ERG)
head(imported_Markers_cleared)
#>  [1] Recording   Step        Description Channel     Result      Eye        
#>  [7] Name        Relative    Time        Voltage    
#> <0 rows> (or 0-length row.names)
ERG<-AutoPlaceMarkers(ERG, Channel.names = pairlist(ERG = "ERG"))
#> ================================================================================
autoplaced_Markers<-Measurements(ERG)
#> Retrieving record values for the given time points.
#> ================================================================================
head(autoplaced_Markers)
#>   Recording Step     Description Channel Result Eye Name Relative       Time
#> 1         1    1 DA 0 01 cd s m      ERG      1  RE    a     <NA> 0.0315 [s]
#> 2         1    1 DA 0 01 cd s m      ERG      1  RE    B        a 0.0515 [s]
#> 3         2    1 DA 0 01 cd s m      ERG      1  LE    a     <NA> 0.0310 [s]
#> 4         2    1 DA 0 01 cd s m      ERG      1  LE    B        a 0.0505 [s]
#> 5         5    2    DA 1 cd s m      ERG      1  RE    a     <NA> 0.0125 [s]
#> 6         5    2    DA 1 cd s m      ERG      1  RE    B        a 0.0355 [s]
#>           Voltage Channel_Name Recording.y
#> 1  -53.17071 [uV]          ERG           1
#> 2  346.34079 [uV]          ERG           1
#> 3  -85.88321 [uV]          ERG           2
#> 4  310.38093 [uV]          ERG           2
#> 5 -206.37706 [uV]          ERG           5
#> 6  547.86889 [uV]          ERG           5

# Calling AutoPlaceAB() directly
X<-ERG@Data[[1]] # get first recording
AutoPlaceAB(X)
#>        Time          Value Relative
#> a 31.5 [ms] -53170.71 [nV]     <NA>
#> B 51.5 [ms] 346340.79 [nV]        a
```

#### Contents

Developed by Moritz Lindner.

Site built with pkgdown 2.0.7.
