## Supplemental Material 1 for "The ERGtools2 package: A Toolset for Processing and Analysing Visual Electrophysiology Data": CheckAvgFxSet.html

Check if all recordings in an ERGExam object have a valid averaging function set. — CheckAvgFxSet • ERGtools2       

Toggle navigation


ERGtools2
0.7.0

- Reference

### Check if all recordings in an ERGExam object have a valid averaging function set.

`CheckAvgFxSet.Rd`

Check if all recordings in an ERGExam object have a valid averaging function set.

```
CheckAvgFxSet(X)
```

#### Arguments

X
:   An ERGExam object.

#### Value

Logical, TRUE if all recordings have a valid averaging function, FALSE otherwise.

#### See also

`AverageFunction<-`, `SetStandardFunctions`

#### Examples

```
data(ERG)
CheckAvgFxSet(ERG)
#> [1] TRUE
ERG<-SetStandardFunctions(ERG)
CheckAvgFxSet(ERG)
#> [1] TRUE
```

#### Contents

Developed by Moritz Lindner.

Site built with pkgdown 2.0.7.
