## Supplemental Material 1 for "The ERGtools2 package: A Toolset for Processing and Analysing Visual Electrophysiology Data": CollectMeasurements.html

Get measurements for plotting — CollectMeasurements • ERGtools2       

Toggle navigation


ERGtools2
0.7.0

- Reference

### Get measurements for plotting

`CollectMeasurements.Rd`

This function extracts measurements and related information, e.g. for plotting or statistics.

```
CollectMeasurements(
  List,
  where = list(),
  Markers = c("a", "B", "N1", "P1"),
  measure.absolute = F
)
```

#### Arguments

List
:   A list of ERG exams.

where
:   A base::list defining selection criteria. Tags/Keys in the names in the list must represent valid column names of Metadata orStimulusTable.

Markers
:   Vector of markers to include in the plot.

measure.absolute
:   Logical, default: FALSE. If absolute amplitudes should be returned instead of amplitudes relative to the reference marker (where given).

#### Value

A data frame with measurements for plotting.#'

#### Examples

```
data(ERG)
ERG<-SetStandardFunctions(ERG)
ERG <- AutoPlaceMarkers(ERG)
#> ================================================================================
CollectMeasurements(list(ERG,ERG), list(Background = "DA", Type = "Flash"))
#> ================================================================================
#>    Step     Description Channel Result Eye Name Relative       Time
#> 1     1 DA 0 01 cd s m      ERG      1  RE    a     <NA> 0.0315 [s]
#> 2     1 DA 0 01 cd s m      ERG      1  RE    B        a 0.0515 [s]
#> 3     1 DA 0 01 cd s m      ERG      1  LE    a     <NA> 0.0310 [s]
#> 4     1 DA 0 01 cd s m      ERG      1  LE    B        a 0.0505 [s]
#> 5     2    DA 1 cd s m      ERG      1  RE    a     <NA> 0.0125 [s]
#> 6     2    DA 1 cd s m      ERG      1  RE    B        a 0.0355 [s]
#> 7     2    DA 1 cd s m      ERG      1  LE    a     <NA> 0.0130 [s]
#> 8     2    DA 1 cd s m      ERG      1  LE    B        a 0.0355 [s]
#> 9     3    DA 3 cd s m      ERG      1  RE    a     <NA> 0.0090 [s]
#> 10    3    DA 3 cd s m      ERG      1  RE    B        a 0.0345 [s]
#> 11    3    DA 3 cd s m      ERG      1  LE    a     <NA> 0.0095 [s]
#> 12    3    DA 3 cd s m      ERG      1  LE    B        a 0.0345 [s]
#> 13    1 DA 0 01 cd s m      ERG      1  RE    a     <NA> 0.0315 [s]
#> 14    1 DA 0 01 cd s m      ERG      1  RE    B        a 0.0515 [s]
#> 15    1 DA 0 01 cd s m      ERG      1  LE    a     <NA> 0.0310 [s]
#> 16    1 DA 0 01 cd s m      ERG      1  LE    B        a 0.0505 [s]
#> 17    2    DA 1 cd s m      ERG      1  RE    a     <NA> 0.0125 [s]
#> 18    2    DA 1 cd s m      ERG      1  RE    B        a 0.0355 [s]
#> 19    2    DA 1 cd s m      ERG      1  LE    a     <NA> 0.0130 [s]
#> 20    2    DA 1 cd s m      ERG      1  LE    B        a 0.0355 [s]
#> 21    3    DA 3 cd s m      ERG      1  RE    a     <NA> 0.0090 [s]
#> 22    3    DA 3 cd s m      ERG      1  RE    B        a 0.0345 [s]
#> 23    3    DA 3 cd s m      ERG      1  LE    a     <NA> 0.0095 [s]
#> 24    3    DA 3 cd s m      ERG      1  LE    B        a 0.0345 [s]
#>            Voltage Recording.y Subject   Group            ExamDate Intensity
#> 1   -53.17071 [uV]           1  CR2170 DEFAULT 2023-08-10 11:18:31      0.01
#> 2   346.34079 [uV]           1  CR2170 DEFAULT 2023-08-10 11:18:31      0.01
#> 3   -81.67991 [uV]           2  CR2170 DEFAULT 2023-08-10 11:18:31      0.01
#> 4   314.19535 [uV]           2  CR2170 DEFAULT 2023-08-10 11:18:31      0.01
#> 5  -209.53673 [uV]           5  CR2170 DEFAULT 2023-08-10 11:18:31      1.00
#> 6   546.54504 [uV]           5  CR2170 DEFAULT 2023-08-10 11:18:31      1.00
#> 7  -210.87246 [uV]           6  CR2170 DEFAULT 2023-08-10 11:18:31      1.00
#> 8   498.93258 [uV]           6  CR2170 DEFAULT 2023-08-10 11:18:31      1.00
#> 9  -233.89249 [uV]           9  CR2170 DEFAULT 2023-08-10 11:18:31      3.00
#> 10  587.60032 [uV]           9  CR2170 DEFAULT 2023-08-10 11:18:31      3.00
#> 11 -249.85830 [uV]          10  CR2170 DEFAULT 2023-08-10 11:18:31      3.00
#> 12  564.03033 [uV]          10  CR2170 DEFAULT 2023-08-10 11:18:31      3.00
#> 13  -53.17071 [uV]           1  CR2170 DEFAULT 2023-08-10 11:18:31      0.01
#> 14  346.34079 [uV]           1  CR2170 DEFAULT 2023-08-10 11:18:31      0.01
#> 15  -81.67991 [uV]           2  CR2170 DEFAULT 2023-08-10 11:18:31      0.01
#> 16  314.19535 [uV]           2  CR2170 DEFAULT 2023-08-10 11:18:31      0.01
#> 17 -209.53673 [uV]           5  CR2170 DEFAULT 2023-08-10 11:18:31      1.00
#> 18  546.54504 [uV]           5  CR2170 DEFAULT 2023-08-10 11:18:31      1.00
#> 19 -210.87246 [uV]           6  CR2170 DEFAULT 2023-08-10 11:18:31      1.00
#> 20  498.93258 [uV]           6  CR2170 DEFAULT 2023-08-10 11:18:31      1.00
#> 21 -233.89249 [uV]           9  CR2170 DEFAULT 2023-08-10 11:18:31      3.00
#> 22  587.60032 [uV]           9  CR2170 DEFAULT 2023-08-10 11:18:31      3.00
#> 23 -249.85830 [uV]          10  CR2170 DEFAULT 2023-08-10 11:18:31      3.00
#> 24  564.03033 [uV]          10  CR2170 DEFAULT 2023-08-10 11:18:31      3.00
#>    Background  Type
#> 1          DA Flash
#> 2          DA Flash
#> 3          DA Flash
#> 4          DA Flash
#> 5          DA Flash
#> 6          DA Flash
#> 7          DA Flash
#> 8          DA Flash
#> 9          DA Flash
#> 10         DA Flash
#> 11         DA Flash
#> 12         DA Flash
#> 13         DA Flash
#> 14         DA Flash
#> 15         DA Flash
#> 16         DA Flash
#> 17         DA Flash
#> 18         DA Flash
#> 19         DA Flash
#> 20         DA Flash
#> 21         DA Flash
#> 22         DA Flash
#> 23         DA Flash
#> 24         DA Flash
```

#### Contents

Developed by Moritz Lindner.

Site built with pkgdown 2.0.7.
