## Supplemental Material 1 for "The ERGtools2 package: A Toolset for Processing and Analysing Visual Electrophysiology Data": dot-SampleERGExam.html

Exemplary ERG Exam — .SampleERGExam • ERGtools2       

Toggle navigation


ERGtools2
0.7.0

- Reference

### Exemplary ERG Exam

`dot-SampleERGExam.Rd`

This data set contains an ERG Exam with ERG and OP channels from DA and LA flash and flicker stimuli..

```
data(ERG)
```

#### Format

An object of class `"ERGExam"`; see ERGExam.

#### Examples

```
data(ERG)
ERG<-SetStandardFunctions(ERG)
ERG
#> An object of class ERGExam
#> Subject:	CR2170, 2023-05-30, Male
#> Exam Date:	 2023-08-10 11:18:31
#> Protocol:	 01-1 MoL_v2_DA long ERG [14806-F || ECN 1685 || 13 July 2021]
#> Steps:		DA 0 01 cd s m 
#> 		DA 1 cd s m 
#> 		DA 3 cd s m 
#> Eyes:		RE	LE
#> Channels:	ERG
#> 		OP
#> 
#> An averge function has been set for this object.
#> Size: 2.3 Mb 
ggERGExam(ERG)
#> Retrieving record values for the given time points.
#> ================================================================================
```

#### Contents

Developed by Moritz Lindner.

Site built with pkgdown 2.0.7.
