## Supplemental Material 1 for "The ERGtools2 package: A Toolset for Processing and Analysing Visual Electrophysiology Data": dot-SampleERGMeasurements.html

Exemplary ERG Measurements object — .SampleERGMeasurements • ERGtools2       

Toggle navigation


ERGtools2
0.7.0

- Reference

### Exemplary ERG Measurements object

`dot-SampleERGMeasurements.Rd`

This data set contains a Measurements object with Markers and corresponding measurements for an hypothetical ERG+VEP exam..

```
data(Measurements.data)
```

#### Format

An object of class `"ERGMeasurements"`; see ERGMeasurements.

#### Examples

```
data(Measurements.data)
Measurements.data
#> ERGMeasurements object:
#> Measurements:
#>   Recording Name ChannelBinding Relative    Time
#> 1         1    a            ERG     <NA> 10 [ms]
#> 2         1    B            ERG        a 15 [ms]
#> 3         2   N1            VEP     <NA> 20 [ms]
#> 4         2   P1            VEP       N1 25 [ms]
#> 5         3    a            ERG     <NA> 30 [ms]
#> 6         3    B            ERG        a 35 [ms]
Markers(Measurements.data)
#>   Name Relative ChannelBinding
#> 1    a     <NA>            ERG
#> 2    B        a            ERG
#> 3   N1     <NA>            VEP
#> 4   P1       N1            VEP
```

#### Contents

Developed by Moritz Lindner.

Site built with pkgdown 2.0.7.
