## Supplemental Material 1 for "The ERGtools2 package: A Toolset for Processing and Analysing Visual Electrophysiology Data": DropRecordings-method.html

Drop specified recordings from an ERGExam Object — DropRecordings-method • ERGtools2       

Toggle navigation


ERGtools2
0.7.0

- Reference

### Drop specified recordings from an ERGExam Object

`DropRecordings-method.Rd`

This method returns a new `ERGExam` object with specific recordings removed.

```
DropRecordings(X, where)
```

#### Arguments

X
:   An ERGExam

where
:   A base::list defining selection criteria to identify recordings to remove.

#### Details

The `DropRecordings` function creates a new `ERGExam` object excluding the recordings
identified by the `where` criteria.

#### See also

EPhysData::Subset

#### Examples

```
data(ERG)
tmp<-DropRecordings(ERG, where=list(Channel="ERG", Intensity=1))
```

#### Contents

Developed by Moritz Lindner.

Site built with pkgdown 2.0.7.
