## Supplemental Material 1 for "The ERGtools2 package: A Toolset for Processing and Analysing Visual Electrophysiology Data": ERGExam.html

ERGExam Class — ERGExam • ERGtools2       

Toggle navigation


ERGtools2
0.7.0

- Reference

### ERGExam Class

`ERGExam.Rd`

A class representing an ERG (Electroretinogram) exam. This class extends the EPhysData::EPhysSet object and all methods valid for EPhysData::EPhysSet can also be applied to ERGExam objects.

#### Slots

`Data`
:   Data A list of EPhysData::EPhysData objects. Each item containing a recording in response to a particular Stimulus ("Step"), from a particular eye and data Channel (e.g. ERG or OP), as defined in the corresponding Metadata.

`Metadata`
:   A data frame containing metadata information associated with the data in the Data slot, each row corresponds to one item in `data`.

    Step
    :   An integer vector containing the step index. A step describes data recorded in response to the same type of stimulus. This column links the Stimulus slot to the metadata

    Eye
    :   A character vector. Possible values "RE" (right eye) and "LE" (left eye).

    Channel
    :   A character vector containing the channel name. This can be "ERG", "VEP" or "OP" for instance.

    Result
    :   A numeric vector containing the indices of individual results contained in an ERG exam. E.g., if a recording to one identical stimulus is performed twice, these would be distinguished by different indices in the Result column. Warning: This is currently experimental

`Stimulus`
:   A data frame containing stimulus information.

    Step
    :   An integer vector row index. Will be removed in future versions.

    Description
    :   A character vector describing the stimulus in a human-readable way.

    Intensity
    :   A numeric vector representing the intensity of the stimulus.

    Background
    :   A character vector describing the adaptation state of the retina for that stimulus (DA or LA).

    Type
    :   A character vector describing the type of the stimulus (e.g. Flash or Flicker).

`Averaged`
:   TRUE if the object contains averaged data, FALES indicates object contains raw traces.

`Measurements`
:   An object of class ERGMeasurements

`ExamInfo`
:   A list containing exam-related information.

    ProtocolName
    :   A character vector indicating the name of the protocol.

    Version
    :   Optional: A character vector indicating the version of the protocol.

    ExamDate
    :   A `POSIXct` The date of the exam.

    Filename
    :   Optional: The filename associated where exam raw data have been imported from.

    RecMode
    :   Optional: A character vector indicating the recording mode.

    Investigator
    :   Optional: A character vector indicating the name of the investigator conducting the exam.

`SubjectInfo`
:   A list containing subject-related information.

    Subject
    :   A character vector indicating the name of the subject

    DOB
    :   A `Date` object indicating the date of birth of the subject

    Gender
    :   Optional: A character vector indicating the gender of the subject

    Group
    :   Optional: A character vector indicating the study group to which the subject belongs.

`Imported`
:   A `POSIXct` timestamp indicating when the object was imported.

#### See also

EPhysData::EPhysData-package EPhysData::EPhysData-class EPhysData::EPhysSet-class

#### Contents

Developed by Moritz Lindner.

Site built with pkgdown 2.0.7.
