## Supplemental Material 1 for "The ERGtools2 package: A Toolset for Processing and Analysing Visual Electrophysiology Data": ERGMeasurements-class.html

ERGMeasurements Class — ERGMeasurements-class • ERGtools2       

Toggle navigation


ERGtools2
0.7.0

- Reference

### ERGMeasurements Class

`ERGMeasurements-class.Rd`

This class represents ERG (Electroretinogram) measurements, which typically consist
of markers placed at specific points in a recording and measurements associated with
each marker. Note: only the time points for the measurement is stored in the object, not the actual values. If called from a parent ERGExam, values are retrieved based on the raw data stored in that object.

#### Slots

`Marker`
:   A data.frame containing marker information. This data.frame should
    have columns 'Name' and 'Relative'. The 'Name' column represents
    the name of each marker and 'Relative' column indicates the relative position of
    each marker with respect to other markers (or NA if no relative marker).

`Measurements`
:   A data.frame containing measurement information. This data.frame
    should have columns 'Recording', 'Marker', and 'Time'. The 'Recording' column
    represents the recording number associated with each measurement, 'Marker' column
    indicates the marker associated with each measurement (using the row index of the
    marker in the Marker data.frame), and 'Time' column represents the time of each
    measurement.

    To create a valid ERGMeasurements object, ensure the following:

    - The Marker slot is a data.frame with columns 'Name' and 'Relative'.
    - The 'Name' column in the Marker data.frame contains names for each marker, which are unique within an individual channel.
    - The 'Relative' column in the Marker data.frame contains valid indices of other markers from the same channel
      or NA if no relative marker.
    - The Measurements slot is a data.frame with columns 'Recording', 'Marker', and 'Time'.
    - The 'Marker' column in the Measurements data.frame contains valid row indices of the
      Marker data.frame.
    - If the 'Marker' column points to a relative marker, there must be already a row containing a measurement from the same recording for the parent marker.
    - The 'Time' column in the Measurements data.frame contains valid time units.

#### See also

EPhysData::EPhysSet Measurements-Methods Get

#### Examples

```
# Create marker data frame
marker_df <- data.frame(
  Name = c("a", "B", "N1", "P1"),
  Relative = c(NA, 1, NA, 3),
  ChannelBinding =c("ERG","ERG","VEP","VEP")
)

# Create measurements data frame
measurements_df <- data.frame(
  Recording = c(1, 1, 2, 2, 3, 3),
  Marker = c(1, 2, 3, 4, 1, 2),
  Time = as_units(c(10, 15, 20, 25, 30, 35), "ms")
)

# Create ERGMeasurements object
erg_obj <- new("ERGMeasurements", Marker = marker_df, Measurements = measurements_df)

# Show the object
erg_obj
#> ERGMeasurements object:
#> Measurements:
#>   Recording Name ChannelBinding Relative    Time
#> 1         1    a            ERG     <NA> 10 [ms]
#> 2         1    B            ERG        a 15 [ms]
#> 3         2   N1            VEP     <NA> 20 [ms]
#> 4         2   P1            VEP       N1 25 [ms]
#> 5         3    a            ERG     <NA> 30 [ms]
#> 6         3    B            ERG        a 35 [ms]

# Check validity of the object
validObject(erg_obj)
#> [1] TRUE
```

#### Contents

Developed by Moritz Lindner.

Site built with pkgdown 2.0.7.
