## Supplemental Material 1 for "The ERGtools2 package: A Toolset for Processing and Analysing Visual Electrophysiology Data": ERGProtocol.html

ERGProtocol class definition — ERGProtocol • ERGtools2       

Toggle navigation


ERGtools2
0.7.0

- Reference

### ERGProtocol class definition

`ERGProtocol.Rd`

This class represents an ERG protocol, as it can be impored from Diagnosys Espion™ software with Export\_Date, Name, nSteps, nChannels, and Step slots.

#### Slots

`Export_Date`
:   POSIXct slot for the export date and time.

`Name`
:   Character slot for the protocol name.

`nSteps`
:   Numeric slot for the number of steps.

`nChannels`
:   Numeric slot for the number of channels.

`Step`
:   List slot for a list of 'ERGStep' objects.

#### Contents

Developed by Moritz Lindner.

Site built with pkgdown 2.0.7.
