## Supplemental Material 1 for "The ERGtools2 package: A Toolset for Processing and Analysing Visual Electrophysiology Data": ERGtools2-package.html

Structured Index and Introduction for the ERGtools2 R Package — ERGtools2-package • ERGtools2       

Toggle navigation


ERGtools2
0.7.0

- Reference

### Structured Index and Introduction for the ERGtools2 R Package

`ERGtools2-package.Rd`

This package contains an environment for working with electroretinogram data. It contains an import method for Diagnosys Espion data, but allows reading in of data coming in other formats with limited coding effort. Standard procedures like averaging, sub-setting an visualization of individual exams are supported.
This package provides the ERGExam class that stores data from a single ERG examination. These may include different recording channels (ERG, OP, VEP, ...), or sequential recordings in response to different stimulus paradigms.
It is usually generated from imported raw data using newERGExam. Data acuired from Diagnosys Espion can be imported directly using ImportEspion.

#### Details

The class ERGExam expands EPhysData::EPhysSet, so **Methods that work on EPhysData::EPhysSet can be applied to ERGExam as well. See: EPhysData::EPhysData-package**.

#### Standard workflow

After setting the functions, the accession methods inherited from EPhysData::EPhysSet-class like EPhysData::as.data.frame-method and EPhysData::GetData with the argument `Raw=F` can be used to return the processed data.

#### Object creation

|  |  |  |
| --- | --- | --- |
| **Function/Method** | **Classes** | **Description** |
| **newERGExam** | ERGExam | Create a new ERGExam object. |
| ImportEspion, ImportEspionInfo, ImportEspionStimTab, and ImportEspionMetadata | ERGExam | Import an ERG exam from a CSV file exported from the Diagnosys Espion software |
| methods:new and ImportEspionProtocol | ERGProtocol | Experimental. Creates a container for ERG recording protocols as exported from exported the Diagnosys Espion software |
| methods:new | ERGMeasurements | Creates a container for ERG Markers and their positions on an ERG trace. Usually not necessary to be called by the user directly. |
| Save, Load.ERGExam | ERGExam | Saves and loads the ERGExam object. |

#### Accession methods

|  |  |  |
| --- | --- | --- |
| **Function/Method** | **Classes** | **Description** |
| **Where** | ERGExam | Returns the recording index for those recordings matching the given criteria. |
| **Subset** | ERGExam | Subsets an ERGExam object into a new object of the same class. |
| **as.data.frame** | ERGExam , ERGProtocol | Returns data frame representing the ERGExam or ERGProtocol object in long format. |
| Stimulus | ERGExam | Returns selected rows of a stimulus table. |
| StimulusDescription, StimulusIntensity, StimulusBackground, StimulusType | ERGExam | Returns details of stimulus description, intensity, background, and type. |
| MarkerNames | ERGExam , ERGMeasurements | Returns the names of markers in the dataset. |
| Markers | ERGExam , ERGMeasurements | Returns marker information from the dataset. |
| **Measurements** | ERGExam , ERGMeasurements | Returns the Measurements table. |
| DOB, ExamDate, GroupName, ProtocolName | ERGExam | Returns information on the exam and the examined subject. |
| Eyes, Steps, Channels, Results | ERGExam | Returns information on the Recordings contained in the dataset. |

#### Processing

|  |  |  |
| --- | --- | --- |
| **Function/Method** | **Classes** | **Description** |
| **SetStandardFunctions** | ERGExam | Sets standard functions for processing ERGExam data, defining default functions for averaging, filtering, and signal rejection based on the stimulus type. |
| FilterFunction<-, Rejected<-, AverageFunction<- | ERGExam | Updates the FilterFunction, Rejected function or AverageFunction for all Recordings in an ERGExam, or only those selected using `where`. |
| AutoPlaceVEP | ERGExam | Automatically sets markers depending on the channel (e.g., ERG, VEP, OP) and stimulus type (Flash, Flicker). |
| AutoPlaceAB, AutoPlaceFlicker, AutoPlaceVEP | EPhysData::EPhysData-class | Automatically sets markers for the respective type of channel (e.g., ERG, VEP, OP) and stimulus (Flash, Flicker). |
| CheckAvgFxSet | ERGExam | Checks if the average function is set correctly for the dataset. |
| **interactiveMeasurements** | ERGExam | Allows for interactive visual placement of markers. |

#### Merging and other object manipulation

|  |  |  |
| --- | --- | --- |
| **Function/Method** | **Classes** | **Description** |
| MergeERGExams | ERGExam | Merges multiple ERGExam objects into one. |
| Measurements<- | ERGExam , ERGMeasurements-class | Adds, updates, or removes Measurements from an ERGExam or ERGMeasurements object. |
| ClearMeasurements | ERGExam | Clears the Measurements slots in an ERGExam object. |
| DropMarker, AddMarker, RenameMarker | ERGExam | Methods for marker modification. |
| StimulusDescription()<- , StimulusIntensity()<-, StimulusBackground()<-, StimulusType()<-, | ERGExam | Updates stimulus description, intensity, background, and type in the stimulus table. |
| UpdateChannelNames | ERGExam | Updates or replaces channel names in the dataset. |
| DropRecordings | ERGExam | Drops specified recordings from the ERGExam object. |
| as.std.channelname, is.std.channelname, erg\_str, op\_str, vep\_str | Function | Standardizes channel names, checks if the channel name is standardized, returns standardized strings for ERG, OP, and VEP. |
| as.std.eyename, eye.haystack, od\_str, os\_str | Function | Standardizes eye names, processes eye data, returns standardized strings for the right eye (OD) and left eye (OS). |

#### Plot methods and interactive methods

|  |  |  |
| --- | --- | --- |
| **Function/Method** | **Classes** | **Description** |
| ggERGTrace | ERGExam | Generates a ggplot plot for a single trace from an ERGExam object. |
| **ggERGExam** | ERGExam | Plots a complete ERGExam object. |
| ggIntensitySequence | ERGExam | Uses ggplot2 to plot intensity sequence for ERG exams. |
| ggStepSequence | ERGExam | Uses ggplot2 to plot step sequence (i.e., sequential recordings within a single protocol) for ERG exams. |
| ggPlotRecordings | ERGExam | Uses ggplot2 to plot ERG traces from multiple ERGExam objects. |
| **interactiveMeasurements** | ERGExam | Interactive visual placement of markers using a Shiny app. |
| **exploreERGExam** | ERGExam | Explores the ERGExam object interactively. |

#### Author

Moritz Lindner

#### Examples

```
# a typical workflow
data(ERG) # load example data, to import own data, see the examples provided for newERGExam and ImportEspion
StimulusTable(ERG) # have a look whats inside
#>   Step     Description Intensity Background  Type
#> 1    1 DA 0 01 cd s m       0.01         DA Flash
#> 2    2    DA 1 cd s m       1.00         DA Flash
#> 3    3    DA 3 cd s m       3.00         DA Flash
Metadata(ERG)
#>    Step Channel Result Eye Channel_Name Recording
#> 1     1     ERG      1  RE          ERG         1
#> 2     1     ERG      1  LE          ERG         2
#> 3     1      OP      1  RE           OP         3
#> 4     1      OP      1  LE           OP         4
#> 5     2     ERG      1  RE          ERG         5
#> 6     2     ERG      1  LE          ERG         6
#> 7     2      OP      1  RE           OP         7
#> 8     2      OP      1  LE           OP         8
#> 9     3     ERG      1  RE          ERG         9
#> 10    3     ERG      1  LE          ERG        10
#> 11    3      OP      1  RE           OP        11
#> 12    3      OP      1  LE           OP        12
ERG<-SetStandardFunctions(ERG)
ggERGTrace(ERG, where = list( Step = as.integer(3), Eye = "RE", Channel ="ERG", Result = as.integer(1))) # pick one and have a look at the traces as imported
#> Retrieving record values for the given time points.
#> ================================================================================
#> Retrieving record values for the given time points.
#> ================================================================================


ERG <- AutoPlaceMarkers(ERG, Channel.names = pairlist(ERG = "ERG_auto")) # automatically place markers
#> ================================================================================

ggERGTrace(ERG, where = list( Step = as.integer(3), Eye = "RE", Channel ="ERG", Result = as.integer(1))) # pick one and have a look at the traces as imported
#> Retrieving record values for the given time points.
#> ================================================================================
#> Retrieving record values for the given time points.
#> ================================================================================
```

#### Contents

Developed by Moritz Lindner.

Site built with pkgdown 2.0.7.
