## Supplemental Material 1 for "The ERGtools2 package: A Toolset for Processing and Analysing Visual Electrophysiology Data": exploreERGExam.html

Explore ERGExam Metadata and Measurements — exploreERGExam • ERGtools2       

Toggle navigation


ERGtools2
0.7.0

- Reference

### Explore ERGExam Metadata and Measurements

`exploreERGExam.Rd`

Launches a Shiny app to explore the content of the Metadata slot of the ERGExam object and allows for filtering by metadata columns Eye and Channel as well as by the Stimulus columns Description, Intensity, Background, and Type. Metadata and Stimulus are matched by the Step column in Metadata, which contains the index of the corresponding row of the data.frame in the Stimulus slot.

```
exploreERGExam(X)
```

#### Arguments

X
:   An ERGExam object.

#### Value

A Shiny app to interactively explore ERGExam metadata and measurements.

#### Examples

```
if (FALSE) {
data(ERG)
exploreERGExam(ERG)
}
```

#### Contents

Developed by Moritz Lindner.

Site built with pkgdown 2.0.7.
