## Supplemental Material 1 for "The ERGtools2 package: A Toolset for Processing and Analysing Visual Electrophysiology Data": Get.html

Accession methods for metadata from ERGExam objects — Get • ERGtools2       

Toggle navigation


ERGtools2
0.7.0

- Reference

### Accession methods for metadata from ERGExam objects

`Get.Rd`

These methods are used to access metadata information from ERGExam objects.

```
Eyes(X)

Channels(X)

Steps(X)

Results(X)

Subject(X)

MarkerNames(X)

ProtocolName(X)

GroupName(X)

ExamDate(X)

DOB(X)
```

#### Arguments

X
:   An ERGExam

#### Value

A vector. For 'StimulusTable()' a data.frame and a function for 'GetFilterFunction()' and 'GetAverageFunction()'.
FIXME: Return is not up to date
FIXME not only for Class ERGExma

#### Details

These methods can be used to access metadata information stored in ERGExam objects.

#### Functions

- `Eyes()`: Returns a the eyes of which the data has been recorded.
- `Channels()`: Returns the Channel names.
- `Steps()`: Returns the steps of the exam
- `Results()`: Returns the indices of individual results contained in an ERG exam.
- `Subject()`: Returns the subject's name
- `MarkerNames()`: Returns the measurement parameter names (e.g: 'a','B','N1','P1').
- `ProtocolName()`: Returns the recording protocol name.
- `GroupName()`: Returns the group name.
- `ExamDate()`: Returns the exam date.
- `DOB()`: Returns the date of birth.

#### Examples

```
# Get Data to work with
data(ERG)

# Accessing eyes from ERGExam object
Eyes(ERG)
#> [1] "RE" "LE"

# Accessing channels from ERGExam object
Channels(ERG)
#> [1] "ERG" "OP" 

# Accessing steps from ERGExam object
Steps(ERG)
#> [1] 1 2 3

# Accessing steps from ERGExam object
Results(ERG)
#> [1] 1

# Accessing subject from ERGExam object
Subject(ERG)
#> [1] "CR2170"

# Accessing protocol name from ERGExam object
ProtocolName(ERG)
#> [1] "01-1 MoL_v2_DA long ERG [14806-F || ECN 1685 || 13 July 2021]"

# Accessing group name from ERGExam object
GroupName(ERG)
#> integer(0)

# Accessing exam date from ERGExam object
ExamDate(ERG)
#> [1] "2023-08-10 11:18:31 CEST"

# Accessing date of birth from ERGExam object
DOB(ERG)
#> [1] "2023-05-30"
```

#### Contents

Developed by Moritz Lindner.

Site built with pkgdown 2.0.7.
