## Supplemental Material 1 for "The ERGtools2 package: A Toolset for Processing and Analysing Visual Electrophysiology Data": ggERGExam.html

Generates a ggplot2 plot from an ERGExam object. — ggERGExam • ERGtools2       

Toggle navigation


ERGtools2
0.7.0

- Reference

### Generates a ggplot2 plot from an ERGExam object.

`ggERGExam.Rd`

This function creates a ggplot2:ggplot plot of an ERGExam object, including line plots of
stimulus response curves for different label categories.

```
ggERGExam(
  X,
  return.as = "grid",
  show.markers = T,
  SetSIPrefix = "auto",
  downsample = 250
)
```

#### Arguments

X
:   An ERGExam object.

return.as
:   Whether to return as a gridExtra::grid.arrange grid (the default: 'return.as = "grid"') or as a list of ggplot2:ggplots ('return.as = "list"').

show.markers
:   Whether to return Marker Postitions as stored in the Measurements slot.

SetSIPrefix
:   Change the SI prefix. Set to `keep`, for not to change anything, to `auto` (default) for using the EPhysData:BestSIPrefix-methods to minimize the number of relevant digits or to any SI prefix to use that. Calls the EPhysData:SetSIPrefix-methods.

downsample
:   Integer giving the desired number of intervals for downsampling. Non-integer values are rounded down. Defaults to 250.

#### Value

A ggplot2:ggplot plot of the ERGExam data.

#### See also

ERGExam ggplot2:ggplot

#### Examples

```
# Example usage:
data(ERG)
ERG<-SetStandardFunctions(ERG)
exam_plot <- ggERGExam(ERG)
#> Retrieving record values for the given time points.
#> ================================================================================
print(exam_plot)

ggERGExam(ERG,SetSIPrefix="auto")
#> Retrieving record values for the given time points.
#> ================================================================================

ggERGExam(ERG,SetSIPrefix="k")
#> Retrieving record values for the given time points.
#> ================================================================================
```

#### Contents

Developed by Moritz Lindner.

Site built with pkgdown 2.0.7.
