## Supplemental Material 1 for "The ERGtools2 package: A Toolset for Processing and Analysing Visual Electrophysiology Data": ggERGTrace.html

Generate a ggplot2 plot for a single item/trace from an ERGExam objects — ggERGTrace • ERGtools2       

Toggle navigation


ERGtools2
0.7.0

- Reference

### Generate a ggplot2 plot for a single item/trace from an ERGExam objects

`ggERGTrace.Rd`

This method generates a ggplot2::ggplot plot for a single trace from anERGExam objects.

```
ggERGTrace(X, where, Interactive = F, SetSIPrefix = "auto")
```

#### Arguments

X
:   An ERGExam

where
:   A base::list defining selection criteria. Tags/Keys in the names in the list must represent valid column names of Metadata orStimulusTable.

Interactive
:   Whether to return an interactive plotly::ggplotly graph

SetSIPrefix
:   Change the SI prefix. Set to `keep`, for not to change anything, to `auto` (default) for using the EPhysData:BestSIPrefix-methods to minimize the number of relevant digits or to any SI prefix to use that. Calls the EPhysData:SetSIPrefix-methods.

#### Value

A ggplot2::ggplot plot visualizing the data from a single trace from an ERGExam object

#### Examples

```
data(ERG)
AverageFunction(ERG, where=pairlist(Step = as.integer(3),Channel = "ERG",Result = as.integer(1))) <- mean
ggERGTrace(ERG, where = list( Step = as.integer(3), Eye = "RE", Channel ="ERG", Result = as.integer(1)))
#> Retrieving record values for the given time points.
#> ================================================================================
#> Retrieving record values for the given time points.
#> ================================================================================

# the information obtained from the interactive plot can be used e.g. to update marker positions.
```

#### Contents

Developed by Moritz Lindner.

Site built with pkgdown 2.0.7.
