## Supplemental Material 1 for "The ERGtools2 package: A Toolset for Processing and Analysing Visual Electrophysiology Data": ggIntensitySequence.html

Uses ggplot2 to plot intensity sequence for ERG exams — ggIntensitySequence • ERGtools2       

Toggle navigation


ERGtools2
0.7.0

- Reference

### Uses ggplot2 to plot intensity sequence for ERG exams

`ggIntensitySequence.Rd`

This function generates a ggplot2:ggplot plot of intensity sequence data for ERG exams.

```
ggIntensitySequence(
  List,
  where = list(Background = "DA", Type = "Flash"),
  Markers = c("a", "B", "N1", "P1"),
  Parameter = "Amplitude",
  wrap_by = "Channel",
  point.size = 1,
  theme.base.size = 8
)
```

#### Arguments

List
:   A list of ERG exams.

Markers
:   Vector of markers to include in the plot.

Parameter
:   The parameter to plot ("Amplitude" or "Time").

wrap\_by
:   Wrapping parameter for facetting ("Channel" or NULL).

Background
:   Background condition for the exams.

Type
:   Type of exam (e.g., "Flash").

Channel
:   The channel to plot (e.g., "ERG").

#### Value

A ggplot2 plot object.

#### See also

ggplot2:ggplot

#### Examples

```
# Example usage:
data(ERG)
ERG<-SetStandardFunctions(ERG)
ERG <- AutoPlaceMarkers(ERG)
#> ================================================================================
data <- list(ERG, ERG)
ggIntensitySequence(data, where= list(Background = "DA", Type = "Flash", Channel = "ERG"))
#> ================================================================================
```

#### Contents

Developed by Moritz Lindner.

Site built with pkgdown 2.0.7.
