## Supplemental Material 1 for "The ERGtools2 package: A Toolset for Processing and Analysing Visual Electrophysiology Data": ggPlotRecordings.html

Uses ggplot2 to plot ERG traces from multiple ERGExam objects — ggPlotRecordings • ERGtools2       

Toggle navigation


ERGtools2
0.7.0

- Reference

### Uses ggplot2 to plot ERG traces from multiple ERGExam objects

`ggPlotRecordings.Rd`

This function generates a ggplot2:ggplot plot of ERG traces from multiple ERGExam objects.

```
ggPlotRecordings(
  List,
  where = list(),
  wrap_by = "Channel",
  scales = "free_y",
  downsample = 250
)
```

#### Arguments

List
:   A list of ERG exams.

where
:   A base::list defining selection criteria. Tags/Keys in the names in the list must represent valid column names of Metadata orStimulusTable.

wrap\_by
:   Wrapping parameter for facetting ("Channel" or NULL).

scales
:   Passed on to ggplot2:facet\_grid.

downsample
:   Integer giving the desired number of intervals for downsampling. Non-integer values are rounded down. Defaults to 250.

#### Value

A ggplot2 plot object.

#### Examples

```
# Example usage:
data(ERG)
ERG<-SetStandardFunctions(ERG)
ERG <- AutoPlaceMarkers(ERG)
#> ================================================================================
data <- list(ERG, ERG)
ggPlotRecordings(data, where = list(Background = "DA", Type = "Flash", Channel = "ERG"))
#> Running 'ggPlotRecordings()'. This may take a while for long ERGExam lists. 
#> Error in str2lang(x): <text>:2:0: unexpected end of input
#> 1: Intensity ~ 
#>    ^
```

#### Contents

Developed by Moritz Lindner.

Site built with pkgdown 2.0.7.
