## Supplemental Material 1 for "The ERGtools2 package: A Toolset for Processing and Analysing Visual Electrophysiology Data": ggStepSequence.html

Uses ggplot2 to plot step sequence for multiple ERG exams — ggStepSequence • ERGtools2       

Toggle navigation


ERGtools2
0.7.0

- Reference

### Uses ggplot2 to plot step sequence for multiple ERG exams

`ggStepSequence.Rd`

This function generates a ggplot2:ggplot plot of step sequence (i.e. sequential recordings within a single protocol) data for multiple ERG exams.

```
ggStepSequence(
  List,
  where = list(Background = "DA", Type = "Flash"),
  Markers = c("N1", "P1"),
  wrap_by = "Channel"
)
```

#### Arguments

List
:   A list of ERG exams.

where
:   A base::list defining selection criteria. Tags/Keys in the names in the list must represent valid column names of Metadata orStimulusTable.

Markers
:   Vector of markers to include in the plot (e.g. c("a","B)).

wrap\_by
:   Wrapping parameter for facetting ("Channel" or NULL).

#### Value

A ggplot2:ggplot plot object.

#### See also

ggplot2:ggplot

#### Examples

```
# Example usage:
data(ERG)
ERG<-SetStandardFunctions(ERG)
ERG <- AutoPlaceMarkers(ERG)
#> ================================================================================
data <- list(ERG, ERG)
ggStepSequence(data, where = list(Background = "DA", Type = "Flash", Channel = "ERG"), Markers = c("a", "B"))
#> ================================================================================
```

#### Contents

Developed by Moritz Lindner.

Site built with pkgdown 2.0.7.
