## Supplemental Material 1 for "The ERGtools2 package: A Toolset for Processing and Analysing Visual Electrophysiology Data": ImportEspion.html

Import ERG data and measurements from an Espion CSV file — ImportEspion • ERGtools2       

Toggle navigation


ERGtools2
0.7.0

- Reference

### Import ERG data and measurements from an Espion CSV file

`ImportEspion.Rd`

This function imports ERG recordings from \*.csv files exported from the Diagnosys Espion™ software and creates an ERGExam object.

```
ImportEspion(
  filename,
  sep = "\t",
  Import = list("Averaged", "Measurements"),
  Protocol = NULL,
  where = NULL
)

ImportEspionInfo(filename, sep = "\t")

ImportEspionMetadata(filename, sep = "\t", Protocol = NULL)

ImportEspionStimTab(filename, sep = "\t", Protocol = NULL)
```

#### Arguments

filename
:   Path to the file. Currently supported `'.csv'` or `'.txt'` files exported from the Diagnosys Espion™ software that contain at least:
    An area with 1) a contents table, 2) a header table, 3) a stimulus table, and 4) a marker table.
    This function is only tested on files with the default horizontal table arrangement but should function for files with a vertical arrangement as well.
    Behaves similarly to the argument `file` from utils::read.table().

sep
:   The field separator character. Values on each line of the file are separated by this character.
    If sep = "" (the default for read.table), the separator is ‘white space’, which includes spaces, tabs, newlines, or carriage returns.

Import
:   A list of character vectors specifying which parts of the data to import.
    Possible elements are "Raw" - for importing the raw recordings, "Averaged" - for importing the averaged recordings ("Results"), and "Measurements" - for importing the measured markers. Either of "Raw" or "Averaged" must be selected.

Protocol
:   An S4 object of class ERGProtocol() or a list thereof.

where
:   A base::list defining selection criteria. Tags/Keys in the names in the list must represent valid column names of Metadata orStimulusTable.

#### Value

For ImportEspion: A ERGExam object.

For ImportEspionInfo: A named list containing the exam info stored in an exported Espion ERG exam.

For ImportEspionMetadata: A data.frame() containing the metadata for an ERG Recording.

For ImportEspionStimTab: A data.frame() containing the stimulus information.

#### Functions

- `ImportEspionInfo()`: Read the Exam information stored in an Espion CSV file
- `ImportEspionMetadata()`: Read the Metadata information stored in an Espion Exam file
- `ImportEspionStimTab()`: Read the Stimulus information stored in an Espion Exam file

#### See also

ERGExam ERGProtocol()

#### Examples

```
if (FALSE) {
# Import a *.csv file exported from the Diagnosys Espion™ software.
ERG_Experiment <- ImportEspion("test.csv")
}

if (FALSE) {
# Import exam information from a *.csv file exported from the Diagnosys Espion™ software.
ImportEspionInfo("test.txt")
}

if (FALSE) {
# Import a *.txt file exported from the Diagnosys Espion™ software.
Metadata <- ImportEspionMetadata("test.txt")
}

if (FALSE) {
# Import stimulus information from a *.csv file exported from the Diagnosys Espion™ software.
ImportEspionInfo("test.txt")
}
```

#### Contents

Developed by Moritz Lindner.

Site built with pkgdown 2.0.7.
