## Supplemental Material 1 for "The ERGtools2 package: A Toolset for Processing and Analysing Visual Electrophysiology Data": ImportEspionProtocol.html

Import an Espion stimulus protocol from a CSV file. — ImportEspionProtocol • ERGtools2       

Toggle navigation


ERGtools2
0.7.0

- Reference

### Import an Espion stimulus protocol from a CSV file.

`ImportEspionProtocol.Rd`

This function imports an Espion stimulus protocol from a CSV file and returns it as a Protocol S4 object.

```
ImportEspionProtocol(filename)
```

#### Arguments

filename
:   The path to the CSV file containing the protocol data.

#### Value

An instance of the Protocol S4 class.

#### Contents

Developed by Moritz Lindner.

Site built with pkgdown 2.0.7.
