## Supplemental Material 1 for "The ERGtools2 package: A Toolset for Processing and Analysing Visual Electrophysiology Data": index.html

Function reference • ERGtools2       

Toggle navigation


ERGtools2
0.7.0

- Reference

### Reference

| All functions | |
| --- | --- |
|  |  |
| --- | --- |
| `AutoPlaceMarkers()` `AutoPlaceAB()` `AutoPlaceFlicker()` `AutoPlaceVEP()` | AutoPlaceMarkers for ERG/VEP Recordings |
| `CheckAvgFxSet()` | Check if all recordings in an ERGExam object have a valid averaging function set. |
| `CollectMeasurements()` | Get measurements for plotting |
| `DropRecordings()` | Drop specified recordings from an ERGExam Object |
| `ERGExam` `ERGExam-class` | ERGExam Class |
| `ERGMeasurements-class` | ERGMeasurements Class |
| `ERGProtocol` | ERGProtocol class definition |
| `ERGtools2-package` | Structured Index and Introduction for the ERGtools2 R Package |
| `Eyes()` `Channels()` `Steps()` `Results()` `Subject()` `MarkerNames()` `ProtocolName()` `GroupName()` `ExamDate()` `DOB()` | Accession methods for metadata from ERGExam objects |
| `ImportEspion()` `ImportEspionInfo()` `ImportEspionMetadata()` `ImportEspionStimTab()` | Import ERG data and measurements from an Espion CSV file |
| `ImportEspionProtocol()` | Import an Espion stimulus protocol from a CSV file. |
| `Save(<ERGExam>)` `Load.ERGExam()` | Load/Save ERGExam objects to or from HDF5 files |
| `AddMarker()` `DropMarker()` `RenameMarker()` `RenameMarker()` `Markers()` | Access and Modify the Markers Stored in an ERGExam or ERGMeasurements Object |
| `Measurements()` `` `Measurements<-`() `` `DropMeasurement()` `newERGMeasurements()` `ClearMeasurements()` | Access and Modify the Marker Measurements Stored in an ERGExam or ERGMeasurements Object |
| `MergeERGExams()` | Merge ERGExams |
| `Stimulus()` `StimulusDescription()` `` `StimulusDescription<-`() `` `StimulusIntensity()` `` `StimulusIntensity<-`() `` `StimulusBackground()` `` `StimulusBackground<-`() `` `StimulusType()` `` `StimulusType<-`() `` | Extract or Replace Parts of an ERGExam object |
| `Subset(<ERGExam>)` | Subset from ERGExam Object |
| `UpdateChannelNames()` | Update channel names in an ERGExam object |
| `` `FilterFunction<-`(<ERGExam>) `` `` `Rejected<-`(<ERGExam>) `` `` `AverageFunction<-`(<ERGExam>) `` `SetStandardFunctions()` | Set processing functions for ERGExam objects |
| `Where()` | Get index of one or several recordings and corresponding measurements |
| `as.data.frame(<ERGMeasurements>)` `as.data.frame(<ERGMarker>)` `as.data.frame(<ERGChannel>)` `as.data.frame(<ERGStep>)` `as.data.frame` | as.data.frame for ERGProtocol, ERGStep, ERGChannel and ERGMarker |
| `ERG` | Exemplary ERG Exam |
| `Measurements.data` | Exemplary ERG Measurements object |
| `exploreERGExam()` | Explore ERGExam Metadata and Measurements |
| `ggERGExam()` | Generates a ggplot2 plot from an ERGExam object. |
| `ggERGTrace()` | Generate a ggplot2 plot for a single item/trace from an ERGExam objects |
| `ggIntensitySequence()` | Uses ggplot2 to plot intensity sequence for ERG exams |
| `ggPlotRecordings()` | Uses ggplot2 to plot ERG traces from multiple ERGExam objects |
| `ggStepSequence()` | Uses ggplot2 to plot step sequence for multiple ERG exams |
| `interactiveMeasurements()` | Interactively perform measurements on an ERGExam object |
| `newERGExam()` | Create an instance of the ERGExam class |
| `show(<ERGProtocol>)` | Show method for ERGProtocol class |
| `validERGMeasurements()` | Check if there is a measurement for each marker with a non-NA Relative value in the same recording where Relative is NA |

#### Contents

Developed by Moritz Lindner.

Site built with pkgdown 2.0.7.
