## Supplemental Material 1 for "The ERGtools2 package: A Toolset for Processing and Analysing Visual Electrophysiology Data": interactiveMeasurements.html

Interactively perform measurements on an ERGExam object — interactiveMeasurements • ERGtools2       

Toggle navigation


ERGtools2
0.7.0

- Reference

### Interactively perform measurements on an ERGExam object

`interactiveMeasurements.Rd`

This method allows interactive marker placement on ERGExam objects using shiny:shiny-package

```
interactiveMeasurements(X, Channel = "ERG", Eye = Eyes(X))
```

#### Arguments

X
:   An ERGExam

#### Value

An object of class 'ERGExam' with the new measurements added.

#### Examples

```
data(ERG)
ERG<-SetStandardFunctions(ERG)
if (FALSE) {
ERG<-interactiveMeasurements(ERG, Channel="OP",Eye="RE")
}
Measurements(ERG)
#> Retrieving record values for the given time points.
#> ================================================================================
#>    Recording Step     Description Channel Result Eye Name Relative       Time
#> 1          1    1 DA 0 01 cd s m      ERG      1  RE    a     <NA> 0.0315 [s]
#> 2          1    1 DA 0 01 cd s m      ERG      1  RE    B        a 0.0515 [s]
#> 3          2    1 DA 0 01 cd s m      ERG      1  LE    a     <NA> 0.0315 [s]
#> 4          2    1 DA 0 01 cd s m      ERG      1  LE    B        a 0.0505 [s]
#> 5          5    2    DA 1 cd s m      ERG      1  RE    a     <NA> 0.0130 [s]
#> 6          5    2    DA 1 cd s m      ERG      1  RE    B        a 0.0355 [s]
#> 7          6    2    DA 1 cd s m      ERG      1  LE    a     <NA> 0.0135 [s]
#> 8          6    2    DA 1 cd s m      ERG      1  LE    B        a 0.0355 [s]
#> 9          9    3    DA 3 cd s m      ERG      1  RE    a     <NA> 0.0095 [s]
#> 10         9    3    DA 3 cd s m      ERG      1  RE    B        a 0.0345 [s]
#> 11        10    3    DA 3 cd s m      ERG      1  LE    a     <NA> 0.0100 [s]
#> 12        10    3    DA 3 cd s m      ERG      1  LE    B        a 0.0345 [s]
#>            Voltage Channel_Name Recording.y
#> 1   -53.17071 [uV]          ERG           1
#> 2   346.34079 [uV]          ERG           1
#> 3   -79.03807 [uV]          ERG           2
#> 4   303.53579 [uV]          ERG           2
#> 5  -202.88360 [uV]          ERG           5
#> 6   544.37543 [uV]          ERG           5
#> 7  -208.97144 [uV]          ERG           6
#> 8   491.28036 [uV]          ERG           6
#> 9  -232.04583 [uV]          ERG           9
#> 10  585.75366 [uV]          ERG           9
#> 11 -240.35971 [uV]          ERG          10
#> 12  568.45035 [uV]          ERG          10
if (FALSE) {
ERG<-interactiveMeasurements(ERG, Channel="ERG")
}
Measurements(ERG)
#> Retrieving record values for the given time points.
#> ================================================================================
#>    Recording Step     Description Channel Result Eye Name Relative       Time
#> 1          1    1 DA 0 01 cd s m      ERG      1  RE    a     <NA> 0.0315 [s]
#> 2          1    1 DA 0 01 cd s m      ERG      1  RE    B        a 0.0515 [s]
#> 3          2    1 DA 0 01 cd s m      ERG      1  LE    a     <NA> 0.0315 [s]
#> 4          2    1 DA 0 01 cd s m      ERG      1  LE    B        a 0.0505 [s]
#> 5          5    2    DA 1 cd s m      ERG      1  RE    a     <NA> 0.0130 [s]
#> 6          5    2    DA 1 cd s m      ERG      1  RE    B        a 0.0355 [s]
#> 7          6    2    DA 1 cd s m      ERG      1  LE    a     <NA> 0.0135 [s]
#> 8          6    2    DA 1 cd s m      ERG      1  LE    B        a 0.0355 [s]
#> 9          9    3    DA 3 cd s m      ERG      1  RE    a     <NA> 0.0095 [s]
#> 10         9    3    DA 3 cd s m      ERG      1  RE    B        a 0.0345 [s]
#> 11        10    3    DA 3 cd s m      ERG      1  LE    a     <NA> 0.0100 [s]
#> 12        10    3    DA 3 cd s m      ERG      1  LE    B        a 0.0345 [s]
#>            Voltage Channel_Name Recording.y
#> 1   -53.17071 [uV]          ERG           1
#> 2   346.34079 [uV]          ERG           1
#> 3   -79.03807 [uV]          ERG           2
#> 4   303.53579 [uV]          ERG           2
#> 5  -202.88360 [uV]          ERG           5
#> 6   544.37543 [uV]          ERG           5
#> 7  -208.97144 [uV]          ERG           6
#> 8   491.28036 [uV]          ERG           6
#> 9  -232.04583 [uV]          ERG           9
#> 10  585.75366 [uV]          ERG           9
#> 11 -240.35971 [uV]          ERG          10
#> 12  568.45035 [uV]          ERG          10
```

#### Contents

Developed by Moritz Lindner.

Site built with pkgdown 2.0.7.
