## Supplemental Material 1 for "The ERGtools2 package: A Toolset for Processing and Analysing Visual Electrophysiology Data": LoadSave-methods.html

Load/Save ERGExam objects to or from HDF5 files — LoadSave • ERGtools2       

Toggle navigation


ERGtools2
0.7.0

- Reference

### Load/Save ERGExam objects to or from HDF5 files

`LoadSave-methods.Rd`

These functions load and save ERGExam objects from or into HDF5 files.

```
# S4 method for ERGExam
Save(X, filename, overwrite = F)

Load.ERGExam(filename)
```

#### Arguments

X
:   An ERGExam object.

filename
:   Path the data is read from or written to.

overwrite
:   Should existing files be overwritten?

#### Value

- Save: Does not return any values.
- Load: An ERGExam object.

#### Functions

- `Load.ERGExam()`: Load EPhysData:EPhysData or EPhysData:EPhysSet objects from an HDF5 file

#### Examples

```
data(ERG)
ERG<-SetStandardFunctions(ERG)
fn <- tempfile()
Save(ERG, fn, overwrite=T)
#> ================================================================================
data(ERG)
ERG<-SetStandardFunctions(ERG)
fn <- tempfile()
Save(ERG, fn, overwrite=T)
#> ================================================================================
Load.ERGExam(fn)
#> =========================================================================
#> Error in value[[3L]](cond): The function stored in the 'Rejected' slot could not be applied. A likely reason is that the function is malformed or does not fit to the data stored in the object. Object has: 5 trials. Function string is: 'function (x, rejection.cutoff = 1)  {     if ("units" %in% class(x)) {         x <- drop_units(x)     }     scale <- apply(x, 2, function(x) {         scale(filter.detrend(x), center = F)     })     CoV <- apply(scale, 1, function(x) {         log1p(sd(x)/abs(mean(x)))     })     smoother <- (2/floor((length(CoV)/10)/2)) + 1     CoV <- scale(runmed(CoV, smoother), center = T)     RoV <- CoV > 0     dat_ROV <- x[RoV, ]     dat_ROV <- apply(dat_ROV, 1, function(x) {         abs(x - mean(x))/sd(x)     })     spread <- scale(abs((apply(dat_ROV, 1, mean))))     rejected <- spread > rejection.cutoff     return(rejected) }' and returned error message is 'Error in filter.detrend(x): could not find function "filter.detrend"
#> '
```

#### Contents

Developed by Moritz Lindner.

Site built with pkgdown 2.0.7.
