## Supplemental Material 1 for "The ERGtools2 package: A Toolset for Processing and Analysing Visual Electrophysiology Data": Marker-Methods.html

Access and Modify the Markers Stored in an ERGExam or ERGMeasurements Object — Marker-Methods • ERGtools2       

Toggle navigation


ERGtools2
0.7.0

- Reference

### Access and Modify the Markers Stored in an ERGExam or ERGMeasurements Object

`Marker-Methods.Rd`

These methods enable accession and modification of the Markers stored in an ERGExam object or the underlying ERGMeasurements object directly.

```
AddMarker(X, Marker, Relative = NA, ChannelBinding, update.empty.relative = F)

DropMarker(X, Marker, ChannelBinding, drop.dependent = FALSE)

RenameMarker(X, Marker, ChannelBinding, New.Name)

RenameMarker(X, Marker, ChannelBinding, New.Name)

Markers(X)
```

#### Arguments

X
:   An ERGExam or ERGMeasurements object.

Marker
:   Name of the marker.

Relative
:   For AddMarker and Measurements<- only. Index of the marker the new marker is relative to.

ChannelBinding
:   For AddMarker and DropMarker only. Channel to which the marker belongs. Usually defined by the Parent ERGExam object. Set to NA if not required.

update.empty.relative
:   Logical. If an empty relative value should be overwritten by an otherways matching marker.

drop.dependent
:   For DropMarker and DropMeasurement only. Logical. If TRUE, dependent markers will also be dropped. If FALSE, fails if there are markes dependent on that removed.

New.Name
:   The new name to assign to the marker.

#### Value

An updated version of the object, except for Markers, which returns a data.frame representing the Markers stored in the the ERGExam or the ERGMeasurements object.

An updated ERGMeasurements object with the marker renamed.

#### Functions

- `AddMarker()`: Add a new Marker
- `DropMarker()`: Drop marker by name and channel
- `RenameMarker()`: Renames a marker within an ERGMeasurements object according to the specified channel binding.
- `RenameMarker()`: Renames a marker within an ERGMeasurements object according to the specified channel binding.
- `Markers()`: Get Markers.

#### See also

EPhysData::EPhysSet-class Measurements-Methods Get

#### Examples

```
# Add a new Marker to an ERGMeasurements object#'
data(Measurements.data)
new_erg_measurements <- AddMarker(Measurements.data, Marker = "Marker1", Relative = NA, ChannelBinding = "ChannelA")
new_erg_measurements
#> ERGMeasurements object:
#> Measurements:
#>   Recording Name ChannelBinding Relative    Time
#> 2         1    a            ERG     <NA> 10 [ms]
#> 3         1    B            ERG        a 15 [ms]
#> 4         2   N1            VEP     <NA> 20 [ms]
#> 5         2   P1            VEP       N1 25 [ms]
#> 6         3    a            ERG     <NA> 30 [ms]
#> 7         3    B            ERG        a 35 [ms]
#> 
#> Unused Markers:
#>      Name Relative ChannelBinding
#> 5 Marker1     <NA>       ChannelA
Markers(new_erg_measurements)
#>      Name Relative ChannelBinding
#> 1       a     <NA>            ERG
#> 2       B        a            ERG
#> 3      N1     <NA>            VEP
#> 4      P1       N1            VEP
#> 5 Marker1     <NA>       ChannelA

# Add a new Marker to an ERGMeasurements object
data(ERG)
new_erg <- AddMarker(ERG, Marker = "Marker1", Relative = NA, ChannelBinding = "ERG")
# but this would fail:
# new_erg <- DropMarker(new_erg, Marker = "Marker1")
Markers(new_erg)
#>      Name Relative ChannelBinding
#> 1       a     <NA>            ERG
#> 2       B        a            ERG
#> 3 Marker1     <NA>            ERG

# Drop a Marker from an ERGMeasurements object
data(Measurements.data)
Measurements.data
#> ERGMeasurements object:
#> Measurements:
#>   Recording Name ChannelBinding Relative    Time
#> 1         1    a            ERG     <NA> 10 [ms]
#> 2         1    B            ERG        a 15 [ms]
#> 3         2   N1            VEP     <NA> 20 [ms]
#> 4         2   P1            VEP       N1 25 [ms]
#> 5         3    a            ERG     <NA> 30 [ms]
#> 6         3    B            ERG        a 35 [ms]
Markers(Measurements.data)
#>   Name Relative ChannelBinding
#> 1    a     <NA>            ERG
#> 2    B        a            ERG
#> 3   N1     <NA>            VEP
#> 4   P1       N1            VEP

new_erg_measurements <- DropMarker(Measurements.data, Marker = "N1", ChannelBinding = "VEP", drop.dependent = TRUE)
new_erg_measurements
#> ERGMeasurements object:
#> Measurements:
#>   Recording Name ChannelBinding Relative    Time
#> 1         1    a            ERG     <NA> 10 [ms]
#> 2         1    B            ERG        a 15 [ms]
#> 3         3    a            ERG     <NA> 30 [ms]
#> 4         3    B            ERG        a 35 [ms]
Markers(new_erg_measurements)
#>   Name Relative ChannelBinding
#> 1    a     <NA>            ERG
#> 2    B        a            ERG

data(Measurements.data)
RenameMarker(Measurements.data,"a","ERG","X")
#> ERGMeasurements object:
#> Measurements:
#>   Recording Name ChannelBinding Relative    Time
#> 1         1    X            ERG     <NA> 10 [ms]
#> 2         1    B            ERG        X 15 [ms]
#> 3         2   N1            VEP     <NA> 20 [ms]
#> 4         2   P1            VEP       N1 25 [ms]
#> 5         3    X            ERG     <NA> 30 [ms]
#> 6         3    B            ERG        X 35 [ms]

data(Measurements.data)  # Assume Measurements.data is a pre-loaded ERGMeasurements object
updated_measurements <- RenameMarker(Measurements.data, Marker = "a", ChannelBinding = "ERG", New.Name = "NewMarkerName")
updated_measurements
#> ERGMeasurements object:
#> Measurements:
#>   Recording          Name ChannelBinding      Relative    Time
#> 1         1 NewMarkerName            ERG          <NA> 10 [ms]
#> 2         1             B            ERG NewMarkerName 15 [ms]
#> 3         2            N1            VEP          <NA> 20 [ms]
#> 4         2            P1            VEP            N1 25 [ms]
#> 5         3 NewMarkerName            ERG          <NA> 30 [ms]
#> 6         3             B            ERG NewMarkerName 35 [ms]
```

#### Contents

Developed by Moritz Lindner.

Site built with pkgdown 2.0.7.
