## Supplemental Material 1 for "The ERGtools2 package: A Toolset for Processing and Analysing Visual Electrophysiology Data": Measurements-Methods.html

Access and Modify the Marker Measurements Stored in an ERGExam or ERGMeasurements Object — Measurements-Methods • ERGtools2       

Toggle navigation


ERGtools2
0.7.0

- Reference

### Access and Modify the Marker Measurements Stored in an ERGExam or ERGMeasurements Object

`Measurements-Methods.Rd`

These methods enable modification of the marker measurements (hereafter short: measurements) stored in an ERGExam object or the underlying ERGMeasurements object directly.
**Note:** the `Measurements(X, ...) <- value` methods require that all arguments are provided. Set to NULL if not needed.

```
Measurements(X, where = NULL, Marker = NULL, quiet = F, ...)

# S4 method for ERGMeasurements
Measurements(X, where = NULL, Marker = NULL, quiet = F)

# S4 method for ERGExam
Measurements(
  X,
  where = NULL,
  Marker = NULL,
  quiet = F,
  measure.absolute = F,
  TimesOnly = F
)

Measurements(
  X,
  Marker,
  where,
  create.marker.if.missing = T,
  Relative = NULL,
  ChannelBinding = NULL
) <- value

# S4 method for ERGMeasurements
Measurements(
  X,
  Marker,
  where,
  create.marker.if.missing = T,
  Relative = NULL,
  ChannelBinding = NULL
) <- value

# S4 method for ERGExam
Measurements(
  X,
  Marker,
  where = NULL,
  create.marker.if.missing = T,
  Relative = NULL,
  ChannelBinding = NULL
) <- value

DropMeasurement(X, Marker, where, ChannelBinding = NULL)

# S4 method for ERGExam
DropMeasurement(X, Marker, where = NULL, ChannelBinding = NULL)

newERGMeasurements(data, update.empty.relative = F)

ClearMeasurements(X)
```

#### Arguments

X
:   An ERGExam or ERGMeasurements object.

where
:   A base::list defining selection criteria. Tags/Keys in the names in the list must represent valid column names of Metadata orStimulusTable.

Marker
:   Name of the marker.

quiet
:   For Measurements only. Logical. If TRUE, suppresses warnings and progress bars.

measure.absolute
:   Logical, default: FALSE. If absolute amplitudes should be returned instead of amplitudes relative to the reference marker (where given).

TimesOnly
:   For Measurements only. Logical. If TRUE, only fetches times, not amplitudes

create.marker.if.missing
:   Logical. If TRUE and the marker does not exist, it will be created. Dies nothing if value is set to `NULL`

Relative
:   For AddMarker and Measurements<- only. Index of the marker the new marker is relative to.

ChannelBinding
:   For AddMarker and DropMarker only. Channel to which the marker belongs. Usually defined by the Parent ERGExam object. Set to NA if not required.

value
:   For Measurements<- only. Plain numeric value or value of class units (units::units with a time unit set. If `NULL` it will remove the indicated row.

data
:   For newERGMeasurements only. A data Measurementsframe containing measurements data with columns:
    `Channel`, `Name`, `Recording`, `Time`, and `Relative`.

update.empty.relative
:   Logical. If an empty relative value should be overwritten by an otherways matching marker.

#### Value

An updated version of the object, except for Measurements, which returns a data.frame representing the Measurements stored in the Measurements slot of the ERGExam object or the ERGMeasurements object-

#### Functions

- `Measurements()`: Get Measurements table
- `Measurements(ERGMeasurements)`: Get Measurements table from an ERGMeasurements object
- `Measurements(ERGExam)`: Get Measurements table from an ERGExam object
- `Measurements(
  X,
  Marker,
  where,
  create.marker.if.missing = T,
  Relative = NULL,
  ChannelBinding = NULL
  ) <- value`: Update, add, or remove a line to the Measurements slot
- `Measurements(ERGMeasurements) <- value`: Update, add, or remove a measurement from an ERGMeasurements object
- `Measurements(ERGExam) <- value`: Update, add, or remove a measurement from the Measurements slot of an ERGExam object
- `DropMeasurement()`: Remove a measurement from the Measurements slot of an ERGExam object
- `DropMeasurement(ERGExam)`: Remove a measurement from the Measurements slot of an ERGExam object
- `newERGMeasurements()`: #' Create an ERGMeasurements object from data
- `ClearMeasurements()`: Clears the Measurements slots in an link[=ERGExam]ERGExam object (both, markers and measurements.

#### See also

EPhysData::EPhysSet-class Measurements-Methods Get

#### Examples

```
data(Measurements.data)
# Get Measurements table for specific recording and marker in ERGMeasurements object
Markers(Measurements.data)
#>   Name Relative ChannelBinding
#> 1    a     <NA>            ERG
#> 2    B        a            ERG
#> 3   N1     <NA>            VEP
#> 4   P1       N1            VEP
Measurements(X = Measurements.data, where = 1, Marker = "a", quiet = TRUE)
#>   Recording Name ChannelBinding Relative    Time
#> 1         1    a            ERG     <NA> 10 [ms]
#' but this would fail with an informative error message
#' Measurements(X = Measurements.data, where = 1, Marker = "X", quiet = F)

# Get Measurements table for specific recording and marker in ERGExam object
data(ERG)
ERG<-SetStandardFunctions(ERG)
Measurements(ERG)
#> Retrieving record values for the given time points.
#> ================================================================================
#>    Recording Step     Description Channel Result Eye Name Relative       Time
#> 1          1    1 DA 0 01 cd s m      ERG      1  RE    a     <NA> 0.0315 [s]
#> 2          1    1 DA 0 01 cd s m      ERG      1  RE    B        a 0.0515 [s]
#> 3          2    1 DA 0 01 cd s m      ERG      1  LE    a     <NA> 0.0315 [s]
#> 4          2    1 DA 0 01 cd s m      ERG      1  LE    B        a 0.0505 [s]
#> 5          5    2    DA 1 cd s m      ERG      1  RE    a     <NA> 0.0130 [s]
#> 6          5    2    DA 1 cd s m      ERG      1  RE    B        a 0.0355 [s]
#> 7          6    2    DA 1 cd s m      ERG      1  LE    a     <NA> 0.0135 [s]
#> 8          6    2    DA 1 cd s m      ERG      1  LE    B        a 0.0355 [s]
#> 9          9    3    DA 3 cd s m      ERG      1  RE    a     <NA> 0.0095 [s]
#> 10         9    3    DA 3 cd s m      ERG      1  RE    B        a 0.0345 [s]
#> 11        10    3    DA 3 cd s m      ERG      1  LE    a     <NA> 0.0100 [s]
#> 12        10    3    DA 3 cd s m      ERG      1  LE    B        a 0.0345 [s]
#>            Voltage Channel_Name Recording.y
#> 1   -53.17071 [uV]          ERG           1
#> 2   346.34079 [uV]          ERG           1
#> 3   -79.03807 [uV]          ERG           2
#> 4   303.53579 [uV]          ERG           2
#> 5  -202.88360 [uV]          ERG           5
#> 6   544.37543 [uV]          ERG           5
#> 7  -208.97144 [uV]          ERG           6
#> 8   491.28036 [uV]          ERG           6
#> 9  -232.04583 [uV]          ERG           9
#> 10  585.75366 [uV]          ERG           9
#> 11 -240.35971 [uV]          ERG          10
#> 12  568.45035 [uV]          ERG          10
Measurements(ERG, measure.absolute = T)
#> Retrieving record values for the given time points.
#> ================================================================================
#>    Recording Step     Description Channel Result Eye Name Relative       Time
#> 1          1    1 DA 0 01 cd s m      ERG      1  RE    a     <NA> 0.0315 [s]
#> 2          1    1 DA 0 01 cd s m      ERG      1  RE    B        a 0.0515 [s]
#> 3          2    1 DA 0 01 cd s m      ERG      1  LE    a     <NA> 0.0315 [s]
#> 4          2    1 DA 0 01 cd s m      ERG      1  LE    B        a 0.0505 [s]
#> 5          5    2    DA 1 cd s m      ERG      1  RE    a     <NA> 0.0130 [s]
#> 6          5    2    DA 1 cd s m      ERG      1  RE    B        a 0.0355 [s]
#> 7          6    2    DA 1 cd s m      ERG      1  LE    a     <NA> 0.0135 [s]
#> 8          6    2    DA 1 cd s m      ERG      1  LE    B        a 0.0355 [s]
#> 9          9    3    DA 3 cd s m      ERG      1  RE    a     <NA> 0.0095 [s]
#> 10         9    3    DA 3 cd s m      ERG      1  RE    B        a 0.0345 [s]
#> 11        10    3    DA 3 cd s m      ERG      1  LE    a     <NA> 0.0100 [s]
#> 12        10    3    DA 3 cd s m      ERG      1  LE    B        a 0.0345 [s]
#>            Voltage Channel_Name Recording.y
#> 1   -53.17071 [uV]          ERG           1
#> 2   293.17008 [uV]          ERG           1
#> 3   -79.03807 [uV]          ERG           2
#> 4   224.49772 [uV]          ERG           2
#> 5  -202.88360 [uV]          ERG           5
#> 6   341.49184 [uV]          ERG           5
#> 7  -208.97144 [uV]          ERG           6
#> 8   282.30891 [uV]          ERG           6
#> 9  -232.04583 [uV]          ERG           9
#> 10  353.70783 [uV]          ERG           9
#> 11 -240.35971 [uV]          ERG          10
#> 12  328.09065 [uV]          ERG          10
Measurements(ERG, TimesOnly = T)
#>    Recording Step     Description Channel Result Eye Name Relative       Time
#> 1          1    1 DA 0 01 cd s m      ERG      1  RE    a     <NA> 0.0315 [s]
#> 2          1    1 DA 0 01 cd s m      ERG      1  RE    B        a 0.0515 [s]
#> 3          2    1 DA 0 01 cd s m      ERG      1  LE    a     <NA> 0.0315 [s]
#> 4          2    1 DA 0 01 cd s m      ERG      1  LE    B        a 0.0505 [s]
#> 5          5    2    DA 1 cd s m      ERG      1  RE    a     <NA> 0.0130 [s]
#> 6          5    2    DA 1 cd s m      ERG      1  RE    B        a 0.0355 [s]
#> 7          6    2    DA 1 cd s m      ERG      1  LE    a     <NA> 0.0135 [s]
#> 8          6    2    DA 1 cd s m      ERG      1  LE    B        a 0.0355 [s]
#> 9          9    3    DA 3 cd s m      ERG      1  RE    a     <NA> 0.0095 [s]
#> 10         9    3    DA 3 cd s m      ERG      1  RE    B        a 0.0345 [s]
#> 11        10    3    DA 3 cd s m      ERG      1  LE    a     <NA> 0.0100 [s]
#> 12        10    3    DA 3 cd s m      ERG      1  LE    B        a 0.0345 [s]
#>    Voltage Channel_Name Recording.y
#> 1  NA [uV]          ERG           1
#> 2  NA [uV]          ERG           1
#> 3  NA [uV]          ERG           2
#> 4  NA [uV]          ERG           2
#> 5  NA [uV]          ERG           5
#> 6  NA [uV]          ERG           5
#> 7  NA [uV]          ERG           6
#> 8  NA [uV]          ERG           6
#> 9  NA [uV]          ERG           9
#> 10 NA [uV]          ERG           9
#> 11 NA [uV]          ERG          10
#> 12 NA [uV]          ERG          10

# Set Measurements for specific recording and marker in ERGMeasurements object
Markers(Measurements.data)
#>   Name Relative ChannelBinding
#> 1    a     <NA>            ERG
#> 2    B        a            ERG
#> 3   N1     <NA>            VEP
#> 4   P1       N1            VEP
Measurements(X = Measurements.data, Marker = "B" ,where = 1)<-99
# this would fail, as marker does not exist
# Measurements.data<-Measurements(X = Measurements.data, Marker = "c" , where = 1, Relative = "a")<-150
# But this would work
Measurements(X = Measurements.data, Marker = "c" ,where = 1,Relative = "a", ChannelBinding="ERG")<-150
# Set Measurements table for specific recording and marker in ERGExam object
ERG<-SetStandardFunctions(ERG)
Measurements(ERG)
#> Retrieving record values for the given time points.
#> ================================================================================
#>    Recording Step     Description Channel Result Eye Name Relative       Time
#> 1          1    1 DA 0 01 cd s m      ERG      1  RE    a     <NA> 0.0315 [s]
#> 2          1    1 DA 0 01 cd s m      ERG      1  RE    B        a 0.0515 [s]
#> 3          2    1 DA 0 01 cd s m      ERG      1  LE    a     <NA> 0.0315 [s]
#> 4          2    1 DA 0 01 cd s m      ERG      1  LE    B        a 0.0505 [s]
#> 5          5    2    DA 1 cd s m      ERG      1  RE    a     <NA> 0.0130 [s]
#> 6          5    2    DA 1 cd s m      ERG      1  RE    B        a 0.0355 [s]
#> 7          6    2    DA 1 cd s m      ERG      1  LE    a     <NA> 0.0135 [s]
#> 8          6    2    DA 1 cd s m      ERG      1  LE    B        a 0.0355 [s]
#> 9          9    3    DA 3 cd s m      ERG      1  RE    a     <NA> 0.0095 [s]
#> 10         9    3    DA 3 cd s m      ERG      1  RE    B        a 0.0345 [s]
#> 11        10    3    DA 3 cd s m      ERG      1  LE    a     <NA> 0.0100 [s]
#> 12        10    3    DA 3 cd s m      ERG      1  LE    B        a 0.0345 [s]
#>            Voltage Channel_Name Recording.y
#> 1   -53.17071 [uV]          ERG           1
#> 2   346.34079 [uV]          ERG           1
#> 3   -79.03807 [uV]          ERG           2
#> 4   303.53579 [uV]          ERG           2
#> 5  -202.88360 [uV]          ERG           5
#> 6   544.37543 [uV]          ERG           5
#> 7  -208.97144 [uV]          ERG           6
#> 8   491.28036 [uV]          ERG           6
#> 9  -232.04583 [uV]          ERG           9
#> 10  585.75366 [uV]          ERG           9
#> 11 -240.35971 [uV]          ERG          10
#> 12  568.45035 [uV]          ERG          10
# Example data frame
data <- data.frame(
  Channel = c("ERG", "ERG", "ERG", "ERG"),
  Name = c("Marker1", "Marker2", "Marker1", "Marker2"),
  Recording = c(1, 1, 2, 2),
  Time = c(10, 12, 15, 18),
  Relative = c(NA, "Marker1", NA, "Marker1")
)
# Create ERGMeasurements object
newERGMeasurements(data)
#> ERGMeasurements object:
#> Measurements:
#>   Recording    Name ChannelBinding Relative   Time
#> 1         1 Marker1            ERG     <NA> 10 [s]
#> 2         1 Marker2            ERG  Marker1 12 [s]
#> 3         2 Marker1            ERG     <NA> 15 [s]
#> 4         2 Marker2            ERG  Marker1 18 [s]
```

#### Contents

Developed by Moritz Lindner.

Site built with pkgdown 2.0.7.
