## Supplemental Material 1 for "The ERGtools2 package: A Toolset for Processing and Analysing Visual Electrophysiology Data": merge2ERGMeasurements.html

Merge two ERGMeasurements objects — merge2ERGMeasurements • ERGtools2       

Toggle navigation


ERGtools2
0.7.0

- Reference

### Merge two ERGMeasurements objects

`merge2ERGMeasurements.Rd`

This function merges two ERGMeasurements objects by combining their Marker and Measurements data frames.
It also provides options to update recording indices and handle marker mismatches.

```
merge2ERGMeasurements(
  obj1,
  obj2,
  increment.obj2.recording.index.by = NULL,
  obj2.recording.index.old = NULL,
  obj2.recording.index.new = NULL
)
```

#### Arguments

obj1
:   An object of class ERGMeasurements.

obj2
:   Another object of class ERGMeasurements.

increment.obj2.recording.index.by
:   Depreciated. Numeric indicating by what number to increment the recording indices by. Could be e.g. the number of recordings in object 1.

obj2.recording.index.old, obj2.recording.index.old
:   Two vectors of the same length indicating how the obj2 recording indices hould be updated.

#### Value

An object of class ERGMeasurements, representing the merged data.

#### Examples

```
# load the example Measurements object
data(Measurements.data)

# create a second Measurements object
# Create marker data frame
marker_df <- data.frame(
  Name = c("N1", "P1", "a", "C"),
  Relative = c(NA, 1, NA, 3),
  ChannelBinding =c("VEP","VEP","ERG","ERG")
)

# Create measurements data frame
measurements_df <- data.frame(
  Recording = c(1, 1, 4, 4, 3, 3),
  Marker = c(1, 2, 3, 4, 1, 2),
  Time = as_units(c(10, 40, 19, 26, 34, 31), "ms")
)

# Create ERGMeasurements object
Measurements.data2 <- new("ERGMeasurements", Marker = marker_df, Measurements = measurements_df)
# Show the object
Measurements.data2
#> ERGMeasurements object:
#> Measurements:
#>   Recording Name ChannelBinding Relative    Time
#> 1         1   N1            VEP     <NA> 10 [ms]
#> 2         1   P1            VEP       N1 40 [ms]
#> 3         3   N1            VEP     <NA> 34 [ms]
#> 4         3   P1            VEP       N1 31 [ms]
#> 5         4    a            ERG     <NA> 19 [ms]
#> 6         4    C            ERG        a 26 [ms]

# now merge both objects
merge2ERGMeasurements(Measurements.data,Measurements.data2, max(Measurements(Measurements.data)[,"Recording"]))
#> ERGMeasurements object:
#> Measurements:
#>    Recording Name ChannelBinding Relative    Time
#> 1          1    a            ERG     <NA> 10 [ms]
#> 2          1    B            ERG        a 15 [ms]
#> 3          2   N1            VEP     <NA> 20 [ms]
#> 4          2   P1            VEP       N1 25 [ms]
#> 5          3    a            ERG     <NA> 30 [ms]
#> 6          3    B            ERG        a 35 [ms]
#> 7          4   N1            VEP     <NA> 10 [ms]
#> 8          4   P1            VEP       N1 40 [ms]
#> 9          6   N1            VEP     <NA> 34 [ms]
#> 10         6   P1            VEP       N1 31 [ms]
#> 11         7    a            ERG     <NA> 19 [ms]
#> 12         7    C            ERG        a 26 [ms]

# other example

marker_df3 <- data.frame(
Name = c("N2", "P2", "b", "D", "E"),
Relative = c(NA, 1, NA, 3, 3),
ChannelBinding = c("VEP", "VEP", "ERG", "ERG", "ERG")
)

measurements_df3 <- data.frame(
  Recording = c(1, 1, 4, 4, 5, 5, 3, 3),
  Marker = c(1, 2, 3, 4, 3, 5, 1, 2),
  Time = as_units(c(10, 40, 19, 26, 34, 31, 15, 25), "ms")
)

Measurements.data3 <- new("ERGMeasurements", Marker = marker_df3, Measurements = measurements_df3)

merge2ERGMeasurements(Measurements.data2,Measurements.data3,max(Measurements(Measurements.data2)[,"Recording"]))
#> ERGMeasurements object:
#> Measurements:
#>    Recording Name ChannelBinding Relative    Time
#> 1          1   N1            VEP     <NA> 10 [ms]
#> 2          1   P1            VEP       N1 40 [ms]
#> 3          3   N1            VEP     <NA> 34 [ms]
#> 4          3   P1            VEP       N1 31 [ms]
#> 5          4    a            ERG     <NA> 19 [ms]
#> 6          4    C            ERG        a 26 [ms]
#> 7          5   N2            VEP     <NA> 10 [ms]
#> 8          5   P2            VEP       N2 40 [ms]
#> 9          7   N2            VEP     <NA> 15 [ms]
#> 10         7   P2            VEP       N2 25 [ms]
#> 11         8    b            ERG     <NA> 19 [ms]
#> 12         8    D            ERG        b 26 [ms]
#> 13         9    b            ERG     <NA> 34 [ms]
#> 14         9    E            ERG        b 31 [ms]
```

#### Contents

Developed by Moritz Lindner.

Site built with pkgdown 2.0.7.
