## Supplemental Material 1 for "The ERGtools2 package: A Toolset for Processing and Analysing Visual Electrophysiology Data": MergeERGExams.html

Merge ERGExams — MergeERGExams • ERGtools2       

Toggle navigation


ERGtools2
0.7.0

- Reference

### Merge ERGExams

`MergeERGExams.Rd`

Merges two or more ERGExam objects into a single ERGExam object.

```
MergeERGExams(ERGExam_list, mergemethod = "Append")
```

#### Arguments

ERGExam\_list
:   A list of ERGExam objects to be merged with `X`.

mergemethod
:   . Depreciated.

#### Value

An ERGExam object representing the merged data. the file names, protocol names and dates from the individual files merged are stored as additional columns in the Stimulus table. Use `Metadata()` to see those.

#### Examples

```
# Merge two ERGExams
data(ERG)
# make some ERGExams objects that differ
exam1<-Subset(ERG,where=list(Step=as.integer(c(1,2,3))))
exam2<-Subset(ERG,where=list(Eye="RE"))
# merge
merged_exam <- MergeERGExams(list(exam1, exam2))#'
#> Merging exams is experimental.
```

#### Contents

Developed by Moritz Lindner.

Site built with pkgdown 2.0.7.
