## Supplemental Material 1 for "The ERGtools2 package: A Toolset for Processing and Analysing Visual Electrophysiology Data": newERGExam.html

Create an instance of the ERGExam class — newERGExam • ERGtools2       

Toggle navigation


ERGtools2
0.7.0

- Reference

### Create an instance of the ERGExam class

`newERGExam.Rd`

This function creates an instance of the ERGExam class.

```
newERGExam(
  Data,
  Metadata,
  Stimulus,
  Averaged = FALSE,
  Measurements = new("ERGMeasurements"),
  ExamInfo,
  SubjectInfo
)
```

#### Arguments

Data
:   A list of EPhysData::EPhysData objects.

Metadata
:   A data frame containing metadata information associated with the data, each row corresponds to one list item.

    Step
    :   An integer vector pointing to a row index of the Stimulus table.

    Eye
    :   A character vector containing the possible values "RE" (right eye) and "LE" (left eye).

    Channel
    :   A character vector containing the unique names of the third level of `Data`.

Stimulus
:   A data frame containing stimulus information associated with the data.

    Description
    :   A character vector describing the stimuli.

    Background
    :   An integer vector representing the background intensity of the stimulus (unitless).

    Intensity
    :   An numeric vector representing the intensity of the stimulus (unitless).

Averaged
:   A list of averaged data.

Measurements
:   An object of class ERGMeasurements.

ExamInfo
:   A list containing exam-related information.

    ProtocolName
    :   A character vector indicating the name of the protocol.

    Version
    :   Optional: A character vector indicating the version of the protocol.

    ExamDate
    :   A `POSIXct` timestamp representing the date of the exam.

    Filename
    :   Optional: A character vector indicating the filename associated with the exam data.

    RecMode
    :   Optional: A character vector indicating the recording mode during the exam.

    Investigator
    :   Optional: A character vector indicating the name of the investigator conducting the exam.

SubjectInfo
:   A list containing subject-related information.

    Subject
    :   A character vector (required) indicating the name of the patient.

    DOB
    :   A `Date` object (required) indicating the date of birth of the patient.

    Gender
    :   Optional: A character vector indicating the gender of the patient.

    Group
    :   Optional: A character vector indicating the group to which the patient belongs.

#### Value

An object of class `ERGExam`.

#### See also

ERGExam

#### Examples

```
# Create example data and metadata
Data <-
  list(
    makeExampleEPhysData(sample_points = 100, replicate_count = 3),
    makeExampleEPhysData(sample_points = 100, replicate_count = 3),
    makeExampleEPhysData(sample_points = 100, replicate_count = 6),
    makeExampleEPhysData(sample_points = 100, replicate_count = 6)
  )  # List of EPhysData objects
Metadata <-
  data.frame(
    Step = as.integer(c(1,1,2,2)),
    Eye = c("RE", "LE", "LE", "LE"),
    Channel = c("Ch1", "Ch1", "Ch1", "Ch2"),
    Result = as.integer(c(1,1,1,1))
  )
Stimulus <-
  data.frame(
             Step = as.integer(c(1,1,2,2)),
             Description = c("Stim1", "Stim2"),
             Intensity = as.numeric(c(1,10)),
             Background = c("DA","DA"),
             Type = c("Flash","Flash"))  # Example stimulus data
ExamInfo <-
  list(
    ProtocolName = "ERG Protocol",
    Version = "1.0",
    ExamDate = as.POSIXct("2023-08-14"),
    Filename = "exam_data.csv"
  )
SubjectInfo <-
  list(
    Subject = "John Doe",
    DOB = as.Date("1990-05-15"),
    Gender = "Male",
    Group = "Control"
  )
ergExam <-
  newERGExam(
    Data = Data,
    Metadata = Metadata,
    Stimulus = Stimulus,
    ExamInfo = ExamInfo,
    SubjectInfo = SubjectInfo
  )
```

#### Contents

Developed by Moritz Lindner.

Site built with pkgdown 2.0.7.
