## Supplemental Material 1 for "The ERGtools2 package: A Toolset for Processing and Analysing Visual Electrophysiology Data": show.ERGProtocol.html

Show method for ERGProtocol class — show.ERGProtocol • ERGtools2       

Toggle navigation


ERGtools2
0.7.0

- Reference

### Show method for ERGProtocol class

`show.ERGProtocol.Rd`

Show method for ERGProtocol class

```
# S4 method for ERGProtocol
show(object)
```

#### Arguments

object
:   An instance of the ERGProtocol class.

...
:   Additional arguments (not used).

#### Contents

Developed by Moritz Lindner.

Site built with pkgdown 2.0.7.
