## Supplemental Material 1 for "The ERGtools2 package: A Toolset for Processing and Analysing Visual Electrophysiology Data": StimulusTableMethods.html

Extract or Replace Parts of an ERGExam object — StimulusTableMethods • ERGtools2       

Toggle navigation


ERGtools2
0.7.0

- Reference

### Extract or Replace Parts of an ERGExam object

`StimulusTableMethods.Rd`

Methods acting on ERGExam to extract or replace parts.

```
Stimulus(X, where = NULL, full = F)

StimulusDescription(X, where = NULL)

StimulusDescription(X, where = NULL) <- value

StimulusIntensity(X, where = NULL)

StimulusIntensity(X, where = NULL) <- value

StimulusBackground(X, where = NULL)

StimulusBackground(X, where = NULL) <- value

StimulusType(X, where = NULL)

StimulusType(X, where = NULL) <- value
```

#### Arguments

X
:   An ERGExam

where
:   A base::list defining selection criteria. Tags/Keys in the names in the list must represent valid column names of Metadata orStimulusTable.

full
:   For `Stimulus` only. Whether to return the full stimulus table (i.e. also any additional data that might have been added by the user or when merging single ERGExam using MergeERGExams) or only the main columns "Step", "Description", "Intensity", "Background" and "Type". Default is false.

value
:   For '<-' methods only. Vector of the same length as Stimuli selected by where containing the values to be assigned.

#### Value

For '<-' methods: an updated ERGExam object. For others: a vector containing the extracted values.

#### Functions

- `Stimulus()`: Returns selected rows of a stimulus table.
- `StimulusDescription()`: Returns the description of one or more selected stimuli.
- `StimulusDescription(X, where = NULL) <- value`: Sets the description of one or more selected stimuli.
- `StimulusIntensity()`: Returns the intensity value of one or more selected stimuli.
- `StimulusIntensity(X, where = NULL) <- value`: Sets the intensity value for one or more selected stimuli.
- `StimulusBackground()`: Returns the background value of one or more selected stimuli.
- `StimulusBackground(X, where = NULL) <- value`: Sets the background value of one or more selected stimuli.
- `StimulusType()`: Returns the type of one or more selected stimuli.
- `StimulusType(X, where = NULL) <- value`: Sets the type value of one or more selected stimuli.

#### Examples

```
# Load the ERG dataset
data(ERG)

# Extracting selected rows of the stimulus table
selected_rows <- Stimulus(ERG, where = list(Step = as.integer(c(1, 2, 3))))
head(selected_rows)
#>   Step     Description Intensity Background  Type
#> 1    1 DA 0 01 cd s m       0.01         DA Flash
#> 2    2    DA 1 cd s m       1.00         DA Flash
#> 3    3    DA 3 cd s m       3.00         DA Flash

# Extracting the description of selected stimuli
descriptions <- StimulusDescription(ERG, where = list(Background = "DA"))
descriptions
#> [1] "DA 0 01 cd s m " "DA 1 cd s m "    "DA 3 cd s m "   

# Extracting the intensity value of selected stimuli
intensity <- StimulusIntensity(ERG, where = list(Type = "Flash"))
intensity
#> [1] 0.01 1.00 3.00

# Extracting the background value of selected stimuli
background <- StimulusBackground(ERG, where = list(Step = c(1:7)))
background
#> [1] "DA" "DA" "DA"

# Extracting the type of selected stimuli
types <- StimulusType(ERG, where = list(Step = as.integer(c(1, 2, 3))))
types
#> [1] "Flash" "Flash" "Flash"

# Assigning new values to the description of selected stimuli
Stimulus(ERG)
#>   Step     Description Intensity Background  Type
#> 1    1 DA 0 01 cd s m       0.01         DA Flash
#> 2    2    DA 1 cd s m       1.00         DA Flash
#> 3    3    DA 3 cd s m       3.00         DA Flash
new_descriptions <- c("New Desc 1", "New Desc 2", "New Desc 2")
StimulusDescription(ERG, where = list(Type = "Flash")) <- new_descriptions
# Now let's check if the descriptions have been updated
updated_descriptions <- Stimulus(ERG)
updated_descriptions
#>   Step Description Intensity Background  Type
#> 1    1  New Desc 1      0.01         DA Flash
#> 2    2  New Desc 2      1.00         DA Flash
#> 3    3  New Desc 2      3.00         DA Flash
```

#### Contents

Developed by Moritz Lindner.

Site built with pkgdown 2.0.7.
