## Supplemental Material 1 for "The ERGtools2 package: A Toolset for Processing and Analysing Visual Electrophysiology Data": Subset-method.html

Subset from ERGExam Object — Subset-method • ERGtools2       

Toggle navigation


ERGtools2
0.7.0

- Reference

### Subset from ERGExam Object

`Subset-method.Rd`

This method subsets an `ERGExam` object into a new object of the same class.

```
# S4 method for ERGExam
Subset(
  X,
  Time = NULL,
  TimeExclusive = FALSE,
  Trials = NULL,
  Raw = TRUE,
  where = NULL,
  ...
)
```

#### Arguments

X
:   An ERGExam

Time
:   Numeric vector of length 2 representing the time range for data extraction.
    Default is the entire time range (i.e., keep all data).

TimeExclusive
:   Keep only the two time points stated under 'Time', not the range.

Trials
:   Specifies which of the repeated measurements (if any) to use for extraction.
    It can be either a numeric vector specifying the indices of the repeated measurements
    or a logical vector of the same length as repeats stored,
    where `TRUE` indicates using that column for extraction. Default is the inverse of the `Rejected-method`(X) vector.

Raw
:   Logical indicating whether to get raw data or processed (filtered, averaged) data.

where
:   A base::list defining selection criteria. Tags/Keys in the names in the list must represent valid column names of Metadata orStimulusTable.

#### Details

The `Subset` function creates a new `ERGExam` object containing a subset of the data from the original object, based on the provided parameters.

#### See also

EPhysData::Subset

#### Examples

```
data(ERG)
Subset(ERG,where=list(Channel="ERG"))
#> An object of class ERGExam
#> Subject:	CR2170, 2023-05-30, Male
#> Exam Date:	 2023-08-10 11:18:31
#> Protocol:	 01-1 MoL_v2_DA long ERG [14806-F || ECN 1685 || 13 July 2021]
#> Steps:		DA 0 01 cd s m 
#> 		DA 1 cd s m 
#> 		DA 3 cd s m 
#> Eyes:		RE	LE
#> Channels:	ERG
#> 
#> An averge function has been set for this object.
#> Size: 626.7 Kb
```

#### Contents

Developed by Moritz Lindner.

Site built with pkgdown 2.0.7.
