## Supplemental Material 1 for "The ERGtools2 package: A Toolset for Processing and Analysing Visual Electrophysiology Data": UpdateChannelNames.html

Update channel names in an ERGExam object — UpdateChannelNames • ERGtools2       

Toggle navigation


ERGtools2
0.7.0

- Reference

### Update channel names in an ERGExam object

`UpdateChannelNames.Rd`

This method updates channel names in an ERGExam object. It changes the channel names
specified in the 'from' argument to the corresponding names given in the 'to' argument for the
provided ERGExam object.

```
UpdateChannelNames(X, from, to, Steps = Steps(X), Eyes = Eyes(X))
```

#### Arguments

X
:   An ERGExam

from
:   A character vector with the original channel names to be replaced.

to
:   A character vector with the new channel names to replace the original ones.

Steps
:   (Optional) A character vector specifying the steps to consider for the update.
    Defaults to all available steps in the ERGExam object.

Eyes
:   (Optional) A character vector specifying the eyes to consider for the update.
    Defaults to all available eyes in the ERGExam object.

#### Value

The updated ERGExam object with modified channel names.

#### Examples

```
if (FALSE) {
# Create an example ERGExam object
erg_data <- makeExampleERGSteps()

# Original channel names
original_names <- c("CH1", "CH2", "CH3")

# New channel names corresponding to the original names
new_names <- c("Red_Channel", "Green_Channel", "Blue_Channel")

# Update the channel names in the ERGExam object
updated_erg <- UpdateChannelNames(erg_data, from = original_names, to = new_names)
}
```

#### Contents

Developed by Moritz Lindner.

Site built with pkgdown 2.0.7.
