## Supplemental Material 1 for "The ERGtools2 package: A Toolset for Processing and Analysing Visual Electrophysiology Data": UpdateProcessingMethods.html

Set processing functions for ERGExam objects — UpdateProcessingMethods • ERGtools2       

Toggle navigation


ERGtools2
0.7.0

- Reference

### Set processing functions for ERGExam objects

`UpdateProcessingMethods.Rd`

These methods are used to set filter, rejection and averaging functions for ERGExam objects.
They allow setting the functions to a subset of recordings based on the specified conditions.

```
# S4 method for ERGExam
FilterFunction(X, where) <- value

# S4 method for ERGExam
Rejected(X, where) <- value

# S4 method for ERGExam
AverageFunction(X, where) <- value

SetStandardFunctions(
  X,
  Stimulus.type.names = pairlist(Flash = "Flash", Flicker = "Flicker")
)
```

#### Arguments

X
:   An ERGExam object

where
:   A base::list defining selection criteria. Tags/Keys in the names in the list must represent valid column names of Metadata orStimulusTable.

value
:   A value (usually a function) to set.

Stimulus.type.names
:   A base::pairlist specifying the names identifying the different stimulus types, e.g., `Flash="Flash"` or `Flash="Blitz"`.

#### Value

An updated ERGExan object.

#### Functions

- `FilterFunction(ERGExam) <- value`: Update the FilterFunction for all Recordings in an ERGExam, or only those slected using `where`.
- `Rejected(ERGExam) <- value`: Update the Rejected for all Recordings in an ERGExam, or only those slected using `where`.
- `AverageFunction(ERGExam) <- value`: Update the AverageFunction for all Recordings in an ERGExam, or only those slected using `where`.
- `SetStandardFunctions()`: This method is used to set standard functions for processing ERGExam data. It defines default functions for averaging, filtering, and signal rejection based on the stimulus type.

#### See also

EPhysData::Get\_Set\_EPhysData, EPhysMethods::autoreject.by.distance, EPhysMethods::autoreject.by.signalfree, EPhysMethods::filter.bandpass, EPhysMethods::filter.detrend,

#### Examples

```
data(ERG)
ERG<-SetStandardFunctions(ERG)
ggERGTrace(ERG,where=list(Intensity=1,Channel="ERG",Eye="RE"))
#> Retrieving record values for the given time points.
#> ================================================================================
#> Retrieving record values for the given time points.
#> ================================================================================


AverageFunction(ERG,where=list(Intensity=1))<-min
ggERGTrace(ERG,where=list(Intensity=1,Channel="ERG",Eye="RE"))
#> Retrieving record values for the given time points.
#> ================================================================================
#> Retrieving record values for the given time points.
#> ================================================================================
```

#### Contents

Developed by Moritz Lindner.

Site built with pkgdown 2.0.7.
