## Supplemental Material 1 for "The ERGtools2 package: A Toolset for Processing and Analysing Visual Electrophysiology Data": validERGMeasurements.html

Check if there is a measurement for each marker with a non-NA Relative value
in the same recording where Relative is NA — validERGMeasurements • ERGtools2       

Toggle navigation


ERGtools2
0.7.0

- Reference

### Check if there is a measurement for each marker with a non-NA Relative value in the same recording where Relative is NA

`validERGMeasurements.Rd`

Check if there is a measurement for each marker with a non-NA Relative value
in the same recording where Relative is NA

```
validERGMeasurements(object)
```

#### Contents

Developed by Moritz Lindner.

Site built with pkgdown 2.0.7.
