## Supplemental Material 1 for "The ERGtools2 package: A Toolset for Processing and Analysing Visual Electrophysiology Data": Where.html

Get index of one or several recordings and corresponding measurements — Where • ERGtools2       

Toggle navigation


ERGtools2
0.7.0

- Reference

### Get index of one or several recordings and corresponding measurements

`Where.Rd`

This method gets the index of a recording stored in an ERGExam object.

```
Where(X, where, expected.length = NULL)
```

#### Arguments

X
:   An ERGExam

where
:   A base::list defining selection criteria. Tags/Keys in the names in the list must represent valid column names of Metadata orStimulusTable.

expected.length
:   The number of elements that is expected to match the the selection criteria. This can be a helpful validity check. Ignored if NULL.

#### Value

A numeric vector of containing the index of a recording defined by the parameters, or NULL if entry was not found.

#### Details

This method gets the indices of recordings stored in an ERGExam object based on the selection citeria definde in `where`.

#### Examples

```
data(ERG)
Where(ERG,list(Channel="ERG", Intensity=1))
#> [1] 5 6
```

#### Contents

Developed by Moritz Lindner.

Site built with pkgdown 2.0.7.
