## Supplemental Material 1 for "The ERGtools2 package: A Toolset for Processing and Analysing Visual Electrophysiology Data": 0ERGMeasurements-class.R

```
validERGMeasurements <- function(object) {
  # Check if Marker slot is a data.frame with required columns
  if (!inherits(object@Marker, "data.frame") ||
      !all(c("Name", "Relative") %in% names(object@Marker))) {
    return("Marker slot must be a data.frame with columns 'Name' and 'Relative'")
  }

  # Check if Relative column contains valid values
  if (!all(object@Marker$Relative %in% c(NA, seq_len(nrow(object@Marker))))) {
    return("Relative column must contain valid row indices of 'Marker' or 'NA'")
  }

  # Check if each row's Relative value is valid
  for (i in seq_len(nrow(object@Marker))) {
    if (!is.na(object@Marker$Relative[i])) {
      if (object@Marker$Relative[i] == i) {
        return(paste0("Marker '", i,"' points to marker '", object@Marker$Name[object@Marker$Relative[i]], "' (Index: ", object@Marker$Relative[i], ") Relative must not point back onto the same marker."))
      }
    }
  }

  # Check if Measurements slot is a data.frame with required columns
  if (!inherits(object@Measurements, "data.frame") ||
      !all(c("Recording", "Marker", "Time") %in% names(object@Measurements))) {
    return(
      "Measurements slot must be a data.frame with columns 'Recording', 'Marker', and 'Time'"
    )
  }

  # Check if Marker column contains valid row indices
  valid_indices <- seq_len(nrow(object@Marker))
  if (!all(object@Measurements$Marker %in% valid_indices)) {
    return("Marker column in Measurements must contain valid row indices of Marker data frame")
  }

  # Check if Time column contains valid time units
  if (!inherits(object@Measurements$Time, "units") ||
      isFALSE(tryCatch(
        set_units(object@Measurements$Time, "s"),
        error = function(e)
          FALSE
      ))) {
    return("Time column must contain valid time units")
  }

  # check if Measurements entries are unique
  if (nrow(unique(object@Measurements[, c("Recording", "Marker")])) != nrow(object@Measurements[, c("Recording", "Marker")])) {
    return("Each Marker may only be measured once for an individual Recording.")
  }

  #' Check if there is a measurement for each marker with a non-NA Relative value
  #' in the same recording where Relative is NA
  markers <- object@Marker
  measurements <- object@Measurements
  for (i in seq_len(nrow(markers))) {
    if (!is.na(markers$Relative[i])) {
      parent <- markers$Relative[i]
      for (j in unique(measurements$Recording[measurements$Marker == i])) { # for each recording, where the current marker is set
          if (!(parent %in% measurements$Marker[measurements$Recording == j])) { # is also the parent marker in the same recording
            return(paste0(
              "No measurement for a parent marker was found for the relative marker '",
              markers$Name[i],
              "' in Recording '",
              j,
              "'."
            ))
        }
      }
      measurements[measurements$Marker == parent,]
    }
  }

  return(TRUE)
}
#' ERGMeasurements Class
#'
#' This class represents ERG (Electroretinogram) measurements, which typically consist
#' of markers placed at specific points in a recording and measurements associated with
#' each marker. Note: only the time points for the measurement is stored in the object, not the actual values. If called from a parent  \linkS4class{ERGExam}, values are retrieved based on the raw data  stored in that object.
#'
#' @slot Marker A data.frame containing marker information. This data.frame should
#'   have columns 'Name' and 'Relative'. The 'Name' column represents
#'   the name of each marker and 'Relative' column indicates the relative position of
#'   each marker with respect to other markers (or NA if no relative marker).
#'
#' @slot Measurements A data.frame containing measurement information. This data.frame
#'   should have columns 'Recording', 'Marker', and 'Time'. The 'Recording' column
#'   represents the recording number associated with each measurement, 'Marker' column
#'   indicates the marker associated with each measurement (using the row index of the
#'   marker in the Marker data.frame), and 'Time' column represents the time of each
#'   measurement.
#'
#' To create a valid ERGMeasurements object, ensure the following:
#' \itemize{
#'   \item The Marker slot is a data.frame with columns 'Name' and 'Relative'.
#'   \item The 'Name' column in the Marker data.frame contains names for each marker, which are unique within an individual channel.
#'   \item The 'Relative' column in the Marker data.frame contains valid indices of other markers from the same channel
#'     or NA if no relative marker.
#'   \item The Measurements slot is a data.frame with columns 'Recording', 'Marker', and 'Time'.
#'   \item The 'Marker' column in the Measurements data.frame contains valid row indices of the
#'     Marker data.frame.
#'   \item If the 'Marker' column points to a relative marker, there must be already a row containing a measurement from the same recording for the parent marker.
#'   \item The 'Time' column in the Measurements data.frame contains valid time units.
#' }
#' @name ERGMeasurements-class
#' @seealso \link[EPhysData:EPhysSet]{EPhysData::EPhysSet} \link{Measurements-Methods} \link{Get}
#' @importFrom units as_units
#' @examples
#' # Create marker data frame
#' marker_df <- data.frame(
#'   Name = c("a", "B", "N1", "P1"),
#'   Relative = c(NA, 1, NA, 3),
#'   ChannelBinding =c("ERG","ERG","VEP","VEP")
#' )
#'
#' # Create measurements data frame
#' measurements_df <- data.frame(
#'   Recording = c(1, 1, 2, 2, 3, 3),
#'   Marker = c(1, 2, 3, 4, 1, 2),
#'   Time = as_units(c(10, 15, 20, 25, 30, 35), "ms")
#' )
#'
#' # Create ERGMeasurements object
#' erg_obj <- new("ERGMeasurements", Marker = marker_df, Measurements = measurements_df)
#'
#' # Show the object
#' erg_obj
#'
#' # Check validity of the object
#' validObject(erg_obj)
#' @exportClass ERGMeasurements
setClass("ERGMeasurements",
         slots = list(
           Marker = "data.frame",
           Measurements = "data.frame"
         ),
         prototype = list(
           Marker = data.frame(
             Name = character(),
             Relative = numeric(),
             ChannelBinding = character()
           ),
           Measurements = data.frame(
             Recording = numeric(),
             Marker = numeric(),
             Time = as_units(integer(), "s")
           )
         ),
         validity = validERGMeasurements
)

#' @noMd
setMethod("show", "ERGMeasurements",
          function(object) {
            # Merge Marker and Measurements data frames
            merged_df <- Measurements(object)
            cat("ERGMeasurements object:\nMeasurements:\n")
            print(merged_df)
            if(any(!(1:nrow(object@Marker) %in% object@Measurements$Marker))){
              cat("\nUnused Markers:\n")
              Empty.Markers<-!(1:nrow(object@Marker) %in% object@Measurements$Marker)
              print(Markers(object)[Empty.Markers,])
            }

          })
```
