## Supplemental Material 1 for "The ERGtools2 package: A Toolset for Processing and Analysing Visual Electrophysiology Data": 0ERGProtocol-class.R

```
#' @noMd
ERGMarker <- setClass(
  "ERGMarker",
  slots = c(
    Name = "character",
    RelativeTo = "character"
  ),
  prototype = list(
    Name = NA_character_,
    RelativeTo = NA_character_
  )
)


#' @noMd
ERGChannel<-setClass(
  "ERGChannel",
  slots = c(
    Name = "character",
    Eye = "character",
    LowFreqCutoff = "character",
    HighFreqCutoff = "character",
    Inverted = "logical",
    Enabled = "logical",
    Markers = "list"
  ),
  validity = function(object) {
    markers_valid <- all(sapply(object@Markers, function(m) inherits(m, "ERGMarker")))
    eye_valid <- is.std.eyename("both")
    # marker_warnings <- sapply(object@Markers, function(m) tryCatch({
    #   validObject(m, test=T)
    #   NULL
    # }, warning = function(w) w$message, error = function(e) e$message))
    # marker_warnings <- marker_warnings[marker_warnings != ""]

    if (!markers_valid || !eye_valid) {
      msg <- paste("Validity check failed for ERGChannel object with Name:", object@Name)
      if (!markers_valid) {
        msg <- paste(msg, "\nMarkers slot should contain only objects of class 'ERGMarker'")
        # Collect warnings or errors from markers
      }
      # if (length(marker_warnings) > 0) {
      #   msg <- paste(msg, "\nMarker validity issues:", paste(marker_warnings, collapse = "; "))
      # }
      if (!eye_valid) {
        msg <- paste(msg, "\nEye slot should be 'RE', 'LE' or 'BE' but is ", object@Eye, ".")
      }
      warning(msg)
      return(FALSE)
    } else {
      return(TRUE)
    }
    if (!all(sapply(object@Markers, function(m) inherits(m, "ERGMarker")))) {
      "Markers slot should contain only objects of class 'ERGMarker'"
    } else {
      TRUE
    }
  },
  prototype = list(
    Name = character(),
    Eye = NA_character_,
    LowFreqCutoff = NA_character_,
    HighFreqCutoff = NA_character_,
    Inverted = NA,
    Enabled = NA,
    Markers = list()
  )
)

#' @noMd
ERGStep<-setClass(
  "ERGStep",
  slots = c(
    Description = "character",
    Adaptation = "character",
    Resultsperrun = "numeric",
    Timebetweenresults = "character",
    SampleFrequency = "character",
    PreTriggerTime = "character",
    PostTriggerTime = "character",
    InterSweepDelay = "character",
    Sweepsperresult = "numeric",
    Sweepsperresultmin = "numeric",
    Glitchremoval = "logical",
    Driftremoval = "logical",
    Baselineenabled = "logical",
    Baselinepretrigger = "character",
    Baselinerange = "character",
    Channels = "list"  # Slot for a list of 'ERGChannel' objects
  ),
  validity = function(object) {
    channels_valid <- all(sapply(object@Channels, function(c) inherits(c, "ERGChannel")))
    description_valid <- nchar(object@Description) > 0
    # Collect warnings or errors from channels
    channel_warnings <- sapply(object@Channels, function(c) tryCatch({
      validObject(c,test=T)
      ""
    }, warning = function(w) w$message, error = function(e) e$message))
    channel_warnings <- channel_warnings[channel_warnings != ""]

    if (!channels_valid || !description_valid || length(channel_warnings) > 0) {
      msg <- paste("Validity check failed for ERGStep object with Description:", object@Description)
      if (!channels_valid) {
        msg <- paste(msg, "\nChannels slot should contain only objects of class 'ERGChannel'")
      }
      if (length(channel_warnings) > 0) {
        msg <- paste(msg, "\nChannel validity issues:", paste(channel_warnings, collapse = "; "))
      }
      if (!description_valid) {
        msg <- paste(msg, "\nDescription slot should not be empty")
      }
      warning(msg)
      return(FALSE)
    } else {
      return(TRUE)
    }
  },
  prototype = list(
    Description = "",
    Adaptation = "",
    Resultsperrun = -1,
    Timebetweenresults = "",
    SampleFrequency = "",
    PreTriggerTime = "",
    PostTriggerTime = "",
    InterSweepDelay = "",
    Sweepsperresult = -1,
    Sweepsperresultmin = -1,
    Glitchremoval = FALSE,
    Driftremoval = FALSE,
    Baselineenabled = FALSE,
    Baselinepretrigger = "",
    Baselinerange = "",
    Channels = list()  # Empty list as a prototype for 'ERGChannels'
  )
)


#' ERGProtocol class definition
#'
#' This class represents an ERG protocol, as it can be impored from Diagnosys Espion™ software with Export_Date, Name, nSteps, nChannels, and Step slots.
#'
#' @slot Export_Date POSIXct slot for the export date and time.
#' @slot Name Character slot for the protocol name.
#' @slot nSteps Numeric slot for the number of steps.
#' @slot nChannels Numeric slot for the number of channels.
#' @slot Step List slot for a list of 'ERGStep' objects.
#' @name ERGProtocol
#' @rdname ERGProtocol
ERGProtocol<-setClass(
  "ERGProtocol",
  slots = c(
    Export_Date = "POSIXct",
    Name = "character",
    nSteps = "numeric",
    nChannels = "numeric",
    Step = "list"  # Slot for a list of 'Channel' objects
  ),
  validity = function(object) {
    steps_valid <- all(sapply(object@Step, function(c) inherits(c, "ERGStep")))
    name_valid <- nchar(object@Name) > 0
    # Collect warnings or errors from steps
    step_warnings <- sapply(object@Step, function(c) tryCatch({
      validObject(c,test=T)
      ""
    }, warning = function(w) w$message, error = function(e) e$message))
    step_warnings <- step_warnings[step_warnings != ""]

    if (!steps_valid || !name_valid || length(step_warnings) > 0) {
      msg <- paste("Validity check failed for ERGProtocol object with Name:", object@Name)
      if (!steps_valid) {
        msg <- paste(msg, "\nStep slot should contain only objects of class 'ERGStep'")
      }
      if (length(step_warnings) > 0) {
        msg <- paste(msg, "\nStep validity issues:", paste(step_warnings, collapse = "; "))
        message(step_warnings)
      }
      if (!name_valid) {
        msg <- paste(msg, "\nName slot should not be empty")
      }
      warning(msg)
      return(FALSE)
    } else {
      return(TRUE)
    }
  },
  prototype = list(
    Export_Date = as.POSIXct(NA),
    Name = NA_character_,
    nSteps = NA_real_,
    nChannels = NA_real_,
    Step = list()  # Empty list as a prototype for 'ERGStep'
  )
)

#' Show method for ERGProtocol class
#'
#' @param object An instance of the ERGProtocol class.
#' @param ... Additional arguments (not used).
#' @name show.ERGProtocol
#' @noMd
setMethod("show", signature = "ERGProtocol", function(object) {
  cat("ERGProtocol Object\n")
  cat("Export Date: ", object@Export_Date, "\n")
  cat("Name: ", object@Name, "\n")
  cat("Number of Steps: ", object@nSteps, "\n")
  cat("Number of Channels: ", object@nChannels, "\n")
  for (i in seq_along(object@Step)) {
    cat("Step ", i, ":\n")
    cat("\tDescription: ", object@Step[[i]]@Description, "\n")
    cat("\tAdaptation: ", object@Step[[i]]@Adaptation, "\n")
    cat("\tSample Frequency: ", object@Step[[i]]@SampleFrequency, "\n")
    cat("\tPre-Trigger Time: ", object@Step[[i]]@PreTriggerTime, "\n")
    cat("\tPost-Trigger Time: ", object@Step[[i]]@PostTriggerTime, "\n")
    cat("\tInter-Sweep Delay: ", object@Step[[i]]@InterSweepDelay, "\n")
    for (j in seq_along(object@Step[[i]]@Channels)) {
      cat("\tChannel ", j, ":\n")
      cat("\t\tName: ", object@Step[[i]]@Channels[[j]]@Name, "\n")
      cat("\t\tEye: ", object@Step[[i]]@Channels[[j]]@Eye, "\n")
      cat("\t\tLow Freq Cutoff: ", object@Step[[i]]@Channels[[j]]@LowFreqCutoff, "\n")
      cat("\t\tHigh Freq Cutoff: ", object@Step[[i]]@Channels[[j]]@HighFreqCutoff, "\n")
      cat("\t\tInverted: ", object@Step[[i]]@Channels[[j]]@Inverted, "\n")
      cat("\t\tEnabled: ", object@Step[[i]]@Channels[[j]]@Enabled, "\n")
      for (k in seq_along(object@Step[[i]]@Channels[[j]]@Markers)) {
        cat("\tMarker ", k, ":\n")
        cat("\t\t\tName: ", object@Step[[i]]@Channels[[j]]@Markers[[k]]@Name, "\n")
        cat("\t\t\tRelativeTo: ", object@Step[[i]]@Channels[[j]]@Markers[[k]]@RelativeTo, "\n")

      }
    }
  }
})

setMethod("show", signature = "ERGProtocol", function(object) {

  print_hierarchical_list <- function(hierarchical_list, level = 0) {
    for (i in 1:length(hierarchical_list)) {
      item<-hierarchical_list[[i]]
      name<-names(hierarchical_list)[[i]]
      if (is.list(item)) {
        # If the item is a list, recursively print its contents with increased indentation.
        cat(rep("\t", level), name, ": ",i, "\n", sep = "")
        print_hierarchical_list(item, level + 1)
      } else {
        # If the item is not a list, print it with the current indentation level.
        cat(rep("\t", level), name, ": ", item, "\n", sep = " ")
      }
    }
  }

  moveRedundant <- function(list) {
    if (length(list) == 1) {
      # If there's only one item in the list, return that item
      return(list[[1]])
    } else {
      # Check if all items in the list are identical
      is_redundant <- all(unlist(lapply(list, identical, list[[1]])))
      if (is_redundant) {
        # If all items are identical, return the first item
        return(list[[1]])
      } else {
        # If not all items are identical, return the list as is
        return(list)
      }
    }
  }

  protocol_list <- list(
    Export_Date = object@Export_Date,
    Name = object@Name,
    nSteps = object@nSteps,
    nChannels = object@nChannels,
    Steps = list()
  )

  for (i in seq_along(object@Step)) {
    step_list <- list(
      Description = object@Step[[i]]@Description,
      Adaptation = object@Step[[i]]@Adaptation,
      SampleFrequency = object@Step[[i]]@SampleFrequency,
      PreTriggerTime = object@Step[[i]]@PreTriggerTime,
      PostTriggerTime = object@Step[[i]]@PostTriggerTime,
      InterSweepDelay = object@Step[[i]]@InterSweepDelay,
      Channels = list()
    )

    for (j in seq_along(object@Step[[i]]@Channels)) {
      channel_list <- list(
        Name = object@Step[[i]]@Channels[[j]]@Name,
        Eye = object@Step[[i]]@Channels[[j]]@Eye,
        LowFreqCutoff = object@Step[[i]]@Channels[[j]]@LowFreqCutoff,
        HighFreqCutoff = object@Step[[i]]@Channels[[j]]@HighFreqCutoff,
        Inverted = object@Step[[i]]@Channels[[j]]@Inverted,
        Enabled = object@Step[[i]]@Channels[[j]]@Enabled,
        Markers = list()
      )

      for (k in seq_along(object@Step[[i]]@Channels[[j]]@Markers)) {
        marker_list <- list(
          Name = object@Step[[i]]@Channels[[j]]@Markers[[k]]@Name,
          RelativeTo = object@Step[[i]]@Channels[[j]]@Markers[[k]]@RelativeTo
        )
        channel_list$Markers[[k]] <- marker_list
      }

      step_list$Channels[[j]] <- channel_list
    }
    #step_list <- lapply(step_list, moveRedundant)
    protocol_list$Steps[[i]] <- step_list
  }

  #protocol_list <- lapply(protocol_list, moveRedundant)
  #protocol_list
  print_hierarchical_list(protocol_list)
})
```
