## Supplemental Material 1 for "The ERGtools2 package: A Toolset for Processing and Analysing Visual Electrophysiology Data": 1ERGExam-class.R

```
#' Check the validity of an ERGExam object
#'
#' This function validates the structure and integrity of an \code{ERGExam} object by ensuring that all required fields
#' are present, correctly formatted, and contain appropriate values. It checks the validity of subject information,
#' metadata, and measurements, ensuring consistency across the various components of the object.
#'
#' @param object An \code{ERGExam} object to validate.
#' @param non.fatal A string specifying the severity level for certain notices. Can be "Error", "Warning", or "Information".
#'   Defaults to "Error".
#'
#' @details
#' This function performs the following checks:
#' \itemize{
#'   \item Ensures essential subject identifiers (e.g., subject name, date of birth) and exam information (e.g., exam date) are present.
#'   \item Verifies that the \code{Metadata} slot contains required columns: "Step", "Eye", "Channel", and "Repeat". It also checks their class types.
#'   \item Ensures that the \code{Metadata} slot does not contain any missing values in its essential columns and that reserved column names are not used.
#'   \item Checks the validity of \code{Eye} entries in the \code{Metadata} slot against predefined valid identifiers.
#'   \item Ensures that data consistency is maintained across steps, eyes, and repeats in the \code{Data} slot for certain fields like "Rejected", "filter.fx", and "average.fx".
#'   \item Validates the \code{Measurements} slot to ensure it contains a valid \code{ERGMeasurements} object and that channel binding matches the metadata.
#'   \item Checks the \code{Stimulus} slot for required columns and the presence of valid data in essential columns like "Step", "Description", and "StimulusEnergy".
#'   \item Ensures temporal consistency among the \code{DOB}, \code{ExamDate}, and \code{Imported} dates.
#' }
#'
#' The function generates \code{Notice} messages if any issues are found, which could include errors, warnings, or information based on the provided \code{non.fatal} argument.
#'
#' @return Returns \code{TRUE} if the \code{ERGExam} object passes all validation checks. Otherwise, it issues \code{Notice} messages with specific instructions for resolving issues.
#'
#' @examples
#' \dontrun{
#' validERGExam(myERGExam)
#' }
#'
#' @importFrom EPhysData Rejected FilterFunction AverageFunction
#' @importFrom stats var
#' @importFrom cli cli_abort cli_inform cli_alert_warning cli_warn
#' @export
validERGExam <- function(object, non.fatal = "Error") {

  # essential identifiers
  #Essential fields
  if (!(
    nzchar(object@SubjectInfo$Subject) &&
    !is.na(object@SubjectInfo$DOB) &&
    all(!is.na(object@ExamInfo$ExamDate))
  )) {
    cli_abort("x Subject Name, DOB and ExamDate must be provided.")
  }

  # Check if the metadata has the required columns and are of correct format
  required_columns <- c("Step", "Eye", "Channel","Repeat")
  if (!all(required_columns %in% names(object@Metadata))) {
    missing_columns <- setdiff(required_columns, names(object@Metadata))
    Notice(object,
            what = c("Error"),
            notice_text = c("i The metadata must be a data.frame with the columns: 'Step', 'Eye', 'Channel', and 'Repeat'.",
                            "x The following required columns are missing: {missing_columns}."),
            help_page = "ERGtools2::ERGExam")
  }
  if (!(any(c("integer", "numeric") %in% class(object@Metadata$Step))) ||
      !(any(c("integer", "numeric") %in% class(object@Metadata$Repeat)))) {
    Notice(object,
           what = c("Error"),
           notice_text = c("x Metadata columns 'Step' and 'Repeat' must be of class 'integer'.",
                           "i However, they are currently of class:",
                           "i 'Step': {class(object@Metadata$Step)}",
                           "i 'Repeat': {class(object@Metadata$Repeat)}",
                           "x Please ensure both columns are of class 'integer' before proceeding."),
           help_page = "ERGtools2::ERGExam")
  }
  if (!("character" %in% class(object@Metadata$Channel)) ||
      !("character" %in% class(object@Metadata$Eye))) {
    Notice(object,
           what = c("Error"),
           notice_text = c("x Metadata columns 'Channel' and 'Eye' must be of class 'character'.",
                           "i However, they are currently of class:",
                           "i 'Channel': {class(object@Metadata$Channel)}",
                           "i 'Eye': {class(object@Metadata$Eye)}",
                           "x Please ensure both columns are of class 'character' before proceeding."),
           help_page = "ERGtools2::ERGExam")
  }

  missing_indices <- which(is.na(object@Metadata[, c("Step", "Eye", "Channel", "Repeat")]), arr.ind = TRUE)
  missing_info <- apply(missing_indices, 1, function(index) {
    paste0("Row ", index[1], ", Column '", colnames(object@Metadata)[index[2]], "'")
  })

  if (length(missing_indices) > 0) {
    Notice(object,
           what = c(non.fatal),
           notice_text = c("! The essential columns of the Metadata slot ('Step', 'Eye', 'Channel', 'Repeat') contain missing values.",
                           "i Missing values detected at: {paste(missing_info, collapse = '; ')}.",
                           "! Run {.run Metadata({deparse(substitute(x))})} to identify and address these missing values in order to avoid potential issues in downstream methods."),
           help_page = "ERGtools2::ERGExam")
  }

  reserved_columns <- c(
    "Description", "StimulusEnergy", "Background", "Type", "Name",
    "ChannelBinding", "Relative", "Time"
  )

  used_reserved_columns <- intersect(colnames(object@Metadata), reserved_columns)

  if (length(used_reserved_columns) > 0) {
    Notice(object,
           what = c(non.fatal),
           notice_text = c("! The following reserved column names are being used in 'Metadata': {paste(used_reserved_columns, collapse = ', ')}.",
                           "i 'Description', 'StimulusEnergy', 'Background', 'Type', 'Name', 'ChannelBinding', 'Relative', and 'Time' are reserved column names.",
                           "! These names should not be used as extra column names in 'Metadata' to avoid potential conflicts."),
           help_page = "ERGtools2::ERGExam")
  }

  # Check if Eye entries valid
  invalid_eyes <- setdiff(unique(object@Metadata$Eye), eye.haystack())

  if (length(invalid_eyes) > 0) {
    Notice(object,
           what = c("Error"),
           notice_text = c("x Eye identifiers in the 'Metadata' slot are invalid.",
                           "i Valid identifiers must match any of the values returned by `eye.haystack()`.",
                           "x The following invalid eye identifiers were found: {paste(invalid_eyes, collapse = ', ')}."),
           help_page = "ERGtools2::ERGExam")

  }

  # Rejected (in EPhysRaw) - must be the same across all channels within one step and eye
  for (s in unique(object@Metadata$Step)) {
    for (e in unique(object@Metadata$Eye[object@Metadata$Step == s])) {
      for (r in unique(object@Metadata$Repeat[object@Metadata$Step == s &
                                              object@Metadata$Eye == e])) {
        ids.equal <-
          object@Metadata$Step == s & object@Metadata$Eye == e & object@Metadata$Repeat == r
        feature.list <- lapply(object@Data[ids.equal], Rejected)
        if (length(unique(unlist(lapply(
          feature.list, length
        )))) != 1) {
          Notice(object,
                 what = c("E"),
                 where = list(Step = s, Eye = e, Repeat = r),
                 notice_text = c("x Unequal amount of recordings for the different channels."),
                 help_page = "ERGtools2::ERGExam")
        }
        feature.df <- tryCatch(
          as.data.frame(do.call(cbind, feature.list)),
          error = function(er) {
            Notice(object,
                   what = c("E"),
                   where = list(Step = s, Eye = e, Repeat = r),
                   notice_text = c("x 'Rejected' slots are filled with vectors of unequal length."),
                   help_page = "ERGtools2::ERGExam")
          }
        )
        if (!all(apply(feature.df, 1, function(x) {
          if (length(x) > 1) {
            var(x) == 0
          } else{
            TRUE
          }
        }))) {
          Notice(object,
                 what = c("W"), #this will be replaced by "non.fatal" in future
                 where = list(Step = s, Eye = e, Repeat = r),
                 notice_text = c("! 'Rejected' vectors are not identical across channels."),
                 help_page = "ERGtools2::ERGExam")
        }
      }
    }
  }

  # filter.fx (inEPhysRaw) - must be the same across all eyes within one channel and step
  for (s in unique(object@Metadata$Step)) {
    for (c in unique(object@Metadata$Channel[object@Metadata$Step == s])) {
      ids.equal <-
        which(object@Metadata$Step == s &
                object@Metadata$Channel == c)
      feature.list <-
        lapply(object@Data[ids.equal], function(x) {
          deparse1(FilterFunction(x))
        })
      feature.df <- tryCatch(
        as.data.frame(do.call(cbind, feature.list)),
        error = function(e) {
          Notice(object,
                 what = c("E"),
                 where = list(Step = s, Eye = e, Repeat = r),
                 notice_text = c("x 'filter.fx' slots are filled with vectors of unequal length. "),
                 help_page = "ERGtools2::ERGExam")
        }
      )
      if (!all(apply(feature.df, 1, function(x) {
        length(unique(x)) == 1
      }))) {
        Notice(object,
               what = c("E"),
               where = list(Step = s, Eye = e, Repeat = r),
               notice_text = c("x 'filter.fx' vectors are not identical across both eyes."),
               help_page = "ERGtools2::ERGExam")
      }
    }
  }

  # average.fx (inEPhysRaw) - must be the same across all eyes within one channel and step
  for (s in unique(object@Metadata$Step)) {
    for (c in unique(object@Metadata$Channel[object@Metadata$Step == s])) {
      ids.equal <-
        which(object@Metadata$Step == s &
                object@Metadata$Channel == c)
      feature.list <-
        lapply(object@Data[ids.equal], function(x) {
          deparse1(AverageFunction(x))
        })
      feature.df <- tryCatch(
        as.data.frame(do.call(cbind, feature.list)),
        error = function(e) {
          Notice(object,
                 what = c("E"),
                 where = list(Step = s, Eye = e, Repeat = r),
                 notice_text = c("x average.fx' slots are filled with vectors of unequal length."),
                 help_page = "ERGtools2::ERGExam")
        }
      )
      if (!all(apply(feature.df, 1, function(x) {
        length(unique(x)) == 1
      }))) {
        Notice(object,
               what = c("E"),
               where = list(Step = s, Eye = e, Repeat = r),
               notice_text = c("x 'average.fx' vectors are not identical across both eyes."),
               help_page = "ERGtools2::ERGExam")
      }
    }
  }

  # Averaged slot
  if (object@Averaged){
    Trials<-unique(unlist(lapply(object@Data,function(x){dim(x)[2]})))
    if(length(Trials)!=1 || Trials != 1){
      Notice(object,
             what = c("I"),
             notice_text = c("i Object does not appear to contain averaged data. Multiple repeats observed. This message is likely irrelevant, as the 'Averaged' is obsolete."),
             help_page = "ERGtools2::ERGExam")
    }
  }

  # Measurements slot
  if (length(object@Measurements) != 0) {
    if (any(!("ERGMeasurements" %in% class(object@Measurements)),
            !validObject(object@Measurements))) {
      Notice(object,
             what = c("E"),
             notice_text = c("x 'Measurements' slot must contain a valid ERGMeasurements object."),
             help_page = "ERGtools2::ERGMeasurements-class")
    }

    # check if ChannelBinding matches Channel of Recording
    measurements<-Measurements(object@Measurements)
    for (r in unique(measurements$Recording)){
      cb<-unique(measurements$ChannelBinding[measurements$Recording ==r])
      if(length(cb)>1){
        Notice(object,
               what = c("E"),
               notice_text = c("x Measurements object malformed: Multiple Channel Bindings detected for recording {.val {r}}.",
                               "i The detected Channel Bindings are: {.val {cb}}.",
                               "i Expected exactly one Channel Binding per recording."),
               help_page = "ERGtools2::ERGMeasurements-class")
      }
      expected_channel <- Metadata(object)$Channel[r]
      if (expected_channel != cb) {
        Notice(object,
               what = c("E"),
               notice_text = c("x Channel mismatch detected: The Channel Binding stored in the Measurements slot ({.val {cb}}) does not match the channel of recording {.val {r}} as stored in the parent object ({.val {expected_channel}}).",
                               "i Please check the consistency between the Measurements slot and the parent object's metadata."),
               help_page = "ERGtools2::ERGMeasurements-class")
      }
    }
  }

  # Stimulus slot

  stimulus_steps <- object@Stimulus$Step
  metadata_steps <- unique(Metadata(object)$Step)
  if (!all(stimulus_steps %in% metadata_steps)) {
    # Identify which Step entries in Stimulus are missing from Metadata
    missing_steps <- stimulus_steps[!stimulus_steps %in% metadata_steps]
    Notice(
      object,
      what = c("E"),
      notice_text = c(
        "x Mismatch between Stimulus and Metadata Steps detected.",
        "i Step entries in Stimulus table are: {.val {stimulus_steps}}.",
        "i Steps contained in Metadata are: {.val {metadata_steps}}."
      ),
      help_page = "ERGtools2::ERGExam"
    )

    if (length(missing_steps) > 0) {
      Notice(
        object,
        what = c(non.fatal),
        notice_text = c(
          "! The following Steps in the Stimulus table are missing from Metadata: {.val {missing_steps}}."
        ),
        help_page = "ERGtools2::ERGExam"
      )
    }
    Notice(
      object,
      what = c("E"),
      notice_text = c(
        "x All stimuli described must correspond to a Step as defined in 'Metadata'."
      ),
      help_page = "ERGtools2::ERGExam"
    )
  }

  essential_columns <- c("Step", "Description", "StimulusEnergy", "Background", "Type")

  if (!all(essential_columns %in% colnames(object@Stimulus))) {
   missing<-colnames(object@Stimulus)[!(essential_columns %in% colnames(object@Stimulus))]
   Notice(
     object,
     what = c("E"),
     notice_text = c(
       "x Data.frame in the Stimulus slot does not contain all necessary columns.",
       "i The essential columns of the Stimulus slot are: ('Step', 'Description', 'StimulusEnergy', 'Background', 'Type').",
       "i The following column is missing: {.val {missing}}."
     ),
     help_page = "ERGtools2::ERGExam"
   )
  }

  missing_values <- is.na(object@Stimulus[, essential_columns])
  if (any(missing_values)) {
    columns_with_na <- essential_columns[apply(missing_values, 2, any)]
    rows_with_na <- which(rowSums(missing_values) > 0)
    Notice(
      object,
      what = c(non.fatal),
      notice_text = c(
        "x Missing values detected in essential columns of the Stimulus slot.",
        "i The essential columns of the Stimulus slot ('Step', 'Description', 'StimulusEnergy', 'Background', 'Type') should not contain missing values.",
        "i The following columns contain missing values: {.val {columns_with_na}}.",
        "i Missing values were found in the following rows: {.val {rows_with_na}}.",
        "i Use '{.run Stimulus({deparse(substitute(x))})}' to check and adjust manually to ensure downstream methods won't fail."
      ),
      help_page = "ERGtools2::ERGExam"
    )
  }

  reserved_columns <- c(
    "Channel", "Repeat", "Eye", "Name", "ChannelBinding", "Relative", "Time"
  )
  used_reserved_columns <- colnames(object@Stimulus)[colnames(object@Stimulus) %in% reserved_columns]
  if (length(used_reserved_columns) > 0) {
    Notice(
      object,
      what = c(non.fatal),
      notice_text = c(
        "! Reserved column names detected in the Stimulus slot. These are: {.val {used_reserved_columns}}",
        "i 'Channel', 'Repeat', 'Eye', 'Name', 'ChannelBinding', 'Relative', and 'Time' are reserved column names.",
        "i These names should not be used as extra column names in 'Stimulus'. Please rename them to avoid conflicts."
      ),
      help_page = "ERGtools2::ERGExam"
    )
  }

  if (!("integer" %in% class(object@Stimulus$Step)) ||
      !(any(c("numeric","integer") %in% class(object@Stimulus$StimulusEnergy)))) {
    Notice(
      object,
      what = c("E"),
      notice_text = c(
        "i Stimulus slot columns 'Step' and 'StimulusEnergy' must be of class 'integer' and 'numeric' or 'integer', respectivley. They are: {.val {class(object@Stimulus$Step)}} and {.val {class(object@Stimulus$StimulusEnergy)}}."
      ),
      help_page = "ERGtools2::ERGExam"
    )
  }

  if (!("character" %in% class(object@Stimulus$Description)) ||
      !("character" %in% class(object@Stimulus$Background)) ||
    !("character" %in% class(object@Stimulus$Type))) {
    Notice(
      object,
      what = c("E"),
      notice_text = c(
        "i Stimulus slot columns 'Description', 'Background' and 'Type' must be of class 'character'. They are: {.val {class(object@Stimulus$Description)}}, {.val {class(object@Stimulus$Background)}} and {.val {class(object@Stimulus$Type)}}, respectivley."
      ),
      help_page = "ERGtools2::ERGExam"
    )
  }

  # Dates
  dob <- object@SubjectInfo$DOB
  exam_date <- object@ExamInfo$ExamDate
  imported_date <- object@Imported

  if (("Date" %in% class(dob)) &&
      ("POSIXct" %in% class(exam_date)) &&
      ("POSIXct" %in% class(imported_date))) {
    if (!(as.POSIXct(dob) < min(exam_date) &&
          max(exam_date) < min(imported_date))) {
      Notice(
        object,
        what = c(non.fatal),
        notice_text = c(
          "i Temporal sequence of 'DOB' ({.val {dob}}), 'ExamDate' ({.val {exam_date}}), and 'Imported' ({.val {imported_date}}) is impossible."
        ),
        help_page = "ERGtools2::ERGExam"
      )
    }
  } else {
    Notice(
      object,
      what = c("E"),
      notice_text = c(
        "i DOB, ExamDate, and Imported should be of class Date, POSIXct, and POSIXct. They are [{.val {class(dob)}}], [{.val {class(exam_date)}}], and [{.val {class(imported_date)}}], respectively."
      ),
      help_page = "ERGtools2::ERGExam"
    )
  }

  if(object@Imported<"2024-02-01"){
    Notice(
      object,
      what = c("I"),
      notice_text = c(
        "i This ERGExam object is outdated. Consider re-importing in case of errors." #Please run {.run UpdateERGExam({deparse(substitute(x))})}."
      )
    )
  }
  TRUE
}

#' ERGExam Class
#'
#' A class representing an ERG (Electroretinogram) exam. This class extends the \link[EPhysData:EPhysSet]{EPhysData::EPhysSet} object and all methods valid for \link[EPhysData:EPhysSet]{EPhysData::EPhysSet} can also be applied to \linkS4class{ERGExam} objects.
#'
#' @slot Data Data A list of \link[EPhysData:EPhysData]{EPhysData::EPhysData} objects. Each item containing a recording in response to a particular Stimulus ("Step"), from a particular eye and data Channel (e.g. ERG or OP), as defined in the corresponding Metadata.
#' @slot Metadata  A data frame containing metadata information associated with the data in the Data slot, each row corresponds to one item in \code{data}.
#' \describe{
#'   \item{Step}{An integer vector containing the step index. A step describes data recorded in response to the same type of stimulus. This column links the Stimulus slot to the metadata}
#'   \item{Eye}{A character vector. Possible standard values "RE" (right eye) , "LE" (left eye), and "BE" (bhot eyes). Possible non-standard values can be listed running \code{od_str}, \code{os_str} and \code{ou_str}, resp.. See: \link[=as.std.eyename]{as.std.eyename}.}
#'   \item{Channel}{A character vector containing the channel name. This can be "ERG", "VEP" or "OP" for instance.}
#'   \item{Repeat}{A numeric vector containing the indices of individual repeats contained in an ERG exam. E.g., if a recording to one identical stimulus is performed twice, these would be distinguished by different indices in the 'repeat' column. Warning: This is currently experimental}
#' }
#'
#' @slot Stimulus
#' A data frame containing stimulus information.
#' \describe{
#'   \item{Step}{An integer vector row index. Will be removed in future versions.}
#'   \item{Description}{A character vector describing the stimulus in a human-readable way.}
#'   \item{StimulusEnergy}{A numeric vector representing the energy of the stimulus.}
#'   \item{Background}{A character vector describing the adaptation state of the retina for that stimulus (DA or LA).}
#'   \item{Type}{A character vector describing the type of the stimulus (e.g. Flash or Flicker).}
#' }
#'
#' @slot Averaged
#' TRUE if the object contains averaged data, FALES indicates object contains raw traces.
#'
#' @slot Measurements
#' An object of class \linkS4class{ERGMeasurements}
#'
#' @slot ExamInfo
#' A list containing exam-related information.
#' \describe{
#'   \item{ProtocolName}{A character vector indicating the name of the protocol.}
#'   \item{Version}{Optional: A character vector indicating the version of the protocol.}
#'   \item{ExamDate}{A \code{POSIXct} The date of the exam.}
#'   \item{Filename}{Optional: The filename associated where exam raw data have been imported from.}
#'   \item{RecMode}{Optional: A character vector indicating the recording mode.}
#'   \item{Investigator}{Optional: A character vector indicating the name of the investigator conducting the exam.}
#' }
#'
#' @slot SubjectInfo
#' A list containing subject-related information.
#' \describe{
#'   \item{Subject}{A character vector indicating the name of the subject}
#'   \item{DOB}{A \code{Date} object indicating the date of birth of the subject}
#'   \item{Gender}{Optional: A character vector indicating the gender of the subject}
#'   \item{Group}{Optional: A character vector indicating the study group to which the subject belongs.}
#' }
#'
#' @slot Imported
#' A \code{POSIXct} timestamp indicating when the object was imported.
#'
#' @name ERGExam
#' @seealso \link[EPhysData:EPhysData]{EPhysData::EPhysData-package} \link[EPhysData:EPhysData]{EPhysData::EPhysData-class} \link[EPhysData:EPhysSet]{EPhysData::EPhysSet-class}
#' @importClassesFrom EPhysData EPhysData EPhysSet
#' @importFrom units as_units
#' @aliases ERGExam-class
#' @exportClass ERGExam
ERGExam <- setClass(
  "ERGExam",
  contains = "EPhysSet",
  slots = c(
    Data = "list",
    # Data is a list of EPhysData objects
    Metadata = "data.frame",
    # Metadata is a data frame
    Stimulus = "data.frame",
    # Stimulus is a data frame
    Averaged = "logical",
    # Averaged is a logical
    Measurements = "ERGMeasurements",
    # Measurements is a data frame
    ExamInfo = "list",
    # ExamInfo is a list
    SubjectInfo = "list",
    # SubjectInfo is a list
    Imported = "POSIXct"
  ),
  prototype = list(
    Data = list(NULL),
    Metadata = data.frame(
      Step = integer(),
      Eye = character(),
      Channel = character(),
      stringsAsFactors = FALSE
    ),
    Stimulus = data.frame(
      Step = integer(),
      Description = character(),
      StimulusEnergy = as_units(numeric(), unitless),
      Background = character(),
      Type = character()
    ),
    Averaged = FALSE,
    Measurements =  new("ERGMeasurements"),
    ExamInfo = list(
      ProtocolName = character(),
      Version = character(),
      ExamDate = as.POSIXct(integer()),
      Filename = character(),
      RecMode = character(),
      Investigator = character()
    ),
    SubjectInfo = list(
      Subject = character(),
      DOB = as.Date(x = integer(0), origin = "1970-01-01"),
      Gender = character(),
      Group = character()
    ),
    Imported = as.POSIXct(integer())
  ),
  validity = function(object) {
    validERGExam(object)
  }
)

#' Create an instance of the ERGExam class
#'
#' @description This function creates an instance of the \linkS4class{ERGExam} class.
#'
#' @param Data Data A list of \link[EPhysData:EPhysData]{EPhysData::EPhysData} objects. Each item containing a recording in response to a particular Stimulus ("Step"), from a particular eye and data Channel (e.g. ERG or OP), as defined in the corresponding Metadata.
#' @param Metadata  A data frame containing metadata information associated with the data in the Data slot, each row corresponds to one item in \code{data}.
#' \describe{
#'   \item{Step}{An integer vector containing the step index. A step describes data recorded in response to the same type of stimulus. This column links the Stimulus slot to the metadata}
#'   \item{Eye}{A character vector. Possible standard values "RE" (right eye) , "LE" (left eye), and "BE" (bhot eyes). Possible non-standard values can be listed running \code{od_str}, \code{os_str} and \code{ou_str}, resp.. See: \link[=as.std.eyename]{as.std.eyename}.}
#'   \item{Channel}{A character vector containing the channel name. This can be "ERG", "VEP" or "OP" for instance.}
#'   \item{Repeat}{A numeric vector containing the indices of individual repeats contained in an ERG exam. E.g., if a recording to one identical stimulus is performed twice, these would be distinguished by different indices in the 'repeat' column. Warning: This is currently experimental}
#' }
#' @param Stimulus A data frame containing stimulus information.
#' \describe{
#'   \item{Step}{An integer vector row index. Will be removed in future versions.}
#'   \item{Description}{A character vector describing the stimulus in a human-readable way.}
#'   \item{StimulusEnergy}{A numeric vector representing the energy of the stimulus.}
#'   \item{Background}{A character vector describing the adaptation state of the retina for that stimulus (DA or LA).}
#'   \item{Type}{A character vector describing the type of the stimulus (e.g. Flash or Flicker).}
#' }
#' @param Averaged TRUE if the object was created from averaged data, FALES indicates object contains raw traces.
#' @param Measurements An object of class \linkS4class{ERGMeasurements}.
#' @param ExamInfo A list containing exam-related information.
#' \describe{
#'   \item{ProtocolName}{A character vector indicating the name of the protocol.}
#'   \item{Version}{Optional: A character vector indicating the version of the protocol.}
#'   \item{ExamDate}{A \code{POSIXct} The date of the exam.}
#'   \item{Filename}{Optional: The filename associated where exam raw data have been imported from.}
#'   \item{RecMode}{Optional: A character vector indicating the recording mode.}
#'   \item{Investigator}{Optional: A character vector indicating the name of the investigator conducting the exam.}
#' }
#' @param SubjectInfo A list containing subject-related information.
#' \describe{
#'   \item{Subject}{A character vector indicating the name of the subject}
#'   \item{DOB}{A \code{Date} object indicating the date of birth of the subject}
#'   \item{Gender}{Optional: A character vector indicating the gender of the subject}
#'   \item{Group}{Optional: A character vector indicating the study group to which the subject belongs.}
#' }
#' @param skip.validation Do not test if object is valid. Default is \code{FALSE}. This can be helpful when creating import routines that might sometimes be called on incomplete/faulty datasets.
#' @examples
#' # Create example data and metadata
#' Data <-
#'   list(
#'     makeExampleEPhysData(sample_points = 100, replicate_count = 3),
#'     makeExampleEPhysData(sample_points = 100, replicate_count = 3),
#'     makeExampleEPhysData(sample_points = 100, replicate_count = 6),
#'     makeExampleEPhysData(sample_points = 100, replicate_count = 6)
#'   )  # List of EPhysData objects
#' Metadata <-
#'   data.frame(
#'     Step = as.integer(c(1,1,2,2)),
#'     Eye = c("RE", "LE", "LE", "LE"),
#'     Channel = c("Ch1", "Ch1", "Ch1", "Ch2"),
#'     Repeat = as.integer(c(1,1,1,1))
#'   )
#' Stimulus <-
#'   data.frame(
#'              Step = as.integer(c(1,1,2,2)),
#'              Description = c("Stim1", "Stim2"),
#'              StimulusEnergy = as.numeric(c(1,10)),
#'              Background = c("DA","DA"),
#'              Type = c("Flash","Flash"))  # Example stimulus data
#' ExamInfo <-
#'   list(
#'     ProtocolName = "ERG Protocol",
#'     Version = "1.0",
#'     ExamDate = as.POSIXct("2023-08-14"),
#'     Filename = "exam_data.csv"
#'   )
#' SubjectInfo <-
#'   list(
#'     Subject = "John Doe",
#'     DOB = as.Date("1990-05-15"),
#'     Gender = "Male",
#'     Group = "Control"
#'   )
#' ergExam <-
#'   newERGExam(
#'     Data = Data,
#'     Metadata = Metadata,
#'     Stimulus = Stimulus,
#'     ExamInfo = ExamInfo,
#'     SubjectInfo = SubjectInfo
#'   )
#'
#' @return An object of class \code{ERGExam}.
#' @seealso \linkS4class{ERGExam}
#' @importFrom units as_units
#' @importFrom methods new validObject
#' @importFrom cli cli_abort cli_alert_warning cli_alert_info
#' @export
newERGExam <-
  function(Data,
           Metadata,
           Stimulus,
           Averaged = FALSE,
           Measurements =  new("ERGMeasurements"),
           ExamInfo,
           SubjectInfo,
           skip.validation = F) {
    # Call the default constructor
    obj <- new("ERGExam")

    if ("Step" %in% colnames(Stimulus)) {
      if ((suppressWarnings(all(!is.na(
        as.integer(as.character(Stimulus$Step))
      ))))) {
        Stimulus$Step <- as.integer(Stimulus$Step)
      } else {
        invalid_values <- Stimulus$Step[is.na(as.integer(as.character(Stimulus$Step)))]
        cli_alert_info("Checking the 'Step' column in the stimulus table...")
        cli_abort(
          c(
            "i 'Step' column contains invalid values.",
            "i The following entries could not be converted to integers: {invalid_values}.",
            "x Please ensure all values in the 'Step' column are numeric and contain no non-numeric characters."
          )
        )
      }
    } else {
      cli_alert_warning("No {.emph 'Step'} column in stimulus table encountered. Will be filled with row numbers. ")
      cli_alert_info("Stimulus table column name{? is/s are} {colnames(Stimulus)}")
      Stimulus$Step <- as.integer(1:nrow(Stimulus))
    }


    # Set the values for the slots
    obj@Data <- Data
    Metadata$Eye <- as.std.eyename(Metadata$Eye)
    obj@Metadata <- Metadata
    obj@Stimulus <- Stimulus
    obj@Averaged <- Averaged
    obj@Measurements <- Measurements
    obj@ExamInfo <- ExamInfo
    obj@SubjectInfo <- SubjectInfo

    if (is.null(obj@SubjectInfo$Group)) {
      obj@SubjectInfo$Group<-"DEFAULT"
    }

    # Set default values for other slots
    obj@Imported <- as.POSIXct(Sys.time())

    # Call the validity method to check if the object is valid
    if (skip.validation){
      if (validObject(obj)) {
        return(obj)
      }
    } else {
      return(obj)
    }
  }

#' @noMd
setAs("EPhysSet", "ERGExam", function(from) {
  new(to, Data = from@Data, Metadata = Metadata(from))
})

#' @noMd
#' @importFrom utils object.size
#' @importFrom cli cli_text cli_alert_warning cli_alert_info cli_alert_success col_green cli_ul
setMethod("show",
          "ERGExam",
          function(object) {
            cli_text("An object of class {col_green('ERGExam')}")
            cli_text("{col_green('Subject:')} {Subject(object)}, {as.character(DOB(object))}, {object@SubjectInfo$Gender}")
            cli_text("{col_green('Exam Date:')} {as.character(ExamDate(object))}")
            cli_text("{col_green('Protocol:')} {object@ExamInfo$ProtocolName}")
            cli_text("{col_green('Steps:')}")
            cli_ul(StimulusDescription(object))
            cli_text("{col_green('Eyes:')} {Eyes(object)}")
            cli_text("{col_green('Channels:')}")
            cli_ul({Channels(object)})

            if (length(object@Data) == 0) {
              cli_alert_warning("Raw data not stored in object.")
            }
            if (object@Averaged) {
              cli_alert_info("Object contains imported averaged traces.")
            }
            if (length(object@Measurements) == 0) {
              cli_alert_info("No measurements stored in object.")
            } else {
              if (object@Averaged) {
                cli_alert_success("*Measurements imported from external source.*")
              }
            }
            if (!all(unlist(lapply(object@Data, function(x) {
              dim(x)[2] == 1
            })))) {
              suppressWarnings({
                fxSet <- CheckAvgFxSet(object)
              })

              if (!fxSet) {
                cli_alert_warning("An average function has not been set for this object.")
              } else {
                cli_alert_success("An average function has been set for this object.")
              }
            } else {
              cli_alert_success("This object seems to contain averaged data or single traces.")
            }

            cli_text("Size: {format(object.size(object), 'auto')}\n")
          })
```
