## Supplemental Material 1 for "The ERGtools2 package: A Toolset for Processing and Analysing Visual Electrophysiology Data": as.data.frame.R

```
#' as.data.frame for ERGProtocol, ERGStep, ERGChannel and ERGMarker
#'
#' Converts an \link{ERGExam}, \link{ERGMeasurements-class}, \link{ERGProtocol}, ERGStep, ERGChannel or ERGMarker object to a data frame format.
#' Note that the \code{as.data.frame} method for \link{ERGExam} objects is inherited from the \link[EPhysData]{EPhysData-package} package. See: \link[EPhysData:as.data.frame]{EPhysData::as.data.frame-method}.
#' @param x An \link{ERGProtocol}, ERGStep, ERGChannel or ERGMarker object.
#' @param ... currently unused.
#' @details
#' as.data.frame convert various ERG-related objects (\link{ERGExam}, \link{ERGMeasurements}, \link{ERGProtocol}, ERGMarker, ERGChannel, and ERGStep) to data frame formats, facilitating data manipulation and analysis. Each method extracts relevant information from the respective object and organizes it into a structured data frame.
#'
#' - For \link{ERGExam} it is inherited from the \link[EPhysData]{EPhysData-package}.
#' - For \link{ERGMeasurements-class} it is an alias for Measurements() (see: \link{Measurements-Methods}).
#'
#' The ERGProtocol-related methods are experimental. These include:
#' - For ERGMarker objects, the resulting data frame contains two columns: Marker.Name and Marker.Relative.to, representing the marker's name and its relative position.
#' - For ERGChannel objects, the resulting data frame includes channel properties such as name, eye, frequency cutoffs, and inversion status, along with information about associated markers.
#' - For ERGStep objects, the resulting data frame includes step properties such as description, adaptation, and recording parameters, along with information about associated channels.
#' - For ERGProtocol objects, the resulting data frame includes protocol properties such as name and export date, along with information about associated steps.
#'
#' These data frames provide comprehensive representations of the respective ERG-related objects, allowing for further analysis, visualization, or integration with other data.

#' @return A data frame representing the \link{ERGExam}, \link{ERGMeasurements}, \link{ERGProtocol}, ERGStep, ERGChannel or ERGMarker object in long format.
#' @examples
#' data(ERG)
#' ERG <- SetStandardFunctions(ERG)
#' ERG <- Subset(ERG,where=list(Step=as.integer(1),Eye="RE")) # converting the whole object would return a huhge data.frame
#' head(as.data.frame(ERG))
#' @importFrom tidyr pivot_longer starts_with
#' @importFrom units drop_units as_units
#' @seealso \link{ERGExam}, \link{ERGMeasurements-class}, \link{ERGProtocol}, \link[EPhysData]{EPhysData-package}, \link{Measurements-Methods}
#' @name as.data.frame
NULL

#' @describeIn as.data.frame Method for ERGMeasurements
#' @exportMethod as.data.frame
setMethod("as.data.frame",
          "ERGMeasurements",
          function(x,
                   ...) {
            return(Measurements(x))
          })
#' @describeIn as.data.frame Method for ERGMarker
#' @exportMethod as.data.frame
setMethod("as.data.frame",
          "ERGMarker",
          function(x,
                   ...) {
            out <-
              data.frame(Marker.Name = character(), Marker.Relative.to = character())
            out[1, ] <- c(x@Name, x@RelativeTo)
            return(out)
          })

#' @describeIn as.data.frame Method for ERGChannel
#' @exportMethod as.data.frame
setMethod("as.data.frame",
          "ERGChannel",
          function(x,
                   ...) {
            markers <- lapply(x@Markers, as.data.frame)
            markers <- do.call(rbind, markers)
            out <-
              data.frame(
                Channel.Name = character(),
                Eye = character(),
                LowFreqCutoff = numeric(),
                HighFreqCutoff = numeric(),
                Channel.Inverted = logical(),
                Channel.Enabled = logical()
              )
            nMarkers <- nrow(markers)
            if (length(nMarkers) < 1) {
              nMarkers <- 1
              marker <- ERGMarker()
              marker@Name <- ""
              markers <- as.data.frame(marker)
            }
            out <- out[1:nMarkers,]
            out$Channel.Name[1:nMarkers] <- as.character(x@Name)
            out$Eye[1:nMarkers]  <- x@Eye
            out$LowFreqCutoff[1:nMarkers] <- x@LowFreqCutoff
            out$HighFreqCutoff[1:nMarkers] <- x@HighFreqCutoff
            out$Channel.Inverted[1:nMarkers] <- x@Inverted
            out$Channel.Enabled[1:nMarkers] <- x@Enabled

            return(cbind(out, markers))
          })

#' @describeIn as.data.frame Method for ERGStep
#' @exportMethod as.data.frame
setMethod("as.data.frame",
          "ERGStep",
          function(x,
                   ...) {
            channels <- lapply(x@Channels, as.data.frame)
            channels <- do.call(rbind, channels)
            nChannels <- nrow(channels)

            # Create a data frame for ERGStep slots and repeat values based on nMarkers
            step_df <- data.frame(
              Description = rep(as.character(x@Description), each = nChannels),
              Adaptation = rep(as.character(x@Adaptation), each = nChannels),
              Resultsperrun = rep(as.numeric(x@Resultsperrun), each = nChannels),
              Timebetweenresults = rep(as.character(x@Timebetweenresults), each = nChannels),
              SampleFrequency = rep(as.character(x@SampleFrequency), each = nChannels),
              PreTriggerTime = rep(as.character(x@PreTriggerTime), each = nChannels),
              PostTriggerTime = rep(as.character(x@PostTriggerTime), each = nChannels),
              InterSweepDelay = rep(as.character(x@InterSweepDelay), each = nChannels),
              Sweepsperresult = rep(as.numeric(x@Sweepsperresult), each = nChannels),
              Sweepsperresultmin = rep(as.numeric(x@Sweepsperresultmin), each = nChannels),
              Glitchremoval = rep(as.logical(x@Glitchremoval), each = nChannels),
              Driftremoval = rep(as.logical(x@Driftremoval), each = nChannels),
              Baselineenabled = rep(as.logical(x@Baselineenabled), each = nChannels),
              Baselinepretrigger = rep(as.character(x@Baselinepretrigger), each = nChannels),
              Baselinerange = rep(as.character(x@Baselinerange), each = nChannels)
            )

            # Combine the ERGStep data frame with the channel data frame
            result_df <- cbind(step_df, channels)
            return(result_df)
          })
#' @describeIn as.data.frame Method for ERGProtocol
#' @exportMethod as.data.frame
as.data.frame <- setMethod("as.data.frame",
                           signature = "ERGProtocol",
                           function(x, ...) {
                             steps <- lapply(x@Step, as.data.frame)
                             steps <- do.call(rbind, steps)
                             nSteps <- nrow(steps)

                             # Create a data frame for ERGProtocol slots and repeat values based on nChannels
                             protocol_df <- data.frame(
                               ProtocolName = rep(as.character(x@Name), each = nSteps),
                               Export_Date = rep(as.character(x@Export_Date), each = nSteps),
                               row.names = 1:nSteps
                             )

                             # Combine the ERGProtocol data frame with the step data frame
                             result_df <- cbind(protocol_df, steps)

                             return(result_df)
                           })
```
