## Supplemental Material 1 for "The ERGtools2 package: A Toolset for Processing and Analysing Visual Electrophysiology Data": AutoPlaceMarkers.R

```
#' AutoPlaceMarkers for ERG/VEP Recordings
#'
#' These methods automatically place markers for ERG/VEP recordings in an \linkS4class{ERGExam} object.
#'
#' @inheritParams SetStandardFunctions
#' @param X An \linkS4class{ERGExam} object for \code{AutoPlaceMarkers()} or an \link[EPhysData:EPhysData-class]{EPhysData::EPhysData-class} object for the lower level methods \link[=AutoPlaceAB]{AutoPlaceAB}, \link[=AutoPlaceFlicker]{AutoPlaceFlicker} or \link[=AutoPlaceVEP]{AutoPlaceVEP}.
#' @param Channel.names A \code{pairlist} specifying channel identifiers
#' @param robust.peak.filter.bands A numeric vector of  of class units (\link[units:units]{units::units}  and length 2 specifying the lower and upper bounds of the frequency band used for initial peak identification. Must be in unit H).
#' @param true.peak.tolerance A vector of class units (\link[units:units]{units::units}  and length 2 specifying the tolerance range around true peaks. Must be time values (i.e. a unit convertible into 'seconds').
#' @param Param.list A named list to override default parameters for the lower-level marker placement methods. The list must contain named elements corresponding to one or more of the lower-level methods: \link[=AutoPlaceAB]{AutoPlaceAB}, \link[=AutoPlaceFlicker]{AutoPlaceFlicker}, or \link[=AutoPlaceVEP]{AutoPlaceVEP}. Each element in the list must be a named list containing specific parameters for that method, such as `robust.peak.filter.bands` or `true.peak.tolerance`. If not provided, the default values for each method will be used. If a method is not specified in `Param.list`, its default parameters will be applied.
#' @importFrom cli cli_progress_bar cli_progress_update cli_progress_done
#'
#' @details These methods are used to automatically place markers for ERGs/VEPs.\cr\cr
#' \code{AutoPlaceMarkers()} sets markers depending on the channel (E.g. ERG, VEP, OP,...) and stimulus type (Flash, Flicker), defined via the \code{Channel.names} and \code{Stimulus.type.names} arguments. Markers are placed using the lower level methods \link[=AutoPlaceAB]{AutoPlaceAB}, \link[=AutoPlaceFlicker]{AutoPlaceFlicker} or \link[=AutoPlaceVEP]{AutoPlaceVEP} function depending on the provided combination of \code{Channel.names} and \code{Stimulus.type.names}.\cr\cr
#'  \link[=AutoPlaceAB]{AutoPlaceAB}, \link[=AutoPlaceFlicker]{AutoPlaceFlicker} and \link[=AutoPlaceVEP]{AutoPlaceVEP} are the lower level functions which perform the actual marker placement on the \link[EPhysData:EPhysData-class]{EPhysData::EPhysData-class} objects contained in the \linkS4class{ERGExam} object. Those methods are usually not called directly by a user, unless she/he wants to perform or re-run marker placement only on certain recordings while leaving previously set markers unchanged for the others.
#'  There working principle is that they apply robust peak filtering within defined frequency bands (low frequency band by default) to locate the gross position of the most prominent peaks peaks and then look for the peak in data using the preset filter function (\link{FilterFunction}) to accurately identify the actual peak position. \cr\cr
#'  Currently, supported are:
#' * a and B waves for Flash ERG
#' * N1, P1 (and Frequency) for Flicker ERGs
#' * P1, N1, and P2 for Flash ERGs \cr\cr
#'
#' @return An updated \code{ERGExam} object with markers placed.
#'
#' @examples
#' # Preparing data
#' data(ERG)
#' ERG<-SetStandardFunctions(ERG)
#' imported_Markers<-Measurements(ERG)
#' head(imported_Markers)
#' ERG<-ClearMeasurements(ERG)
#' imported_Markers_cleared<-Measurements(ERG)
#' head(imported_Markers_cleared)
#' # Using the top-level AutoPlaceMarkers method without manually setting peak detection parameters
#' ERG<-AutoPlaceMarkers(ERG, Channel.names = pairlist(ERG = "ERG"))
#' autoplaced_Markers<-Measurements(ERG)
#' head(autoplaced_Markers)
#' ggERGExam(ERG)
#' # Using the top-level AutoPlaceMarkers method without manually setting peak detection parameters
#' ERG <-
#'   AutoPlaceMarkers(
#'     ERG,
#'     Channel.names = pairlist(ERG = "ERG"),
#'     Param.list <- list(
#'       AutoPlaceAB = list (
#'         robust.peak.filter.bands = as_units(c(5, 75), "Hz"),
#'         true.peak.tolerance = as_units(c(5, 5), "ms")
#'       )
#'     )
#'   )
#' autoplaced_Markers<-Measurements(ERG)
#' head(autoplaced_Markers)
#' ggERGExam(ERG)
#'
#' # Calling AutoPlaceAB() directly
#' X<-ERG@Data[[1]] # get first recording
#' AutoPlaceAB(X)

#' @name AutoPlaceMarkers
NULL
#' @describeIn AutoPlaceMarkers Automatically sets markers depending on the channel (E.g. ERG, VEP, OP,...) and stimulus type (Flash, FLicker).
#' @exportMethod AutoPlaceMarkers
setGeneric(
  name = "AutoPlaceMarkers",
  def = function(X,
                 Channel.names = pairlist(ERG = "ERG",
                                          VEP = "VEP"),
                 Stimulus.type.names = pairlist(Flash = "Flash",
                                                Flicker = "Flicker"),
                 Param.list = list()) {
    standardGeneric("AutoPlaceMarkers")
  }
)

#' @noMd
setMethod(
  "AutoPlaceMarkers",
  signature = "ERGExam",
  definition = function(X,
                        Channel.names = pairlist(ERG = "ERG",
                                                 VEP = "VEP"),
                        Stimulus.type.names = pairlist(Flash = "Flash",
                                                       Flicker = "Flicker"),
                        Param.list = list()) {


    stopifnot(CheckAvgFxSet(X))

    default.param.list <- list(
      AutoPlaceAB = list (
        robust.peak.filter.bands = as_units(c(5, 75), "Hz"),
        true.peak.tolerance = as_units(c(70, 30), "ms")
      ),
      AutoPlaceFlicker = list (
        robust.peak.filter.bands = as_units(c(.5, 300), "Hz"),
        true.peak.tolerance = as_units(c(12, 15), "ms")
      ),
      AutoPlaceVEP = list (
        robust.peak.filter.bands = as_units(c(1, 75), "Hz"),
        true.peak.tolerance = as_units(c(80, 20), "ms")
      )
    )
    if (length(Param.list)!=0){

      if (!all(unique(unlist(lapply(Param.list, names))) %in% "robust.peak.filter.bands",
               "true.peak.tolerance")) {
        Notice(X,
               what = "E",
               notice_text = "AutoPlaceMarkers failed. Param.list contains the subitems that do not match the names of the parameters of the three low-level marker placement methods ({.fun AutoPlaceAB}, {.fun AutoPlaceFlicker} or {.fun AutoPlaceVEP}). Allowed values are: 'robust.peak.filter.bands' and 'true.peak.tolerance'",
               help_page = "ERGtools2::AutoPlaceMarkers")
      }

      for (n in names(Param.list)) {
        if(n %in% names(default.param.list)){
          if (all(c("robust.peak.filter.bands","true.peak.tolerance") %in% names(default.param.list[[n]]))){
            if (all(lapply(default.param.list[[n]], length))) {
              default.param.list[[n]] <- Param.list[[n]]
            } else {
              Notice(X,
                     what = "E",
                     notice_text = "AutoPlaceMarkers failed. For list entry {.val {n}} in Param.list two vectors ('robust.peak.filter.bands' and 'true.peak.tolerance') of length two are expected. however, lengths were {.val lapply(default.param.list[[n]], length)}",
                     help_page = "ERGtools2::AutoPlaceMarkers")
            }
          } else {
            Notice(X,
                   what = "E",
                   notice_text = "AutoPlaceMarkers failed. For list entry {.val {n}} in Param.list two vectors named 'robust.peak.filter.bands' and 'true.peak.tolerance' are expected. however, names encounterd were {.val names(default.param.list[[n]])}",
                   help_page = "ERGtools2::AutoPlaceMarkers")
          }
        } else {
          Notice(X,
                 what = "E",
                 notice_text = "AutoPlaceMarkers failed. Param.list contains the named list item {.val {n}}. However, list item names must be either of the names of the three low-level marker placement methods ({.fun AutoPlaceAB}, {.fun AutoPlaceFlicker} or {.fun AutoPlaceVEP})",
                 help_page = "ERGtools2::AutoPlaceMarkers")
        }
        Param.list<-default.param.list
      }
    }

    markerlist <- list()
    Md <- merge(Metadata(X), StimulusTable(X))
    cli_progress_bar("Placing markers", total = length(X@Data),  clear = TRUE, auto_terminate = T)
    for (i in 1:length(X@Data)) {
      x <- X@Data[[i]]
      update <- F
      # FLASH ERGs
      tryCatch({
        if (Md$Channel[i] %in% Channel.names$ERG &
            Md$Type[i]  %in%  Stimulus.type.names$Flash) {
          Markers <-
            AutoPlaceAB(
              x,
              robust.peak.filter.bands = default.param.list$AutoPlaceAB$robust.peak.filter.bands,
              true.peak.tolerance = default.param.list$AutoPlaceAB$true.peak.tolerance
            )
          if (nrow(Markers) > 0) {
            update <- T
          }
        }
        # Flicker ERGs
        if (Md$Channel[i] %in% Channel.names$ERG &
            Md$Type[i]  %in%  Stimulus.type.names$Flicker) {
          Markers <- AutoPlaceFlicker(
            x,
            robust.peak.filter.bands = default.param.list$AutoPlaceFlicker$robust.peak.filter.bands,
            true.peak.tolerance = default.param.list$AutoPlaceFlicker$true.peak.tolerance
          )
          if (nrow(Markers) > 0) {
            update <- T
          }
        }
        # Flash VEP
        if (Md$Channel[i] %in% Channel.names$VEP &
            Md$Type[i]  %in%  Stimulus.type.names$Flash) {
          Markers <- AutoPlaceVEP(
            x,
            robust.peak.filter.bands = default.param.list$AutoPlaceVEP$robust.peak.filter.bands,
            true.peak.tolerance = default.param.list$AutoPlaceVEP$true.peak.tolerance
          )
          if (nrow(Markers) > 0) {
            update <- T
          }
        }

        if (update) {
          Markers$Name <- rownames(Markers)
          if (nrow(Markers[is.na(Markers$Relative),]) > 0) {
            for (m in (1:nrow(Markers))[is.na(Markers$Relative)]) {
              Measurements(
                X,
                Marker = Markers$Name[m],
                where = i,
                create.marker.if.missing = T,
                Relative = Markers$Relative[m],
                ChannelBinding = Md$Channel[i]
              ) <- Markers$Time[m]
            }
            for (m in (1:nrow(Markers))[!is.na(Markers$Relative)]) {
              Measurements(
                X,
                Marker = Markers$Name[m],
                where = i,
                create.marker.if.missing = T,
                Relative = Markers$Relative[m],
                ChannelBinding = Md$Channel[i]
              ) <- Markers$Time[m]
            }
          }
        }
      },
      error = function(e) {
        Notice(X,
               where = i,
               what = "E",
               notice_text = "Auto placement of markers failed with error message: {.emph {e}}")
      })
      cli_progress_update()
    }
    cli_progress_done()
    return(X)
  }
)

#' @describeIn AutoPlaceMarkers places the a and B waves on Flash ERG data stored in an an \link[EPhysData:EPhysData-class]{EPhysData::EPhysData-class} object.
#' @exportMethod AutoPlaceAB
setGeneric(
  name = "AutoPlaceAB",
  def = function(X,
                 robust.peak.filter.bands = as_units(c(5, 75), "Hz"),
                 true.peak.tolerance = as_units(c(70, 30), "ms")) {
    standardGeneric("AutoPlaceAB")
  }
)
#' @importFrom EPhysMethods filter.bandpass freq.to.w
#' @noMd
setMethod(
  "AutoPlaceAB",
  signature = "EPhysData",
  definition = function(X,
                        robust.peak.filter.bands = as_units(c(5, 75), "Hz"),
                        true.peak.tolerance = as_units(c(70, 30), "ms")) {
    if (!("units" %in% class(true.peak.tolerance))) {
      stop("'true.peak.tolerance' must be of class units.")
    }
    if (!("units" %in% class(robust.peak.filter.bands))) {
      message("Its recommended that 'robust.peak.filter.bands' is of class units. Auto-setting it to unit 'Hz'")
      robust.peak.filter.bands<-as_units(robust.peak.filter.bands,"Hz")
    }

    if (!units.is.convertible(robust.peak.filter.bands, "Hz")) {
      stop("'robust.peak.filter.bands' must be of convertible to Hertz")
    }
    if (!units.is.convertible(true.peak.tolerance, "s")) {
      stop("'true.peak.tolerance' must be of convertible to seconds.")
    }

    robust.peak.filter.bands<-convert_to_unit(robust.peak.filter.bands,"Hz")
    true.peak.tolerance<-convert_to_unit(true.peak.tolerance,"s")

    dat <- as.data.frame(X, Raw = F)
    sample.rate <- mean(diff(TimeTrace(X)))
    sample.rate <- set_units(sample.rate,"s")

    cutoff <-
      freq.to.w(x = robust.peak.filter.bands, time.trace = TimeTrace(X))
    dat$Filtered <- filter.bandpass(dat$Value, cutoff[1], cutoff[2])
    B_estimate <- which.max(dat$Filtered)
    a_estimate <- which.min(dat$Filtered[1:B_estimate])

    search_left <-
      round(drop_units(true.peak.tolerance[1] / (sample.rate)))
    search_right <-
      round(drop_units(true.peak.tolerance[2] / (sample.rate)))

    B_pos <- tryCatch({
      which.max(dat$Value[(B_estimate - search_left):(B_estimate + search_right)]) + B_estimate - search_left
    }, error = function(e) {
      NULL
    })

    a_pos <- tryCatch({
      which.min(dat$Value[max((a_estimate - search_left),0):min((a_estimate + search_right),length(dat$Value))]) + max((a_estimate - search_left),0)
    }, error = function(e) {
      NULL
    })

    if (!is.null(a_pos)) {
      a_time <- dat$Time[a_pos]
      a_amp <- dat$Value[a_pos]
    } else {
      a_time <- as_units(NA, deparse_unit(dat$Time))
      a_amp <- as_units(NA, deparse_unit(dat$Value))
    }

    if (!is.null(B_pos) && !is.null(a_pos)) {
      B_time <- dat$Time[B_pos]
      B_amp <- dat$Value[B_pos] - dat$Value[a_pos]
    } else {
      B_time <- as_units(NA, deparse_unit(dat$Time))
      B_amp <- as_units(NA, deparse_unit(dat$Value))
    }


    out <- data.frame(
      Time = c(a_time, B_time),
      Value = c(a_amp, B_amp),
      Relative = c(NA, "a")
    )
    rownames(out) <- c("a", "B")
    out <-
      out[!is.na(out$Value), ] # workaround as downstream fx cannnot handle na
    return(out)
  }
)

#' @describeIn AutoPlaceMarkers places the N1 and P1 markers and determines 1/frequency (period) for Flicker ERG data stored in an an \link[EPhysData:EPhysData-class]{EPhysData::EPhysData-class} object
#' @exportMethod AutoPlaceFlicker
setGeneric(
  name = "AutoPlaceFlicker",
  def = function(X,
                 robust.peak.filter.bands = as_units(c(.5, 300), "Hz"),
                 true.peak.tolerance = as_units(c(12, 15), "ms")) {
    standardGeneric("AutoPlaceFlicker")
  }
)
#' @importFrom EPhysMethods filter.bandpass fastfourier freq.to.w convert_to_unit
#' @noMd
setMethod(
  "AutoPlaceFlicker",
  signature = "EPhysData",
  definition = function(X,
                        robust.peak.filter.bands = as_units(c(.5, 300), "Hz"),
                        true.peak.tolerance = as_units(c(12, 15), "ms")) {
    if (!("units" %in% class(true.peak.tolerance))) {
      stop("'true.peak.tolerance' must be of class units.")
    }
    if (!("units" %in% class(robust.peak.filter.bands))) {
      message("Its recommended that 'robust.peak.filter.bands' is of class units. Auto-setting it to unit 'Hz'")
      robust.peak.filter.bands<-as_units(robust.peak.filter.bands,"Hz")
    }

    if (!units.is.convertible(robust.peak.filter.bands, "Hz")) {
      stop("'robust.peak.filter.bands' must be of convertible to Hertz")
    }
    if (!units.is.convertible(true.peak.tolerance, "s")) {
      stop("'true.peak.tolerance' must be of convertible to seconds.")
    }

    robust.peak.filter.bands<-convert_to_unit(robust.peak.filter.bands,"Hz")
    true.peak.tolerance<-convert_to_unit(true.peak.tolerance,"s")

    cutoff <-
      freq.to.w(x = robust.peak.filter.bands, time.trace <-
                  TimeTrace(X))
    sample.rate <- mean(diff(TimeTrace(X)))
    sample.rate <- set_units(sample.rate, "s")

    dat <- GetData(X, Raw = T)
    dat <- filter.bandpass(dat, cutoff[1], cutoff[2])
    fft <- fastfourier(dat, samp.freq = 1 / sample.rate)
    fft_short <- lapply(fft, function(x) {
      x$freq<-as_units(x$freq,"Hz")
      x <- x[x$freq < robust.peak.filter.bands[2]/2,] # flicker freq can only be detected if at least half the max bandpass frequency
      x <- x[x$freq > max(robust.peak.filter.bands[1]*2,set_units(1/max(TimeTrace(X)),"Hz")*2),] # flicker freq can only be detected if at least oble the min beandpass frequnecy and if at least two peaks fit into one trace
      x$freq_rounded <- round(x$freq)
      x$fur <- abs(x$fur)
      averaged_data <-
        aggregate(fur ~ freq_rounded, data = x, FUN = mean)
      names(averaged_data)[names(averaged_data) == "freq_rounded"] <-
        "freq"
      x <- averaged_data
      x
    })
    fur <- lapply(fft_short, function(x) {
      x$fur
    })

    fur <- do.call(cbind, fur)
    fur <- apply(fur, 1, median)
    domfreq <- fft_short[[1]]$freq[which.max(fur)]
    # 50Hz reject
    if (domfreq > as_units(47,"Hz") & domfreq < as_units(53,"Hz")) {
      fur[which.max(fur)] <- NA
      domfreq <- fft_short[[1]]$freq[which.max(fur)]
    }

    #todo: Implement options to chech for loss of power over repeats

    peak_interval <- 1/sample.rate / set_units(domfreq,"1/s")
    peak_interval <-
      round(drop_units(set_units(1 / domfreq, "s") / sample.rate))

    # a matrix for averaging, based on the peak interval calculated from the fourier transform
    dat<-as.data.frame(X,Raw=F)
    dat$Filtered <- filter.bandpass(dat$Value, cutoff[1], cutoff[2])

    start <- which(dat$Time == as_units(0, "ms"))
    end <- start + peak_interval
    posmtx <-
      matrix(nrow = (end - start),
             ncol = floor(nrow(dat) / peak_interval))
    for (i in 1:nrow(posmtx)) {
      tmp <- seq(i + start, nrow(dat), peak_interval)
      if (length(tmp) < ncol(posmtx)) {
        tmp <- c(tmp, rep(NA, ncol(posmtx) - length(tmp)))
      }
      posmtx[i, ] <- tmp[1:ncol(posmtx)]#[1:2]
    }

    posmtx<-posmtx[, apply(posmtx, 2, function(x) {
      !any(is.na(x))
    })]

    peak.avg <- apply(posmtx, 1, function(x) {
      mean(dat$Filtered[x], na.rm = F)
    })

    P1_estimate <- dat$Time[which.max(peak.avg) + start]
    P1_estimate_idx <- which(dat$Time == P1_estimate)
    N1_estimate <- dat$Time[which.min(peak.avg[1:P1_estimate_idx]) + start]
    N1_estimate_idx <- which(dat$Time == N1_estimate)

    #error below w search_left and right ms to interval

    search_left <-
      round(drop_units(
        set_units(true.peak.tolerance[1], deparse_unit(sample.rate)) / (sample.rate)
      ))
    search_right <-
      round(drop_units(
        set_units(true.peak.tolerance[2], deparse_unit(sample.rate)) / (sample.rate)
      ))

    P1_pos <- tryCatch({
      which.max(dat$Value[max((P1_estimate_idx - search_left), 1):min((P1_estimate_idx + search_right), length(dat$Value))]) +
        max((P1_estimate_idx - search_left), 1)
    }, error = function(e) {
      NULL
    })
    N1_pos <- tryCatch({
      which.min(dat$Value[max((N1_estimate_idx - search_left), 1):(P1_pos)]) + max((N1_estimate_idx - search_left), 1)
    }, error = function(e) {
      NULL
    })


    P1_time <- dat$Time[P1_pos]
    P1_amp <- dat$Value[P1_pos] - dat$Value[N1_pos]
    N1_time <- dat$Time[N1_pos]
    N1_amp <- dat$Value[N1_pos]

    out <-
      data.frame(
        Time = c(
          N1_time,
          P1_time,
          set_units(set_units(1 / domfreq, "s"), units(P1_time), mode = "standard")
        ),
        Value = c(N1_amp, P1_amp, as_units(NA, "V")),
        Relative = c(NA, "N1", NA)
      )
    rownames(out) <- c("N1", "P1", "Period_length")
    return(out)
  }
)

#' @describeIn AutoPlaceMarkers places the P1, N1 and P2 markers for Flash VEP data stored in an an \link[EPhysData:EPhysData-class]{EPhysData::EPhysData-class} object
#' @exportMethod AutoPlaceVEP
setGeneric(
  name = "AutoPlaceVEP",
  def = function(X,
                 robust.peak.filter.bands = as_units(c(3, 75), "Hz"),
                 true.peak.tolerance = as_units(c(80, 20), "ms")) {
    standardGeneric("AutoPlaceVEP")
  }
)
#' @importFrom EPhysMethods filter.bandpass freq.to.w
#' @importFrom quantmod findPeaks
#' @noMd
setMethod(
  "AutoPlaceVEP",
  signature = "EPhysData",
  definition = function(X,
                        robust.peak.filter.bands = as_units(c(1, 75), "Hz"),
                        true.peak.tolerance = as_units(c(80, 20), "ms")) {
    if (!("units" %in% class(true.peak.tolerance))) {
      stop("'true.peak.tolerance' must be of class units.")
    }
    if (!("units" %in% class(robust.peak.filter.bands))) {
      message("Its recommended that 'robust.peak.filter.bands' is of class units. Auto-setting it to unit 'Hz'")
      robust.peak.filter.bands<-as_units(robust.peak.filter.bands,"Hz")
    }

    if (!units.is.convertible(robust.peak.filter.bands, "Hz")) {
      stop("'robust.peak.filter.bands' must be of convertible to Hertz")
    }
    if (!units.is.convertible(true.peak.tolerance, "s")) {
      stop("'true.peak.tolerance' must be of convertible to seconds.")
    }

    robust.peak.filter.bands<-convert_to_unit(robust.peak.filter.bands,"Hz")
    true.peak.tolerance<-convert_to_unit(true.peak.tolerance,"s")

    dat <- as.data.frame(X, Raw = F)
    sample.rate <- mean(diff(TimeTrace(X)))

    cutoff <-
      freq.to.w(x = robust.peak.filter.bands, time.trace <- TimeTrace(X))
    dat$Filtered <- filter.bandpass(dat$Value, cutoff[1], cutoff[2])
    N1_estimate <- which.min(dat$Filtered)
    P1_estimate <- findPeaks(dat$Filtered[1:N1_estimate])
    P1_estimate <- P1_estimate[length(P1_estimate)]

    P2_estimate <-
      findPeaks(dat$Filtered[N1_estimate:length(dat$Filtered)])[1] + N1_estimate

    search_left <-
      drop_units(true.peak.tolerance[1] / (1 / sample.rate))
    search_right <-
      drop_units(true.peak.tolerance[2] / (1 / sample.rate))

    N1_pos <- tryCatch({
      which.min(dat$Value[(N1_estimate - search_left):(N1_estimate + search_right)]) +
        N1_estimate - search_left
    }, error = function(e)
      NULL)

    P1_pos <- tryCatch({
      which.max(dat$Value[(P1_estimate - search_left):(N1_pos)]) + #P1_estimate + search_right
        P1_estimate - search_left
    }, error = function(e)
      NULL)
    P2_pos <- tryCatch({
      which.max(dat$Value[(N1_pos):(P2_estimate + search_right)]) + # P2_estimate - search_left
        N1_pos #P2_estimate - search_left
    }, error = function(e)
      NULL)

    if (!is.null(P1_pos)) {
      P1_time <- dat$Time[P1_pos]
      P1_amp <- dat$Value[P1_pos]
    } else{
      P1_time <- as_units(NA, deparse_unit(dat$Time))
      P1_amp <- as_units(NA, deparse_unit(dat$Value))
    }
    if (!is.null(N1_pos)) {
      N1_time <- dat$Time[N1_pos]
      N1_amp <- dat$Value[N1_pos] - P1_amp
    } else{
      N1_time <- as_units(NA, deparse_unit(dat$Time))
      N1_amp <- as_units(NA, deparse_unit(dat$Value))
    }
    if (!is.null(P2_pos)) {
      P2_time <- dat$Time[P2_pos]
      P2_amp <- dat$Value[P2_pos] - P1_amp
    } else{
      P2_time <- as_units(NA, deparse_unit(dat$Time))
      P2_amp <- as_units(NA, deparse_unit(dat$Value))
    }

    out <- data.frame(
      Time = c(P1_time, N1_time, P2_time),
      Value = c(P1_amp, N1_amp, P2_amp),
      Relative = c(NA, "P1", "P1")
    )
    rownames(out) <- c("P1", "N1", "P2")
    return(out)
  }
)

#' @importFrom units set_units
#' @keywords internal
units.is.convertible <- function(x, to) {
  # this function will mote to EPhysMethods in future
  tryCatch({
    set_units(x, "s")
    TRUE
  }, error = function(e) {
    FALSE
  })
}
```
