## Supplemental Material 1 for "The ERGtools2 package: A Toolset for Processing and Analysing Visual Electrophysiology Data": Checks.R

```
#' Check if all recordings in an ERGExam object have a valid averaging function set.
#'
#' @param X An ERGExam object.
#'
#' @return Logical, TRUE if all recordings have a valid averaging function, FALSE otherwise.
#'
#' @examples
#' data(ERG)
#' CheckAvgFxSet(ERG)
#' ERG<-SetStandardFunctions(ERG)
#' CheckAvgFxSet(ERG)
#'
#' @seealso
#' \code{\link{AverageFunction<-}}, \code{\link{SetStandardFunctions}}
#'
#' @export
setGeneric(
  name = "CheckAvgFxSet",
  def = function(X)
  {
    standardGeneric("CheckAvgFxSet")
  }
)
#' @noMd
setMethod("CheckAvgFxSet",
          "ERGExam",
          function(X) {
            fx.set <- unlist(lapply(X@Data, function(x) {
              suppressWarnings({
                dat <-
                  GetData(x, Raw = FALSE, Time = c(min(TimeTrace(x)), TimeTrace(x)[min(5, length(TimeTrace(x)))]), Trials = c(1:(min(5, dim(x)[2]))))
              })
              return(as.logical(ncol(dat) == 1))
            }))
            if (!all(fx.set)) {
              Notice(
                X,
                what = c("Warning"),
                where = which(!fx.set),
                notice_text = c(
                  "No valid averaging function found. Result of averaging Data must be a vector. "
                ),
                help_page = "ERGtools2::SetStandardFunctions"
              )
              return(FALSE)
            } else{
              return(TRUE)
            }
          })
```
