## Supplemental Material 1 for "The ERGtools2 package: A Toolset for Processing and Analysing Visual Electrophysiology Data": CollectMeasurements.R

```
#' Get measurements for plotting
#'
#' This function extracts measurements and related information, e.g. for plotting or statistics.
#'
#' @param List A list of ERG exams.
#' @inheritParams Where
#' @inheritParams Measurements
#' @param Markers Vector of markers to include in the plot.
#'
#' @return A data frame with measurements for plotting.
#' @importFrom EPhysData AverageFunction `AverageFunction<-` Rejected `Rejected<-` FilterFunction `FilterFunction<-`
#' @importFrom cli cli_abort cli_progress_bar cli_progress_update cli_progress_done
#' @examples
#' data(ERG)
#' ERG<-SetStandardFunctions(ERG)
#' ERG <- AutoPlaceMarkers(ERG)
#' CollectMeasurements(list(ERG,ERG), list(Background = "DA", Type = "Flash"))
#' @export
CollectMeasurements <- function(List,
                                where = list(),
                                Markers = c("a", "B", "N1", "P1"),
                                measure.absolute = F) {
  # compatibility f single Exam
  quiet=F
  if (!all(unlist((lapply(List, function(x) {
    inherits(x, "ERGExam")
  }))))) {
    if (inherits(List, "ERGExam")) {
      return(
        Measurements(
          List,
          where = where,
          Marker = Markers,
          measure.absolute = measure.absolute
        )
      )
    }
    cli_abort("'List' is not a list of ERGExams.")
  }
  # subset object
  List <- lapply(List, function(x) {
    if(!CheckAvgFxSet(x)){
      Notice(x,
             what = c("Error"),
             notice_text = c("x Average functions must be set for all objects in the list."),
             help_page = "EPhysData::AverageFunction")
    }
    tryCatch({
      x <- Subset(x, where = where, Raw = T)
      return(x)
    }, error = function(e){
      Notice(x,
             what = c("Error"),
             notice_text = c("x Fetching Metadata and Stimulus values failed with error message {e} evoked by {.fun Subset}"),
             help_page = "EPhysData::Subset")
    })
  })
  if(!quiet){
    cli_progress_bar("Collecting measurements", total = length(List),  clear = TRUE)
  }
  results <- list()
  for (i in 1: length(List)){
    x<-List[[i]]
    tryCatch({
      df <- Measurements(x, measure.absolute = measure.absolute, quiet = T)
      markers<-unique(df$Name)
      for(m in markers){
        if(length(df$Recording[df$Name==m])!=length(unique(df$Recording))){

          unique_recordings <- unique(df$Recording)
          filtered_recordings <- unique(df$Recording[df$Name == m])
          missing_recordings <- setdiff(unique_recordings, filtered_recordings)
          Notice(x,
                 what = c("W"),
                 where = missing_recordings,
                 notice_text = c("x Marker {m} missing for at least one of the recordings."),
                 help_page = "EPhysData::Measurements")
        }
      }

      df<-df[,!(colnames(df) %in% ExtraMetaColumns(x))]
      if(nrow(df)==0){
        Notice(x,
               what = c("W"),
               where = missing_recordings,
               notice_text = c("x No measurements found."),
               help_page = "ERGtools2::Measurements")
      }

      if(nrow(df)>0){
        if (length(x@SubjectInfo$Group)==0) {
          x@SubjectInfo$Group<-"DEFAULT"
        }
        df$Subject <- Subject(x)
        df$Group <- GroupName(x)
        df$ExamDate <- min(x@ExamInfo$ExamDate)
      }else{
        df$Subject <- character()
        df$Group <- character()
        df$ExamDate <- as.Date(x = integer(0), origin = "1970-01-01")
      }

      df <-
        merge(df,
              StimulusTable(x),
              by.x = c("Step","Description"),
              by.y = c("Step","Description"))
      results[[i]]<-df
    }, error = function(e){
      Notice(x,
             what = c("W"),
             notice_text = c("x Fetching Measurements failed with error message '{e}'."),
             help_page = "ERGtools2::Measurements")
    })
    if (!quiet) {
      cli_progress_update()
    }
  }

  if (!quiet) {
    cli_progress_done()
  }
  results <- do.call(rbind.data.frame, results)
  # type conversion and column names
  colnames(results)[colnames(results) == "cd.s.m"] <- "StimulusEnergy"
  results$Step.y <- NULL
  results$Step <- iconv(results$Step, "ASCII//TRANSLIT", sub = '')

  # subsetting list
  results <-
    results[results$Name %in% Markers, ]
  return(results)
}

#' @export
#' @noRd
Collect_Measurements<-CollectMeasurements
```
