## Supplemental Material 1 for "The ERGtools2 package: A Toolset for Processing and Analysing Visual Electrophysiology Data": DropRecordings.R

```
#' Drop specified recordings from an ERGExam Object
#'
#' This method returns a new \code{ERGExam} object with specific recordings removed.
#'
#' @inheritParams Get
#' @param where A \link[base:list]{base::list} defining selection criteria to identify recordings to remove.
#' @inheritParams Subset-method
#' @details The \code{DropRecordings} function creates a new \code{ERGExam} object excluding the recordings
#'          identified by the \code{where} criteria.
#' @seealso \link[EPhysData:Subset]{EPhysData::Subset}
#' @importFrom EPhysData Subset newEPhysSet Metadata
#' @importFrom methods validObject as
#' @name DropRecordings-method
#' @examples
#' data(ERG)
#' tmp<-DropRecordings(ERG, where=list(Channel="ERG", StimulusEnergy=1))
#' @exportMethod DropRecordings
setGeneric(
  name = "DropRecordings",
  def = function(X,
                 where) {
    standardGeneric("DropRecordings")
  }
)

#' @noMd
setMethod("DropRecordings",
          signature(X = "ERGExam"),
          function(X,
                   where = NULL) {

            # Identify recordings to exclude
            toExclude <- Where(X, where)
            if (is.numeric(toExclude)) {
              if (!all(toExclude %in% 1:nrow(Metadata(X)))) {
                stop("Not all values in 'where' are valid indices for data in 'X'.")
              }
              # Create a logical vector to select non-excluded recordings
              Recording <- !(1:length(X) %in% toExclude)
            } else {
              stop("Invalid result from 'Where'. Expected numeric indices.")
            }

            # subset measurements slot
            indexupdate <-
              as.data.frame(cbind(cumsum(Recording)[Recording], which(Recording)))
            colnames(indexupdate) <- c("new", "old")

            measurements<-X@Measurements@Measurements[X@Measurements@Measurements$Recording %in%  (1:length(Recording))[Recording],]

            measurements <-
              merge(measurements, indexupdate, by.x = "Recording", by.y = "old")
            measurements$Recording <- measurements$new
            measurements$new <- NULL

            X@Measurements@Measurements<-measurements
            newMeasurements<-X@Measurements
            newMetadata <- Metadata(X)[Recording, , drop = FALSE]
            newStimulus <- StimulusTable(X)[StimulusTable(X)$Step %in% newMetadata$Step, , drop = FALSE]

            # Create the new ERGExam object
            newX <- new("ERGExam",
                        Data = X@Data[Recording], # Assuming you want to subset Data this way
                        Metadata = newMetadata,
                        Stimulus = newStimulus,
                        Averaged = X@Averaged,
                        Measurements = newMeasurements,
                        ExamInfo = X@ExamInfo,
                        SubjectInfo = X@SubjectInfo,
                        Imported = X@Imported)
            if(validObject(newX)){
              return(newX)
            } else {
              stop("Invalid ERGExam object created.")
            }
          })
```
