## Supplemental Material 1 for "The ERGtools2 package: A Toolset for Processing and Analysing Visual Electrophysiology Data": ERGExam-data.R

```
#' Exemplary ERG Exam
#'
#' This data set contains an ERG Exam with ERG and OP channels from DA and LA flash and flicker stimuli..
#'
#' @docType data
#'
#' @name .SampleERGExam
#'
#' @usage data(ERG)
#'
#' @format An object of class \code{"ERGExam"}; see \linkS4class{ERGExam}.
#'
#' @keywords datasets
#'
#' @examples
#' data(ERG)
#' ERG<-SetStandardFunctions(ERG)
#' ERG
#' ggERGExam(ERG)
NULL
```
