## Supplemental Material 1 for "The ERGtools2 package: A Toolset for Processing and Analysing Visual Electrophysiology Data": ERGMeasurements-data.R

```
#' Exemplary ERG Measurements object
#'
#' This data set contains a Measurements object with Markers and corresponding measurements for an hypothetical ERG+VEP exam..
#'
#' @docType data
#'
#' @name .SampleERGMeasurements
#'
#' @usage data(Measurements.data)
#'
#' @format An object of class \code{"ERGMeasurements"}; see \linkS4class{ERGMeasurements}.
#'
#' @keywords datasets
#'
#' @examples
#' data(Measurements.data)
#' Measurements.data
#' Markers(Measurements.data)
NULL
```
