## Supplemental Material 1 for "The ERGtools2 package: A Toolset for Processing and Analysing Visual Electrophysiology Data": ERGtools2-package.html

ERGtools2-package.R


### ERGtools2-package.R

###### moritz

#### 2024-06-21

Structured Index and Introduction for the ERGtools2 R Package

This package contains an environment for working with
electroretinogram data. It contains an import method for Diagnosys
Espion data, but allows reading in of data also from other manufacturers
with limited coding effort. Standard procedures like averaging,
subsetting an visualization of individual exams are supported. This
package provides the 4class{ERGExam} class that stores data from a
single ERG examination. These may include different recording channels
(ERG, OP, VEP, …), or sequential recordings in repsonse to different
stimulus paradigms. It is usually generated from imported raw data using
. Data acuired from Diagnosys Espion can be imported directly using
.

The class 4class{ERGExam} expands , so .

```
## usethis namespace: start
```

@section Standard workflow:

After setting the functions, the accession methods inherited from
like and with the argument can be used to return the processed data.

@section Object creation: \* for
objects. \* , , , and for objects. \* and for objects. Experimental. \* for
objects. \* data(ERG) () Load example ERG recording \*
data(Measurements.data) () Load example Measurements data

@section Accession methods: \* Returns
the recording index for those recordings matching the given criteria. \*
This method subsets an (for object into a new object of the same class.
\* Returns data frame representing the or object in long format. When
used with the argument on an process (i.e. filtered, averaged) data is
returned. See also: .

- , , , : Returning information on the exam and the examined
  subject.
- , , , : Returning information on the Recordings contained in the
  dataset.
- Stimulus(): Returns selected rows of a stimulus table.
- StimulusDescription(), StimulusIntensity(),
  StimulusBackground(), StimulusType().

- Returns the Measurements table.

@section Processing: \* Update the
FilterFunction for all Recordings in an 4class{ERGExam}, or only those
slected using . \* Update the Rejected for all Recordings in an
4class{ERGExam}, or only those slected using . \* Update the
AverageFunction for all Recordings in an 4class{ERGExam}, or only those
slected using . \* This method is used to set standard functions for
processing 4class{ERGExam} data. It defines default functions for
averaging, filtering, and signal rejection based on the stimulus type. \*
Automatically sets markers depending on the channel (E.g. ERG, VEP,
OP,…) and stimulus type (Flash, FLicker). \* Place the a and B waves on
Flash ERG data stored in an an object. \* Place the N1 and P1 markers and
determines 1/frequency (period) for Flicker ERG data stored in an an
object. \* Place the P1, N1 and P2 markers for Flash VEP data stored in
an an object. \* \* : Interactive visual placement of markers.

@section Merging and other object
manipulation: \* (for objects) \* Add, update or remove Measurements from
an or object. \* \* \* StimulusDescription()<-,
StimulusIntensity()<-, StimulusBackground()<-, StimulusType()<-
\* Update or replace channel names. \* Clear the Measurements slots in an
object.

@section Plot methods: \* Generate a
plot for a single trace from an 4class{ERGExam} objects. \* Plot a
complete 4class{ERGExam} object. \* Uses to plot intensity sequence for
ERG exams \* Uses to plot step sequence (i.e. sequential recordings
within a single protocol) for ERG exams \* Uses ggplot2 to plot ERG
traces from multiple ERGExam objects \* : Interactive visual placement of
markers using a Shiny app.

TODO as.std.channelname(channel\_str, clear.unmatched = F)
is.std.channelname(channel\_str) erg\_str() op\_str() vep\_str() TODO
as.std.eyename(eye\_str) eye.haystack()od\_str() os\_str() TODO Save Load
TODO DropRecordings TODO RenameMarker TODO exploreERGExam

@examples # a typical workflow
data(ERG) # load example data StimulusTable(ERG) # have a look whats
inside Metadata(ERG) ERG<-SetStandardFunctions(ERG) ggERGTrace(ERG,
where = list( Step = as.integer(3), Eye = “RE”, Channel =“ERG”, Repeat =
as.integer(1))) # pick one and have a look at the traces as imported

ERG <- AutoPlaceMarkers(ERG, Channel.names = pairlist(ERG =
“ERG\_auto”)) # automatically place markers

ggERGTrace(ERG, where = list( Step = as.integer(3), Eye = “RE”,
Channel =“ERG”, Repeat = as.integer(1))) # pick one and have a look at
the traces as imported

@author @aliases ERGtools2-package @docType package @name ERGtools2-package

```
NULL
```

```
## NULL
```
