## Supplemental Material 1 for "The ERGtools2 package: A Toolset for Processing and Analysing Visual Electrophysiology Data": ERGtools2-package.R

```
#' Structured Index and Introduction for the ERGtools2 R Package
#'
#' This package contains an environment for working with electroretinogram data. It contains an import method for Diagnosys Espion data, but allows reading in of data coming in other formats with limited coding effort. Standard procedures like averaging, sub-setting an visualization of individual exams are supported.
#' This package provides the  \linkS4class{ERGExam} class that stores data from a single ERG examination. These may include different recording channels (ERG, OP, VEP, ...), or sequential recordings in response to different stimulus paradigms.
#' It is usually generated from imported raw data using \link[=newERGExam]{newERGExam}. Data acuired from Diagnosys Espion can be imported directly using \link[=ImportEspion]{ImportEspion}.\cr\cr
#'
#' The class \linkS4class{ERGExam} expands \link[EPhysData:EPhysSet]{EPhysData::EPhysSet}, so \strong{Methods that work on \link[EPhysData:EPhysSet]{EPhysData::EPhysSet} can be applied to \linkS4class{ERGExam} as well. See: \link[EPhysData:EPhysData-package]{EPhysData::EPhysData-package}}.
#'
## usethis namespace: start
#' @section Standard workflow:
#'
#' After setting the functions, the accession methods inherited from \link[EPhysData:EPhysSet-class]{EPhysData::EPhysSet-class} like \link[EPhysData:as.data.frame-method]{EPhysData::as.data.frame-method} and \link[EPhysData:GetData]{EPhysData::GetData} with the argument \code{Raw=F} can be used to return the processed data.\cr\cr
#'
#' @section Object creation:
#' \tabular{lll}{
#'   \strong{Function/Method} \tab \strong{Classes} \tab \strong{Description} \cr
#'   \strong{\link[=newERGExam]{newERGExam}}  \tab \linkS4class{ERGExam}  \tab Create a new ERGExam object.\cr
#'   \link[=ImportEspion]{ImportEspion}, \link[=ImportEspionInfo]{ImportEspionInfo}, \link[=ImportEspionStimTab]{ImportEspionStimTab}, and \link[=ImportEspionMetadata]{ImportEspionMetadata} \tab \linkS4class{ERGExam}  \tab Import an ERG exam from a CSV file exported from the Diagnosys Espion software \cr
#'   \link[methods:new]{methods:new} and \link[=ImportEspionProtocol]{ImportEspionProtocol} \tab \link[=ERGProtocol]{ERGProtocol} \tab Experimental. Creates a container for ERG recording protocols as exported from exported the Diagnosys Espion software \cr
#'   \link[methods:new]{methods:new} \tab \linkS4class{ERGMeasurements} \tab Creates a container for ERG Markers and their positions on an ERG trace. Usually not necessary to be called by the user directly. \cr
#'   \link[=Save]{Save}, \link[=Load.ERGExam]{Load.ERGExam}  \tab \linkS4class{ERGExam} \tab Saves and loads the ERGExam object. \cr
#' }
#'
#' @section Accession methods:
#' \tabular{lll}{
#'   \strong{Function/Method} \tab \strong{Classes} \tab \strong{Description} \cr
#'   \strong{\link[=Where]{Where}} \tab \linkS4class{ERGExam}  \tab Returns the recording index for those recordings matching the given criteria. \cr
#'   \strong{\link[=Subset]{Subset}} \tab \linkS4class{ERGExam}  \tab Subsets an ERGExam object into a new object of the same class. \cr
#'   \strong{\link[=as.data.frame]{as.data.frame}} \tab \linkS4class{ERGExam} , \link[=ERGProtocol]{ERGProtocol} \tab Returns data frame representing the ERGExam or ERGProtocol object in long format. \cr
#'   \link[=Stimulus]{Stimulus} \tab \linkS4class{ERGExam}  \tab Returns selected rows of a stimulus table. \cr
#'   \link[=StimulusDescription]{StimulusDescription}, \link[=StimulusEnergy]{StimulusEnergy}, \link[=StimulusBackground]{StimulusBackground}, \link[=StimulusType]{StimulusType} \tab \linkS4class{ERGExam}  \tab Returns details of stimulus description, energy, background, and type. \cr
#'   \link[=MarkerNames]{MarkerNames} \tab \linkS4class{ERGExam} , \linkS4class{ERGMeasurements} \tab Returns the names of markers in the dataset. \cr
#'   \link[=Markers]{Markers} \tab \linkS4class{ERGExam} , \linkS4class{ERGMeasurements} \tab Returns marker information from the dataset. \cr
#'   \strong{\link[=Measurements]{Measurements}} \tab \linkS4class{ERGExam} , \linkS4class{ERGMeasurements} \tab Returns the Measurements table. \cr
#'   \link[=DOB]{DOB}, \link[=ExamDate]{ExamDate}, \link[=GroupName]{GroupName}, \link[=ProtocolName]{ProtocolName} \tab \linkS4class{ERGExam}  \tab Returns information on the exam and the examined subject. \cr
#'   \link[=Eyes]{Eyes}, \link[=Steps]{Steps}, \link[=Channels]{Channels}, \link[=Repeats]{Repeats} \tab \linkS4class{ERGExam}  \tab Returns information on the Recordings contained in the dataset. \cr
#' }
#'
#' @section Processing:
#' \tabular{lll}{
#'   \strong{Function/Method} \tab \strong{Classes} \tab \strong{Description} \cr
#'   \strong{\link[=SetStandardFunctions]{SetStandardFunctions}} \tab \linkS4class{ERGExam} \tab Sets standard functions for processing ERGExam data, defining default functions for averaging, filtering, and signal rejection based on the stimulus type. \cr
#'   \link[=FilterFunction<-]{FilterFunction<-}, \link[=Rejected<-]{Rejected<-}, \link[=AverageFunction<-]{AverageFunction<-} \tab \linkS4class{ERGExam} \tab Updates the FilterFunction, Rejected function or AverageFunction for all Recordings in an ERGExam, or only those selected using \code{where}. \cr
#'   \link[=AutoPlaceMarkers]{AutoPlaceVEP} \tab \linkS4class{ERGExam} \tab Automatically sets markers depending on the channel (e.g., ERG, VEP, OP) and stimulus type (Flash, Flicker). \cr
#'   \link[=AutoPlaceAB]{AutoPlaceAB}, \link[=AutoPlaceFlicker]{AutoPlaceFlicker}, \link[=AutoPlaceVEP]{AutoPlaceVEP}  \tab \link[EPhysData:EPhysData-class]{EPhysData::EPhysData-class} \tab Automatically sets markers for the respective type of channel (e.g., ERG, VEP, OP) and stimulus (Flash, Flicker). \cr
#'   \link[=CheckAvgFxSet]{CheckAvgFxSet} \tab \linkS4class{ERGExam} \tab Checks if the average function is set correctly for the dataset. \cr
#'   \strong{\link[=interactiveMeasurements]{interactiveMeasurements}} \tab \linkS4class{ERGExam} \tab Allows for interactive visual placement of markers. \cr
#' }
#'
#' @section Merging and other object manipulation:
#' \tabular{lll}{
#'   \strong{Function/Method} \tab \strong{Classes} \tab \strong{Description} \cr
#'   \link[=MergeERGExams]{MergeERGExams} \tab \linkS4class{ERGExam}  \tab Merges multiple ERGExam objects into one. \cr
#'   \link[=Measurements<-]{Measurements<-} \tab \linkS4class{ERGExam} , \link[=ERGMeasurements]{ERGMeasurements-class} \tab Adds, updates, or removes Measurements from an ERGExam or ERGMeasurements object. \cr
#'   \link[=ClearMeasurements]{ClearMeasurements} \tab \linkS4class{ERGExam}  \tab Clears the Measurements slots in an ERGExam object. \cr
#'   \link[=DropMarker]{DropMarker}, \link[=AddMarker]{AddMarker}, \link[=RenameMarker]{RenameMarker} \tab \linkS4class{ERGExam} \tab Methods for marker modification. \cr
#'   \link[=StimulusDescription()<-]{StimulusDescription()<-} , \link[=StimulusEnergy()<-]{StimulusEnergy()<-}, \link[=StimulusBackground()<-]{StimulusBackground()<-}, \link[=StimulusType()<-]{StimulusType()<-}, \tab \linkS4class{ERGExam} \tab Updates stimulus description, intensity, background, and type in the stimulus table. \cr
#'   \link[=UpdateChannelNames]{UpdateChannelNames} \tab \linkS4class{ERGExam} \tab Updates or replaces channel names in the dataset. \cr
#'   \link[=DropRecordings]{DropRecordings} \tab \linkS4class{ERGExam}  \tab Drops specified recordings from the ERGExam object. \cr
#'   \link[=as.std.channelname]{as.std.channelname},  \link[=is.std.channelname]{is.std.channelname}, \link[=erg_str]{erg_str}, \link[=op_str]{op_str}, \link[=vep_str]{vep_str} \tab Function \tab Standardizes channel names, checks if the channel name is standardized, returns standardized strings for ERG, OP, and VEP. \cr
#'   \link[=as.std.eyename]{as.std.eyename}, \link[=eye.haystack]{eye.haystack}, \link[=od_str]{od_str}, \link[=os_str]{os_str} \tab Function \tab Standardizes eye names, processes eye data, returns standardized strings for the right eye (OD) and left eye (OS). \cr
#' }
#'
#' @section Plot methods and interactive methods:
#' \tabular{lll}{
#'   \strong{Function/Method} \tab \strong{Classes} \tab \strong{Description} \cr
#'   \link[=ggERGTrace]{ggERGTrace} \tab \linkS4class{ERGExam} \tab Generates a ggplot plot for a single trace from an ERGExam object. \cr
#'   \strong{\link[=ggERGExam]{ggERGExam}} \tab \linkS4class{ERGExam} \tab Plots a complete ERGExam object. \cr
#'   \link[=ggIntensitySequence]{ggIntensitySequence} \tab \linkS4class{ERGExam} \tab Uses ggplot2 to plot intensity sequence for ERG exams. \cr
#'   \link[=ggStepSequence]{ggStepSequence} \tab \linkS4class{ERGExam} \tab Uses ggplot2 to plot step sequence (i.e., sequential recordings within a single protocol) for ERG exams. \cr
#'   \link[=ggPlotRecordings]{ggPlotRecordings} \tab \linkS4class{ERGExam} \tab Uses ggplot2 to plot ERG traces from multiple ERGExam objects. \cr
#'   \strong{\link[=interactiveMeasurements]{interactiveMeasurements}} \tab \linkS4class{ERGExam} \tab Interactive visual placement of markers using a Shiny app. \cr
#'   \strong{\link[=exploreERGExam]{exploreERGExam}} \tab \linkS4class{ERGExam} \tab Explores the ERGExam object interactively. \cr
#' }
#'
#' @examples
#' # a typical workflow
#' data(ERG) # load example data, to import own data, see the examples provided for newERGExam and ImportEspion
#' StimulusTable(ERG) # have a look whats inside
#' Metadata(ERG)
#' ERG<-SetStandardFunctions(ERG)
#' ggERGTrace(ERG, where = list( Step = as.integer(3), Eye = "RE", Channel ="ERG", Repeat = as.integer(1))) # pick one and have a look at the traces as imported
#'
#' ERG <- AutoPlaceMarkers(ERG, Channel.names = pairlist(ERG = "ERG_auto")) # automatically place markers
#'
#' ggERGTrace(ERG, where = list( Step = as.integer(3), Eye = "RE", Channel ="ERG", Repeat = as.integer(1))) # pick one and have a look at the traces as imported
#'
#' @author \href{https://www.lindnerlab.de}{Moritz Lindner}
#' @aliases ERGtools2-package
#' @docType _PACKAGE
#' @name ERGtools2-package
NULL
```
