## Supplemental Material 1 for "The ERGtools2 package: A Toolset for Processing and Analysing Visual Electrophysiology Data": exploreERGExam.R

```
#' Explore ERGExam Metadata and Measurements
#'
#' Launches a Shiny app to explore the content of the Metadata slot of the ERGExam object and allows for filtering by metadata columns Eye and Channel as well as by the Stimulus columns Description, StimulusEnergy, Background, and Type. Metadata and Stimulus are matched by the Step column in Metadata, which contains the index of the corresponding row of the data.frame in the Stimulus slot.
#'
#' @param X An ERGExam object.
#' @return A Shiny app to interactively explore ERGExam metadata and measurements.
#' @importFrom shiny fluidPage titlePanel sidebarLayout sidebarPanel conditionalPanel selectInput checkboxInput mainPanel h4 hr br req observeEvent reactive stopApp onSessionEnded shinyApp runApp plotOutput
#' @importFrom DT dataTableOutput renderDataTable datatable
#' @importFrom shinyjs useShinyjs extendShinyjs js
#' @export
#' @examples
#' \dontrun{
#' data(ERG)
#' exploreERGExam(ERG)
#' }
setGeneric(
  name = "exploreERGExam",
  def = function(X) {
    standardGeneric("exploreERGExam")
  }
)


#' @noMd
setMethod("exploreERGExam",
          "ERGExam",
          function(X) {
            shiny.style <- HTML(
              "
                                    .highlight-row {
                                      border: 2px solid #3498db;
                                      padding: 10px;
                                      border-radius: 5px;
                                      margin: 10px 0;
                                    }
                                 "
            )

            jscode <- "shinyjs.closeWindow = function() { window.close(); }"


            ui <- fluidPage(
              tags$head(tags$style(shiny.style)),
              useShinyjs(),
              extendShinyjs(text = jscode, functions = c("closeWindow")),
              titlePanel("Explore ERGExam Metadata"),
              sidebarLayout(
                sidebarPanel(
                  h4("Filter recordings"),
                  selectInput("eye", "Eye:", choices = c("All", unique(X@Metadata$Eye))),
                  selectInput("channel", "Channel:", choices = c("All", unique(
                    X@Metadata$Channel
                  ))),
                  selectInput("description", "Description:", choices = c("All", unique(
                    X@Stimulus$Description
                  ))),
                  selectInput("intensity", "StimulusEnergy:", choices = c("All", as.character(
                    unique(X@Stimulus$StimulusEnergy)
                  ))),
                  selectInput("background", "Background:", choices = c("All", unique(
                    X@Stimulus$Background
                  ))),
                  selectInput("type", "Type:", choices = c("All", unique(X@Stimulus$Type))),
                  hr(),
                  h4("Remove Recording"),
                  actionButton("removeRecBtn", "Selected only"),
                  br(),
                  "All of same...",
                  br(),
                  actionButton("removeEyeBtn", "Eye"),
                  actionButton("removeChBtn", "Channel"),
                  actionButton("removeStimBtn", "Stimulus Description"),
                  br(),
                  "...as the Selected.",
                  h4("Close"),
                  actionButton("CloseSaveBtn", "Close & Save"),
                  actionButton("CloseBtn", "Discard changes "),
                  h4("Settings"),
                  checkboxInput("showGraph", "Show Graph", value = FALSE),
                  width = 2
                ),
                mainPanel(
                  h4("Recordings stored in object"),
                  dataTableOutput("filteredData"),
                  hr(),
                  fluidRow(
                    column(
                      6,
                      h4("Measurements for selected recording"),
                      dataTableOutput("measurementsData"),
                    ),
                    column(
                      6,
                      conditionalPanel(
                        condition = "input.showGraph == true",
                        h4("Graph for selected recording"),
                        plotOutput("graphOutput")
                      )
                    )
                  )
                )
              )
            )

            metadata <- Metadata(X)
            stimulus <- Stimulus(X)
            measurements <-
              Measurements(X, TimesOnly = F, quiet = T)
            AnyMeasuremnts<-nrow(measurements)>0
            metadata$IDX <- 1:nrow(metadata)
            server <- function(input, output, session) {
              filteredData <- reactiveVal(metadata)

              observeEvent({
                input$eye
                input$channel
                input$description
                input$intensity
                input$background
                input$type
              }, {
                data <- metadata
                if (input$eye != "All") {
                  data <- data[data$Eye == input$eye,]
                }
                if (input$channel != "All") {
                  data <- data[data$Channel == input$channel,]
                }

                filteredStimulus <- stimulus
                if (input$description != "All") {
                  filteredStimulus <-
                    filteredStimulus[filteredStimulus$Description == input$description,]
                }
                if (input$intensity != "All") {
                  filteredStimulus <-
                    filteredStimulus[filteredStimulus$StimulusEnergy == as.numeric(input$intensity),]
                }
                if (input$background != "All") {
                  filteredStimulus <-
                    filteredStimulus[filteredStimulus$Background == input$background,]
                }
                if (input$type != "All") {
                  filteredStimulus <-
                    filteredStimulus[filteredStimulus$Type == input$type,]
                }

                filteredMetadata <-
                  data[data$Step %in% filteredStimulus$Step,]

                mergedData <-
                  merge(filteredMetadata,
                        filteredStimulus,
                        by = "Step",
                        all.x = TRUE)[, c(
                          "IDX",
                          "Channel",
                          "Eye",
                          "StimulusEnergy",
                          "Background",
                          "Type",
                          "Description",
                          "Repeat"
                        )]
                if (AnyMeasuremnts) {
                  measurementsCount <-
                    as.data.frame(table(measurements$Recording))
                  colnames(measurementsCount) <-
                    c("IDX", "MeasurementsCount")
                  # Merge the measurements count data frame with the mergedData data frame
                  mergedData <-
                    merge(
                      mergedData,
                      measurementsCount,
                      by.x = "IDX",
                      by.y = "IDX",
                      all.x = TRUE
                    )
                } else{
                  mergedData$MeasurementsCount <- 0
                }
                # Replace NA values in the Count column with 0
                mergedData$MeasurementsCount[is.na(mergedData$MeasurementsCount)] <-
                  0

                out<<-mergedData$IDX
                filteredData(mergedData)
              })

              output$filteredData <- renderDataTable({
                filteredData()
              }, selection = "single",
              options = list(dom = 't'))
              if (AnyMeasuremnts) {
                output$measurementsData <- renderDataTable({
                  req(input$filteredData_rows_selected)

                  selectedRow <-
                    as.numeric(filteredData()[input$filteredData_rows_selected, "IDX"])
                  selectedMeasurements <-
                    measurements[measurements$Recording == selectedRow, c("Name", "Relative", "Time", "Voltage")]

                  datatable(selectedMeasurements)
                }, selection = "single",
                options = list(dom = 't'))
              }

              output$graphOutput <- renderPlot({
                req(input$filteredData_rows_selected)
                req(input$showGraph)

                selectedRow <-
                  as.numeric(filteredData()[input$filteredData_rows_selected, "IDX"])
                ggERGTrace(X, where = selectedRow) #where = list(Recording = selectedRow))
              })
              # observe the remove buttons

              observeEvent(input$removeRecBtn, {
                if(!is.null(input$filteredData_rows_selected)){
                  selectedRow <- input$filteredData_rows_selected
                  updatedData <- filteredData()[-selectedRow, ]
                  filteredData(updatedData)
                } else {
                  showModal(
                    modalDialog(
                      title = "Nothing selected.",
                      "No Recording has been selected.",
                      easyClose = TRUE,
                      footer = NULL
                    )
                  )
                }
              })

              observeEvent(input$removeChBtn, {
                if(!is.null(input$filteredData_rows_selected)){
                  selectedRow <- input$filteredData_rows_selected
                  selectedCh <- filteredData()[selectedRow, "Channel"]
                  selectedRow <- filteredData()[filteredData()$Channel == selectedCh, "IDX"]
                  updatedData <- filteredData()[!(filteredData()$IDX %in% selectedRow), ]
                  filteredData(updatedData)
                } else {
                  showModal(
                    modalDialog(
                      title = "Nothing selected.",
                      "No Recording has been selected.",
                      easyClose = TRUE,
                      footer = NULL
                    )
                  )
                }
              })

              observeEvent(input$removeEyeBtn, {
                if(!is.null(input$filteredData_rows_selected)){
                  selectedRow <- input$filteredData_rows_selected
                  selectedEye <- filteredData()[selectedRow, "Eye"]
                  selectedRow <- filteredData()[filteredData()$Eye == selectedEye, "IDX"]
                  updatedData <- filteredData()[!(filteredData()$IDX %in% selectedRow), ]
                  filteredData(updatedData)
                } else {
                  showModal(
                    modalDialog(
                      title = "Nothing selected.",
                      "No Recording has been selected.",
                      easyClose = TRUE,
                      footer = NULL
                    )
                  )
                }
              })

              observeEvent(input$removeStimBtn, {
                if(!is.null(input$filteredData_rows_selected)){
                  selectedRow <- input$filteredData_rows_selected
                  selectedStim <- filteredData()[selectedRow, "Description"]
                  selectedRow <- filteredData()[filteredData()$Description == selectedStim, "IDX"]
                  updatedData <- filteredData()[!(filteredData()$IDX %in% selectedRow), ]
                  filteredData(updatedData)
                } else {
                  showModal(
                    modalDialog(
                      title = "Nothing selected.",
                      "No Recording has been selected.",
                      easyClose = TRUE,
                      footer = NULL
                    )
                  )
                }
              })

              # Observe the finish button
              observeEvent(input$CloseBtn, {
                showModal(modalDialog(
                  title = "Close",
                  "Do you want to close without saveing?",
                  footer = tagList(
                    actionButton("confirmClose", "Close"),
                    modalButton("Cancel")
                  )
                ))
              })
              observeEvent(input$confirmClose, {
                js$closeWindow()
                stopApp(out)
              })
              observeEvent(input$CloseSaveBtn, {
                showModal(modalDialog(
                  title = "Close & Save",
                  "You will now close the dialog and return an updated ERGExam object",
                  footer = tagList(
                    actionButton("confirmSaveClose", "Close & Save"),
                    modalButton("Cancel")
                  )
                ))
              })
              observeEvent(input$confirmSaveClose, {
                out<-filteredData()$IDX
                js$closeWindow()
                stopApp(out)
              })

              onSessionEnded(function() {
                js$closeWindow()
                stopApp(NULL)
              })

            }

            SetMarkersApp <- shinyApp(ui = ui, server = server)

            # run)
            out <- runApp(SetMarkersApp, launch.browser = T)
            if(!is.null(out)){
              Subset(X,where=out)
            } else {
              warning("exploreERGExam seems to have crashed or Browser Tab was closed. Returning original object.")
              X
            }
          })
```
