## Supplemental Material 1 for "The ERGtools2 package: A Toolset for Processing and Analysing Visual Electrophysiology Data": ExportImport.R

```
#' #' @importFrom xml2 xml_new_root
#'
#' setGeneric(
#'   name = "Export",
#'   def = function(X,
#'                  filename,
#'                  type,
#'                  overwrite)
#'   {
#'     standardGeneric("Export")
#'   }
#' )
#'
#' #' @noMd
#' setMethod("Export",
#'           "ERGExam",
#'           function(X,
#'                    filename,
#'                    type = "elvisml",
#'                    overwrite = F,
#'                    ExtraInfo = list()) {
#'
#'             doc = xml_new_root("ElVis", version = "1.0", created =  as.POSIXct(Sys.time()))
#'
#'             subjects <- xml_add_child(doc, "Subjects")
#'             subject <- xml_add_child(subjects, "Subject", id = Subject(X), normal = "true")
#'             xml_add_child(subject, "Lastname", Subject(X))
#'             xml_add_child(subject, "Birthdate", DOB(X))
#'             xml_add_child(subject, "Gender", X@SubjectInfo$Gender)
#'             xml_add_child(subject, "Group", GroupName(X))
#'
#'             if(length(ExtraInfo)==0){
#'               Notice(X,what = "E", notice_text = "")
#'             }
#'             protocol <- xml_add_child(protocols, "Protocol", id = "P001")
#'             xml_add_child(protocol, "Name", "Visual Acuity Test")
#'             xml_add_child(protocol, "Type", "Standard")
#'             xml_add_child(protocol, "Description", "A test for visual acuity")
#'             xml_add_child(protocol, "StimulusDevice", "Mini-Ganzfeld")
#'
#'
#'
#'             xml_structure(doc)
#'
#'
#'             con <- url("https://static-content.springer.com/esm/art%3A10.1007%2Fs10633-017-9618-6/MediaObjects/10633_2017_9618_MOESM1_ESM.xsd")
#'             con <- file("/tmp/10633_2017_9618_MOESM1_ESM.xsd")
#'
#'
#'             schema <- read_xml(con, package = "xml2")
#'
#'             xml_validate(doc, schema) # https://link.springer.com/article/10.1007/s10633-017-9618-6#Sec21    https://www.r-bloggers.com/2017/01/using-xml-schema-and-xslt-in-r/
#'
#'           })
```
