## Supplemental Material 1 for "The ERGtools2 package: A Toolset for Processing and Analysing Visual Electrophysiology Data": Get-methods.R

```
#' Accession methods for metadata from ERGExam objects
#'
#' These methods are used to access metadata information from \linkS4class{ERGExam} objects.
#' @param X An \linkS4class{ERGExam}
#' @details These methods can be used to access metadata information stored in \linkS4class{ERGExam} objects. \cr \cr
#' @return A vector. For 'StimulusTable()' a data.frame and a function for 'GetFilterFunction()' and 'GetAverageFunction()'.
#' FIXME: Return is not up to date
#' FIXME not only for Class ERGExma

#' @examples
#' # Get Data to work with
#' data(ERG)
#'
#' # Accessing eyes from ERGExam object
#' Eyes(ERG)
#'
#' # Accessing channels from ERGExam object
#' Channels(ERG)
#'
#' # Accessing steps from ERGExam object
#' Steps(ERG)
#'
#' # Accessing steps from ERGExam object
#' Repeats(ERG)
#'
#' # Accessing subject from ERGExam object
#' Subject(ERG)
#'
#' # Accessing protocol name from ERGExam object
#' ProtocolName(ERG)
#'
#' # Accessing group name from ERGExam object
#' GroupName(ERG)
#'
#' # Accessing exam date from ERGExam object
#' ExamDate(ERG)
#'
#' # Accessing date of birth from ERGExam object
#' DOB(ERG)
#'
#' @name Get
NULL

#' @describeIn Get Returns a the eyes of which the data has been recorded.
#' @exportMethod Eyes
setGeneric(
  name = "Eyes",
  def = function(X)
  {
    standardGeneric("Eyes")
  }
)
#' @noMd
setMethod("Eyes",
          "ERGExam",
          function(X) {
            unique(X@Metadata$Eye)
          })

#' @describeIn Get Returns the Channel names.
#' @exportMethod Channels
#' @noMd
setGeneric(
  name = "Channels",
  def = function(X)
  {
    standardGeneric("Channels")
  }
)
#' @noMd
setMethod("Channels",
          "ERGExam",
          function(X) {
            unique(X@Metadata$Channel)
          })
#' @noMd
setMethod("Channels",
          "ERGMeasurements",
          function(X) {
            unique(X@Marker$Channel)
          })

#' @describeIn Get Returns the steps of the exam
#' @exportMethod Steps
#' @noMd
setGeneric(
  name = "Steps",
  def = function(X)
  {
    standardGeneric("Steps")
  }
)
#' @noMd
setMethod("Steps",
          "ERGExam",
          function(X) {
            unique(X@Metadata$Step)
          })

#' @describeIn Get Returns the indices of individual repeats contained in an ERG exam.
#' @exportMethod Repeats
#' @noMd
setGeneric(
  name = "Repeats",
  def = function(X)
  {
    standardGeneric("Repeats")
  }
)
#' @noMd
setMethod("Repeats",
          "ERGExam",
          function(X) {
            res<-unique(X@Metadata$Repeat)
            if(length(res)!=1 || res!=1){
              warning("Using multiple Repeats is still experimental. When creating the ERGExam, set Repeats to 1, if not needed.")
            }
            return(res)
          })

#' @describeIn Get Returns the subject's name
#' @exportMethod Subject
#' @noMd
setGeneric(
  name = "Subject",
  def = function(X)
  {
    standardGeneric("Subject")
  }
)
#' @noMd
setMethod("Subject",
          "ERGExam",
          function(X) {
            X@SubjectInfo$Subject
          })

#' @describeIn Get Returns the measurement parameter names (e.g: 'a','B','N1','P1').
#' @family ERGMeasurements functions
#' @family ERGExam functions
#' @exportMethod MarkerNames
#' @noMd
setGeneric(
  name = "MarkerNames",
  def = function(X)
  {
    standardGeneric("MarkerNames")
  }
)
#' @noMd
setMethod("MarkerNames",
          "ERGMeasurements",
          function(X) {
            unique(X@Marker$Name)
          })
#' @noMd
setMethod("MarkerNames",
          "ERGExam",
          function(X) {
            MarkerNames(X@Measurements)
          })

#' @describeIn Get Returns the recording protocol name.
#' @exportMethod ProtocolName
#' @noMd
setGeneric(
  name = "ProtocolName",
  def = function(X)
  {
    standardGeneric("ProtocolName")
  }
)

#' @noMd
setMethod("ProtocolName",
          "ERGExam",
          function(X) {
            X@ExamInfo$ProtocolName
          })

#' @describeIn Get Returns the group name.
#' @exportMethod GroupName
#' @noMd
setGeneric(
  name = "GroupName",
  def = function(X)
  {
    standardGeneric("GroupName")
  }
)

#' @noMd
setMethod("GroupName",
          "ERGExam",
          function(X) {
            X@SubjectInfo$Group
          })

#' @describeIn Get Returns the exam date.
#' @exportMethod ExamDate
#' @noMd
setGeneric(
  name = "ExamDate",
  def = function(X)
  {
    standardGeneric("ExamDate")
  }
)

#' @noMd
setMethod("ExamDate",
          "ERGExam",
          function(X) {
            X@ExamInfo$ExamDate
          })

#' @describeIn Get Returns the date of birth.
#' @exportMethod DOB
#' @noMd
setGeneric(
  name = "DOB",
  def = function(X)
  {
    standardGeneric("DOB")
  }
)

#' @noMd
setMethod("DOB",
          "ERGExam",
          function(X) {
            X@SubjectInfo$DOB
          })
```
