## Supplemental Material 1 for "The ERGtools2 package: A Toolset for Processing and Analysing Visual Electrophysiology Data": ggERGExam.R

```
#' Generates a ggplot2 plot from an ERGExam object.
#'
#' This function creates a \link[ggplot2:ggplot]{ggplot2:ggplot} plot of an ERGExam object, including line plots of
#' stimulus response curves for different label categories.
#'
#' @param X An ERGExam object.
#' @param return.as Whether to return as a \link[gridExtra:grid.arrange]{gridExtra::grid.arrange} grid (the default: 'return.as = "grid"') or as a list of \link[ggplot2:ggplot]{ggplot2:ggplot}s ('return.as = "list"').
#' @param show.markers Whether to return Marker Postitions as stored in the Measurements slot.
#' @param SetSIPrefix Change the SI prefix. Set to \code{keep}, for not to change anything, to \code{auto} (default) for using the \link[EPhysData:BestSIPrefix-methods]{EPhysData:BestSIPrefix-methods} to minimize the number of relevant digits or to any SI prefix to use that. Calls the \link[EPhysData:SetSIPrefix-methods]{EPhysData:SetSIPrefix-methods}.
#' @param downsample 	Integer giving the desired number of intervals for downsampling. Non-integer values are rounded down. Defaults to 250.
#'
#' @return Depending on the \code{return.as} parameter, this function returns either a \link[gridExtra:grid.arrange]{gridExtra::grid.arrange} object containing the plot(s) or a list of \link[ggplot2:ggplot]{ggplot2:ggplot} objects.
#'
#' @details The function arranges the plots based on the data characteristics:
#' \itemize{
#'   \item **Rows:** If there are multiple channels (e.g., ERG, VEP) recorded, each channel is plotted in a separate row. If the `ERGExam` object contains data from both eyes, separate rows will also be used for each eye.
#'   \item **Columns:** The plots are arranged into columns based on the adaptation state (e.g., dark-adapted, light-adapted) and stimulus type.
#' }
#' If both channels and eye recordings are present, the plot will use a facet grid with channels in rows and eyes in columns. If only one eye or one channel is present, a single plot is returned.
#'
#' @export
#' @examples
#' # Example usage:
#' data(ERG)
#' ERG<-SetStandardFunctions(ERG)
#' exam_plot <- ggERGExam(ERG)
#' print(exam_plot)
#' ggERGExam(ERG,SetSIPrefix="auto")
#' ggERGExam(ERG,SetSIPrefix="k")
#'
#' @importFrom ggplot2 ggplot aes geom_line facet_grid vars ggtitle facet_wrap element_line guide_legend
#' @importFrom ggpubr theme_pubr
#' @importFrom gridExtra grid.arrange
#' @importFrom grid textGrob
#' @importFrom EPhysData as.data.frame lapply
#' @importFrom stringr str_detect
#' @importFrom units set_units
#' @importFrom tidyr %>%
#' @importFrom dplyr group_by mutate ungroup group_by_at row_number
#' @seealso \linkS4class{ERGExam} \link[ggplot2:ggplot]{ggplot2:ggplot}
#'
#' @exportMethod ggERGExam
setGeneric(
  name = "ggERGExam",
  def = function(X, return.as = "grid", show.markers = T, SetSIPrefix="auto", downsample = 250) {
    standardGeneric("ggERGExam")
  }
)
#' @noMd
setMethod(
  "ggERGExam",
  signature = "ERGExam",
  definition = function(X, return.as = "grid", show.markers = T, SetSIPrefix="auto", downsample = 250) {

    # internal fx
    count_down_Repeats <- function(df) {
      # Get all column names except "Repeat" and "Filename"
      column_names <- setdiff(names(df), c("Repeat", "Filename","Recording"))

      # Group by these columns, then mutate "Repeat" within each group
      df %>%
        group_by_at(column_names) %>%
        mutate(Repeat = row_number()) %>%
        ungroup()  # Make sure to ungroup at the end
    }

    # ggplot workaround
    Time<-Value<-Eye<-Channel<-colourby<-NULL

    stopifnot(CheckAvgFxSet(X))
    suppressMessages(X <- Downsample(X, n = 250))
    X <- Subset(X, Raw = F)
    X<-SetSIPrefix(X,SetSIPrefix)

    Metadata(X)<-as.data.frame(count_down_Repeats(Metadata(X)))

    dat <- as.data.frame(X)
    stimtab <- StimulusTable(X)
    dat <- merge(dat, stimtab, by = "Step")

    # Measurements
    if(show.markers){
      mes<-Measurements(X, measure.absolute = T)
      colnames(mes)[colnames(mes)=="Voltage"]<-"Value"
      mes<-merge(mes,Stimulus(X))
    }

    plotrows <- list()
    for (b in unique(dat$Background)) {
      for (t in unique(dat$Type[dat$Background == b])) {
        rows_to_keep <- which(dat$Background == b & dat$Type == t)
        curr <- dat[rows_to_keep, ]
        if (nrow(curr) > 0) {
          curr.mes <-mes[mes$Background == b &
                           mes$Type == t, ]
          if (any(curr$Repeat != 1)) {
            for (s in unique(curr$Step[curr$Background == b &
                                       curr$Type == t])) {
              tmp <- curr[curr$Step == s,]
              if (!(length(unique(tmp$Repeat)) > 1)) {
                curr$Repeat[curr$Background == b & curr$Type == t] <- 1
              }
            }
          }
          curr <- curr[curr$Repeat == 1,]

          defining <-
            unique(curr[, c("StimulusEnergy", "Description")])
          defining <-
            apply(defining, 2, function(x) {
              length(unique(x))
            }) == nrow(defining)
          if (defining["StimulusEnergy"]) {
            colourby = "StimulusEnergy"
          } else{
            colourby = "Description"
          }
          curr$colourby <- as.ordered(curr[, colourby])
          ID <- paste0(b, "-", t)
          plotrows[[ID]] <- ggplot(curr,
                                   aes(x = Time,
                                       y = Value,
                                       colour = colourby)) +
            geom_line() +
            #ggtitle(paste0(Subject(X), ", ", ExamDate(X), "\n", ProtocolName(X))) +
            guides(color = guide_legend(title = colourby)) +
            theme_pubr(base_size = 8)
            if (length(unique(curr$Channel)) * length(unique(curr$Eye)) > 1) {
              plotrows[[ID]] <- plotrows[[ID]] +
              facet_grid(Channel ~ Eye, scales = "free")
            }
            plotrows[[ID]] <- plotrows[[ID]] +
              ggtitle((ID))

          # add Measurements

          if(show.markers){
            if(nrow(curr.mes)>0){
              # Measurements
              if (show.markers) {
                plotrows[[ID]] <- plotrows[[ID]] +
                  geom_point(data = curr.mes, aes(group = Name), color = "black")
              }
            }
          }
        }
      }

    }
    # repeated recordings
    repeated <-
      unique(dat[dat$Repeat > 1, c("Step", "Channel")])
    if (nrow(repeated) > 0) {
      message("Repeated recordings detected")
      for (r in 1:nrow(repeated)) {
        curr <-
          dat[dat$Step == repeated$Step[r] &
                dat$Channel == repeated$Channel[r],]
        if (length(unique(curr$Repeat)) > 1) {
          ID <-
            paste0("Repeated: ",
                   unique(curr$Description),
                   "-",
                   unique(curr$Channel))
          plotrows[[ID]] <- ggplot(curr,
                                   aes(
                                     x = Time,
                                     y = Value,
                                     colour = as.ordered(Repeat)
                                   )) +
            geom_line() +
            #ggtitle(paste0(Subject(X), ", ", ExamDate(X), "\n", ProtocolName(X))) +
            theme_pubr(base_size = 8)
          if (length(unique(curr$Eye)) > 1) {
            plotrows[[ID]]  <-
              plotrows[[ID]] + facet_grid(~ Eye, scales = "free")
          }

          plotrows[[ID]] <- plotrows[[ID]] + ggtitle((ID))
        }
      }
    }
    if(return.as == "grid"){
      if (length(plotrows) > 1) {
        return(grid.arrange(
          grobs = plotrows,
          ncol = round(length(plotrows)),
          top = textGrob(Subject(X))
        ))
      } else {
        return(plotrows[[1]])
      }
    }
    if(return.as == "list"){
      return(plotrows)
    }
  }
)
```
