## Supplemental Material 1 for "The ERGtools2 package: A Toolset for Processing and Analysing Visual Electrophysiology Data": ggERGTrace-methods.R

```
#' Generate a ggplot2 plot for a single item/trace from an ERGExam objects
#'
#' This method generates a \link[ggplot2:ggplot]{ggplot2::ggplot} plot for a single trace from an\linkS4class{ERGExam} objects.
#'
#' @inheritParams Subset-method
#' @param Raw Whether to show also raw traces in the background or not.
#' @param SetSIPrefix Change the SI prefix. Set to \code{keep}, for not to change anything, to \code{auto} (default) for using the \link[EPhysData:BestSIPrefix-methods]{EPhysData:BestSIPrefix-methods} to minimize the number of relevant digits or to any SI prefix to use that. Calls the \link[EPhysData:SetSIPrefix-methods]{EPhysData:SetSIPrefix-methods}.
#' @param Interactive Whether to return an interactive \link[plotly:ggplotly]{plotly::ggplotly}  graph
#' @return A \link[ggplot2:ggplot]{ggplot2::ggplot} plot visualizing the data from a single trace from an \linkS4class{ERGExam} object
#' @importFrom ggplot2 geom_label aes
#' @importFrom plotly ggplotly
#' @importFrom units deparse_unit set_units
#' @importFrom EPhysData ggEPhysData Rejected
#' @examples
#' data(ERG)
#' AverageFunction(ERG, where=pairlist(Step = as.integer(3),Channel = "ERG",Repeat = as.integer(1))) <- mean
#' ggERGTrace(ERG, where = list( Step = as.integer(3), Eye = "RE", Channel ="ERG", Repeat = as.integer(1)))
#' # the information obtained from the interactive plot can be used e.g. to update marker positions.
#' @name ggERGTrace
#' @exportMethod ggERGTrace
setGeneric(
  name = "ggERGTrace",
  def = function(X,
                 where,
                 Interactive = F,
                 SetSIPrefix="auto",
                 Raw = T) {
    standardGeneric("ggERGTrace")
  }
)

#' @noRd
setMethod("ggERGTrace",
          "ERGExam",
          function(X,
                   where,
                   Interactive = F,
                   SetSIPrefix="auto",
                   Raw = T) {
            which <-
              Where(X, where = where, expected.length = 1)

            if (length(which) == 0) {
              stop("No item matches selection.")
            }
            if (length(which) > 1) {
              stop("Multiple items match the selection criteria.")
            }
            sel <- X[[which]]

            un <- Measurements(X,
                               where = which)$Voltage[1]
            sel@Data <-
              set_units(sel@Data, deparse_unit(un), mode = "standard")
            out <- ggEPhysData(sel, Raw = Raw, SetSIPrefix = SetSIPrefix)
            mes<-Measurements(X,
                              where = which)
            if (any(mes$Recording == which)) {
              mes <-
                mes[mes$Recording == which, ]

              for (i in 1:nrow(mes)) {
                if (!is.na(mes$Relative[i])) {
                  mes$Voltage[i] <-
                    mes$Voltage[i] + mes$Voltage[mes$Name == mes$Relative[i]]
                }
              }


              if (!Interactive) {
                out <-
                  out + geom_label(
                    data = mes,
                    aes(x = Time, y = Voltage, label = Name),
                    colour = "darkblue",
                    inherit.aes = F
                  )
              }else{
                out <-
                  out + geom_text(
                    data = mes,
                    aes(x = Time, y = Voltage, label = Name),
                    colour = "darkblue",
                    inherit.aes = F
                  )
              }

            }
            if(!Interactive){
              return(out)
            }else{
              return(ggplotly(out))
            }

          })
```
