## Supplemental Material 1 for "The ERGtools2 package: A Toolset for Processing and Analysing Visual Electrophysiology Data": ggPlotList.R

```
#' Uses ggplot2 to plot intensity sequence for ERG exams
#'
#' This function generates a \link[ggplot2:ggplot]{ggplot2:ggplot} plot of intensity sequence data for ERG exams.
#'
#' @param List A list of ERG exams.
#' @param Background Background condition for the exams.
#' @param Type Type of exam (e.g., "Flash").
#' @param Channel The channel to plot (e.g., "ERG").
#' @param Markers Vector of markers to include in the plot.
#' @param Parameter The parameter to plot ("Amplitude" or "Time").
#' @param wrap_by Wrapping parameter for facetting ("Channel" or NULL).
#'
#' @return A ggplot2 plot object.
#'
#' @importFrom units as_units deparse_unit
#' @importFrom ggplot2 ggplot aes geom_line geom_point geom_errorbar scale_x_log10 facet_wrap labs guides
#' @importFrom ggpubr theme_pubr
#' @importFrom tidyr %>%
#' @importFrom dplyr summarise group_by n
#' @seealso  \link[ggplot2:ggplot]{ggplot2:ggplot}
#'
#' @examples
#' # Example usage:
#' data(ERG)
#' ERG<-SetStandardFunctions(ERG)
#' ERG <- AutoPlaceMarkers(ERG)
#' data <- list(ERG, ERG)
#' ggIntensitySequence(data, where= list(Background = "DA", Type = "Flash", Channel = "ERG"))
#'
#' @export
ggIntensitySequence <-
  function(List,
           where = list(Background = "DA",
                        Type = "Flash"),
           Markers = c("a", "B", "N1", "P1"),
           Parameter =  "Amplitude",
           wrap_by = "Channel",
           point.size = 1,
           theme.base.size = 8) {

    # dplyr workaround
    Group<-Name<-StimulusEnergy<-Voltage<-sd<-Time<-Amplitude<-sem<-ImplicitTime<-NULL

    # Extract Measurements and related info
    results <- CollectMeasurements(
      List = List,
      where = where,
      Markers = Markers
    )

    # Stats

    if (Parameter == "Amplitude") {
      results <-
        results %>% group_by(Group, Name, StimulusEnergy, Channel) %>% summarise(
          Amplitude = mean(Voltage),
          sem = sd(Voltage) / sqrt(n()),
          n =
            n()
        )
      results$sem <-
        as_units(results$sem, deparse_unit(results$Amplitude))
    }
    if(Parameter =="Time"){
      results <-
        results %>% group_by(Group, Name, StimulusEnergy, Channel) %>% summarise(
          ImplicitTime = mean(Time),
          sem = sd(Time, na.rm = T) / sqrt(n()),
          n =
            n()
        )

      results$sem <-
        as_units(results$sem, deparse_unit(results$ImplicitTime))

    }

    #results$Group<-paste0(results$Group," (n=",results$n,")")
    # PLot

    if (wrap_by == "Channel") {
      if(Parameter == "Amplitude"){
        plt <- ggplot(data = results,
                      aes(
                        x = StimulusEnergy,
                        y = Amplitude ,
                        color = Group,
                        shape = Name
                      )) +
          geom_line() +
          geom_point(size = point.size) +
          geom_errorbar(aes(ymin = Amplitude - sem, ymax = Amplitude + sem), width = 0.2)

      }
      if(Parameter == "Time"){
        plt <- ggplot(data = results,
                      aes(
                        x = StimulusEnergy,
                        y = ImplicitTime,
                        color = Group,
                        shape = Name
                      )) +
          geom_line() +
          geom_point(size = point.size) +
          geom_errorbar(aes(ymin = ImplicitTime - sem, ymax = ImplicitTime + sem), width = 0.2)

      }

      if (length(unique(results$Channel))!=1) {
        plt <- plt +
          facet_wrap(~ Channel,
                     scales = "free")
      }
      plt<-plt+
        theme_pubr(base_size = theme.base.size) +
        scale_x_log10() +
        labs(x = "Stimulus Energy [cd*s/m^2]", shape = "Marker")

      if(length(unique(results$Group))==1){
        plt<-plt+
          guides( color = "none")
      }
      if(length(unique(results$Name))==1){
        plt<-plt+
          guides( shape = "none")
      }
    }
    return(plt)
  }

#' @export
#' @noRd
PlotIntensitySequence<-ggIntensitySequence


#' Uses ggplot2 to plot step sequence for multiple ERG exams
#'
#' This function generates a \link[ggplot2:ggplot]{ggplot2:ggplot} plot of step sequence (i.e. sequential recordings within a single protocol) data for multiple ERG exams.
#'
#' @param List A list of ERG exams.
#' @inheritParams Where
#' @param Markers Vector of markers to include in the plot (e.g. c("a","B)).
#' @param wrap_by Wrapping parameter for facetting ("Channel" or NULL).
#'
#' @return A \link[ggplot2:ggplot]{ggplot2:ggplot} plot object.
#'
#' @importFrom ggplot2 ggplot aes geom_line geom_point geom_errorbar scale_x_log10 facet_wrap labs
#' @importFrom ggpubr theme_pubr
#' @importFrom tidyr %>%
#' @importFrom dplyr group_by summarise
#' @seealso \link[ggplot2:ggplot]{ggplot2:ggplot}
#'
#' @examples
#' # Example usage:
#' data(ERG)
#' ERG<-SetStandardFunctions(ERG)
#' ERG <- AutoPlaceMarkers(ERG)
#' data <- list(ERG, ERG)
#' ggStepSequence(data, where = list(Background = "DA", Type = "Flash", Channel = "ERG"), Markers = c("a", "B"))
#'
#' @export
ggStepSequence <-
  function(List,
           where = list(Background = "DA",
                        Type = "Flash"),
           Markers = c("N1", "P1"),
           wrap_by = "Channel")
  {

    #dplyr workaround
    Step<-Group<-Name<-StimulusEnergy<-Voltage<-sd<-Time<-Amplitude<-sem<-ImplicitTime<-NULL


    # Extract Measurements and related info
    results <- CollectMeasurements(
      List = List,
      where = where,
      Markers = Markers
    )

    # Stats

    results <-
      results %>% group_by(Group, Name, Step , Channel) %>% summarise(Amplitude = mean(Voltage),
                                                                      sem = sd(Voltage) / sqrt(n()),
                                                                      n=n())

    results$sem <-
      as_units(results$sem, deparse_unit(results$Amplitude))

    results$Group<-paste0(results$Group," (n=",results$n,")")

    # PLot

    if (wrap_by == "Channel") {
      plt <- ggplot(data = results,
                    aes(
                      x = Step,
                      y = Amplitude,
                      color = Group,
                      shape = Name
                    )) +
        geom_errorbar(aes(ymin = Amplitude - sem, ymax = Amplitude + sem), width = 0.2) +
        geom_point(size = 2) +
        facet_wrap( ~ Channel,
                    scales = "free") +
        theme_pubr(base_size = 8) +
        labs(x = "", shape = "Marker")
    }
    return(plt)
  }

#' @export
#' @noRd
PlotStepSequence<-ggStepSequence


#' Uses ggplot2 to plot ERG traces from multiple ERGExam objects
#'
#' This function generates a \link[ggplot2:ggplot]{ggplot2:ggplot} plot of ERG traces from multiple ERGExam objects.
#'
#' @param List A list of ERG exams.
#' @inheritParams Where
#' @inheritParams ggERGExam
#' @param wrap_by Wrapping parameter for facetting ("Channel" or NULL).
#' @param scales Passed on to \link[ggplot2:facet_grid]{ggplot2:facet_grid}.
#'
#' @return A ggplot2 plot object.
#'
#' @importFrom ggplot2 geom_hline geom_line facet_grid scale_color_manual theme geom_ribbon scale_fill_manual guides
#' @importFrom ggpubr theme_pubclean
#' @importFrom stringr str_detect
#' @importFrom units drop_units as_units
#' @importFrom cli cli_abort cli_inform cli_progress_bar cli_progress_update cli_progress_done
#' @examples
#' # Example usage:
#' data(ERG)
#' ERG<-SetStandardFunctions(ERG)
#' ERG <- AutoPlaceMarkers(ERG)
#' data <- list(ERG, ERG)
#' ggPlotRecordings(data, where = list(Background = "DA", Type = "Flash", Channel = "ERG"))
#'
#' @export
ggPlotRecordings <- function(List,
                             where = list(),
                             wrap_by = "Channel",
                             scales = "free_y",
                             downsample = 250){

  # ggplot workaround
  Time<-Value<-Eye<-Repeat<-NULL
  commoncolnames<-c("Step","Channel","Repeat","Eye","Repeat","Time")
  cli_inform(c(
    "Running {.fun ggPlotRecordings}. This may take a while for long ERGExam lists."
  ))

  results<-list()
  cli_progress_bar("Preparing data", total = length(List),  clear = TRUE)
  for (h in seq_along(List)) {
    x <- List[[h]]
    tryCatch({
      x<-Subset(x,where=where, Raw = T)
      x<-Downsample(x, n = downsample)
      x<-SetSIPrefix(x,"u")

      # SD
      sdev <- x
      AverageFunction(sdev, where = 1:length(sdev)) <- sd
      sdev <- Subset(sdev, Raw = F)
      df.sdev <- as.data.frame(sdev)
      df.sdev<-df.sdev[,c(commoncolnames,"Value")]
      colnames(df.sdev)[colnames(df.sdev)=="Value"]<-"sdev"

      df.sdev$n<-NA
      for (i in 1:length(x)) {
        curr.md <- Metadata(x)[i, ]
        df.sdev[c(
          df.sdev$Step == curr.md$Step &
          df.sdev$Channel == curr.md$Channel &
          df.sdev$Repeat == curr.md$Repeat &
          df.sdev$Eye == curr.md$Eye
        ),"n"] <- dim(x[[i]])[2]
      }

      # Avg as per set fx
      x <- Subset(x, Raw = F)
      df <- as.data.frame(x)
      df<-df[,c(commoncolnames,"Value")]

      df<-merge(df,df.sdev,by=commoncolnames)
      df$SEM[df$n>1]<-df$sdev[df$n>1]/df$n[df$n>1]
      df$SEM[df$n==1]<-NA
      df$SEM<-as_units(df$SEM,deparse_unit(df$Value))

      tab <- StimulusTable(x)
      df$Subject <- Subject(x)
      if (length(x@SubjectInfo$Group)==0) {
        x@SubjectInfo$Group<-"DEFAULT"
      }
      df$Group <- x@SubjectInfo$Group
      df$ExamDate <- min(x@ExamInfo$ExamDate)
      tab$Description <-
        iconv(tab$Description, "ASCII//TRANSLIT", sub = '')
      df <- merge(df, tab)
      results[[h]] <- df
    }, error = function (e){
      Notice(object,
             what = c("Error"),
             notice_text = c("x Gathering data failed with Error message: {.val {e}}"),
             help_page = "ERGtools2::ERGExam")
    })
    cli_progress_update()
  }
  cli_progress_done()

  results <- do.call(rbind.data.frame, results)
  # type conversion and column names
  colnames(results)[str_detect(colnames(results), "cd.s.m")] <-
    "StimulusEnergy"
  results$Step <- iconv(results$Step, "ASCII//TRANSLIT", sub = '')

  # define faceting variables
  get_non_unary_columns <- function(df, columns) {
    non_unary_columns <- columns[sapply(df[columns], function(x) length(unique(x)) > 1)]
    return(non_unary_columns)
  }
  rows <- get_non_unary_columns(results, c("Type", "Background", "StimulusEnergy", "Channel"))
  if(length(rows)==0){
    rows<-"1"
  }
  cols <- get_non_unary_columns(results, c("Group", "Subject"))
  if(length(cols)==0){
    cols<-"1"
  }
  facet_formula <- as.formula(paste(
    paste(rows, collapse = "+"),
    paste(cols, collapse = "+"),
    sep = " ~ "
  ))

  # plot
  ggplot(data = results, aes(
    x = Time,
    y = Value,
    ymin = Value - SEM,
    ymax = Value + SEM,
    color = Eye,
    fill = Eye,
    linetype = as.factor(Repeat)
  )) +
    geom_hline(yintercept = as_units(0,"uV"), colour = "gray") +
    geom_line() +
    geom_ribbon(alpha=0.2, colour=NA) +
    facet_grid(facet_formula,
               scales = scales
    ) +
    theme_pubclean(base_size = 8) +
    #labs(x = paste0("Time [", si_x, "]"), y = paste0("Voltage [", si_y, "]")) +
    scale_color_manual(values = c("RE" = "darkred", "LE" = "darkblue","OD" = "darkred", "OS" = "darkblue")) +
    scale_fill_manual(values = c("RE" = "darkred", "LE" = "darkblue","OD" = "darkred", "OS" = "darkblue")) +
    theme(panel.grid.major = element_line(linewidth = .1))+
    guides(fill = "none")
}

#' @export
#' @noRd
PlotRecordings<-ggPlotRecordings
```
