## Supplemental Material 1 for "The ERGtools2 package: A Toolset for Processing and Analysing Visual Electrophysiology Data": ImportEspion.R

```
#' Import ERG data and measurements from an Espion CSV file
#'
#'
#' This function imports ERG recordings from *.csv files exported from the Diagnosys Espion™ software and creates an \link[=ERGExam]{ERGExam} object.
#'
#' @param filename Path to the file. Currently supported \code{'.csv'} or \code{'.txt'} files exported from the Diagnosys Espion™ software that contain at least:
#'   An area with 1) a contents table, 2) a header table, 3) a stimulus table, and 4) a marker table.
#'   This function is only tested on files with the default horizontal table arrangement but should function for files with a vertical arrangement as well.
#'   Behaves similarly to the argument \code{file} from \link[utils:read.table]{utils::read.table()}.
#' @param sep The field separator character. Values on each line of the file are separated by this character.
#'   If sep = "" (the default for read.table), the separator is ‘white space’, which includes spaces, tabs, newlines, or carriage returns.
#' @param Import A list of character vectors specifying which parts of the data to import.
#'   Possible elements are "Raw" - for importing the raw recordings, "Averaged" - for importing the averaged recordings ("Results"), and "Measurements" - for importing the measured markers. Either of "Raw" or "Averaged" must be selected.
#' @param Protocol An S4 object of class \link[Protocol]{ERGProtocol()} or a list thereof.
#' @inheritParams Where
#' @inheritParams newERGExam
#' @seealso \linkS4class{ERGExam} \link[Protocol]{ERGProtocol()}
#'
#' @examples
#' \dontrun{
#' # Import a *.csv file exported from the Diagnosys Espion™ software.
#' ERG_Experiment <- ImportEspion("test.csv")
#' }
#'
#' @return For ImportEspion: A \linkS4class{ERGExam} object.
#'
#' @importFrom data.table fread
#' @importFrom units as_units
#' @importFrom utils read.csv txtProgressBar setTxtProgressBar
#' @importFrom stats na.exclude
#' @importFrom stringr str_detect str_remove str_trim str_replace_all regex
#' @importFrom EPhysData newEPhysData
#' @importFrom cli cli_warn cli_abort cli_alert_info cli_progress_bar cli_progress_update cli_progress_done
#' @name ImportEspion
#' @export
ImportEspion <- function(filename,
                         sep = "\t",
                         Import = list ("Averaged", "Measurements"),
                         Protocol = NULL,
                         where = NULL,
                         skip.validation = F) {
  #(NA.EXCLUDE REMOVED!!!! DOES THAT WORK??)
  message(paste("Importing", filename))

  # Checks

  if (!file.exists(filename)) {
    cli_abort("File {.file {filename}} does not exist.")
  }
  if (sep != "\t") {
    cli_alert_info("Import of files using separaotrs other than '\t' is experimental.")
  }

  if (is.null(Protocol)) {
    cli_warn(c(
      "Provision of the protocol is recommended and may be essential if the marker table is not provided or does not include markers for each step and channel of the recording."
    ))
  } else{
    if (!inherits(Protocol, "ERGProtocol")) {
      if (is.list(Protocol)) {
        if (!(all(unlist(lapply(Protocol, function(x) {
          inherits(x, "ERGProtocol")
        }))))) {
          cli_abort(c(
            "'{.strong Protocol}' must be an object of class 'Protocol' or a list thereof."
          ))
        }
      } else{
        cli_abort(c(
          "'{.strong Protocol}' must be an object of class 'Protocol' or a list thereof."
        ))
      }
    }
  }

  contains_raw <- "Raw" %in% Import
  contains_averaged <- "Averaged" %in% Import
  if (!((contains_raw || contains_averaged) && !(contains_raw && contains_averaged))) {
    cli_abort(c(
      "Exactly one of '{.strong Raw}' and '{.strong Averaged}' must be selected for import."
    ))
  }

  if (!(all(Import %in% list("Raw", "Averaged", "Measurements")))) {
    cli_abort(c(
      "'Import' must be one or several of the following: '{.val Raw}', '{.val Averaged}', '{.val Measurements}'."
    ))
  }

  # load
  if (read.csv(filename,
               header = F,
               sep = sep,
               nrow = 1)[[1]] != "Contents Table") {
    cli_abort("{.file {filename}} does not begin with a table of content.")
  }
  # get Table of content
  toc <- get_toc(filename, sep = sep)

  required_tables <- c("Header Table", "Marker Table", "Stimulus Table", "Data Table")
  missing_tables <- required_tables[!(required_tables %in% rownames(toc))]

  if (length(missing_tables) > 0) {
    cli_abort(c(
      "'Header Table', 'Marker Table', 'Stimulus Table', and 'Data Table' must all be included in the data set (even if they should not be imported).",
      "The following table(s) are missing: {.val {missing_tables}}."
    ))
  }


  # get protocol info
  recording_info <- ImportEspionInfo(filename)
  required_fields <-
    c("Protocol",
      "Version",
      "Dateperformed",
      "Testmethod",
      "Animal",
      "DOB")
  missing_fields <-
    required_fields[!(required_fields %in% names(recording_info))]

  if (length(missing_fields) > 0) {
    cli_abort(
      c(
        "Table of content incomplete. Have these data been exported as anonymous? This is currently unsupported.",
        "The following field(s) are missing: {.val {missing_fields}}."
      )
    )

  }


  # Get Protocol info
  if (!is.null(Protocol)) {
    tmp1 <- sub(" \\[.*", "", recording_info$Protocol)
    if (is.list(Protocol)) {
      idx <- which(tmp1 == unlist(lapply(Protocol, function(x) {
        x@Name
      })))
      if (length(idx) == 0) {
        cli_abort(c("Required protocol not in the list."))
      }
      if (length(idx) > 1) {
        cli_abort(c("Duplicate protocol entry in the 'Protocols' list."))
      }
      Protocol <- Protocol[[idx]]
    } else{
      if (Protocol@Name != tmp1) {
        cli_abort(c("Provided protocol is not the required one."))
      }
    }
  }

  # Get stimulus information
  stim_info <- ImportEspionStimTab(filename, sep, Protocol)

  # get Metadata
  Metadata <-
    ImportEspionMetadata(filename, sep = sep, Protocol = Protocol)

  Metadata$Channel_idx<-Metadata$Channel
  Metadata$Channel<-Metadata$Channel_Name

  # prepare for loading only requested relevant
  if(!is.null(where)){
    where.idx <- Where.generic(Metadata,
                               nrow(Metadata),
                               stim_info,
                               where = where)
    if(length(where.idx)==0){
      cli_abort(c(
        "'{.strong where}' selection does not return any data for the given file.",
        "Try using {.run ImportEspionMetadata({filename})} and {.run ImportEspionStimTab({filename})} to find available key-value pairs for '{.strong where}'."
      ))
    }
  }else{
    where.idx<-1:nrow(Metadata)
  }

  # Get Data
  if (("Data Table" %in% rownames(toc))) {
    tmp <- toc # modify to only get header of data table
    tmp$Right <-
      tmp$Left + 5 # maximum width, if results are included
    Data_Header <-
      na.exclude(get_content(filename, tmp, "Data Table", sep = sep))
    Data_Header$Eye <- tmp$Eye
    rownames(Data_Header)<-NULL
    #Data_Header<-Data_Header[where.idx,]
    STEPS = vector("list", nrow(Data_Header))
  }

  if (("Data Table" %in% rownames(toc)) &&
      any(c("Raw", "Averaged") %in% Import)) {
    cli_progress_bar("Importing", total = dim(Data_Header)[1],  clear = TRUE, auto_terminate = T)
    for (i in 1:dim(Data_Header)[1]) {
      cli_progress_update()
      tryCatch({
        if (as.numeric(Data_Header$Chan[i]) == 1) {
          #if(i==1 || (Data_Header$Step[i]!=Data_Header$Step[i-1])){ # only the first for new step contains a time frame
          # get time trace
          TimeTrace <- get_trace(filename, toc, Data_Header, i, "TimeTrace")
          #}
        }
        if (i %in% where.idx){ # only import, if selected by where
          if (("Result" %in% colnames(Data_Header)) &&
              ("Averaged" %in% Import)) {
            # get Averages / "Results"
            resulttrace <-
              get_trace(filename, toc, Data_Header, i, "ResultTrace")

            STEPS[[i]] <-
              newEPhysData(Data = resulttrace,
                           TimeTrace = TimeTrace)
          }

          if ("Trials" %in% colnames(Data_Header) &&
              ("Raw" %in% Import)) {
            trialtraces <- get_trace(filename, toc, Data_Header, i, "TrialTrace")
            STEPS[[i]] <-
              newEPhysData(Data = trialtraces,
                           TimeTrace = TimeTrace)
          }
        }
      }, error = function (e) {
        cli_abort(c(
          "Importing data failed for file '{.file {basename(filename)}}' with the following error message:",
          "{.val {e}}",
          "Failed at Step '{.strong {Data_Header$Step[i]}}', Channel '{.strong {Data_Header$Channel[i]}}', Repeat '{.strong {Data_Header$Repeat[i]}}'."
        ))
      })
    }
    cli_progress_done()
  }
  STEPS<-STEPS[where.idx]

  #subset Metadata and stim_info to required
  Metadata<-Metadata[where.idx,]
  rownames(Metadata)<-NULL
  stim_info<-stim_info[stim_info$Step %in% Metadata$Step,]

  # Get Measurements
  if ("Measurements" %in% Import) {
    measurements <- get_measurements(filename, toc, sep)
    measurements$Relative <- NA
    md.tmp<-Metadata
    md.tmp$Recording <- 1:nrow(md.tmp)
    md.tmp$Channel<-md.tmp$Channel_idx
    measurements <-
      merge(measurements,
            md.tmp[, c("Step", "Channel", "Repeat", "Eye", "Recording","Channel_Name")],
            by = c("Step", "Channel", "Repeat", "Eye"))
    measurements$Channel<-measurements$Channel_Name
    measurements$Channel_Name<-NULL
    measurements$Channel_idx<-NULL
    rm(md.tmp)

    if (all(is.null(unique(measurements$Group)))) {
      measurements$Group <- NULL
    }
    if(length(unique(measurements$Group))==1){
      recording_info$Group <- unique(measurements$Group)
    }

    colnames(measurements)[colnames(measurements) == "Marker"] <-
      "Name"

    M <- newERGMeasurements(measurements, update.empty.relative = T)

    grp = unique(measurements$Group)

  } else{
    M <- newERGMeasurements(data.frame(Channel=character(),Name=character(),Recording=numeric(),Time=numeric(),Relative=character()))
    grp <- NULL
  }

  # Dump empty traces

  EmptyTraces<-unlist((lapply(STEPS, function(x) {
    sum(drop_units(x@Data))
  })))==0

  STEPS<-STEPS[!EmptyTraces]
  Metadata<-Metadata[!EmptyTraces,]
  rownames(Metadata)<-NULL
  stim_info<-stim_info[stim_info$Step %in% unique(Metadata$Step),]
  rownames(stim_info)<-NULL
  Metadata$Channel_Name<-NULL
  Metadata$Channel_idx<-NULL

  # Subject and Exam data

  DOB <-
    as.Date(as.character(recording_info$DOB), format =    "%d/%m/%Y", origin = "1970-01-01")

  ExamDate <-
    as.POSIXct(strptime(recording_info$Dateperformed, format =
                          "%d/%m/%Y %H:%M:%S"))
  ExamDate <-
    as.POSIXct.numeric(as.numeric(ExamDate), origin = "1970-01-01 00:00.00 UTC")

  # return the object
  newERGExam(
    Data = STEPS,
    Metadata = Metadata,
    Stimulus = stim_info,
    Measurements = M,
    ExamInfo = list(
      ProtocolName = recording_info$Protocol,
      Version = recording_info$Version,
      ExamDate = as.POSIXct(
        strptime(recording_info$Dateperformed, format =
                   "%d/%m/%Y %H:%M:%S")
      ),
      Filename = filename,
      RecMode = recording_info$Testmethod,
      Investigator = recording_info$Investigator
    ),
    SubjectInfo = list(
      Subject = recording_info$Animal,
      DOB = as.Date(as.character(recording_info$DOB), format =    "%d/%m/%Y", origin = "1970-01-01"),
      Gender = recording_info$Gender,
      Group = grp
    ),
    skip.validation = skip.validation
  )
}
```
