## Supplemental Material 1 for "The ERGtools2 package: A Toolset for Processing and Analysing Visual Electrophysiology Data": ImportEspionInfo.R

```
#' @describeIn ImportEspion Read the Exam information stored in an Espion CSV file
#' @examples
#' \dontrun{
#' # Import exam information from a *.csv file exported from the Diagnosys Espion™ software.
#' ImportEspionInfo("test.txt")
#' }
#'
#' @return For ImportEspionInfo: A named list containing the exam info stored in an exported Espion ERG exam.
#' @importFrom stringr str_remove_all
#' @importFrom utils read.csv
#' @importFrom cli cli_abort cli_alert_info
#' @export
ImportEspionInfo <- function(filename,
                             sep = "\t") {
  if (!file.exists(filename)) {
    stop("File ", filename, " does not exist")
  }
  if (sep != "\t") {
    cli_alert_info("Import of files using separaotrs other than '\t' is experimental.")
  }
  if (read.csv(filename,
               header = F,
               sep = sep,
               nrow = 1)[[1]] != "Contents Table") {
    cli_abort(c(
      "{.file {filename}} does not begin with a table of content."
    ))
  }

  # get Table of content
  toc <- get_toc(filename, sep = sep)

  if (!all(c("Header Table") %in% (rownames(toc)))) {
    cli_abort(c(
      "'{.strong Header Table}' must be included in the data set."
    ))
  }
  recording_info <-
    get_content(filename, toc, "Header Table", sep = sep)
  out.list<-as.list(recording_info$Value)
  names(out.list)<-str_remove_all(make.names(recording_info$Parameter,unique = T),"[.]")
  return(out.list)
}
```
