## Supplemental Material 1 for "The ERGtools2 package: A Toolset for Processing and Analysing Visual Electrophysiology Data": ImportEspionMetadata.R

```
#' @describeIn ImportEspion Read the Metadata information stored in an Espion Exam file
#' @examples
#' \dontrun{
#' # Import a *.txt file exported from the Diagnosys Espion™ software.
#' Metadata <- ImportEspionMetadata("test.txt")
#' }
#'
#' @return For ImportEspionMetadata: A \link[data.frame]{data.frame()} containing the metadata for an ERG Recording.
#' @importFrom units as_units
#' @importFrom utils read.csv
#' @importFrom cli cli_warn cli_abort cli_inform
#' @export
ImportEspionMetadata <- function(filename,
                                 sep = "\t",
                                 Protocol = NULL) {
  # Checks
  if (is.null(Protocol)) {
    cli_warn(c(
      "Provision of the protocol is recommended and may be essential if the marker table is not provided or does not include markers for each step and channel of the recording."
    ))
  } else{
    if (!inherits(Protocol, "ERGProtocol")) {
      if (is.list(Protocol)) {
        if (!(all(unlist(lapply(Protocol, function(x) {
          inherits(x, "ERGProtocol")
        }))))) {
          cli_abort(c(
            "'{.strong Protocol}' must be an object of class 'Protocol' or a list thereof."
          ))
        }
      } else{
        cli_abort(c(
          "'{.strong Protocol}' must be an object of class 'Protocol' or a list thereof."
        ))
      }
    }
  }

  if (!file.exists(filename)) {
    cli_abort("File {.file {filename}} does not exist.")
  }
  if (sep != "\t") {
    cli_inform("Import of files using separaotrs other than '\t' is experimental.")
  }

  # load
  if (read.csv(filename,
               header = F,
               sep = sep,
               nrow = 1)[[1]] != "Contents Table") {
    cli_abort("{.file {filename}} does not begin with a table of content.")
  }
  # get Table of content
  toc <- get_toc(filename, sep = sep)

  if (!c("Header Table") %in% (rownames(toc))) {
    cli_abort(c(
      "'{.strong Header Table}' must be included in the data set."
    ))
  }

  # get protocol info
  recording_info <- ImportEspionInfo(filename)
  if (!("Protocol" %in%  names(recording_info))) {
    cli_abort(c(
      "Table of content incomplete.",
      "Have these data been exported as anonymous? This is currently unsupported."
    ))

  }

  # Get Protocol info
  if (!is.null(Protocol)) {
    tmp1 <- sub(" \\[.*", "", recording_info$Protocol)
    if (is.list(Protocol)) {
      idx <- which(tmp1 == unlist(lapply(Protocol, function(x) {
        x@Name
      })))
      if (length(idx) == 0) {
        cli_abort(c(
          "Required protocol not in the list."
        ))
      }
      if (length(idx) > 1) {
        cli_abort(c(
          "Duplicate protocol entry in the 'Protocols' list."
        ))
      }
      Protocol <- Protocol[[idx]]
    } else{
      if (Protocol@Name != tmp1) {
        cli_abort(c(
          "Provided protocol is not the required one."
        ))
      }
    }
  }

  #import Metadata
  if ("Data Table" %in% rownames(toc)) {
    tmp <- toc # modify to only get header of data table
    tmp$Right <-
      tmp$Left + 5 # maximum width, if results are included
    Data_Header <-
      na.exclude(get_content(filename, tmp, "Data Table", sep = sep))
  }

  Metadata <- Data_Header[, c("Step", "Chan", "Result")]
  colnames(Metadata)[colnames(Metadata) == "Chan"] <- "Channel"
  colnames(Metadata)[colnames(Metadata) == "Result"] <- "Repeat"

  # if Protocol avaliable, get info from there, if not, try Measurements/Marker table
  if (!is.null(Protocol)) {
    Metadata$Eye <- "Unspecified"
    Metadata$LowFreqCutoff <- as_units(NA, "Hz")
    Metadata$HighFreqCutoff <- as_units(NA, "Hz")
    Metadata$Inverted <- FALSE
    Metadata$Channel_Name <- as.character(NA)
    for (i in 1:nrow(Metadata)) {
      Metadata$Channel_Name[i] <-
        Protocol@Step[[Metadata$Step[i]]]@Channels[[Metadata$Channel[i]]]@Name
      Metadata$Eye[i] <-
        Protocol@Step[[Metadata$Step[i]]]@Channels[[Metadata$Channel[i]]]@Eye
      Metadata$LowFreqCutoff[i] <-
        str_to_unit(Protocol@Step[[Metadata$Step[i]]]@Channels[[Metadata$Channel[i]]]@LowFreqCutoff)
      Metadata$HighFreqCutoff[i] <-
        str_to_unit(Protocol@Step[[Metadata$Step[i]]]@Channels[[Metadata$Channel[i]]]@HighFreqCutoff)
      Metadata$Inverted[i] <-
        Protocol@Step[[Metadata$Step[i]]]@Channels[[Metadata$Channel[i]]]@Inverted
    }

    for (c in unique(Metadata$Channel)) {
      new.chname <-
        as.std.channelname(unique(Metadata$Channel_Name[Metadata$Channel == c]), clear.unmatched =
                             T)
      if (length(new.chname) > 1 & !all(is.na(new.chname))) {
        new.chname <- new.chname[!is.na(new.chname)]
        if (length(new.chname) == 1) {
          cli_inform(c(
            "Several channel names detected for channel '{c}'.",
            "Inferring by markers to: '{new.chname}'."
          ))

        } else {
          relevantsteps <- unique(Metadata$Step[Metadata$Channel == c])
          curr.markers <- lapply(relevantsteps, function(x) {
            as.data.frame(Protocol@Step[[Metadata$Step[x]]]@Channels[[Metadata$Channel[c]]])$Marker.Name
          })
          new.chname <-
            inferre.channel.names.from.markers(unique(unlist(curr.markers)))
          cli_inform(c(
            "Several channel names detected for channel '{c}'.",
            "Inferring by markers to: '{new.chname}'."
          ))
        }
      }
      Metadata$Channel_Name[Metadata$Channel == c] <- new.chname
    }

  } else {
    # if marker table contained in file
    if (!c("Marker Table" %in% (rownames(toc)))) {
      cli_abort(c(
        "'{.strong Marker Table}' must be included in the data set if the protocol is missing."
      ))
    }
    measurements <- get_measurements(filename, toc, sep)
    measurements$Channel_Name <- as.character(NA)
    for (c in unique(measurements$Channel)) {
      curr.markers <- unique(measurements$Marker[measurements$Channel == c])
      measurements$Channel_Name[measurements$Channel == c] <-
        inferre.channel.names.from.markers(curr.markers)
    }
    Metadata$id  <- 1:nrow(Metadata)
    Metadata <-
      merge(
        Metadata,
        unique(measurements[, c("Step", "Channel", "Repeat", "Eye", "Channel_Name")]),
        by = c("Step", "Channel", "Repeat"),
        all.x = T
      )
    Metadata <- Metadata[order(Metadata$id),]
    Metadata$id<-NULL
    Metadata$Eye <- as.std.eyename(Metadata$Eye, warn.only=T)
  }

  return(Metadata)
}
```
