## Supplemental Material 1 for "The ERGtools2 package: A Toolset for Processing and Analysing Visual Electrophysiology Data": ImportEspionProtocol.R

```
#' Import an Espion stimulus protocol from a CSV file.
#'
#' This function imports an Espion stimulus protocol from a CSV file and returns it as a Protocol S4 object.
#'
#' @param filename The path to the CSV file containing the protocol data.
#' @return An instance of the Protocol S4 class.
#'
#' @importFrom data.table fread
#' @importFrom stringr str_detect str_remove str_trim str_replace_all regex
#' @importFrom methods new
#' @export
ImportEspionProtocol<-function(filename){
  Header<-fread(filename,sep = ",",header = F,fill=T,strip.white = T, nrows=85, data.table=F)
  Name=getSingleColumnChar(Header,"Protocol Name")
  Exported<-as.POSIXct(getSingleColumnChar(Header,"Export date"))
  nSteps=getSingleColumnNumeric(Header,"Total steps")
  nChannels=getSingleColumnNumeric(Header,"Total channels")

  message("Methods handling Espion Protocols are experimental.")
  # import Step Parameters
  rn <-
    fread(
      filename,
      sep = ",",
      header = F,
      fill = T,
      skip=85,
      strip.white = T,
      select = c(1),
      data.table = F
    )$V1

  StepParam <-
    getTable(
      filename,
      start = which(rn == "Step Parameters") + 85,
      end = which(rn == "")[1] + 85,
      nCols = nSteps
    )
  keep <-
    c(
      "Description",
      "Adaptation required",
      "Results per run",
      "Time between results",
      "Sample frequency",
      "Sweep pre-trigger time",
      "Sweep post trigger time",
      "Inter sweep delay",
      "Sweeps per result",
      "Min no. of sweeps/results",
      "Glitch removal",
      "Drift removal",
      "Baseline enabled",
      "Baseline pre-trigger",
      "Baseline range"
    )
  StepParam<-StepParam[keep, , drop=F]
  StepParam<-unlist(apply(StepParam, 2, list), recursive = F, use.name = F)
  StepParam<-lapply(StepParam, function(x) {
    x <- as.data.frame(x, col.names = FALSE)
  })

  # import Channel Parameters

  ChannelList<-list()
  MarkerList<-list()
  for (i in 1:nChannels) {
    start <- which(rn == paste0("Chan ", i, " Parameters")) + 85
    end <- if (any(rn == paste0("Chan ", i + 1, " Parameters"))) {
      which(rn == paste0("Chan ", i + 1, " Parameters")) + 85
    } else {
      length(rn) + 85
    }

    ChParam <-
      getTable(filename,
               start = start[1],
               end = end[1],
               nCols = nSteps)
    markers_start <-
      which(str_detect(rownames(ChParam), "Marker"))
    markers_names <-
      which(str_detect(rownames(ChParam), "Name"))
    if (length(markers_names)>1){
      markers_names <- markers_names[markers_names > markers_start]
      markers_relto <-
        which(str_detect(rownames(ChParam), "Amplitude is relative to"))


      markers_names <- ChParam[markers_names, , drop=F]
      markers_relto <- ChParam[markers_relto, , drop=F]
    }else{
      markers_names<-NULL
      markers_relto<-NULL
    }

    keep <-
      c(
        "Name",
        "Enabled",
        "Eye being tested",
        "Filter low frequency cutoff",
        "Filter high frequency cutoff",
        "Invert input polarity"
      )
    ChParam <- ChParam[keep, , drop=F]

    if(length(markers_names)>1){
      MarkerList[[i]]<-list(Names=markers_names,RelTo=markers_relto)
    } else {
      MarkerList[[i]]<-list(Names=NULL,RelTo=NULL)
    }
    ChannelList[[i]]<-as.data.frame(ChParam)
  }

  # convert into S4 Structure
  Protocol<-new("ERGProtocol")
  Protocol@Name <- enc2utf8(Name)
  Protocol@Export_Date<-Exported
  Protocol@nSteps<-nSteps
  Protocol@nChannels<-nChannels

  Steps<-list()
  for (s in 1:Protocol@nSteps){
    currStep<-new("ERGStep")
    currStep@Description <-
      str_replace_all(enc2utf8(StepParam[[s]]$x[rownames(StepParam[[s]]) == "Description"]), regex("[^A-Za-z0-9]+"), " ")
    currStep@Adaptation <-
      StepParam[[s]]$x[rownames(StepParam[[s]]) == "Adaptation required"]

    currStep@Resultsperrun <-as.numeric(
      StepParam[[s]]$x[rownames(StepParam[[s]]) == "Results per run"]
    )

    currStep@Timebetweenresults <-
      StepParam[[s]]$x[rownames(StepParam[[s]]) == "Time between results"]
    currStep@Sweepsperresult <-as.numeric(
      StepParam[[s]]$x[rownames(StepParam[[s]]) == "Sweeps per result"]
    )
    currStep@Sweepsperresultmin <- as.numeric(
      StepParam[[s]]$x[make.names(rownames(StepParam[[s]])) == "Min.no..of.sweeps.results"]
    )
    currStep@Glitchremoval <-
      StepParam[[s]]$x[rownames(StepParam[[s]]) == "Glitch removal"]=="On"
    currStep@Driftremoval <-
      StepParam[[s]]$x[rownames(StepParam[[s]]) == "Drift removal"]=="On"
    currStep@Baselineenabled <-
      StepParam[[s]]$x[rownames(StepParam[[s]]) == "Baseline enabled"]=="On"
    currStep@Baselinepretrigger <-
      StepParam[[s]]$x[rownames(StepParam[[s]]) == "Baseline pre-trigger"]
    currStep@Baselinerange <-
      StepParam[[s]]$x[rownames(StepParam[[s]]) == "Baseline range"]

    currStep@SampleFrequency <-
      StepParam[[s]]$x[rownames(StepParam[[s]]) == "Sample frequency"]
    currStep@PreTriggerTime <-
      StepParam[[s]]$x[rownames(StepParam[[s]]) == "Sweep pre-trigger time"]
    currStep@PostTriggerTime <-
      StepParam[[s]]$x[rownames(StepParam[[s]]) == "Sweep post trigger time"]
    currStep@InterSweepDelay <-
      StepParam[[s]]$x[rownames(StepParam[[s]]) == "Inter sweep delay"]

    for (c in 1:Protocol@nChannels) {
      currChan<-new("ERGChannel")
      curr<-ChannelList[[c]][,s, drop=F]
      currChan@Name<-curr["Name",1]
      orig.ch.name<-currChan@Name
      currChan@Name<-as.std.channelname(orig.ch.name)
      currChan@Eye<-tryCatch({
        as.std.eyename(curr["Eye being tested",1])
      }, error = function (e){
        inchanneldesc <- str_detect(orig.ch.name, eye.haystack())
        if(sum(inchanneldesc, na.rm = T) == 1){
          extr<-as.std.eyename(orig.ch.name, exact = F)
          message("Eye identifier ('", extr, "') in '", basename(filename), "', Step '", s, "', Channel '", c, "' (Name: '", currChan@Name, "') was extracted from original channel name ('", orig.ch.name, "'). ")
          extr
        } else {
          as.character(NA)
          warning("No valid Eye identifier in  '", basename(filename), "', Step '", s, "' Channel '", c, "'. Channel Name: '", currChan@Name, "'. Error message: ", e, ". Consider correcting manually. ")
        }
      })
      currChan@Enabled<-curr["Enabled",1]=="On"
      currChan@LowFreqCutoff<-curr["Filter low frequency cutoff",1]
      currChan@HighFreqCutoff<-curr["Filter high frequency cutoff",1]
      currChan@Inverted<-curr["Enabled",1]=="On"
      for (m in 1:length(MarkerList[[c]]$Names[, s])) {
        currMarker<-new("ERGMarker")
        if(!is.null(MarkerList[[c]]$Names[m, s])){
          currMarker@Name<-MarkerList[[c]]$Names[m, s]
          currMarker@RelativeTo<-MarkerList[[c]]$RelTo[m, s]
          currChan@Markers[[m]]<-currMarker
        }
      }

      if(!is.std.channelname(currChan@Name)){
        old<-currChan@Name
        inferred.name<-inferre.channel.names.from.markers(unlist(lapply(currChan@Markers, function(x){x@Name})))
        if(inferred.name!="Unknown"){
          currChan@Name<-inferred.name
        } else {
          if(currChan@Name==""){
            currChan@Name<-inferred.name
          }
        }
        message("No valid Channel name in  '", basename(filename), "', Step '", s, "' Channel '", c, "'. Original channel name '", old, "' was changed to ",  currChan@Name, ".")
      }

      currStep@Channels[[c]]<-currChan
    }
    Steps[[s]]<-currStep
  }
  Protocol@Step<-Steps

  return(Protocol)
}

#' @noMd
getSingleColumnChar<-function(table,key){
  value<-table[table$V1==key,]
  value<-value[1,!is.na(value[1,])]
  paste(unlist(value[,-1]), collapse = ', ',sep = "_")
}
#' @noMd
getSingleColumnNumeric<-function(table,key){
  value<-table[table$V1==key,]
  as.numeric(value[2])
}
#' @noMd
getTable<-function(filename,start,end,nCols){
  tab<-fread(
    filename,
    sep = ",",
    header = T,
    fill = T,
    skip=start-1,
    nrows=end-start,
    strip.white = T,
    blank.lines.skip=T,
    select = 1:(1+nCols),
    data.table = F
  )
  rownames(tab)<-make.unique(tab[,1])
  tab<-tab[,-1 , drop=F]
  return(tab)
}
```
