## Supplemental Material 1 for "The ERGtools2 package: A Toolset for Processing and Analysing Visual Electrophysiology Data": ImportEspionStimTab.R

```
#' @describeIn ImportEspion Read the Stimulus information stored in an Espion Exam file
#' @importFrom utils read.csv
#' @importFrom cli cli_warn cli_abort cli_inform
#' @examples
#' \dontrun{
#' # Import stimulus information from a *.csv file exported from the Diagnosys Espion™ software.
#' ImportEspionInfo("test.txt")
#' }
#' @return For ImportEspionStimTab: A \link[data.frame]{data.frame()} containing the stimulus information.
#' @export
ImportEspionStimTab <- function(filename,
                         sep = "\t",
                         Protocol = NULL) {
  # Checks
  if (!is.null(Protocol)) {
    if (!inherits(Protocol, "ERGProtocol")) {
      if (is.list(Protocol)) {
        if (!(all(unlist(lapply(Protocol, function(x) {
          inherits(x, "ERGProtocol")
        }))))) {
          cli_abort(c(
            "'{.strong Protocol}' must be an object of class 'Protocol' or a list thereof."
          ))
        }
      } else{
        cli_abort(c(
          "'{.strong Protocol}' must be an object of class 'Protocol' or a list thereof."
        ))
      }
    }
  }

  if (!file.exists(filename)) {
    stop("File ", filename, " does not exist")
  }
  if (sep != "\t") {
    cli_inform("Import of files using separaotrs other than '\t' is experimental.")
  }

  # load
  if (read.csv(filename,
               header = F,
               sep = sep,
               nrow = 1)[[1]] != "Contents Table") {
    cli_abort("{.file {filename}} does not begin with a table of content.")
  }
  # get Table of content
  toc <- get_toc(filename, sep = sep)

  missing_tables <- c("Header Table", "Stimulus Table")[!(c("Header Table", "Stimulus Table") %in% rownames(toc))]
  if (length(missing_tables) > 0) {
    cli_abort(c(
      "'{.strong Header Table}' and '{.strong Stimulus Table}' must be included in the data set (even if they should not be imported).",
      "The following table(s) are missing: {.val {missing_tables}}."
    ))
  }

  # get protocol info
  recording_info <- ImportEspionInfo(filename)
  if (!"Protocol" %in%  names(recording_info)) {
    cli_abort(c(
      "Table of content incomplete.",
      "Does not contain protocol name."
    ))
  }

  # Get Protocol info
  if (!is.null(Protocol)) {
    tmp1 <- sub(" \\[.*", "", recording_info$Protocol)
    if (is.list(Protocol)) {
      idx <- which(tmp1 == unlist(lapply(Protocol, function(x) {
        x@Name
      })))
      if (length(idx) == 0) {
        cli_abort("Required protocol not in the list.")
      }
      if (length(idx) > 1) {
        cli_abort("Duplicate protocol entry in the 'Protocols' list.")
      }
      Protocol <- Protocol[[idx]]
    } else{
      if (Protocol@Name != tmp1) {
        cli_abort("Provided protocol is not the required one.")

      }
    }
  }

  # Get stimulus information
  stim_info <- get_stim_info(filename, toc, sep)

  if (!is.null(Protocol)) {
    stim_info$PreTriggerTime<-as_units(NA,"s")
    stim_info$PostTriggerTime<-as_units(NA,"s")
    stim_info$InterSweepDelay<-as_units(NA,"s")
    stim_info$Baselinerange<-as_units(NA,"s")
    for (i in 1:nrow(stim_info)) {
      stim_info$Background[i] <-
        Protocol@Step[[stim_info$Step[i]]]@Adaptation
      stim_info$PreTriggerTime[i] <-
        str_to_unit(Protocol@Step[[stim_info$Step[i]]]@PreTriggerTime)

      stim_info$PostTriggerTime[i] <-
        str_to_unit(Protocol@Step[[stim_info$Step[i]]]@PostTriggerTime)

      stim_info$InterSweepDelay[i] <-
        str_to_unit(Protocol@Step[[stim_info$Step[i]]]@InterSweepDelay)
      stim_info$Driftremoval[i] <-
        Protocol@Step[[stim_info$Step[i]]]@Driftremoval
      stim_info$Baselineenabled[i] <-
        Protocol@Step[[stim_info$Step[i]]]@Baselineenabled
      stim_info$Baselinerange[i] <-
        str_to_unit(Protocol@Step[[stim_info$Step[i]]]@Baselinerange)
    }
  }

  return(stim_info)
}
```
