## Supplemental Material 1 for "The ERGtools2 package: A Toolset for Processing and Analysing Visual Electrophysiology Data": INT_as.std.X.R

```
#' Convert eye identifier strings to standard notation
#'
#' @param eye_str Character vector of eye identifier strings.
#' @param exact If \code{TRUE}: Require exact match. If \code{FALSE}: Whole word match is sufficient.
#' @param warn.only Invalid eye identifier strings will only cause a warning, not an error.
#' @return A character vector with standardized eye identifiers ('RE' for right eye, 'LE' for left eye, 'BE' for both eyes).
#' @examples
#' as.std.eyename(c("RE", "OD", "OS", "Right","OU"))
#' @export
as.std.eyename <- function(eye_str,
                           exact = T,
                           warn.only = F) {
  stopifnot(is.logical(exact))
  stopifnot(is.logical(warn.only))
  if (!is.std.eyename(eye_str, exact = exact, warn.only = warn.only)) {
    return(NULL)
  }
  if (exact) {
    # Exact match conversion
    eye_str <- unlist(lapply(eye_str, function(x) {
      if (x %in% od_str()) {
        return("RE")  # Right Eye
      }
      if (x %in% os_str()) {
        return("LE")  # Left Eye
      }
      if (x %in% ou_str()) {
        return("BE")  # Both Eyes
      }
      return(paste0("Unknown_", as.character(x)))  # Handle unknown values
    }))
  } else {
    # Inexact match conversion
    pattern <-
      paste0("(^|[^A-Za-z])(",
             paste(od_str(), collapse = "|"),
             ")([^A-Za-z]|$)")
    eye_str[grepl(pattern, eye_str)] <- "RE"

    pattern <-
      paste0("(^|[^A-Za-z])(",
             paste(os_str(), collapse = "|"),
             ")([^A-Za-z]|$)")
    eye_str[grepl(pattern, eye_str)] <- "LE"
  }
  return(eye_str)
}

#' @return For is.std.eyename: Logical vector indicating whether each element in \code{eye_str} contains a word that describes a standard eye name.
#' @examples
#' \dontrun{
#' is.std.eyename(c("RE", "BE", "LE", "Nonstandard"))
#' }
#' @describeIn as.std.eyename Check if eye name strings are standard eye names
#' @export
is.std.eyename <- function(eye_str,
                           exact = T,
                           warn.only = F) {
  stopifnot(is.logical(exact))
  stopifnot(is.logical(warn.only))
  if(!warn.only){
    stopifnot(is.character(eye_str))
  }
  if (exact) {
    if (!any(is.character(eye_str)) ||
        !all(eye_str %in% c(od_str(), os_str(), ou_str()))) {
      eye_str_nf <- eye_str[!(eye_str %in% c(od_str(), os_str()))]
      if (warn.only) {
        warning(paste0(eye_str_nf, sep = ", "),
                " is not a / are no valid eye identifier(s).")
      } else{
        stop(paste0(eye_str_nf, sep = ", "),
             " is not a / are no valid eye identifier(s).")
      }
      return(FALSE)
    }
  } else {
    # Inexact match can have a different implementation
    pattern <-
      paste0("(^|[^A-Za-z])(",
             paste(c(od_str(), os_str(), ou_str()), collapse = "|"),
             ")([^A-Za-z]|$)")
    if (!any(grepl(pattern, eye_str))) {
      cli_warn(c("No valid eye identifiers found in {.val {eye_str}}."))
      return(FALSE)
    }
  }
  return(TRUE)
}

#' Convert non-standard channel names to standard channel names
#'
#' @param channel_str Character vector of channel name strings to be converted.
#' @param clear.unmatched Logical indicating whether to clear unmatched strings.
#'
#' @return A character vector with standardized channel names.
#' @examples
#' \dontrun{
#' as.std.channelname(c("ERG_auto", "C-wave", "OPs", "Nonstandard"), clear.unmatched = TRUE)
#' }
#' @keywords internal
as.std.channelname<-function(channel_str, clear.unmatched=F){
  found<-is.std.channelname(channel_str)

  if(clear.unmatched){
    channel_str[!found]<-as.character(NA)
  }

  #ERG
  pattern <-
    paste0("(^|[^A-Za-z])(",
           paste(erg_str(), collapse = "|"),
           ")([^A-Za-z]|$)")
  channel_str[found][grepl(pattern,channel_str[found])]<-"ERG"

  #OP
  pattern <-
    paste0("(^|[^A-Za-z])(",
           paste(op_str(), collapse = "|"),
           ")([^A-Za-z]|$)")
  channel_str[found][grepl(pattern,channel_str[found])]<-"OP"

  #VEP
  pattern <-
    paste0("(^|[^A-Za-z])(",
           paste(vep_str(), collapse = "|"),
           ")([^A-Za-z]|$)")
  channel_str[found][grepl(pattern,channel_str[found])]<-"VEP"

  return(channel_str)
}

#' @return For is.std.channelname: Logical vector indicating whether each element in \code{channel_str} contains a word that describes a standard channel name.
#' @examples
#' \dontrun{
#' is.std.channelname(c("ERG_auto", "C-wave", "OPs", "Nonstandard"))
#' }
#' @describeIn as.std.channelname Check if channel name strings are standard channel names
#' @keywords internal
is.std.channelname <- function(channel_str) {
  if (!is.character(channel_str)) {
    stop(paste0(channel_str, sep = ", "),
         " is /are no character vectors.")
  }

  all_keywords <- c(erg_str(), op_str(), vep_str())

  pattern <-
    paste0("(^|[^A-Za-z])(",
           paste(all_keywords, collapse = "|"),
           ")([^A-Za-z]|$)")

  # Check if any word in channel_str matches the pattern
  return(grepl(pattern, channel_str))
}


#' @examples
#' \dontrun{
#' eye.haystack()
#' }#'
#' @describeIn as.std.eyename Return a vector containing all standard eye identifier strings
#' @noMd
#' @export
eye.haystack <- function() {
  return(c(od_str(),
           os_str(),
           ou_str()))
}

#' @examples
#' \dontrun{
#' od_str()
#' }
#' @describeIn as.std.eyename Return a vector containing all standard right eye identifier strings
#' @noMd
#' @export
od_str <- function() {
  c("OD",
    "RE",
    "Rechts",
    "Right",
    "Dexter",
    "od",
    "re",
    "rechts",
    "right",
    "dexter")
}

#' @examples
#' \dontrun{
#' os_str()
#' }
#' @describeIn as.std.eyename Return a vector containing all standard left eye identifier strings
#' @noMd
#' @export
os_str <- function() {
  c("OS",
    "LE",
    "Links",
    "Left",
    "Sinister",
    "os",
    "le",
    "links",
    "left",
    "sinister")
}

#' @examples
#' \dontrun{
#' ou_str()
#' }
#' @describeIn as.std.eyename Return a vector containing all standard identifier strings identifying recordings acuired from/for both eyes
#' @noMd
#' @export
ou_str <- function() {
  c("OU",
    "BE",
    "EB",
    "Both",
    "Beide",
    "uo",
    "be",
    "eb",
    "both",
    "beide")
}

#' @examples
#' \dontrun{
#' erg_str()
#' }
#' @describeIn as.std.channelname Get standard ERG channel name strings
#' @noMd
#' @keywords internal
erg_str <- function() {
  c("ERG",
    "ERGs",
    "ERG_auto",
    "C-wave",
    "PhNp")
}

#' @examples
#' \dontrun{
#' op_str()
#' }
#'
#' @describeIn as.std.channelname Get standard oscillatory potentials channel name strings
#' @noMd
#' @keywords internal
op_str <- function() {
  c("OP",
    "OPs")
}

#' @examples
#' \dontrun{
#' vep_str()
#' }
#'
#' @describeIn as.std.channelname Get standard VEP channel name strings
#' @noMd
#' @keywords internal
vep_str <- function() {
  c("VEP",
    "VEPs")
}
```
