## Supplemental Material 1 for "The ERGtools2 package: A Toolset for Processing and Analysing Visual Electrophysiology Data": INT_Espion_functions.R

```
get_toc <- function(filename, sep = "\t") {
  toc <- fread(
    filename,
    sep = sep,
    data.table = F,
    skip = 1,
    header = T,
    blank.lines.skip = T,
    select = c("Table", "Left", "Right", "Top", "Bottom"),
    encoding = "UTF-8"
  )

  HeaderTabPos <-
    which(toc$Table == "Header Table") # if vertical table, chop off after end of Header
  if (length(HeaderTabPos) > 1) {
    # "Header Table" found twice, so its a vertical table
    EndOfTOC <-
      min(which(toc$Table == "")[which(toc$Table == "") > HeaderTabPos[1]]) # this is the first blank line after TOC
    toc <- toc[1:EndOfTOC, ]
  }

  toc = toc[!toc$Table == "", ]
  tmp <- toc$Table
  toc$Table <- NULL
  toc <-
    data.frame(apply(toc, 2, function(x)
      as.numeric(as.character(x))))
  rownames(toc) <- tmp

  # check if Header and Stimulus Info present
  if (!sum(rownames(toc) %in% c("Header Table", "Stimulus Table")) == 2) {
    stop("Table of content incomplete.")
  }

  return(toc)
}

#' @importFrom data.table fread
get_content <- function(filename, toc, what, sep = "\t") {
  #Sys.setlocale("LC_ALL", "C")
  recording_info <- fread(
    filename,
    sep = sep,
    select = toc[what, "Left"]:toc[what, "Right"],
    nrows = toc[what, "Bottom"] - toc[what, "Top"] +
      1,
    skip = toc[what, "Top"] - 1,
    data.table = F,
    header = F,
    encoding = "UTF-8"
  )

  if (!(what %in% (rownames(toc)))) {
    stop("'", what, "' not fount in Table of Content.")
  }
  tmp <-  fread(
    filename,
    sep = sep,
    select = toc[what, "Left"]:toc[what, "Right"],
    nrows = 1,
    skip = toc[what, "Top"] -
      3,
    data.table = F,
    header = F,
    encoding = "UTF-8"
  )

  tmp <-
    make.unique(make.names(iconv(tmp, "ASCII//TRANSLIT", sub = '')))

  colnames(recording_info) <- tmp

  return(recording_info)
}

#' @importFrom stringr str_detect
#' @keywords internal
get_stim_info <- function(filename, toc, sep) {
  stim_info <-
    as.data.frame(get_content(filename, toc, "Stimulus Table", sep = sep))
  colnames(stim_info)[str_detect(colnames(stim_info), "cd.")] <-
    "StimulusEnergy"
  stim_info$Background <- NA
  stim_info$Background[grepl("LA", stim_info[, "Description"], fixed = TRUE, useBytes = TRUE)] <-
    "LA"
  stim_info$Background[grepl("DA", stim_info[, "Description"], fixed = TRUE, useBytes = TRUE)] <-
    "DA"
  stim_info$Type <- NA
  stim_info$Type[stim_info$Background == "DA"] <- "Flash"
  stim_info$Type[grepl("Flash",
                       stim_info[, "Description"],
                       fixed = TRUE,
                       useBytes = TRUE)] <- "Flash"
  stim_info$Type[grepl("Single",
                       stim_info[, "Description"],
                       fixed = TRUE,
                       useBytes = TRUE)] <- "Flash"
  stim_info$Type[grepl("Flicker",
                       stim_info[, "Description"],
                       fixed = TRUE,
                       useBytes = TRUE)] <- "Flicker"

  stim_info$Description <- str_replace_all(enc2utf8(stim_info$Description), regex("[^A-Za-z0-9]+"), " ")

  return(stim_info)
}

#' @importFrom units as_units
#' @keywords internal
get_measurements <- function(filename, toc, sep) {
  measurements <-
    get_content(filename, toc, "Marker Table", sep = sep)
  measurements <-
    measurements[, c("Group", "S", "C", "R", "Eye", "Name.1", "uV", "ms")]
  colnames(measurements)[colnames(measurements) == "S"] <- "Step"
  colnames(measurements)[colnames(measurements) == "C"] <-
    "Channel"
  colnames(measurements)[colnames(measurements) == "Name.1"] <-
    "Marker"
  colnames(measurements)[colnames(measurements) == "R"] <-
    "Repeat"

  TimeUnit <-enc2utf8( colnames(measurements)[colnames(measurements) == "ms"])
  colnames(measurements)[colnames(measurements) == "ms"] <- "Time"
  measurements$Time <- as_units(measurements$Time, TimeUnit)

  voltageunit <-
    colnames(measurements)[colnames(measurements) == "uV"]
  colnames(measurements)[colnames(measurements) == "uV"] <-
    "Voltage"
  measurements$Voltage <-
    as_units(measurements$Voltage, voltageunit)
  return(measurements)
}

#' @importFrom data.table fread
#' @importFrom units as_units
#' @keywords internal
get_trace <- function(filename, toc, Data_Header, IDX, what) {
  if (what == "TimeTrace") {
    sel = Data_Header[IDX, "Column"]
  }
  if (what == "ResultTrace") {
    sel = Data_Header[IDX, "Column.1"]
  }
  if (what == "TrialTrace") {
    sel = c((Data_Header[IDX, "Column.1"] +
               1):(Data_Header[IDX, "Column.1"] + Data_Header[IDX, "Trials"]))
  }

  ERRORSTRING<-paste0("While importing ",what," for Step '",Data_Header[IDX, "Step"],"', Channel '",Data_Header[IDX, "Chan"],"', Repeat '",Data_Header[IDX, "Repeat"],"': ")

  tryCatch({

    UNIT <- fread(
      filename,
      select = sel,
      nrows = 1,
      skip = toc["Data Table", "Top"] - 2,
      data.table = F,
      header = F
    )
    UNIT<-unlist(UNIT)
    if (what == "TrialTrace") {
      earlyend <- which(UNIT != UNIT[1])
      if (length(earlyend)!=0){
        earlyend <- min(earlyend)
        warning(
          paste0(ERRORSTRING, " It seems like Trials/Repeats have been rejected in the Espion software and were therefore not included into the export file. It is recommended to export all Trials and reject unwanted Trials inside ERGtools2.")
        )
        UNIT<-UNIT[1:earlyend-1]
        sel<-sel[1:earlyend-1]
      }
    }

    TRACE <-
      fread(
        filename,
        select = sel,
        nrows = toc["Data Table", "Bottom"] -
          toc["Data Table", "Top"],
        skip = toc["Data Table", "Top"] -
          1,
        data.table = F,
        header = F
      )
    if (what %in% c("TimeTrace", "ResultTrace")) {
      TRACE <- TRACE[, 1]
    }

    if (what %in% c("TimeTrace", "ResultTrace")) {
      UNIT <- UNIT[1]
    } else {
      premature.end<-which(UNIT == "Time (ms)")
      if (length(premature.end)!=0){
        TRACE<-TRACE[,1:premature.end-1]
      }
      UNIT <- unique(as.vector(t(UNIT)))
    }

    UNIT <-
      gsub("[\\(\\)]", "", regmatches(UNIT, gregexpr("\\(.*?\\)", UNIT))[[1]])

    if (length(UNIT) > 1) {
      stop(ERRORSTRING, "Distict units detected.")
    }

    if (what == "TrialTrace") {
      TRACE <-
        as.matrix(TRACE[apply(TRACE, 1, function(x) {
          all(!is.na(x))
        }), ])
    } else {
      TRACE <- TRACE[!is.na(TRACE)]
    }

    TRACE <- as_units(TRACE, UNIT)
  }, error = function(e){
    stop(ERRORSTRING, e)
  })

  return(TRACE)
}

#' @keywords internal
inferre.channel.names.from.markers<-function(markers){

  checkfx <- function(l, pos,markers) {
    any(unlist(lapply(l[names(l) %in% pos], function(x) {
      all(x %in% markers)
    }))) &&
      !any(unlist(lapply(l[!(names(l) %in% pos)], function(x) {
        all(x %in% markers)
      })))

  }

  def_markers<-list(
    ERG_flash=c("A","B"),
    ERG_flicker=c("P1","N1"),
    ERG_OP=c("OP1","OP2","OP3"),
    VEP_early=c("P1","N1","P2"),
    VEP_late=c("P100","N75","N135","P200","P300")
  )
  markers<-toupper(markers)

  # ERG
  if (checkfx(def_markers, c("ERG_flash", "ERG_flicker"), markers)) {
    return("ERG")
  }
  # OP
  if (checkfx(def_markers, c("ERG_OP"), markers)) {
    return("OP")
  }
  # VEP
  if (checkfx(def_markers, c("VEP_early", "VEP_late"), markers)) {
    return("VEP")
  }
  return("Unknown")
}
```
