## Supplemental Material 1 for "The ERGtools2 package: A Toolset for Processing and Analysing Visual Electrophysiology Data": INT_ExtraMetaColumns.R

```
#' @keywords internal
#' @noMd
setGeneric(
  name = "ExtraMetaColumns",
  def = function(X)
  {
    standardGeneric("ExtraMetaColumns")
  }
)
#' @keywords internal
#' @noMd
setMethod("ExtraMetaColumns",
          "ERGExam",
          function(X) {
            cols<-colnames(Metadata(X))
            return(cols[!(cols %in% c("Step", "Channel", "Repeat", "Eye"))])
          })
```
