## Supplemental Material 1 for "The ERGtools2 package: A Toolset for Processing and Analysing Visual Electrophysiology Data": INT_str_to_unit.R

```
#' @importFrom units as_units
#' @importFrom stringr str_extract regex str_length
#' @keywords internal
str_to_unit <- function(val) {
  if (str_length(val)>0){
    return(as_units(as.numeric(str_extract(val, regex("[0-9.]+"))), str_extract(val, regex("[^0-9.]+"))))
  } else {
    return(NA)
  }
}
```
