## Supplemental Material 1 for "The ERGtools2 package: A Toolset for Processing and Analysing Visual Electrophysiology Data": interactiveMeasurements.R

```
#' Interactively perform measurements on an ERGExam object
#'
#' This method allows interactive marker placement on \linkS4class{ERGExam} objects using  \link[shiny:shiny]{shiny:shiny-package}
#'
#' @inheritParams  Subset-method
#' @return An object of class 'ERGExam' with the new measurements added.
#' @export
#' @importFrom plotly ggplotly event_data plotlyOutput renderPlotly event_register
#' @importFrom shiny fluidPage titlePanel fluidRow column verticalLayout tags validate HTML h4 actionButton selectInput textInput
#' @importFrom DT dataTableOutput renderDataTable datatable
#' @importFrom shiny modalDialog modalButton showModal removeModal observeEvent reactive stopApp runApp onSessionEnded shinyApp
#' @importFrom shinyjs useShinyjs extendShinyjs
#' @importFrom stringr str_split
#' @importFrom ggplot2 aes geom_line geom_text facet_grid labs theme_minimal element_text geom_vline label_wrap_gen
#' @importFrom units set_units deparse_unit
#' @importFrom utils txtProgressBar setTxtProgressBar
#' @importFrom tidyr %>%
#' @examples
#' data(ERG)
#' ERG<-SetStandardFunctions(ERG)
#' \dontrun{
#' ERG<-interactiveMeasurements(ERG, Channel="OP",Eye="RE")
#' }
#' Measurements(ERG)
#' \dontrun{
#' ERG<-interactiveMeasurements(ERG, Channel="ERG")
#' }
#' Measurements(ERG)
#'
setGeneric(
  name = "interactiveMeasurements",
  def = function(X,
                 Channel = "ERG",
                 Eye = Eyes(X)) {
    standardGeneric("interactiveMeasurements")
  }
)

#' @noMd
setMethod("interactiveMeasurements",
          "ERGExam", {
            function(X,
                     Channel = "ERG",
                     Eye = Eyes(X)) {
              # internal functions
              interact_plot <- function(erg.obj) {
                trace <- as.data.frame(erg.obj)
                trace$Value <- set_units(trace$Value, "uV")

                markers <-
                  Measurements(erg.obj, measure.absolute = T)
                colnames(markers)[colnames(markers) == "Voltage"] <-
                  "Value"
                trace <-
                  merge(trace, Stimulus(erg.obj)[c("Step", "Description")], by = "Step")

                g <- ggplot(trace,
                            aes(
                              x = Time,
                              y = Value,
                              color = factor(Repeat),
                              customdata = key
                            )) +
                  geom_line()
                if (nrow(markers) > 0) {
                  g <- g +
                    geom_vline(data = markers,
                               aes(xintercept = Time),
                               alpha = 0.1) +
                    geom_text(data = markers,
                              aes(
                                x = Time,
                                y = Value,
                                label = Name
                              ),
                              color = "black")
                }
                g <- g +
                  facet_grid(Description ~ Eye,
                             scales = "free_y",
                             labeller = label_wrap_gen(25)) +
                  labs(title = "Trace Plots", x = "Time", y = "Amplitude") +
                  theme_minimal() +
                  theme(legend.position = "none",
                        strip.text.y = element_text(angle = -90)) # Multi-ine labeller https://stackoverflow.com/questions/12673392/how-to-fit-strip-text-x-if-the-heading-string-is-too-long
                return(g)
              }

              shiny.style <- HTML(
                "
                                    .highlight-row {
                                      border: 2px solid #3498db;
                                      padding: 10px;
                                      border-radius: 5px;
                                      margin: 10px 0;
                                    }
                                 "
              )

              # Validity checks
              if (!all(Eye %in% Eyes(X))) {
                stop("Invalid value(s) given for 'Eye'.")
              }

              if (!all(Channel %in% Channels(X))) {
                stop("Invalid value given for 'Channel'.")
              }

              if (length(Channel) != 1) {
                stop("Exactly one valid Channel must be named")
              }

              stopifnot(CheckAvgFxSet(X))

              #Get Relevant subset of the ERGExam and keep avg and filter fx.

              curr <- Subset(X,
                             where = list(Channel = Channel,
                                          Eye =  Eye),
                             Raw = F)


              markers <- Markers(curr)
              if (nrow(markers) == 0) {
                curr <- AutoPlaceMarkers(curr)
                markers <- Markers(curr)
              }
              if (nrow(markers) > 0) {
                for (i in nrow(Markers(curr)):1) {
                  markers$ChannelBinding[i]
                  if (!(markers$ChannelBinding[i] %in% Channel)) {
                    curr <- DropMarker(curr,
                                       markers$Name[i],
                                       markers$ChannelBinding[i],
                                       drop.dependent = T)
                  }
                }
              }


              # prepare ERGExam Object
              curr@Stimulus$Description <-
                iconv(curr@Stimulus$Description, "UTF-8", "UTF-8", sub = '')
              key <-
                as.factor(paste(
                  Metadata(curr)$Step,
                  Metadata(curr)$Eye,
                  Metadata(curr)$Repeat,
                  sep = "_"
                ))
              curr <- AddMetadata(curr, "key", key)


              jscode <- "shinyjs.closeWindow = function() { window.close(); }"
              # Define the UI for the application
              ui <- fluidPage(
                useShinyjs(),
                extendShinyjs(text = jscode, functions = c("closeWindow")),
                tags$head(tags$style(shiny.style)),
                titlePanel(
                  paste0(
                    "Set Markers - Subject: ",
                    Subject(X),
                    ", DOE: ",
                    ExamDate(X),
                    ", Channel = ",
                    Channel
                  )
                ),
                sidebarLayout(
                  sidebarPanel(
                    verticalLayout(
                      fluidRow(class = "highlight-row",
                               actionButton("finishBtn", "Finish")),
                      fluidRow(
                        class = "highlight-row",
                        h4("Select active marker"),
                        dataTableOutput("markerTable")
                      ),
                      fluidRow(
                        class = "highlight-row",
                        h4("Add new marker"),
                        textInput("INP_marker_new", "Name"),
                        selectInput(
                          "INP_marker_new_rel",
                          "Relative to",
                          choices = c("", markers$Name),
                          selected = 1
                        ),
                        actionButton("INP_marker_new_add", "<-- Add new marker")
                      ),

                      fluidRow(
                        class = "highlight-row",
                        actionButton("INP_marker_delete", "Delete selected marker")
                      )
                    ),
                    width = 3
                  ),
                  mainPanel(
                    h4("Traces"),
                    plotlyOutput(
                      "trace_plot",
                      height =  paste0(200 + (100 * length(Steps(
                        curr
                      ))), "px"),
                      width = paste0(120 + (300 * length(Eyes(
                        curr
                      ))), "px")
                    ),
                    hr(),
                    h4("Measurements"),
                    dataTableOutput("measurement_output"),
                    actionButton("INP_measurement_delete", "Delete selected measurement"),
                    width = 9
                  ),
                  position = "right"
                )
              )

              # Define the server for the application
              server <- function(input, output, session) {
                out <- curr
                CURR <- reactiveValues(data = curr)

                # Render the interactive plot
                output$trace_plot <- renderPlotly({
                  ggplotly(interact_plot(CURR$data),
                           width = (80 + (300 * length(Eyes(
                             curr
                           ))))) %>%
                    event_register("plotly_click")
                })

                #Display the available markers
                output$markerTable <-
                  renderDataTable({
                    Markers(CURR$data)[, c("Name", "Relative")]
                  }, selection = "single",
                  options = list(dom = 't'))

                # Observe marker Select
                observeEvent(input$markerTable_rows_selected, {
                  marker.sel.idx <<- isolate(input$markerTable_rows_selected)
                  curr_marker <<-
                    Markers(CURR$data)$Name[marker.sel.idx]
                  curr_marker_ref <<-
                    Markers(CURR$data)$Relative[marker.sel.idx]
                })

                # Observe measurement Select
                observeEvent(input$measurement_output_rows_selected, {
                  measurement.sel.idx <<-
                    isolate(input$measurement_output_rows_selected)
                })

                # Observe new marker position set
                observeEvent(event_data("plotly_click"),
                             {
                               click.coord <- event_data("plotly_click")
                               if (!is.null(click.coord)) {
                                 if (curr_marker != "") {
                                   # get current eye and step for curve_number
                                   click.traceinfo <-
                                     unlist(stringr::str_split(click.coord$customdata, "_"))
                                   if (!is.null(click.traceinfo)) {
                                     click.Step <-
                                       as.numeric(click.traceinfo[1])
                                     click.Eye <- click.traceinfo[2]
                                     click.Repeat <-
                                       as.numeric(click.traceinfo[3])
                                     click.idx <-
                                       IndexOf(
                                         CURR$data,
                                         Step = click.Step,
                                         Eye = click.Eye,
                                         Channel = Channel,
                                         Repeat = click.Repeat
                                       )

                                     existing <- Markers(CURR$data)
                                     canupdate = F
                                     print(existing)
                                     if (!is.na(existing$Relative[existing$Name == curr_marker])) {
                                       print("REL")
                                       if (nrow(
                                         Measurements(
                                           X = CURR$data,
                                           Marker = existing$Relative[existing$Name ==
                                                                      curr_marker],
                                           where = click.idx,
                                           TimesOnly = T
                                         )
                                       ) != 0) {
                                         print("Parent Present")
                                         canupdate = T
                                       }
                                     } else {
                                       canupdate = T
                                     }

                                     if (canupdate) {
                                       Measurements(
                                         X = CURR$data,
                                         Marker = curr_marker,
                                         where = click.idx,
                                         create.marker.if.missing = F,
                                         Relative = NULL,
                                         ChannelBinding = NULL
                                       ) <-
                                         as_units(click.coord$x, deparse_unit(TimeTrace(curr@Data[[click.idx]])))
                                     } else {
                                       showModal(
                                         modalDialog(
                                           title = "Placement of marker failed.",
                                           paste0(
                                             "The marker ",
                                             curr_marker,
                                             " could not be placed on this trace. This might be becuase the marker is relative to another, which is still missing in the trace."
                                           ),
                                           easyClose = TRUE,
                                           footer = NULL
                                         )
                                       )
                                     }

                                     # update the plot
                                     output$trace_plot <-
                                       renderPlotly({
                                         ggplotly(interact_plot(CURR$data),
                                                  width = (80 + (300 * length(
                                                    Eyes(curr)
                                                  ))))
                                       })
                                     # reselect marker table
                                     output$markerTable <-
                                       renderDataTable({
                                         Markers(CURR$data)[, c("Name", "Relative")]
                                       }, selection = list(mode = "single",
                                                           selected = marker.sel.idx),
                                       options = list(dom = 't'))

                                     out <<-
                                       reactiveValuesToList(CURR)$data
                                   }
                                 }
                               }
                             })

                # Observe new Markers
                observeEvent(input$INP_marker_new_add, {
                  new_marker <- isolate(input$INP_marker_new)
                  new_marker_rel <-
                    isolate(input$INP_marker_new_rel)
                  if (!is.null(new_marker) &&
                      new_marker != "" &&
                      !(new_marker %in% Markers(CURR$data)$Name)) {
                    CURR$data <-
                      AddMarker(
                        CURR$data,
                        Marker = new_marker,
                        Relative = which(Markers(CURR$data)$Name == new_marker_rel),
                        ChannelBinding = Channel
                      )
                    updateSelectInput(session,
                                      "INP_marker_new_rel",
                                      choices = c("", Markers(CURR$data)$Name))

                    out <<- reactiveValuesToList(CURR)$data
                  } else {
                    showModal(
                      modalDialog(
                        title = "Adding of new marker failed.",
                        paste0("The new marker could not be added ",
                               if (new_marker == "") {
                                 ": New marker name must not be an empty string,"
                               },
                               if (new_marker %in% Markers(CURR$data)$Name) {
                                 ": Marker name already in list."
                               }),
                        easyClose = TRUE,
                        footer = NULL
                      )
                    )
                  }
                })

                # Observe deleteMarker Button
                observeEvent(input$INP_marker_delete, {
                  showModal(modalDialog(
                    title = "Delete Marker",
                    paste0("Delete selected marker(", curr_marker, ") and all markers relative to it?"),
                    footer = tagList(
                      actionButton("confirmDelete", "Delete"),
                      modalButton("Cancel")
                    )
                  ))
                })
                observeEvent(input$confirmDelete, {
                  CURR$data <-
                    DropMarker(
                      CURR$data,
                      curr_marker,
                      ChannelBinding = Channel,
                      drop.dependent = T
                    )
                  out <<- reactiveValuesToList(CURR)$data
                  removeModal()
                })

                # Observe delete Measurement Button
                observeEvent(input$INP_measurement_delete, {
                  if (!is.null(measurement.sel.idx)) {
                    print(measurement.sel.idx)
                    if (all(measurement.sel.idx %in% 1:nrow(CURR$data@Measurements@Measurements))) {
                      warning("Deleting Measurments interactively is experimental.")
                      # needs to implement checks to ensure data dropped have no depending or call Measurements()<-NULL

                      tmp<-mes[measurement.sel.idx,]
                      tryCatch(
                        Measurements(
                          CURR$data,
                          where = list(
                            Step = tmp$Step,
                            Eye = tmp$Eye,
                            Repeat = tmp$Repeat
                          ),
                          Marker = tmp$Name,
                          ChannelBinding = Channel
                        ) <- NULL,
                        error = function(e) {
                          showModal(
                            modalDialog(
                              title = "Deleting measurement failed.",
                              paste0("Deleting measurement failed with message: ",e),
                              easyClose = TRUE,
                              footer = NULL
                            )
                          )
                        }
                      )
                      out <<- reactiveValuesToList(CURR)$data
                    } else {
                      showModal(
                        modalDialog(
                          title = "Deleting Measurement failed.",
                          "No Measurement was selected.",
                          easyClose = TRUE,
                          footer = NULL
                        )
                      )
                    }
                  } else {
                    showModal(
                      modalDialog(
                        title = "Deleting Measurement failed.",
                        "Selection does not exist.",
                        easyClose = TRUE,
                        footer = NULL
                      )
                    )
                  }
                })

                # Observe the finish button
                observeEvent(input$finishBtn, {
                  showModal(modalDialog(
                    title = "Close & Save",
                    "You will now close the dialog and return an updated ERGExam object",
                    footer = tagList(
                      actionButton("confirmSaveClose", "Close & Save"),
                      actionButton("confirmDiscard", "Close & DISCARD changes"),
                    )
                  ))
                })
                observeEvent(input$confirmSaveClose, {
                  js$closeWindow()
                  stopApp(out)
                })

                observeEvent(input$confirmDiscard, {
                  js$closeWindow()
                  stopApp(curr)
                })
                onSessionEnded(function() {
                  js$closeWindow()
                  stopApp(out)
                })

                #Display the measurement results
                output$measurement_output <-
                  renderDataTable({
                    mes <-Measurements(CURR$data,TimesOnly=T,quiet = T)[,c("Step","Eye","Repeat","Name","Time")]
                    mes<-mes[with(mes, order(Step, Eye, Repeat)), ]
                    mes<<-mes
                    datatable(mes, selection = "single")
                  }, width = "100%", selection = "single",
                  options = list(dom = 't'))
              }


              # Make the application
              SetMarkersApp <- shinyApp(ui = ui, server = server)

              # run
              message("Waiting for Marker placement app to finish....")
              out <- runApp(SetMarkersApp, launch.browser = T)
              message("Finished. Please wait.")

              # merge measurements with unchanged part of object
              old.measurements <-
                Measurements(curr, TimesOnly = T, quiet = T)
              new.measurements <-
                Measurements(out, TimesOnly = T, quiet = T)
              removed.measurements <-
                old.measurements[!(do.call(paste, old.measurements[, c("Step",
                                                                       "Eye",
                                                                       "Recording",
                                                                       "Repeat",
                                                                       "Name",
                                                                       "Channel",
                                                                       "Relative")]) %in% do.call(paste, new.measurements[, c("Step",
                                                                                                                              "Eye",
                                                                                                                              "Recording",
                                                                                                                              "Repeat",
                                                                                                                              "Name",
                                                                                                                              "Channel",
                                                                                                                              "Relative")])), c("Step",
                                                                                                                                                "Eye",
                                                                                                                                                "Recording",
                                                                                                                                                "Repeat",
                                                                                                                                                "Name",
                                                                                                                                                "Channel")]
              ## update and add
              new.measurements <-
                merge(
                  Measurements(out@Measurements),
                  Metadata(out),
                  by.x = "Recording",
                  by.y = 0
                )

              if (nrow(new.measurements) > 0) {
                message("Updating measurements...")
                pb = txtProgressBar(
                  min = 0,
                  max = nrow(new.measurements),
                  initial = 0
                )
                for (m in 1:nrow(new.measurements)) {
                  curr.m <- new.measurements[m, ]
                  Measurements(
                    X,
                    Marker = curr.m$Name,
                    where = Where(
                      X,
                      where = list(
                        Step = curr.m$Step,
                        Eye = curr.m$Eye,
                        Channel = curr.m$Channel,
                        Repeat = curr.m$Repeat
                      )
                    ),
                    create.marker.if.missing = T,
                    Relative = curr.m$Relative,
                    ChannelBinding = curr.m$Channel
                  ) <- curr.m$Time
                  setTxtProgressBar(pb, m)
                }
                close(pb)
              }

              ## drop removed
              if (nrow(removed.measurements) > 0) {
                message("Deleting removed measurements...")
                pb = txtProgressBar(
                  min = 0,
                  max = nrow(new.measurements),
                  initial = 0
                )
                for (m in 1:nrow(removed.measurements)) {
                  curr.m <- removed.measurements[m, ]
                  Measurements(
                    X,
                    Marker = curr.m$Name,
                    where = Where(
                      X,
                      where = list(
                        Step = curr.m$Step,
                        Eye = curr.m$Eye,
                        Channel = curr.m$Channel,
                        Repeat = curr.m$Repeat
                      )
                    ),
                    create.marker.if.missing = T,
                    Relative = curr.m$Relative,
                    ChannelBinding = curr.m$ChannelBinding
                  ) <- NULL
                  setTxtProgressBar(pb, m)
                }
                close(pb)
              }
              if (validObject(X)) {
                return(X)
              }
            }
          })
```
