## Supplemental Material 1 for "The ERGtools2 package: A Toolset for Processing and Analysing Visual Electrophysiology Data": LoadSave.R

```
#' Load/Save ERGExam objects to or from  HDF5 files
#'
#' These functions load and save \linkS4class{ERGExam} objects from or into HDF5 files.
#' @param X An \linkS4class{ERGExam} object.
#' @param filename Path the data is read from or written to.
#' @param overwrite Should existing files be overwritten?
#' @return
#' * Save: Does not return any values.
#' * Load: An \linkS4class{ERGExam} object.
#' @name LoadSave
#' @rdname LoadSave-methods
#' @aliases Load Save
#' @docType methods
NULL

#' @export
#' @importFrom stringr str_detect
#' @importFrom units deparse_unit
#' @importFrom hdf5r H5File
#' @importFrom EPhysData Save
#' @examples
#' data(ERG)
#' ERG<-SetStandardFunctions(ERG)
#' fn <- tempfile()
#' Save(ERG, fn, overwrite=T)
#' @rdname LoadSave-methods
setMethod("Save",
          "ERGExam",
          function(X,
                   filename,
                   overwrite) {

            Save(as(X,"EPhysSet"), filename, overwrite = overwrite) # Verification of file name etc in here, also starts new file

            if(exists("con")){
              if(con$is_valid){
                con$close_all()
              }
            }
            tryCatch({
              con <- H5File$new(filename, mode = "a")
            }, error= function(e){
              if (str_detect(e$message,"unable to create file")){
                out<-e
                stop("File '", filename,"' could not be created. Check path.")
              } else {
                stop(e)
              }
            })

            X@Stimulus$Description<-iconv(X@Stimulus$Description, "UTF-8", "UTF-8", sub = '')
            con$create_dataset("Stimulus",X@Stimulus,gzip_level = 3)
            con[["Averaged"]] <- X@Averaged
            M_SLOT <- con$create_group("Measurements")
            M_SLOT$create_dataset("Marker",X@Measurements@Marker,gzip_level = 9)
            M_SLOT$create_dataset("Measurements",X@Measurements@Measurements,gzip_level = 9)
            M_SLOT[["TimeUnit"]] <- deparse_unit(X@Measurements@Measurements$Time)
            con[["ExamInfo_Content"]] <- names(X@ExamInfo)
            E_SLOT <- con$create_group("ExamInfo")
            for (i in 1:length(X@ExamInfo)){
              E_SLOT[[names(X@ExamInfo)[i]]] <- X@ExamInfo[[i]]
            }

            con[["SubjectInfo_Content"]] <- names(X@SubjectInfo)
            S_SLOT <- con$create_group("SubjectInfo")
            for (i in 1:length(X@SubjectInfo)){
              S_SLOT[[names(X@SubjectInfo)[i]]] <- X@SubjectInfo[[i]]
            }
            con[["Imported"]] <- as.character(X@Imported)
            con$link_delete("Type")
            con[["Type"]] <- "ERGExam"
            con[["ERGExamVersion"]] <- as.character(packageVersion('ERGtools2'))
            con$close_all()
          })


#' @importFrom hdf5r H5File
#' @importFrom utils packageVersion compareVersion getFromNamespace
#' @importFrom EPhysMethods filter.detrend filter.bandpass filter.lin.detrend autoreject.by.absolute.threshold autoreject.by.distance autoreject.by.signalfree
#' @describeIn LoadSave-methods Load \linkS4class{EPhysData:EPhysData} or \linkS4class{EPhysData:EPhysSet} objects from an HDF5 file
#' @examples
#' data(ERG)
#' ERG<-SetStandardFunctions(ERG)
#' fn <- tempfile()
#' Save(ERG, fn, overwrite=T)
#' require(EPhysMethods) # EPhysMethods is required as SetStandardFunctions has written functions from it into the file
#' Load.ERGExam(fn)
#' @export
Load.ERGExam <- function(filename) {
  Load.EPhysData_imp<-getFromNamespace("Load.EPhysData","EPhysData")
  if (file.exists(filename)) {
    con <- H5File$new(filename, mode = "r")
  } else {
    stop("File '", filename, "' does not exist. ")
  }

  if (compareVersion(con$open("EPhysDataVersion")$read(),
                     as.character(packageVersion('EPhysData')))>0) {
    warning("This file was created using a newer version of EPhysData (",con$open("EPhysDataVersion")$read(),")")
  }

  if (con$open("Type")$read() == "ERGExam") {
    if (compareVersion(con$open("ERGExamVersion")$read(),
                       as.character(packageVersion('ERGtools2')))>0) {
      warning("This file was created using a newer version of ERGtools2 (",con$open("ERGExamVersion")$read(),")")
    }
  } else {
    stop("This HDF5 file has no ERGExam stored.")
  }

  Data.con <- con$open("Data")
  DATA <- list()
  pb = txtProgressBar(min = 0,
                      max = length(names(Data.con)),
                      initial = 0)
  j = 0
  for (i in names(Data.con)) {
    Data.curr <- Data.con$open(i)
    DATA[[as.integer(i)]] <- Load.EPhysData_imp(con = Data.curr)
    setTxtProgressBar(pb, j)
    j = j + 1
  }
  close(pb)
  Metadata = con$open("Metadata")$read()
  Stimulus = con$open("Stimulus")$read()
  Averaged = con$open("Averaged")$read()


  M_SLOT <- con$open("Measurements")
  M <- new("ERGMeasurements")
  M@Marker<-M_SLOT$open("Marker")$read()
  M@Measurements<-M_SLOT$open("Measurements")$read()
  M@Measurements$Time<-as_units(M@Measurements$Time, M_SLOT$open("TimeUnit")$read())
  stopifnot(validObject(M))

  ExamInfo_Content = con$open("ExamInfo_Content")$read()
  ExamInfo <- list()
  E_SLOT <- con$open("ExamInfo")
  for (i in ExamInfo_Content) {
    ExamInfo[[i]] <- E_SLOT$open(i)$read()
  }
  ExamInfo$ExamDate <-
    as.POSIXct(ExamInfo$ExamDate, origin = "1970-01-01")
  SubjectInfo_Content = con$open("SubjectInfo_Content")$read()
  SubjectInfo <- list()
  S_SLOT <- con$open("SubjectInfo")
  for (i in SubjectInfo_Content) {
    SubjectInfo[[i]] <- S_SLOT$open(i)$read()
  }
  SubjectInfo$DOB <- as.Date(SubjectInfo$DOB, origin = "1970-01-01")
  Imported = as.POSIXct(con$open("Imported")$read(),origin="1970-01-01")

  out<-newERGExam(
    Data = DATA,
    Metadata = Metadata,
    Stimulus = Stimulus,
    Measurements = M,
    Averaged = Averaged,
    ExamInfo = ExamInfo,
    SubjectInfo = SubjectInfo
  )

  out@Imported <- Imported

  con$close_all()

  if(validObject(out)){
    return(out)
  }

}
```
