## Supplemental Material 1 for "The ERGtools2 package: A Toolset for Processing and Analysing Visual Electrophysiology Data": MarkerMethods.R

```
#' Access and Modify the Markers Stored in an ERGExam or ERGMeasurements Object
#'
#' These methods enable accession and  modification of the Markers stored in an  \linkS4class{ERGExam} object or the underlying \linkS4class{ERGMeasurements} object directly.
#'
#' @param X An \linkS4class{ERGExam} or \linkS4class{ERGMeasurements} object.
#' @param Marker Name of the marker.
#' @param ChannelBinding For AddMarker and DropMarker only. Channel to which the marker belongs. Usually defined by the Parent \linkS4class{ERGExam} object. Set to NA if not required.
#' @param Relative For AddMarker and Measurements<- only. Index of the marker the new marker is relative to.
#' @param drop.dependent For DropMarker and DropMeasurement only. Logical. If TRUE, dependent markers will also be dropped. If FALSE, fails if there are markes dependent on that removed.
#' @param update.empty.relative Logical. If an empty relative value should be overwritten by an otherways matching marker.
#' @return An updated version of the object, except for Markers, which returns a data.frame representing the Markers stored in the the \linkS4class{ERGExam} or the \linkS4class{ERGMeasurements} object.
#' @seealso \link[EPhysData:EPhysSet-class]{EPhysData::EPhysSet-class} \link{Measurements-Methods} \link{Get}
#' @examples
#' # Add a new Marker to an ERGMeasurements object#'
#' data(Measurements.data)
#' new_erg_measurements <- AddMarker(Measurements.data, Marker = "Marker1", Relative = NA, ChannelBinding = "ChannelA")
#' new_erg_measurements
#' Markers(new_erg_measurements)
#'
#' # Add a new Marker to an ERGMeasurements object
#' data(ERG)
#' new_erg <- AddMarker(ERG, Marker = "Marker1", Relative = NA, ChannelBinding = "ERG")
#' # but this would fail:
#' # new_erg <- DropMarker(new_erg, Marker = "Marker1")
#' Markers(new_erg)
#'
#' # Drop a Marker from an ERGMeasurements object
#' data(Measurements.data)
#' Measurements.data
#' Markers(Measurements.data)
#'
#' new_erg_measurements <- DropMarker(Measurements.data, Marker = "N1", ChannelBinding = "VEP", drop.dependent = TRUE)
#' new_erg_measurements
#' Markers(new_erg_measurements)
#'
#' data(Measurements.data)
#' RenameMarker(Measurements.data,"a","ERG","X")
#'
#' @name Marker-Methods
NULL

#' Add a new Marker
#' @describeIn Marker-Methods Add a new Marker

#' @exportMethod AddMarker
setGeneric(
  name = "AddMarker",
  def = function(X,
                 Marker,
                 Relative = NA,
                 ChannelBinding,
                 update.empty.relative=F) {
    standardGeneric("AddMarker")
  }
)

#' @noMd
setMethod("AddMarker",
          "ERGMeasurements",
          function(X, Marker, Relative, ChannelBinding, update.empty.relative=F) {

            if(is.null(Relative)){
              stop("'Relative' must be set, either to a valid marker name or to NA, if the new marker is not a relative marker." )
            }

            if(length(Relative)==0){
              Relative<-NA
            }

            if(is.null(ChannelBinding)){
              stop("'ChannelBinding' should be set to any valid channel of the parent object. However, can be set to NA, if not required." )
            }

            if(length(Relative)>1){
              Relative<-Relative[!is.na(Relative)]
              if(length(Relative)>1){
                Relative<-Relative[1]
                warning("Multiple non-na values for 'Relative' provided. Taking first.")
              } else {
                warning("Multiple values for 'Relative' provided. Removing NA.")
              }
            }

            if (!is.na(Relative)) {
              if (!is.numeric(Relative) &
                  !is.integer(Relative) & !is.character(Relative)) {
                stop("'Relative' must point to an existing marker (by index or name) in the Marker slot.")
              }
              if (is.character(Relative)) {
                if (is.na(ChannelBinding)) {
                  if (length(MarkerNames(X)[MarkerNames(X) == Relative]) != 1) {
                    stop(
                      "'Relative' is a character string, but does not unambigously point to a parent marker. Use marker indices instead or specify 'ChannelBinding',"
                    )
                  } else {
                    Relative <- which(MarkerNames(X) == Relative)
                  }
                } else {
                  if (length(MarkerNames(X)[MarkerNames(X) == Relative &
                                            Markers(X)$ChannelBinding == ChannelBinding]) == 0) {
                    stop(
                      "'Relative' is a character string, but does not point to a parent marker present in the channel defined by 'ChannelBinding',"
                    )
                  }
                  if (length(MarkerNames(X)[MarkerNames(X) == Relative &
                                            Markers(X)$ChannelBinding == ChannelBinding]) != 1) {
                    stop(
                      "'Relative' is a character string, but does not unambigously point to a parent marker present in the channel defined by 'ChannelBinding',"
                    )
                  } else {
                    Relative <-
                      which(MarkerNames(X) == Relative & Markers(X)$ChannelBinding == ChannelBinding)
                  }
                }
              }
              if (!(Relative %in% 1:length(MarkerNames(X)))) {
                stop("'Relative' must point to an existing marker in the Marker slot")
              }
            }

            # Check if the new marker is valid
            existing_markers <-
              unique(X@Marker$Name[X@Marker$ChannelBinding == ChannelBinding])

            if(length(existing_markers)>0){

              if (Marker %in% existing_markers) {
                if (X@Marker$Relative[X@Marker$ChannelBinding == ChannelBinding & X@Marker$Name == Marker] %in% Relative) {
                  message("This marker is already present in the object. doing nothing.")
                  return(X)
                } else {
                  if (update.empty.relative) {
                    message("An empty relative slot is being overwritten.")
                    X@Marker$Relative[X@Marker$ChannelBinding == ChannelBinding & X@Marker$Name == Marker] <-
                      Relative
                    if (validObject(X)) {
                      return(X)
                    }
                  } else {
                    stop(
                      paste0(
                        "A Marker with the same name ('",
                        Marker,
                        "') but incompatible parent markers (Existing: '",
                        X@Marker$Relative[X@Marker$ChannelBinding == ChannelBinding & X@Marker$Name == Marker] ,
                        "', New: '",
                        Relative,
                        "' ) already exists in the Marker slot for the respective Channel ('",
                        ChannelBinding,
                        "')."
                      )
                    )
                  }
                }
              }
            }
            new_row <-
              data.frame(Name = Marker,
                         Relative = Relative,
                         ChannelBinding = ChannelBinding)


            # Add the new row to the Marker slot
            X@Marker <- rbind(X@Marker, new_row)

            # Return the updated object
            if(validObject(X)){
              return(X)
            }
          })
#' @noMd
setMethod("AddMarker", "ERGExam",
          function(X, Marker, Relative, ChannelBinding) {
            if (!(ChannelBinding %in% Channels(X))){
              stop("'ChannelBinding' must describe a valid channel contained in X.")
            }
            X@Measurements <- AddMarker(X@Measurements, Marker, Relative, ChannelBinding)
            validObject(X)
            return(X)
          })

#' Drop marker by name and channel
#'
#' @describeIn Marker-Methods Drop marker by name and channel
#' @exportMethod DropMarker
setGeneric(
  name = "DropMarker",
  def = function(X, Marker, ChannelBinding, drop.dependent = FALSE) {
    standardGeneric("DropMarker")
  }
)

#' @noMd
setMethod("DropMarker",
          "ERGMeasurements",
          function(X, Marker, ChannelBinding = NULL, drop.dependent = FALSE) {
            if (length(Marker) != 1 || !is.character(Marker)) {
              stop("Multiple marker names provided in 'Marker' argument.")
            }

            # Check if the marker exists in the Marker slot
            if( is.null(ChannelBinding)){
              marker_index <-
                which(X@Marker$Name == Marker)
            } else {
              marker_index <-
                which(X@Marker$Name == Marker & X@Marker$ChannelBinding == ChannelBinding)
            }

            if (length(marker_index) == 0) {
              warning("Marker not found in the Marker slot")
              return(X)
            }
            if (length(marker_index) > 1) {
              stop("Multiple markers found for the Name given by 'Marker'.")
            }

            # Find dependent markers recursively
            findDependentMarkers <-
              function(marker_index, dependent) {
                dependent <-
                  which(X@Marker$Relative %in% marker_index  & X@Marker$ChannelBinding %in% X@Marker$ChannelBinding[marker_index])
                if (length(dependent) == 0) {
                  return(marker_index)
                } else {
                  c(marker_index,
                    findDependentMarkers(dependent, dependent))
                }
              }

            dependent <- findDependentMarkers(marker_index, NULL)
            if (!drop.dependent
                && length(dependent) > 1) { # because dependent includes the index marker
              stop(
                "Other markers are relative to the marker that you are trying to delete. Consider using drop.dependent = TRUE."
              )
            }

            # Create a vector to store indices to remove
            indices_to_remove <-
              sort(unique(c(marker_index, dependent)), decreasing = T)
            X@Marker <- X@Marker[-indices_to_remove,]
            rownames(X@Marker)<-NULL

            # Remove the corresponding rows from the Measurements slot
            measurements_to_remove <-
              which(X@Measurements$Marker %in% indices_to_remove)
            if(length(measurements_to_remove)>0){
              X@Measurements <- X@Measurements[-measurements_to_remove, ]
            }
            rownames(X@Measurements)<-NULL

            # Move the pointers in relative accordingly
            for (i in indices_to_remove) {
              X@Marker$Relative[which(X@Marker$Relative > i)] <-
                X@Marker$Relative[which(X@Marker$Relative > i)] - 1
              X@Measurements$Marker[which(X@Measurements$Marker > i)] <-
                X@Measurements$Marker[which(X@Measurements$Marker > i)] - 1
            }

            if(!validObject(X)){
              stop("Dropping markers failed for unknown reason.")
            }
            # Return the updated object
            return(X)
          })

#' @noMd
setMethod("DropMarker",
          "ERGExam",
          function(X, Marker, ChannelBinding = Channels(X), drop.dependent = FALSE) {
            X@Measurements <-
              DropMarker(X@Measurements , Marker, ChannelBinding, drop.dependent)
            if(validObject(X)){
              return(X)
            } else {
              stop("Object validation failed after dropping Markers")
            }
          })

#' @describeIn Marker-Methods Renames a marker within an ERGMeasurements object according to the specified channel binding.
#' @param New.Name The new name to assign to the marker.
#' @examples
#' data(Measurements.data)  # Assume Measurements.data is a pre-loaded ERGMeasurements object
#' updated_measurements <- RenameMarker(Measurements.data, Marker = "a", ChannelBinding = "ERG", New.Name = "NewMarkerName")
#' updated_measurements
#' @exportMethod RenameMarker
setGeneric(
  name = "RenameMarker",
  def = function(X,
                 Marker,
                 ChannelBinding,
                 New.Name) {
    standardGeneric("RenameMarker")
  }
)

#' @noMd
setMethod("RenameMarker",
          "ERGMeasurements",
          function(X, Marker, ChannelBinding, New.Name) {

            if(is.null(ChannelBinding)){
              stop("'ChannelBinding' should be set to any valid channel of the parent object. However, can be set to NA, if not required." )
            }

            if(!(Marker %in% MarkerNames(X))){
              stop("'Marker' does not point to any marker present in 'X'.")
            }
            if(!is.na(ChannelBinding)){
              if (!(Marker %in% MarkerNames(X)[Markers(X)$ChannelBinding ==
                                               ChannelBinding])) {
                stop("'Marker' does not point to any marker with the indicated 'ChannelBinding' present in 'X'.")
              }
              idx <- which(Markers(X)$ChannelBinding ==  ChannelBinding &  MarkerNames(X) == Marker)
              if(length(idx)!=1){
                stop("Combination of 'Marker' and 'ChannelBinding' provided does not point to a single marker. ")
              }
            } else {
              idx <- which(MarkerNames(X) == Marker)
              if(length(idx)!=1){
                stop("'Marker' does not point to a single marker. Consider setting 'ChannelBinding'.")
              }
            }
            X@Marker$Name[idx]<-New.Name
            X
          })


#' Rename a Marker in an ERGMeasurements Object
#'
#' @describeIn Marker-Methods Renames a marker within an ERGMeasurements object according to the specified channel binding.
#' @param New.Name The new name to assign to the marker.
#' @return An updated ERGMeasurements object with the marker renamed.
#' @exportMethod RenameMarker
setGeneric(
  name = "RenameMarker",
  def = function(X, Marker, ChannelBinding, New.Name)
  {
    standardGeneric("RenameMarker")
  }
)
#' @noMd
setMethod("RenameMarker",
          "ERGMeasurements",
          function(X, Marker, ChannelBinding, New.Name) {
            # Check if the channel binding is provided and valid
            if(is.null(ChannelBinding) || !(ChannelBinding %in% unique(X@Marker$ChannelBinding))) {
              stop("'ChannelBinding' should be present in the object. It cannot be NA.")
            }

            # Locate the marker within the specified channel
            idx <- which(X@Marker$Name == Marker & X@Marker$ChannelBinding == ChannelBinding)

            # Check if exactly one marker is identified to be renamed
            if(length(idx) != 1) {
              stop("The combination of 'Marker' and 'ChannelBinding' provided does not point to a single marker. Please check the values.")
            }

            # Perform the renaming
            X@Marker$Name[idx] <- New.Name

            # Validate and return the updated object
            if(!validObject(X)) {
              stop("Object validation failed after renaming marker.")
            }

            return(X)
          })


#' @describeIn Marker-Methods Get Markers.
#' @exportMethod Markers
#' @noMd
setGeneric(
  name = "Markers",
  def = function(X)
  {
    standardGeneric("Markers")
  }
)
#' @noMd
setMethod("Markers",
          "ERGMeasurements",
          function(X) {
            df<-X@Marker
            df$Relative<-df$Name[df$Relative]
            return(df)
          })
#' @noMd
setMethod("Markers",
          "ERGExam",
          function(X) {
            return(Markers(X@Measurements))
          })
```
