## Supplemental Material 1 for "The ERGtools2 package: A Toolset for Processing and Analysing Visual Electrophysiology Data": MeasurementMethots.R

```
#' Access and Modify the Marker Measurements Stored in an ERGExam or ERGMeasurements Object
#'
#' These methods enable modification of the marker measurements (hereafter short: measurements) stored in an \linkS4class{ERGExam} object or the underlying \linkS4class{ERGMeasurements} object directly.
#' \strong{Note:} the \code{Measurements(X, ...) <- value} methods require that all arguments are provided. Set to NULL if not needed.
#'
#' @inheritParams Marker-Methods
#' @inheritParams Where
#' @param value For Measurements<- only. Plain numeric value or value of class units (\link[units:units]{units::units} with a time unit set. If \code{NULL} it will remove the indicated row.
#' @param data For newERGMeasurements only. A data Measurementsframe containing measurements data with columns:
#'   \code{Channel}, \code{Name}, \code{Recording}, \code{Time}, and \code{Relative}.
#' @param measure.absolute Logical, default: FALSE. If absolute amplitudes should be returned instead of amplitudes relative to the reference marker (where given).
#' @param create.marker.if.missing Logical. If TRUE and the marker does not exist, it will be created. Dies nothing if value is set to \code{NULL}
#' @param update.empty.relative Logical. If an empty relative value should be overwritten by an otherways matching marker.
#' @param quiet For Measurements only. Logical. If TRUE, suppresses warnings and progress bars.
#' @param TimesOnly For Measurements only. Logical. If TRUE, only fetches times, not amplitudes
#' @return An updated version of the object, except for Measurements, which returns a data.frame representing the Measurements stored in the Measurements slot of the \linkS4class{ERGExam} object or the \linkS4class{ERGMeasurements} object-
#' @seealso \link[EPhysData:EPhysSet-class]{EPhysData::EPhysSet-class} \link{Measurements-Methods} \link{Get}
#' @examples
#' data(Measurements.data)
#' # Get Measurements table for specific recording and marker in ERGMeasurements object
#' Markers(Measurements.data)
#' Measurements(X = Measurements.data, where = 1, Marker = "a", quiet = TRUE)
#' #' but this would fail with an informative error message
#' #' Measurements(X = Measurements.data, where = 1, Marker = "X", quiet = F)
#'
#' # Get Measurements table for specific recording and marker in ERGExam object
#' data(ERG)
#' ERG<-SetStandardFunctions(ERG)
#' Measurements(ERG)
#' Measurements(ERG, measure.absolute = T)
#' Measurements(ERG, TimesOnly = T)
#'
#' @name Measurements-Methods
NULL

#' Get Measurements table
#' @describeIn Measurements-Methods Get Measurements table

#' @exportMethod Measurements
setGeneric(
  name = "Measurements",
  def = function(X,
                 where = NULL,
                 Marker =  NULL,
                 quiet = F,
                 ...)
  {
    standardGeneric("Measurements")
  }
)

#' @describeIn Measurements-Methods Get Measurements table from an  \linkS4class{ERGMeasurements} object
#' @noMd
setMethod("Measurements", "ERGMeasurements",
          function(X,
                   where = NULL,
                   Marker =  NULL,
                   quiet = F) {
            # integrity checks
            if (is.null(where)) {
              where <- unique(X@Measurements$Recording)
            }
            if (is.null(Marker)) {
              Marker <-  MarkerNames(X)
            }


            if (!is.character(Marker)) {
              stop("'Marker' must be a character string.")
            }
            emptyobject <- !nrow(X@Marker) > 0
            if (!emptyobject) {
              if (!all(Marker %in% MarkerNames(X)) && !quiet) {
                warning("'Marker' is not a valid marker name of a marker contained in the object.")
              }
            }

            # Merge Marker and Measurements data frames
            mark <- X@Marker
            mark$Relative <- mark$Name[mark$Relative]

            merged_df <-
              merge(
                mark,
                X@Measurements,
                by.x = 0,
                by.y = "Marker",
                all.x = TRUE,
                all.y = TRUE
              )
            merged_df <-
              merged_df[c("Recording", "Name", "ChannelBinding", "Relative", "Time")]

            merged_df <-
              merged_df[with(merged_df,
                             order(Recording, ChannelBinding, Relative, na.last =
                                     F)),]
            rownames(merged_df) <- NULL

            if (!emptyobject) {
              if(length(where)!=0){
                if (!all(where %in% merged_df$Recording) && !quiet) {
                  warning("'where' is not a valid Index of a Measurement already contained in the object.")
                }
              }
            }

            return(merged_df[merged_df$Recording %in% where &
                               merged_df$Name %in% Marker,])
          })


#' @describeIn Measurements-Methods Get Measurements table from an \linkS4class{ERGExam} object
#' @importFrom cli cli_progress_bar cli_progress_update cli_progress_done
#' @importFrom EPhysData Metadata AverageFunction
#' @noMd
setMethod("Measurements",
          "ERGExam",
          function(X,
                   where = NULL,
                   Marker =  NULL,
                   quiet = F,
                   measure.absolute = F,
                   TimesOnly = F) {

            if(!is.null(where)){
              where = Where(X, where)
            }

            if(is.null(Marker)){
              Marker<-MarkerNames(X)
            }

            # make sure also parent of relative markers are retrieved
            marker.info <- Markers(X)
            marker.info.full<-marker.info
            marker.info <- marker.info[marker.info$Name %in% Marker, ]
            Fetch<-Marker
            if(!TimesOnly){
              if (any(!is.na(marker.info$Relative))) {
                rel.marker.rows<-marker.info[!is.na(marker.info$Relative),]
                for (i in 1:nrow(rel.marker.rows)){
                  Fetch <-
                    c(Fetch, marker.info.full$Name[marker.info.full$Name == rel.marker.rows$Relative[i] &
                                                     marker.info.full$ChannelBinding == rel.marker.rows$ChannelBinding[i]])
                }

              }
            }

            measurements <- Measurements(X@Measurements, where, Fetch, quiet)
            if (nrow(measurements) != 0) {
              extracols<-ExtraMetaColumns(X)
              measurements <-
                merge(measurements,
                      Metadata(X)[, c("Step", "Channel", "Repeat",  "Eye")],
                      by =
                        "Recording",
                      by.y = 0)
              measurements <-
                merge(measurements, StimulusTable(X), by = "Step")
              measurements <-
                measurements[, c(
                  "Recording",
                  "Step",
                  "Description",
                  "Channel",
                  "Repeat",
                  "Eye",
                  "Name",
                  "Relative",
                  "Time"
                )]

              measurements <-
                measurements[with(measurements,
                                  order(Recording, Step, Channel, Eye, Relative, na.last =
                                          F)), ]
              measurements$Voltage <- as_units(NA, "uV")

              if (!TimesOnly) {
                if (!quiet) {
                  cli_progress_bar("Retrieving record values for the given time points.", total = length(X@Data),  clear = TRUE, auto_terminate = F)
                }
                for (i in 1:nrow(measurements)) {
                  sel <- X[[measurements$Recording[i]]]
                  if (length(AverageFunction(sel)(1:3)) != 1 &&
                      dim(sel)[2] != 1) {
                    Notice(X,
                           where = measurements$Recording[i],
                           what = "E",
                           notice_text = "You need to set an averaging function before performing Marker and Measuremnt operations. E.g. run {.run SetStandardFunctions({deparse(substitute(x))})}")
                  }
                  if (!is.na(measurements$Time[i])) {
                    Voltage <-
                      GetData(sel, Time = TimeTrace(sel)[which.min(abs(TimeTrace(sel) - measurements$Time[i]))], Raw = F)
                    if (!is.na(measurements$Relative[i]) &&
                        !measure.absolute) {
                      rel.marker.time <-
                        measurements$Time[measurements$Recording == measurements$Recording[i] &
                                            measurements$Name == measurements$Relative[i]]
                      Voltage <- Voltage - GetData(sel,
                                                   Time = TimeTrace(sel)[which.min(abs(TimeTrace(sel) - rel.marker.time))],
                                                   Raw = F)
                    }

                    if (all(dim(Voltage) == 1)) {
                      measurements$Voltage[i] <- set_units(Voltage[1, 1], "uV")
                    } else{
                      Notice(X,
                             where = measurements$Recording[i],
                             what = "E",
                             notice_text = "GetData returns multiple values.")
                    }
                  } else {
                    Voltage <- set_units(NA, "uV")
                  }
                  if (!quiet) {
                    cli_progress_update()
                  }
                }

                if (!quiet) {
                  cli_progress_done()
                }
              }
              measurements<-merge(measurements,Metadata(X)[,c(extracols),drop=F],by.x = "Recording", by.y = 0)

              if(!TimesOnly){
                measurements<-measurements[measurements$Name %in% Marker,]
              }


            } else {
              measurements <-
                data.frame(
                  Recording = numeric(),
                  Step = character(),
                  Description = character(),
                  Channel = character(),
                  Repeat = numeric(),
                  Eye = character(),
                  Name = character(),
                  Relative = character(),
                  Time = as_units(numeric(), "ms"),
                  Voltage = as_units(numeric(), "uV")
                )
            }
            return(measurements)
          })

#' Update or add a line to the Measurements slot
#'
#' @describeIn Measurements-Methods Update, add, or remove a line to the Measurements slot
#' @examples
#' # Set Measurements for specific recording and marker in ERGMeasurements object
#' Markers(Measurements.data)
#' Measurements(X = Measurements.data, Marker = "B" ,where = 1)<-99
#' # this would fail, as marker does not exist
#' # Measurements.data<-Measurements(X = Measurements.data, Marker = "c" , where = 1, Relative = "a")<-150
#' # But this would work
#' Measurements(X = Measurements.data, Marker = "c" ,where = 1,Relative = "a", ChannelBinding="ERG")<-150
#' # Set Measurements table for specific recording and marker in ERGExam object
#' ERG<-SetStandardFunctions(ERG)
#' Measurements(ERG)
#' @exportMethod Measurements<-
setGeneric(
  name = "Measurements<-",
  def = function(X,
                 Marker,
                 where,
                 create.marker.if.missing = T,
                 Relative = NULL,
                 ChannelBinding = NULL,
                 value)
  {
    standardGeneric("Measurements<-")
  }
)

#' @describeIn Measurements-Methods Update, add, or remove a measurement from an \linkS4class{ERGMeasurements} object
#' @noMd
#' @importFrom units deparse_unit set_units as_units units_options
setMethod("Measurements<-",
          signature = "ERGMeasurements",
          function(X,
                   Marker,
                   where,
                   create.marker.if.missing = T,
                   Relative = NULL,
                   ChannelBinding = NULL,
                   value) {            #validity checks and marker Marker conversion

            # internal functions
            get.marker.idx<-function(cb,X,Marker){
              if(!is.null(cb)){
                marker.idx <- which(Markers(X)$Name == Marker & Markers(X)$ChannelBinding == cb)
              } else {
                marker.idx <- which(Markers(X)$Name == Marker)
              }
              if(length(marker.idx)!=1){
                stop("Marker is not unabiguously defined by the given paramaters ('Marker' and possibly 'ChannelBinding').")
              }
              return(marker.idx)
            }

            #validity checks
            if (length(Marker) != 1) {
              stop ("'Marker' must contain a single value.")
            }
            if (nrow(Measurements(X, where = where, Marker = Marker, quiet = T)) > 1) {
              stop ("'where' must point to a single or non existent measurement.")
            }
            if (length(value) != 1 && !is.null(value)) {
              stop ("value' must contain a single value or NULL.")
            }
            if (!is.character(Marker)) {
              stop("'Marker' must be a character string.")
            }
            if (!is.numeric(where)) {
              stop("'where' must be numeric.")
            }

            if(!is.null(value)){
              # if the aim is not to delete a measurement (value!=NULL)
              if (!is.numeric(value) && !is.null(value)) {
                stop("'value' must be numeric or NULL.")
              }else{
                refunit<-X@Measurements$Time[1]
                refunit<-deparse_unit(refunit)
                if (!("units" %in% class(value))) {
                  value <-
                    as_units(value, refunit, mode = "standard")
                } else{
                  if(deparse_unit(value)!=refunit){
                    units_options(set_units_mode = "standard")
                    value <- set_units(value, refunit)
                  }
                }
              }

              nrow.same<-nrow(Measurements(X, where = where, Marker = Marker,quiet = T))

              if (nrow.same ==
                  1) {    # Update Measurement
                marker.idx<-get.marker.idx(ChannelBinding,X,Marker)

                if (length(X@Measurements$Time[X@Measurements$Recording == where &
                                               X@Measurements$Marker %in% marker.idx]) != 1) {
                  stop("Unexpected error during marker selection.")
                }
                X@Measurements$Time[X@Measurements$Recording == where &
                                      X@Measurements$Marker %in% marker.idx] <- value
              }
              if (nrow.same ==
                  0) {   # add Measurement
                if (!(Marker %in% MarkerNames(X))) {
                  if (create.marker.if.missing) {
                    X <-
                      AddMarker(
                        X,
                        Marker = Marker,
                        Relative = which(MarkerNames(X)==Relative),
                        ChannelBinding = ChannelBinding
                      )
                  } else{
                    stop(paste0("Marker '", Marker,"' does not exist in object."))
                  }
                }

                marker.idx<-get.marker.idx(ChannelBinding,X,Marker)
                new.measurement <- data.frame(Recording = where,
                                              Marker = marker.idx,
                                              Time = value)
                X@Measurements <-
                  rbind(X@Measurements, new.measurement)

              }
              if (nrow.same > 2) {
                stop(
                  "Malformed ERGMeasurements object. Multiple entries found for the given combination of 'where' and 'Marker'"
                )
              }
            } else {
              # if value is NULL --> delete the selected measurement
              marker.idx<-get.marker.idx(ChannelBinding,X,Marker)
              # are markers dependent on the one to be deleted?
              marker.dept.idx<-which(X@Marker$Relative==marker.idx)
              if(any(X@Measurements$Recording == where &
                     X@Measurements$Marker %in% marker.dept.idx)){
                stop("Measurement cant be deleted because other measurements depend on it (dependent/relative markers).")
              }

              X@Measurements <-
                X@Measurements[!(X@Measurements$Recording == where &
                                   X@Measurements$Marker %in% marker.idx),]
              rownames(X@Measurements)<-NULL

            }

            if (validObject(X)) {
              return(X)
            }else{
              stop("object validation failed")
            }
          })

#' @describeIn Measurements-Methods Update, add, or remove a measurement from the Measurements slot of an  \linkS4class{ERGExam} object
#' @noMd
#' @importFrom units deparse_unit set_units as_units
setMethod("Measurements<-",
          signature = "ERGExam",
          function(X,
                   Marker,
                   where = NULL,
                   create.marker.if.missing = T,
                   Relative = NULL,
                   ChannelBinding = NULL,
                   value){

            Measurements(X@Measurements,
                         Marker = Marker,
                         where = Where(X,where = where),
                         create.marker.if.missing = create.marker.if.missing,
                         Relative = Relative,
                         ChannelBinding = ChannelBinding
            ) <- value

            if (validObject(X)) {
              return(X)
            }else{
              stop("Object validation failed for unknown reason.")
            }

          })
#' @describeIn Measurements-Methods Remove a measurement from the Measurements slot of an \linkS4class{ERGExam} object
#' @noMd
#' @exportMethod DropMeasurement
setGeneric(
  name = "DropMeasurement",
  def = function(X,
                 Marker,
                 where,
                 ChannelBinding = NULL)
  {
    standardGeneric("DropMeasurement")
  }
)
#' @describeIn Measurements-Methods Remove a measurement from the Measurements slot of an \linkS4class{ERGExam} object
#' @noMd
setMethod("DropMeasurement",
          signature = "ERGExam",
          function(X,
                   Marker,
                   where = NULL,
                   ChannelBinding = NULL){
            Measurements(X = X, Marker = Marker, where = where, create.marker.if.missing = T, Relative = NULL, ChannelBinding = ChannelBinding)<-NULL
          })


#' Create an ERGMeasurements object from data
#'
#' @describeIn Measurements-Methods #' Create an ERGMeasurements object from data
#' @examples
#' # Example data frame
#' data <- data.frame(
#'   Channel = c("ERG", "ERG", "ERG", "ERG"),
#'   Name = c("Marker1", "Marker2", "Marker1", "Marker2"),
#'   Recording = c(1, 1, 2, 2),
#'   Time = c(10, 12, 15, 18),
#'   Relative = c(NA, "Marker1", NA, "Marker1")
#' )
#' # Create ERGMeasurements object
#' newERGMeasurements(data)
#'
#' @export newERGMeasurements
newERGMeasurements <- function(data, update.empty.relative = F) {

  if (!is.data.frame(data)) {
    stop("'data' must be a data frame.")
  }
  required_columns <- c("Channel", "Name", "Recording", "Time", "Relative")
  if (!all(required_columns %in% names(data))) {
    stop("Data frame must contain all required columns: ", paste(required_columns, collapse = ", "))
  }

  M <- new("ERGMeasurements")
  # first add independent markers
  for (c in unique(data$Channel[is.na(data$Relative)])) {
    for (m in unique(data$Name[data$Channel == c &
                               is.na(data$Relative)])) {
      rel <- NA
      # data[data$Channel == c &
      #        data$Name == m, ]
      M <- AddMarker(M, m, rel, c, update.empty.relative = update.empty.relative)
      for (r in data$Recording[data$Channel == c &
                               data$Name == m]) {
        suppressWarnings({
          Measurements(
            X = M,
            Marker = m,
            where = r,
            Relative = rel,
            create.marker.if.missing = F,
            ChannelBinding = c
          ) <-
            data$Time[data$Channel == c &
                        data$Name == m &
                        data$Recording == r]
        })
      }
    }
  }
  #then dependent markers
  for (c in unique(data$Channel[!is.na(data$Relative)])) {
    for (m in unique(data$Name[data$Channel == c &
                               !is.na(data$Relative)])) {
      rel <-
        unique(data$Relative[data$Channel == c &
                               data$Name == m])
      if(length(rel)>1){
        rel<-rel[!is.na(rel)]
      }
      if(length(rel)>1){
        rel<-rel[1]
        warning("More than one matching relatve reference found. Taking first none-NA")
      }
      rel.idx<-which(Markers(M)==rel)
      M <- AddMarker(M, m, rel.idx, c,update.empty.relative = update.empty.relative)
      for (r in data$Recording[data$Channel == c &
                               data$Name == m]) {
        Measurements(
          M,
          Marker = m,
          where = r,
          create.marker.if.missing = F,
          ChannelBinding = c
        ) <-
          data$Time[data$Channel == c &
                      data$Name == m &
                      data$Recording == r]
      }
    }
  }
  if (validObject(M)) {
    return(M)
  }
}

#' @describeIn Measurements-Methods Clears the Measurements slots in an  link[=ERGExam]{ERGExam} object (both, markers and measurements.
#' @importFrom methods validObject
#' @exportMethod ClearMeasurements
setGeneric(
  name = "ClearMeasurements",
  def = function(X) {
    standardGeneric("ClearMeasurements")
  }
)
#' @noMd
setMethod("ClearMeasurements",
          signature(X = "ERGExam"),
          function(X) {
            X@Measurements <- new("ERGMeasurements")
            if (validObject(X)) {
              return(X)
            }
          })


#' Measurements-Methods Converts the output of the Measurements method from relative to absolute amplitudes
#' @keywords Internal
#' @noMd
#' @noRd
ConvertMeasurementsToAbsolute <- function(data) {
  for (i in seq_along(data$Voltage)) {
    if (!is.na(data$Relative[i])) {
      match_row <- which(data$Description == data$Description[i] &
                           data$Eye == data$Eye[i] &
                           data$Channel == data$Channel[i] &
                           data$Name == data$Relative[i])

      if (length(match_row) > 0) {
        data$Voltage[i] <- data$Voltage[i] + data$Voltage[match_row]
      }
    }
  }
  #colnames(data)[colnames(data)=="Relative"]<-".Relative"
  return(data)
}
```
