## Supplemental Material 1 for "The ERGtools2 package: A Toolset for Processing and Analysing Visual Electrophysiology Data": MergeERGExams.R

```
#' Merge ERGExams
#'
#' Merges two or more ERGExam objects into a single ERGExam object.
#'
#' @param ERGExam_list A list of ERGExam objects to be merged with \code{X}.
#' @param mergemethod . Depreciated.
#' @return An ERGExam object representing the merged data. the file names, protocol names and dates from the individual files merged are stored as additional columns in the Stimulus table. Use \code{Metadata()} to see those.
#' @noMd
#' @examples
#' # Merge two ERGExams
#' data(ERG)
#' # make some ERGExams objects that differ
#' exam1<-Subset(ERG,where=list(Step=as.integer(c(1,2,3))))
#' exam2<-Subset(ERG,where=list(Eye="RE"))
#' # merge
#' merged_exam <- MergeERGExams(list(exam1, exam2))#'
#' @export
MergeERGExams <- function(ERGExam_list, mergemethod = "Append") {
  message("Merging exams is experimental.")
  if (!("list" %in% class(ERGExam_list))) {
    stop("'ERGExam_list' must be a list of 'ERGExam' objects")
  }
  if (!all(unlist(lapply(ERGExam_list, class)) %in% "ERGExam")) {
    stop("'ERGExam_list' must be a list of 'ERGExam' objects")
  }
  out <- ERGExam_list[[1]]
  rest <- ERGExam_list[2:length(ERGExam_list)]
  for (ex in rest) {
    tryCatch(
      out <- merge2ERGExams(out, ex),
      error = function(e) {
        stop("Mergin Exams failed for ",
             Subject(ex), ", ",
             ExamDate(ex), ", ",
             ProtocolName(ex),
             " with message: ",
             e)
      }
    )
  }
  return(out)

}

#' @noMd
#' @keywords internal

merge2ERGExams <- function(exam1, exam2, mergemethod = "Append") {
  if ((exam1@Averaged != exam2@Averaged)) {
    stop("Objects to merge contain averaged and unaveraged data.")
  }

  # Validity checks
  if (!identical(exam1@SubjectInfo, exam2@SubjectInfo)) {
    stop("SubjectInfo must be identical in both exams.")
  }

  exam1Date <- as.POSIXct(exam1@ExamInfo$ExamDate)
  exam2Date <- as.POSIXct(exam2@ExamInfo$ExamDate)
  if (max(abs(difftime(exam1Date, exam2Date, units = "hours"))) > 3) {
    stop("ExamInfo$ExamDate should differ by a maximum of 3 hours.")
  }

  #
  # make joint stimulus table
  #

  stimtab1 <- StimulusTable(exam1)
  if (is.null(stimtab1$ProtocolName)) {
    stimtab1$ProtocolName <- ProtocolName(exam1)
  }
  stimtab2 <- StimulusTable(exam2)
  if (is.null(stimtab2$ProtocolName)) {
    stimtab2$ProtocolName <- ProtocolName(exam2)
  }

  stimtab1$OrigStep1 <- stimtab1$Step
  stimtab2$OrigStep2 <- stimtab2$Step

  stimtab.merged <-
    merge(
      stimtab2,
      stimtab1,
      by = c(
        "Description",
        "StimulusEnergy",
        "Background",
        "Type",
        "ProtocolName"
      ),
      all = TRUE
    )
  stimtab.merged$Step.x <- NULL
  stimtab.merged$Step.y <- NULL
  stimtab.merged$Step <- 1:nrow(stimtab.merged)

  #
  # make joint metadata table
  #

  md1 <- Metadata(exam1)
  md2 <- Metadata(exam2)

  # add possible extra columns in Metadata
  if (!isTRUE(all.equal(colnames(md1), colnames(md2)))) {
    extra.in.exam1 <- colnames(md1)[!(colnames(md1) %in% colnames(md2))]
    extra.in.exam2 <-
      colnames(md2)[!(colnames(md2) %in% colnames(md1))]
    if (length(extra.in.exam1) > 0) {
      for (e in extra.in.exam1) {
        exam2@Metadata[, e] <- as.numeric(NULL)
        class(exam2@Metadata[, e]) <- class(exam1@Metadata[, e])
      }
    }
    if (length(extra.in.exam2) > 0) {
      for (e in extra.in.exam2) {
        exam1@Metadata[, e] <- as.numeric(NULL)
        class(exam1@Metadata[, e]) <- class(exam2@Metadata[, e])
      }
    }
  }

  # add source info
  if (is.null(md1$ExamDate)) {
    md1$ExamDate <- ExamDate(exam1)
    md1$Filename <- exam1@ExamInfo$Filename
    md1$ProtocolName <- ProtocolName(exam1)
    md1$Version <- exam1@ExamInfo$Version
    md1$Investigator <- exam1@ExamInfo$Investigator
  }

  if (is.null(md2$ExamDate)) {
    md2$ExamDate <- ExamDate(exam2)
    md2$Filename <- exam2@ExamInfo$Filename
    md2$ProtocolName <- ProtocolName(exam2)
    md2$Version <- exam2@ExamInfo$Version
    md2$Investigator <- exam2@ExamInfo$Investigator
  }

  md1$OrigRecording <- 1:nrow(md1)
  md2$OrigRecording <- 1:nrow(md2)

  md1$SRC <- 1
  md2$SRC <- 2

  # update Step
  for (i in 1:nrow(stimtab.merged)) {
    if (!is.na(stimtab.merged$OrigStep1[i])) {
      md1$Step[md1$Step == stimtab.merged$OrigStep1[i]] <-
        stimtab.merged$Step[i]
    }
  }
  for (i in 1:nrow(stimtab.merged)) {
    if (!is.na(stimtab.merged$OrigStep1[i])) {
      md2$Step[md2$Step == stimtab.merged$OrigStep1[i]] <-
        stimtab.merged$Step[i]
    }
  }

  combined <- rbind(md1, md2)
  md.merged <- data.frame()
  # Process each row of the combined data frame
  for (i in 1:nrow(combined)) {
    row <- combined[i,]
    existing_row_index <- which(
      md.merged$Step == row$Step &
        md.merged$Channel == row$Channel &
        md.merged$Eye == row$Eye &
        md.merged$Recording == row$Recording
    )

    if (length(existing_row_index) > 0) {
      row$Repeat <-
        row$Repeat  + max(md.merged$Repeat[existing_row_index])
      md.merged <- rbind(md.merged, row)
    } else {
      md.merged <- rbind(md.merged, row)
    }
  }

  #
  # merge data
  #

  data.merged <- list()
  for (i in 1:nrow(md.merged)) {
    if (md.merged$SRC[i] == 1) {
      data.merged[[i]] <- exam1@Data[[md.merged$OrigRecording[i]]]
    }
    if (md.merged$SRC[i] == 2) {
      data.merged[[i]] <- exam2@Data[[md.merged$OrigRecording[i]]]
    }
  }

  #
  # merge measurements
  #

  idx.updater <-
    data.frame(cbind(
      new = 1:nrow(md.merged),
      orig = md.merged$OrigRecording,
      src = md.merged$SRC
    ))
  idx.updater <- idx.updater[idx.updater$src == 2, c("new", "orig")]

  mergedMeasurements <-
    merge2ERGMeasurements(
      exam1@Measurements,
      exam2@Measurements,
      obj2.recording.index.old = idx.updater$orig,
      obj2.recording.index.new = idx.updater$new
    )

  #
  # Cleanup
  #

  rm(combined)
  stimtab.merged$OrigStep1 <- NULL
  stimtab.merged$OrigStep2 <- NULL
  md.merged$OrigRecording <- NULL
  md.merged$SRC <- NULL

  #
  # Create a new ERGExam instance with merged data and metadata
  #

  examinfo <- exam1@ExamInfo
  examinfo$ProtocolName <-
    paste(exam1@ExamInfo$ProtocolName,
          exam2@ExamInfo$ProtocolName,
          sep = " and ")
  examinfo$Filename <- "Merged Exam"
  mergedExam <- newERGExam(
    Data = data.merged,
    Metadata = md.merged,
    Stimulus = stimtab.merged,
    Averaged = exam1@Averaged,
    Measurements = mergedMeasurements,
    ExamInfo = examinfo,
    SubjectInfo = exam1@SubjectInfo
  )

  return(mergedExam)
}

#' Merge two ERGMeasurements objects
#'
#' This function merges two ERGMeasurements objects by combining their Marker and Measurements data frames.
#' It also provides options to update recording indices and handle marker mismatches.
#'
#' @param obj1 An object of class ERGMeasurements.
#' @param obj2 Another object of class ERGMeasurements.
#' @param increment.obj2.recording.index.by Depreciated. Numeric indicating by what number to increment the recording indices by. Could be e.g. the number of recordings in object 1.
#' @param obj2.recording.index.old,obj2.recording.index.old Two vectors of the same length indicating how the obj2 recording indices hould be updated.
#' @noMd
#' @return An object of class ERGMeasurements, representing the merged data.
#' @keywords internal
#' @examples
#'
#' # load the example Measurements object
#' data(Measurements.data)
#'
#' # create a second Measurements object
#' # Create marker data frame
#' marker_df <- data.frame(
#'   Name = c("N1", "P1", "a", "C"),
#'   Relative = c(NA, 1, NA, 3),
#'   ChannelBinding =c("VEP","VEP","ERG","ERG")
#' )
#'
#' # Create measurements data frame
#' measurements_df <- data.frame(
#'   Recording = c(1, 1, 4, 4, 3, 3),
#'   Marker = c(1, 2, 3, 4, 1, 2),
#'   Time = as_units(c(10, 40, 19, 26, 34, 31), "ms")
#' )
#'
#' # Create ERGMeasurements object
#' Measurements.data2 <- new("ERGMeasurements", Marker = marker_df, Measurements = measurements_df)
#' # Show the object
#' Measurements.data2
#'
#' # now merge both objects
#' merge2ERGMeasurements(Measurements.data,Measurements.data2, max(Measurements(Measurements.data)[,"Recording"]))
#'
#' # other example
#'
#' marker_df3 <- data.frame(
#' Name = c("N2", "P2", "b", "D", "E"),
#' Relative = c(NA, 1, NA, 3, 3),
#' ChannelBinding = c("VEP", "VEP", "ERG", "ERG", "ERG")
#' )
#'
#' measurements_df3 <- data.frame(
#'   Recording = c(1, 1, 4, 4, 5, 5, 3, 3),
#'   Marker = c(1, 2, 3, 4, 3, 5, 1, 2),
#'   Time = as_units(c(10, 40, 19, 26, 34, 31, 15, 25), "ms")
#' )
#'
#' Measurements.data3 <- new("ERGMeasurements", Marker = marker_df3, Measurements = measurements_df3)
#'
#' merge2ERGMeasurements(Measurements.data2,Measurements.data3,max(Measurements(Measurements.data2)[,"Recording"]))
#'
#'
#' @export merge2ERGMeasurements
merge2ERGMeasurements <-
  function(obj1,
           obj2,
           increment.obj2.recording.index.by = NULL,
           obj2.recording.index.old = NULL,
           obj2.recording.index.new = NULL) {
    if (!validObject(obj1) || !validObject(obj2)) {
      stop("One or both of the objects are not valid ERGMeasurements objects.")
    }
    if(length(MarkerNames(obj1))==0 || length(MarkerNames(obj2)==0)){
      if(length(MarkerNames(obj1))==0){
        return(obj2)
      }
      if(length(MarkerNames(obj2))==0){
        return(obj1)
      }
    }

    obj1.m <- obj1@Marker
    rownames(obj1.m) <- NULL
    obj1.m$obj1.idx <- 1:nrow(obj1.m)
    obj2.m <- obj2@Marker
    rownames(obj2.m) <- NULL
    obj2.m$obj2.idx <- 1:nrow(obj2.m)
    pointer.updates.for.obj2 <-
      merge(obj1.m[, c("Name", "ChannelBinding", "obj1.idx")], obj2.m[, c("Name", "ChannelBinding", "obj2.idx")])[, c("obj1.idx", "obj2.idx")]

    #Check parent markers of relatives match
    obj2.rel <- obj2.m$Relative[pointer.updates.for.obj2$obj2.idx]
    obj2.rel <- obj2.rel[!is.na(obj2.rel)]
    pointer.pairs <-
      pointer.updates.for.obj2[match(obj2.rel, pointer.updates.for.obj2$obj2.idx), ]
    if (!(all(is.na(pointer.pairs)))) {
      for (i in 1:nrow(pointer.pairs)) {
        if (!all(obj1.m[pointer.pairs$obj1.idx[i], c("Name", "ChannelBinding")] == obj2.m[pointer.pairs$obj2.idx[i], c("Name", "ChannelBinding")])) {
          obj1.probl <-
            obj1.m[obj1.m$Relative %in% pointer.pairs$obj1.idx[i],]
          obj2.probl <-
            obj2.m[obj2.m$Relative %in% pointer.pairs$obj2.idx[i],]
          stop(
            "At least 1 marker mismatch was detected: Marker '",
            obj1.probl[1, 1],
            "' with ChannelBinding '",
            obj1.probl[1, 3],
            "' from 'obj1' is relative to marker '",
            obj1.m$Name[pointer.pairs$obj1.idx[i]] ,
            "', while the corresponding marker in 'obj2' is relative to marker '",
            obj2.m$Name[pointer.pairs$obj2.idx[i]]  ,
            "'. "
          )
        }
      }
    }

    # for appending
    obj2.idx <-
      which(!(1:nrow(obj2.m) %in% pointer.updates.for.obj2$obj2.idx))
    if (length(obj2.idx) > 0) {
      obj1.idx <- nrow(obj1.m) + 1:length(obj2.idx)
      pointer.appending.for.obj2 <-
        data.frame(obj2.idx = obj2.idx, obj1.idx = obj1.idx)

      # append new and update relative column
      for (i in 1:nrow(pointer.appending.for.obj2)) {
        if (obj2.m$Relative[pointer.appending.for.obj2$obj2.idx[i]] %in% pointer.updates.for.obj2$obj2.idx) {
          obj2.m$Relative[pointer.appending.for.obj2$obj2.idx[i]] <-
            pointer.updates.for.obj2$obj1.idx[pointer.updates.for.obj2$obj2.idx == obj2.m$Relative[pointer.appending.for.obj2$obj2.idx[i]]]
        }
        if (obj2.m$Relative[pointer.appending.for.obj2$obj2.idx[i]] %in% pointer.appending.for.obj2$obj2.idx) {
          obj2.m$Relative[pointer.appending.for.obj2$obj2.idx[i]] <-
            pointer.appending.for.obj2$obj1.idx[pointer.appending.for.obj2$obj2.idx == obj2.m$Relative[pointer.appending.for.obj2$obj2.idx[i]]]
        }
        obj1.m[pointer.appending.for.obj2$obj1.idx[i], ] <-
          obj2.m[pointer.appending.for.obj2$obj2.idx[i], ]
      }
    }

    obj1.m$obj1.idx <- NULL

    # for updating
    obj2.measurements <- obj2@Measurements
    obj2.measurements <-
      merge(
        obj2.measurements,
        pointer.updates.for.obj2,
        by.x = "Marker",
        by.y = "obj2.idx",
        all.x = T
      )

    if (length(obj2.idx) > 0) {
      obj2.measurements <-
        merge(
          obj2.measurements,
          pointer.appending.for.obj2,
          by.x = "Marker",
          by.y = "obj2.idx",
          all.x = T
        )
    }
    obj2.measurements$Marker[!is.na(obj2.measurements$obj1.idx.x)] <-
      obj2.measurements$obj1.idx.x[!is.na(obj2.measurements$obj1.idx.x)]
    obj2.measurements$Marker[!is.na(obj2.measurements$obj1.idx.y)] <-
      obj2.measurements$obj1.idx.y[!is.na(obj2.measurements$obj1.idx.y)]

    # update recording indices.
    if(!is.null(obj2.recording.index.old)){
      if(length(obj2.recording.index.old)!=length(obj2.recording.index.new)){
        stop("'obj2.recording.index.old' and 'obj2.recording.index.new' must have the same length.")
      }
      if(!all(obj2.measurements$Recording) %in% obj2.recording.index.old){
        stop("All recording indices present in 'obj2' must be matched by 'obj2.recording.index.old'.")
      }

      for (i in 1:nrow(obj2.measurements)) {
        obj2.measurements$Recording[i] <-
          obj2.recording.index.new[obj2.recording.index.old == obj2.measurements$Recording[i]]
      }

      obj2.measurements <-
        obj2.measurements[, c("Recording", "Marker", "Time")]

      out <- new("ERGMeasurements")
      out@Marker <- obj1.m
      out@Measurements <- rbind(obj1@Measurements, obj2.measurements)

    } else {
      if (!is.null(increment.obj2.recording.index.by)) {
        obj2.measurements$Recording <-
          obj2.measurements$Recording + increment.obj2.recording.index.by

        obj2.measurements <-
          obj2.measurements[, c("Recording", "Marker", "Time")]

        out <- new("ERGMeasurements")
        out@Marker <- obj1.m
        out@Measurements <- rbind(obj1@Measurements, obj2.measurements)
      }
    }


    if (validObject(out)) {
      return(out)
    } else {
      stop("Merging ERGMeasurements object failed for unknown reaons.")
    }
  }
```
