## Supplemental Material 1 for "The ERGtools2 package: A Toolset for Processing and Analysing Visual Electrophysiology Data": Notice.R

```
#' Provide a Notice for an ERGExam Object
#'
#' The `Notice` function dynamically generates a message (warning, error, or information) based on the contents of an `ERGExam` object. The function formats the message with metadata from the `ERGExam` object, allowing for additional customizable notice text and an optional help page reference.
#'
#' @param X An `ERGExam` object.
#' @param where Specifies the row of metadata to display. This parameter is passed to the `Where` function to identify the appropriate metadata row.
#' @param what A character string specifying the type of notice to issue. Options are `"Warning"`, `"Error"`, or `"Information"`. Depending on the choice, the message will be formatted with a corresponding symbol (`!` for warning and error, `i` for information).
#' @param notice_text Additional lines of text to append to the message. This can be used to provide further details.
#' @param help_page An optional help page reference. If provided, the function adds a suggestion to see the specified help page for more information.
#'
#' @return A dynamically generated message that is either a warning, an error, or an information notice. The message includes subject and exam date information, metadata details from the specified row, and any additional lines of text or help references provided.
#'
#' @examples
#' \dontrun{
#' data(ERG)
#' Notice(ERG, where = 1, what = "Warning", notice_text = c("Check your input data."))
#' Notice(ERG, where = 2, what = "Error", help_page = "ERGtools2::ERGExam-class")
#' }
#'
#' @seealso
#' \code{\link{Where}}, \code{\link{Metadata}}
#'
#' @keywords internal
setGeneric(
  name = "Notice",
  def = function(X,
                 where = NULL,
                 what = c("Warning", "Error", "Information"),
                 notice_text = c(),
                 help_page = NULL) {
    standardGeneric("Notice")
  }
)
#' @importFrom cli cli_inform
#' @noMd
setMethod("Notice",
          "ERGExam",
          function(X,
                   where = NULL,
                   what = c("Warning", "Error", "Information"),
                   notice_text = c(),
                   help_page = NULL) {
            what <- match.arg(what, c("Warning", "Error", "Information"))
            base_message <- c("{what} for: ERGExam Object {.strong { Subject(X)}}, DoE {.emph {ExamDate(X)}}")
            if (!is.null(where)) {
              idx <- Where(X, where)
              md <- Metadata(X)[idx, ]
              base_message <- paste(
                base_message,
                "in Step {.emph {unique(md[,'Step'])}}, Channel {.emph {unique(md[,'Channel'])}}, Repeat {.emph {unique(md[,'Repeat'])}}, Eye {.emph {unique(md[,'Eye'])}} (Simulus {.emph {StimulusDescription(X, where=idx)}})"
              )
            }

            full_message <-
              c(base_message, notice_text)

            if(!is.null(help_page)){
              full_message <-
                c(full_message, "i See {.topic {help_page}} for further informations.")
            }

            # Create a composite environment
            combined_env <- new.env(parent = parent.frame())
            # Assign local variables to the composite environment
            assign("what", what, envir = combined_env)
            if (!is.null(help_page)) {
              assign("help_page", help_page, envir = combined_env)
            }
            assign("X", X, envir = combined_env)
            if (!is.null(where)) {
              assign("idx", idx, envir = combined_env)
              assign("md", md, envir = combined_env)
            }

            if (what == "Warning") {
              cli_warn(full_message, .envir = combined_env)
            } else if (what == "Error") {
              cli_abort(full_message, .envir = combined_env)
            } else if (what == "Information") {
              cli_inform(full_message, .envir = combined_env)
            }
          })
```
