## Supplemental Material 1 for "The ERGtools2 package: A Toolset for Processing and Analysing Visual Electrophysiology Data": SetStandardFunctions.R

```
#' Set processing functions for ERGExam objects
#'
#' These methods are used to set filter, rejection and averaging functions for \link{ERGExam} objects.
#' They allow setting the functions to a subset of recordings based on the specified conditions.
#' @inheritParams EPhysData::Get_Set_EPhysData
#' @param X An \linkS4class{ERGExam} object
#' @inheritParams Where
#' @param Stimulus.type.names A \link[base:pairlist]{base::pairlist} specifying the names identifying the different stimulus types, e.g., \code{Flash="Flash"} or \code{Flash="Blitz"}.
#' @param Channel.hierarchy The hierarchy of channels to choose by which to set the rejection parameter. Explanation: If a recording consists of more than one channel, the same trials should be rejected for each channel to ensure downstream analyses for all channels is performed on a consistent data basis. See also \link[EPhysData:Rejected]{EPhysData::Rejected}
#' @return An updated ERGExan object.
#'
#' @examples
#' data(ERG)
#' ERG<-SetStandardFunctions(ERG)
#' ggERGTrace(ERG,where=list(StimulusEnergy=1,Channel="ERG",Eye="RE"))
#'
#' AverageFunction(ERG,where=list(StimulusEnergy=1))<-min
#' ggERGTrace(ERG,where=list(StimulusEnergy=1,Channel="ERG",Eye="RE"))
#'
#'
#' @seealso \link[EPhysData:Get_Set_EPhysData]{EPhysData::Get_Set_EPhysData}, \link[EPhysMethods:autoreject.by.distance]{EPhysMethods::autoreject.by.distance}, \link[EPhysMethods:autoreject.by.signalfree]{EPhysMethods::autoreject.by.signalfree}, \link[EPhysMethods:filter.bandpass]{EPhysMethods::filter.bandpass}, \link[EPhysMethods:filter.detrend]{EPhysMethods::filter.detrend},
#' @name UpdateProcessingMethods
NULL


#' @describeIn UpdateProcessingMethods Update the FilterFunction for all Recordings in an \linkS4class{ERGExam}, or only those slected using \code{where}.
#' @importFrom EPhysData Rejected
#' @exportMethod FilterFunction<-
setMethod("FilterFunction<-", signature = "ERGExam", function(X, where, value) {
  # one channel:same filter function
  # one channel and eye:same fliter function
  # one channel and eye:same average function
  # FilterFunction set in presence of Rejected function:update rejected from lead channel
  if (any(unlist(lapply(X@Data, function(x) {
    any(Rejected(x))
  })))
  ){
    if (length(X)>1){
      Notice(X,
             what = c("W"),
             notice_text = c("! You are trynig to update the filter functions for an object that already has a Rejected function set (at least for some recordings).",
                             "i This can be problematic as it changes the rejection function that is applied after filtering. Thus it may change which trials are marked as rejected. If other recordings exist for the the same channel and eye trials rejected for those might not match anymore, resulting in potential data integrity issues for downstream analyses."),
             help_page = "ERGtools2::SetStandardFunctions")
    }
  }
  return(functionupdater(X, where, value, "FilterFunction<-"))
})

#' @describeIn UpdateProcessingMethods Update the Rejected for all Recordings in an \linkS4class{ERGExam}, or only those slected using \code{where}.
#' @exportMethod Rejected<-
setMethod("Rejected<-", signature = "ERGExam", function(X, where, value) {
  return(functionupdater(X, where, value, "Rejected<-"))
})

#' @describeIn UpdateProcessingMethods Update the AverageFunction for all Recordings in an \linkS4class{ERGExam}, or only those slected using \code{where}.
#' @exportMethod AverageFunction<-
setMethod("AverageFunction<-", signature = "ERGExam", function(X, where, value) {
  return(functionupdater(X, where, value, "AverageFunction<-"))
})

#' @describeIn UpdateProcessingMethods This method is used to set standard functions for processing \linkS4class{ERGExam} data. It defines default functions for averaging, filtering, and signal rejection based on the stimulus type.
#' @importFrom EPhysMethods filter.detrend autoreject.by.signalfree autoreject.by.distance
#' @exportMethod SetStandardFunctions
setGeneric(
  name = "SetStandardFunctions",
  def = function(X,
                 Stimulus.type.names = pairlist(Flash = "Flash",
                                           Flicker = "Flicker"),
                 Channel.hierarchy = c("VEP","ERG","OP")) {
    standardGeneric("SetStandardFunctions")
  }
)

#' @importFrom EPhysMethods filter.lin.detrend autoreject.by.signalfree autoreject.by.distance
#' @noMd
setMethod(
  "SetStandardFunctions",
  signature = "ERGExam",
  definition = function(X,
                        Stimulus.type.names = pairlist(Flash = "Flash",
                                                  Flicker = "Flicker"),
                        Channel.hierarchy = c("VEP","ERG","OP")) {
    Md <- merge(Metadata(X), StimulusTable(X))
    for (i in 1:nrow(Md)) {
      AverageFunction(X@Data[[i]]) <- mean
      FilterFunction(X@Data[[i]]) <- filter.lin.detrend
    }
    X <- rejectedUpdater(X, where= NULL, Channel.hierarchy, fx = autoreject.by.distance)
    if(validERGExam(X)){
      return(X)
    }
  }
)

#'
#' @importFrom EPhysData Metadata AverageFunction<- Rejected<- FilterFunction<-
#' @keywords internal
#' @noMd
functionupdater <- function(X, where, value, what) {
  Md <- merge(Metadata(X), StimulusTable(X))
  sel <- Where(X, where = where)
  for (i in sel) {
    tryCatch({
      if (what == "FilterFunction<-") {
        FilterFunction(X@Data[[i]]) <- value
      }
      if (what == "Rejected<-") {
        Rejected(X@Data[[i]]) <- value
      }
      if (what == "AverageFunction<-") {
        AverageFunction(X@Data[[i]]) <- value
      }
    }, error = function(e){
      Notice(object,
             what = c("E"),
             where= i,
             notice_text = c("x Running {.val {what}} on ERGExam object faild.",
                             "i Error message {.val {e}}."),
             help_page = "ERGtools2::SetStandardFunctions")
    })

  }
  if(validERGExam(X)){
    return(X)
  }
}

#' @importFrom EPhysData Metadata Rejected<-
#' @keywords internal
#' @noMd
rejectedUpdater <- function(X, where, Channel.hierarchy, fx) {
  # FIXME
  idx <- Where(X, where = where)
  Md <- merge(Metadata(X), StimulusTable(X))
  MD <- Md[idx,]
  for (s in unique(Md$Step)) {
    for (r in unique(Md$Repeat[Md$Step == s])) {
      for (e in unique(Md$Eye[Md$Step == s & Md$Repeat == r])) {
        Channels <- Md$Channel[Md$Step == s & Md$Repeat == r & Md$Eye == e]
        lead.channel <-
          Channels[which.min(as.numeric(ordered(Channels, levels = Channel.hierarchy)))]
        other.channels <- Channels[!(Channels %in% lead.channel)]
        Rejected(X@Data[[Where(X,
                               where = list(
                                 Step = s,
                                 Repeat = r,
                                 Eye = e,
                                 Channel = lead.channel
                               ))]]) <- fx
        for (c in other.channels) {
          Rejected(X@Data[[Where(X, where = list(
            Step = s,
            Repeat = r,
            Eye = e,
            Channel = c
          ))]]) <-
            Rejected(X@Data[[Where(X,
                                   where = list(
                                     Step = s,
                                     Repeat = r,
                                     Eye = e,
                                     Channel = lead.channel
                                   ))]], return.fx = F)
        }
      }
    }
  }
  return(X)
}
```
