## Supplemental Material 1 for "The ERGtools2 package: A Toolset for Processing and Analysing Visual Electrophysiology Data": StimulusTableMethods.R

```
#' Extract or Replace Parts of an ERGExam object
#'
#' Methods acting on \linkS4class{ERGExam} to extract or replace parts.
#' @inheritParams Where
#' @param full For \code{Stimulus} only. Whether to return the full stimulus table (i.e. also any additional data that might have been added by the user or when merging single \linkS4class{ERGExam} using \link{MergeERGExams}) or only the main columns "Step", "Description", "StimulusEnergy", "Background" and "Type". Default is false.
#' @param value For '<-' methods only. Vector of the same length as Stimuli selected by where containing the values to be assigned.
#' @returns For '<-' methods: an updated \linkS4class{ERGExam} object. For others: a vector containing the extracted values.
#' @examples
#' # Load the ERG dataset
#' data(ERG)
#'
#' # Extracting selected rows of the stimulus table
#' selected_rows <- Stimulus(ERG, where = list(Step = as.integer(c(1, 2, 3))))
#' head(selected_rows)
#'
#' # Extracting the description of selected stimuli
#' descriptions <- StimulusDescription(ERG, where = list(Background = "DA"))
#' descriptions
#'
#' # Extracting the intensity value of selected stimuli
#' intensity <- StimulusEnergy(ERG, where = list(Type = "Flash"))
#' intensity
#'
#' # Extracting the background value of selected stimuli
#' background <- StimulusBackground(ERG, where = list(Step = c(1:7)))
#' background
#'
#' # Extracting the type of selected stimuli
#' types <- StimulusType(ERG, where = list(Step = as.integer(c(1, 2, 3))))
#' types
#'
#' # Assigning new values to the description of selected stimuli
#' Stimulus(ERG)
#' new_descriptions <- c("New Desc 1", "New Desc 2", "New Desc 2")
#' StimulusDescription(ERG, where = list(Type = "Flash")) <- new_descriptions
#' # Now let's check if the descriptions have been updated
#' updated_descriptions <- Stimulus(ERG)
#' updated_descriptions
#'
#' @name StimulusTableMethods
NULL

#' @describeIn StimulusTableMethods Returns the stimulus table
#' @exportMethod StimulusTable
#' @noRd
setGeneric(
  name = "StimulusTable",
  def = function(X,
                 full = F)
  {
    standardGeneric("StimulusTable")
  }
)

#' @noRd
setMethod("StimulusTable",
          "ERGExam",
          function(X,
                   full = F) {
            if(!full){
              return(X@Stimulus[, c("Step", "Description", "StimulusEnergy", "Background", "Type")])
            }else{
              return(X@Stimulus)

            }
          })

#' @describeIn StimulusTableMethods Returns the stimulus names. SUPERSEEDED. Use StimulsDescription instead.
#' @keywords internal
#' @noRd
setGeneric(
  name = "StimulusNames",
  def = function(X)
  {
    standardGeneric("StimulusNames")
  }
)
#' @noRd
setMethod("StimulusNames",
          "ERGExam",
          function(X) {
            X@Stimulus$Description
          })

#' @describeIn StimulusTableMethods Returns selected rows of a stimulus table.
#' @exportMethod Stimulus
#' @noMd
setGeneric(
  name = "Stimulus",
  def = function(X,
                 where = NULL,
                 full = F)
  {
    standardGeneric("Stimulus")
  }
)

#' @noMd
setMethod("Stimulus",
          "ERGExam",
          function(X,
                   where = NULL,
                   full = F) {
            idx<-Where(X,where = where)
            step<-unique(Metadata(X)$Step[idx])
            out<-X@Stimulus[X@Stimulus$Step %in% step,]
            if(!full){
              return(out[, c("Step", "Description", "StimulusEnergy", "Background", "Type")])
            }else{
              return(out)

            }
          })


#' @describeIn StimulusTableMethods INTRNAL Gets a column of a stimulus table.
#' @keywords internal
#' @noRd
setGeneric(
  name = "StimulusX",
  def = function(X,
                 where = NULL,
                 what = NULL)
  {
    standardGeneric("StimulusX")
  }
)

#' @keywords internal
#' @noRd
setMethod("StimulusX",
          "ERGExam",
          function(X,
                   where = NULL,
                   what = NULL) {
            if(length(what)!=1 || !is.character(what)){
              stop(paste0("malformatted column selector. 'what' is '", what, "'."))
            }
            if (!(what %in% colnames(StimulusTable(X)))) {
              stop(paste0("Column '", what, "' is not a valid column of the stimuls table."))
            }
            idx<-Where(X,where = where)
            step<-unique(Metadata(X)$Step[idx])
            out<-X@Stimulus[X@Stimulus$Step %in% step, what]
            return(out)
          })

#' @describeIn StimulusTableMethods INTRNAL Sets the value a column of a selected part of the stimulus table.
#' @keywords internal
#' @noRd
setGeneric(
  name = "StimulusX<-",
  def = function(X,
                 where = NULL,
                 what = NULL,
                 value)
  {
    standardGeneric("StimulusX<-")
  }
)

#' @keywords internal
#' @noRd
setMethod("StimulusX<-",
          "ERGExam",
          function(X,
                   where = NULL,
                   what = NULL,
                   value) {
            if(length(what)!=1 || !is.character(what)){
              stop(paste0("malformatted column selector. 'what' is '", what, "'."))
            }
            if (!(what %in% colnames(StimulusTable(X)))) {
              stop(paste0("Column '", what, "' is not a valid column of the stimuls table."))
            }
            if(class(value) != class(StimulusTable(X)[,what])){
              stop(
                paste0(
                  "Incorrect class for 'value'. Is ",
                  class(value),
                  " but column '",
                  what,
                  "' in the stimulus table is of class '",
                  class(StimulusTable(X)[,what])
                  ,
                  "'."
                )
              )
            }
            idx<-Where(X,where = where)
            step<-unique(Metadata(X)$Step[idx])
            if(length(step)!=length(value)  && length(value!=1)){
              stop(paste0("Number of items selected is ", length(step), " but ",length(value), " values are given for replacement."))
            }
            X@Stimulus[X@Stimulus$Step %in% step,what]<-value
            if(validObject(X)){
              return(X)
            }
          })

#' @describeIn StimulusTableMethods Returns the description of one or more selected stimuli.
#' @exportMethod StimulusDescription
setGeneric(
  name = "StimulusDescription",
  def = function(X,
                 where = NULL)
  {
    standardGeneric("StimulusDescription")
  }
)

#' @noMd
setMethod("StimulusDescription",
          "ERGExam",
          function(X,
                   where = NULL) {
            return(StimulusX(X, where = where, what = "Description"))
          })

#' @describeIn StimulusTableMethods Sets the description of one or more selected stimuli.
#' @exportMethod StimulusDescription<-
setGeneric(
  name = "StimulusDescription<-",
  def = function(X,
                 where = NULL,
                 value)
  {
    standardGeneric("StimulusDescription<-")
  }
)

#' @noMd
setMethod("StimulusDescription<-",
          "ERGExam",
          function(X,
                   where = NULL,
                   value) {
            StimulusX(X, where = where, what = "Description")<-value
            return(X)
          })


#' @describeIn StimulusTableMethods DEPRECIATED. Use \code{StimulusEnergy} instead. Returns the energy value of one or more selected stimuli.
#' @exportMethod StimulusIntensity
setGeneric(
  name = "StimulusIntensity",
  def = function(X,
                 where = NULL)
  {
    standardGeneric("StimulusIntensity")
  }
)

#' @noMd
setMethod("StimulusIntensity",
          "ERGExam",
          function(X,
                   where = NULL) {
            return(StimulusX(X, where = where, what = "StimulusEnergy"))
          })

#' @describeIn StimulusTableMethods DEPRECIATED. Use \code{StimulusEnergy<-} instead. Sets the energy value for one or more selected stimuli.
#' @exportMethod StimulusIntensity<-
setGeneric(
  name = "StimulusIntensity<-",
  def = function(X,
                 where = NULL,
                 value)
  {
    standardGeneric("StimulusIntensity<-")
  }
)

#' @noMd
setMethod("StimulusIntensity<-",
          "ERGExam",
          function(X,
                   where = NULL,
                   value) {
            StimulusX(X, where = where, what = "StimulusEnergy")<-value
            return(X)
          })


#' @describeIn StimulusTableMethods Returns the energy value of one or more selected stimuli.
#' @exportMethod StimulusEnergy
setGeneric(
  name = "StimulusEnergy",
  def = function(X,
                 where = NULL)
  {
    standardGeneric("StimulusEnergy")
  }
)

#' @noMd
setMethod("StimulusEnergy",
          "ERGExam",
          function(X,
                   where = NULL) {
            return(StimulusX(X, where = where, what = "StimulusEnergy"))
          })

#' @describeIn StimulusTableMethods Sets the energy value for one or more selected stimuli.
#' @exportMethod StimulusEnergy<-
setGeneric(
  name = "StimulusEnergy<-",
  def = function(X,
                 where = NULL,
                 value)
  {
    standardGeneric("StimulusEnergy<-")
  }
)

#' @noMd
setMethod("StimulusEnergy<-",
          "ERGExam",
          function(X,
                   where = NULL,
                   value) {
            StimulusX(X, where = where, what = "StimulusEnergy")<-value
            return(X)
          })


#' @describeIn StimulusTableMethods Returns the background value of one or more selected stimuli.
#' @exportMethod StimulusBackground
setGeneric(
  name = "StimulusBackground",
  def = function(X,
                 where = NULL)
  {
    standardGeneric("StimulusBackground")
  }
)

#' @noMd
setMethod("StimulusBackground",
          "ERGExam",
          function(X,
                   where = NULL) {
            return(StimulusX(X, where = where, what = "Background"))
          })

#' @describeIn StimulusTableMethods Sets the background value of one or more selected stimuli.
#' @exportMethod StimulusBackground<-
setGeneric(
  name = "StimulusBackground<-",
  def = function(X,
                 where = NULL,
                 value)
  {
    standardGeneric("StimulusBackground<-")
  }
)

#' @noMd
setMethod("StimulusBackground<-",
          "ERGExam",
          function(X,
                   where = NULL,
                   value) {
            StimulusX(X, where = where, what = "Background")<-value
            return(X)
          })


#' @describeIn StimulusTableMethods Returns the type of one or more selected stimuli.
#' @exportMethod StimulusType
setGeneric(
  name = "StimulusType",
  def = function(X,
                 where = NULL)
  {
    standardGeneric("StimulusType")
  }
)

#' @noMd
setMethod("StimulusType",
          "ERGExam",
          function(X,
                   where = NULL) {
            return(StimulusX(X, where = where, what = "Type"))
          })

#' @describeIn StimulusTableMethods Sets the type value of one or more selected stimuli.
#' @exportMethod StimulusType<-
setGeneric(
  name = "StimulusType<-",
  def = function(X,
                 where = NULL,
                 value)
  {
    standardGeneric("StimulusType<-")
  }
)

#' @noMd
setMethod("StimulusType<-",
          "ERGExam",
          function(X,
                   where = NULL,
                   value) {
            StimulusX(X, where = where, what = "Type")<-value
            return(X)
          })
```
