## Supplemental Material 1 for "The ERGtools2 package: A Toolset for Processing and Analysing Visual Electrophysiology Data": stopERGtools2.R

```
#' @keywords internal
#' @noMd
setGeneric(
  name = "stopERGtools2",
  def = function(X, message)
  {
    standardGeneric("stopERGtools2")
  }
)
#' @keywords internal
#' @noMd
setMethod("stopERGtools2",
          "ERGExam",
          function(X, message) {
            return(paste0(Subject(X), ": ", message))
          })
```
