## Supplemental Material 1 for "The ERGtools2 package: A Toolset for Processing and Analysing Visual Electrophysiology Data": Subset.R

```
#' Subset from ERGExam Object
#'
#' This method subsets an \code{ERGExam} object into a new object of the same class.
#'
#' @inheritParams Get
#' @param Time Numeric vector of length 2 representing the time range for data extraction.
#'             Default is the entire time range (i.e., keep all data).
#' @param TimeExclusive Keep only the two time points stated under 'Time', not the range.
#' @param Trials Specifies which of the repeated measurements (if any) to use for extraction.
#'                It can be either a numeric vector specifying the indices of the repeated measurements
#'                or a logical vector of the same length as repeats stored,
#'                where `TRUE` indicates using that column for extraction. Default is the inverse of the \code{\link{Rejected-method}}(X) vector.
#' @param Raw Logical indicating whether to get raw data or processed (filtered, averaged) data.
#' @inheritParams Where
#' @details The \code{Subset} function creates a new \code{ERGExam}  object containing a subset of the data from the original object, based on the provided parameters.
#' @seealso \link[EPhysData:Subset]{EPhysData::Subset}
#' @importFrom EPhysData Subset newEPhysSet Metadata
#' @importFrom methods validObject as
#' @name Subset-method
#' @examples
#' data(ERG)
#' Subset(ERG,where=list(Channel="ERG"))
#' @exportMethod Subset
setMethod("Subset",
          signature(X = "ERGExam"),
          function(X,
                   Time = NULL,
                   TimeExclusive = FALSE,
                   Trials = NULL,
                   Raw = TRUE,
                   where = NULL,
                   ...) {

            where = Where(X, where)

            if (is.numeric(where)) {
              if(length(where)==0){
               stop("No recordings in the object are found for parameters provided to 'where'")
              }
              if (!all(where %in% 1:nrow(Metadata(X)))) {
                stop("Not all values in 'where' ar valid indices for data in 'X'.")
              }
              Recording.numeric <- where
              Recording <-
                (1:length(X) %in% where) # transform to logical index
            } else{
              if (length(Recording) != length(X)) {
                # length does not match
                stop(
                  "'Recording' can only be 'NULL' or a numeric or logical vector of same length as items in 'Metadata(X)'."
                )
              }
            }

            # subset measurements slot
            indexupdate <-
              as.data.frame(cbind(cumsum(Recording)[Recording], which(Recording)))
            colnames(indexupdate) <- c("new", "old")

            measurements<-X@Measurements@Measurements[X@Measurements@Measurements$Recording %in% Recording.numeric ,]

            measurements <-
              merge(measurements, indexupdate, by.x = "Recording", by.y = "old")
            measurements$Recording <- measurements$new
            measurements$new <- NULL

            X@Measurements@Measurements<-measurements
            measurements<-X@Measurements

            # subset data
            Y <- as(X, "EPhysSet")
            Y <- Subset(
              Y,
              Time = Time,
              TimeExclusive = TimeExclusive,
              Trials = Trials,
              SetItems = Recording,
              Raw = Raw,
              Simplify = FALSE
            )

            # subset stimulus
           stimtab <-
              StimulusTable(X)[StimulusTable(X)$Step %in% Metadata(Y)$Step, ]

            # subset Metadata

            metadata<-Metadata(X)[Recording,]
            rownames(metadata)<-NULL

            out<-new("ERGExam",
                Data = Y@Data,
                Metadata = metadata,
                Stimulus = stimtab,
                Averaged = X@Averaged,
                Measurements = measurements,
                ExamInfo = X@ExamInfo,
                SubjectInfo = X@SubjectInfo,
                Imported = X@Imported)

           if(validObject(out)){
             return(out)
           }
          })
```
