## Supplemental Material 1 for "The ERGtools2 package: A Toolset for Processing and Analysing Visual Electrophysiology Data": UpdateChannelNames.R

```
#' Update channel names in an ERGExam object
#'
#' This method updates channel names in an \linkS4class{ERGExam} object. It changes the channel names
#' specified in the 'from' argument to the corresponding names given in the 'to' argument for the
#' provided \linkS4class{ERGExam} object.
#'
#' @inheritParams Subset-method
#' @param from A character vector with the original channel names to be replaced.
#' @param to A character vector with the new channel names to replace the original ones.
#' @param Steps (Optional) A character vector specifying the steps to consider for the update.
#'        Defaults to all available steps in the \linkS4class{ERGExam} object.
#' @param Eyes (Optional) A character vector specifying the eyes to consider for the update.
#'        Defaults to all available eyes in the \linkS4class{ERGExam} object.
#' @return The updated \linkS4class{ERGExam} object with modified channel names.
#' @exportMethod UpdateChannelNames
#' @examples
#' \dontrun{
#' # Create an example ERGExam object
#' erg_data <- makeExampleERGSteps()
#'
#' # Original channel names
#' original_names <- c("CH1", "CH2", "CH3")
#'
#' # New channel names corresponding to the original names
#' new_names <- c("Red_Channel", "Green_Channel", "Blue_Channel")
#'
#' # Update the channel names in the ERGExam object
#' updated_erg <- UpdateChannelNames(erg_data, from = original_names, to = new_names)
#' }
setGeneric(
  name = "UpdateChannelNames",
  def = function(X, from, to, Steps = Steps(X), Eyes = Eyes(X)) {
    standardGeneric("UpdateChannelNames")
  }
)

#' @noMd
setMethod("UpdateChannelNames",
          signature(X = "ERGExam", from = "character", to = "character"),
          function(X, from, to, Steps = NULL, Eyes = NULL) {
            if (is.null(Steps)) {
              Steps <- Steps(X)
            }
            if (is.null(Eyes)) {
              Eyes <- Eyes(X)
            }
            if (!is.character(from) || !is.character(to)) {
              stop("Both 'from' and 'to' arguments must be character vectors.")
            }
            if (length(from) != length(to)) {
              stop("The lengths of 'from' and 'to' vectors must be the same.")
            }
            if (!all(from %in% Metadata(X)$Channel)) {
              warning("Some items in 'from' vector do not exist in the ERGExam object.")
            }
            if (any(duplicated(to))) {
              warning("Duplicate entries found in 'to' vector.")
              dup_to<-unique(to[duplicated(to)])
              for (i in 1:length(dup_to)){
                dup_from<-from[to==dup_to[i]]
                matching_channels <-Metadata(X)$Channel %in% dup_from
                if(length(unique(Metadata(X)$Eye[matching_channels]))<2){
                  stop("Duplicate entries results in non-unique channel names for the same eye.")
                }
              }
            }
            for (i in 1:length(from)) {
              # Temporary variables to store matching results
              matching_channels <- Metadata(X)$Channel %in% from[i]
              matching_steps <- Metadata(X)$Step %in% Steps
              matching_eyes <- Metadata(X)$Eye %in% Eyes

              # Apply the update only where all conditions are met
              X@Metadata$Channel[matching_channels & matching_steps & matching_eyes] <- to[i]
            }

            return(X)
          })
```
