## Supplemental Material 1 for "The ERGtools2 package: A Toolset for Processing and Analysing Visual Electrophysiology Data": UpdateObject.R

```
#' Update an old (v. 0.7) ERGExam object
#'
#' This method can be used to update an \linkS4class{ERGExam} object created in older versions of ERGtools2.
#' @inheritParams Subset-method
#' @return An updated  \linkS4class{ERGExam} object.
#' @name UpdateERGExam
#' @exportMethod UpdateERGExam
setGeneric(
  name = "UpdateERGExam",
  def = function(X) {
    standardGeneric("UpdateERGExam")
  }
)
#' @noMd
setMethod("UpdateERGExam",
          "ERGExam",
          function(X) {
            if ("Intensity" %in% colnames(ERG@Stimulus)) {
              Notice(X, what = "I", notice_text = "i Old Stimulus table column name 'Intensity detected. Will be updated to 'StimulusEnergy'")
              colnames(X@Stimulus)[colnames(X@Stimulus) %in% "Intensity"] <-
                "StimulusEnergy"
            }
            X
          })
```
