## Supplemental Material 1 for "The ERGtools2 package: A Toolset for Processing and Analysing Visual Electrophysiology Data": Where.R

```
#' Get index of one or several recordings and corresponding measurements
#'
#' This method gets the index of a recording stored in an \linkS4class{ERGExam} object.
#' @inheritParams Subset-method
#' @param where A \link[base:list]{base::list} defining selection criteria. Tags/Keys in the names in the list must represent valid column names of \link{Metadata} or\link{StimulusTable}.
#' @param expected.length The number of elements that is expected to match the the selection criteria. This can be a helpful validity check. Ignored if NULL.
#' @details This method gets the indices of recordings stored in an \linkS4class{ERGExam} object based on the selection citeria definde in \code{where}.
#' @return A numeric vector of  containing the index of a recording defined by the parameters, or NULL if entry was not found.
#' @examples
#' data(ERG)
#' Where(ERG,list(Channel="ERG", StimulusEnergy=1))
#'
#' @name Where
#' @exportMethod Where
setGeneric(
  name = "Where",
  def = function(X,
                 where,
                 expected.length = NULL)
  {
    standardGeneric("Where")
  }
)

#' @noMd
setMethod("Where",
          "ERGExam",
          function(X,
                   where,
                   expected.length = NULL) {
            MD <- Metadata(X)
            LENGTH.X <- length(X)
            STIMTAB <- StimulusTable(X)
            return(
              Where.generic(
                MD = MD,
                LENGTH.X = LENGTH.X,
                STIMTAB = STIMTAB,
                where = where,
                expected.length = expected.length
              )
            )
          })

#' @keywords internal
Where.generic <- function(MD,
                          LENGTH.X,
                          STIMTAB,
                          where,
                          expected.length = NULL
                          ) {

  # for backwards compatibility, convert Intensity to StimulusEnergy

  if (!is.list(where)) {
    if (is.null(where)) {
      # if all should be returned
      idx <- 1:nrow(MD)
      if (!is.null(expected.length)) {
        if (is.numeric(expected.length)) {
          if (length(idx) != expected.length) {
            stop(
              paste0(
                length(idx),
                " recordings match the selection criteria but only ",
                expected.length,
                " matches were expected."
              )
            )
          }
        } else {
          stop("'expected.length' must be NULL or numeric")
        }
      }
      return(idx)
    } else {
      if (is.numeric(where) ||
          is.integer(where)) {
        # allow pass through of Recording indices
        if (all(where %in% 1:LENGTH.X)) {
          return(where)
        } else {
          stop("Where is numeric, but not a valid recording index.")
        }
      } else {
        stop("'where' argument must be a list, or NULL for retaining all indices).")
      }
    }
  }
  if (length(where) == 0) {
    # if all should be returned
    idx <- 1:nrow(MD)
  } else {
    if ("Intensity" %in% names(where)) {
      if (!("" %in% c(colnames(STIMTAB), colnames(MD)))) {
        warning(
          "Standard column name 'Intensity' has been replaced by 'StimulusEnergy' in ERGtools2 v0.8.0. Assuming you refer to 'StimulusEnergy'..."
        )
        names(where)[names(where) == "Intensity"] <- "StimulusEnergy"
      }
    }

    if (!(all(names(where) %in% c(colnames(STIMTAB), colnames(MD))))) {
      not.found<-names(where)[!(names(where) %in% c(colnames(STIMTAB), colnames(MD)))]
      stop(
        paste0("Names of 'where' must be valid column names from the object's metadata or stimulus table, but '", not.found,"' was not found.")
      )
    }

    md.sel <- which(names(where) %in% colnames(MD))
    stim.sel <-
      which(names(where) %in% colnames(STIMTAB))


    if (all(length(md.sel) == 0, length(stim.sel) == 0)) {
      stop(
        "Names of 'where' must be valid column names from the object's metadata or stimulus table, but none of the provided names was found"
      )
    }
    if (length(stim.sel) != 0) {
      # for the entries regarding the stimulus table, convert to step
      stim <- STIMTAB
      stim.steps <- !logical(nrow(STIMTAB))
      for (n in names(where)[stim.sel]) {
        if (class(where[[n]]) != class(stim[, n])) {
          if (all( c("numeric","integer") %in% c(class(where[[n]]), class(stim[, n])))){
            if (inherits(stim[, n],"numeric")) {
              where[[n]]<-as.numeric(where[[n]])
            }
            if (inherits(stim[, n],"integer")) {
              where[[n]]<-as.integer(where[[n]])
            }
          } else {
            stop (
              paste0(
                "Entry for '",
                n,
                "' in 'where' is of type '",
                class(where[[n]]),
                "' while content of the corresponding coulumn in the stimulus table is of type '",
                class(stim[, n]),
                "'."
              )
            )
          }
          # check if has correct class

        }
        stim.steps <- stim.steps & stim[, n] %in% where[[n]]
      }
      stim.steps <- STIMTAB$Step[stim.steps]
      if ("Step" %in% names(where)) {
        stim.steps <- stim.steps[stim.steps %in% where$Step]
      } else{
        where$Step <- stim.steps
      }
      md.sel <-
        which(names(where) %in% colnames(MD)) # update md.sel
    }

    if (length(md.sel) != 0) {
      # now for the entries in metadata column
      md <- MD
      md.logidx <- !logical(nrow(MD))
      for (n in names(where)[md.sel]) {
        if (class(where[[n]]) != class(md[, n])) {
          if (all(c("numeric", "integer") %in% c(class(where[[n]]), class(md[, n])))) {
            if (inherits(md[, n], "numeric")) {
              where[[n]] <- as.numeric(where[[n]])
            }
            if (inherits(md, "integer")) {
              where[[n]] <- as.integer(where[[n]])
            }
          } else {
            # check if has correct class
            stop(
              paste0(
                "Entry for '",
                n,
                "' in 'where' is of type '",
                class(where[[n]]),
                "' while content of the corresponding coulumn in the meatadta is of type '",
                class(md[, n]),
                "'."
              )
            )
          }
        }
        md.logidx <- md.logidx & md[, n] %in% where[[n]]
      }
      idx <- which(md.logidx)
    }
  }

  if (!is.null(expected.length)) {
    if (length(idx) != expected.length) {
      stop(
        paste0(
          length(idx),
          " recordings match the selection criteria but only ",
          expected.length,
          " matches were expected."
        )
      )
    }
  }
  return(idx)
}


#' @noMd
#' @exportMethod IndexOf
setGeneric(
  name = "IndexOf",
  def = function(X,
                 Step = Steps(X),
                 Eye = Eyes(X),
                 Channel = Channels(X),
                 Repeat = 1)
  {
    standardGeneric("IndexOf")
  }
)

#' @noMd
setMethod("IndexOf",
          "ERGExam",
          function(X,
                   Step = Steps(X),
                   Eye = Eyes(X),
                   Channel = Channels(X),
                   Repeat = 1) {
            # Check if Step, Eye, and Channel are characters
            if (!is.character(Eye) || !is.character(Channel)) {
              stop("Eye and Channel must be characters.")
            }
            # Check if Repeat is numeric
            if (!is.numeric(Step) || !is.numeric(Repeat)) {
              stop("Step and Repeat must be numeric.")
            }

            if (!all(Step %in% Steps(X)) |
                !all(Eye %in% Eyes(X)) |
                !all(Channel %in% Channels(X))|
                !all(Repeat %in% Repeats(X))) {
              stop("Values for 'Step','Eye', 'Channel' and 'Repeat' must exist in the metadata.")
            }

            Md <- Metadata(X)

            which <-
              which(Md$Step  %in% Step &
                      Md$Eye %in% Eye &
                      Md$Channel %in% Channel &
                      Md$Repeat %in% Repeat)

            # if(length(which)>1){
            #   stop("Multiple recordings match the given parameters. Is the metadata slot of X corrupted?")
            # }
            return(which)
          })
```
