## Supplemental Material 1 for "The ERGtools2 package: A Toolset for Processing and Analysing Visual Electrophysiology Data": README.html

README.knit


### ERGtools2


The ERGtools2 package is an environment for working with
electroretinogram data. It contains an import method for Diagnosys
Espion data, but allows reading in of data coming in other formats with
limited coding effort. Standard procedures like averaging, sub-setting
an visualization of individual exams are supported.

#### Installation

You can install the development version of ERGtools2 from GitHub with:

```
if (!requireNamespace("remotes", quietly = TRUE)){
  install.packages("remotes")
}
remotes::install_github("moritzlindner/ERGtools2")
#> These packages have more recent versions available.
#> It is recommended to update all of them.
#> Which would you like to update?
#> 
#>  1: All                                                 
#>  2: CRAN packages only                                  
#>  3: None                                                
#>  4: EPhysData    (0.9.5        -> f88d9ccb4...) [GitHub]
#>  5: EPhysMethods (0.3.1        -> d8b7a8caa...) [GitHub]
#>  6: rlang        (1.1.2        -> 1.1.4       ) [CRAN]  
#>  7: fastmap      (1.1.1        -> 1.2.0       ) [CRAN]  
#>  8: digest       (0.6.33       -> 0.6.37      ) [CRAN]  
#>  9: htmltools    (0.5.7        -> 0.5.8.1     ) [CRAN]  
#> 10: R6           (2.5.0        -> 2.5.1       ) [CRAN]  
#> 11: fs           (1.5.0        -> 1.6.4       ) [CRAN]  
#> 12: glue         (1.6.2        -> 1.7.0       ) [CRAN]  
#> 13: cli          (3.6.2        -> 3.6.3       ) [CRAN]  
#> 14: sass         (0.4.0        -> 0.4.9       ) [CRAN]  
#> 15: mime         (0.10         -> 0.12        ) [CRAN]  
#> 16: memoise      (2.0.0        -> 2.0.1       ) [CRAN]  
#> 17: cachem       (1.0.5        -> 1.1.0       ) [CRAN]  
#> 18: Rcpp         (1.0.9        -> 1.0.13      ) [CRAN]  
#> 19: promises     (1.2.0.1      -> 1.3.0       ) [CRAN]  
#> 20: later        (1.2.0        -> 1.3.2       ) [CRAN]  
#> 21: stringi      (1.7.8        -> 1.8.4       ) [CRAN]  
#> 22: xfun         (0.34         -> 0.47        ) [CRAN]  
#> 23: tinytex      (0.31         -> 0.52        ) [CRAN]  
#> 24: fontawesome  (0.5.0        -> 0.5.2       ) [CRAN]  
#> 25: bslib        (0.3.1        -> 0.8.0       ) [CRAN]  
#> 26: highr        (0.9          -> 0.11        ) [CRAN]  
#> 27: evaluate     (0.23         -> 0.24.0      ) [CRAN]  
#> 28: yaml         (2.2.1        -> 2.3.10      ) [CRAN]  
#> 29: rmarkdown    (2.18         -> 2.28        ) [CRAN]  
#> 30: knitr        (1.33         -> 1.48        ) [CRAN]  
#> 31: sys          (3.4          -> 3.4.2       ) [CRAN]  
#> 32: askpass      (1.1          -> 1.2.0       ) [CRAN]  
#> 33: openssl      (1.4.4        -> 2.2.1       ) [CRAN]  
#> 34: curl         (4.3.1        -> 5.2.2       ) [CRAN]  
#> 35: colorspace   (2.0-1        -> 2.1-1       ) [CRAN]  
#> 36: viridisLite  (0.4.0        -> 0.4.2       ) [CRAN]  
#> 37: RColorBrewer (1.1-2        -> 1.1-3       ) [CRAN]  
#> 38: munsell      (0.5.0        -> 0.5.1       ) [CRAN]  
#> 39: labeling     (0.4.2        -> 0.4.3       ) [CRAN]  
#> 40: farver       (2.1.0        -> 2.1.2       ) [CRAN]  
#> 41: MatrixModels (0.5-0        -> 0.5-3       ) [CRAN]  
#> 42: SparseM      (1.81         -> 1.84-2      ) [CRAN]  
#> 43: utf8         (1.2.1        -> 1.2.4       ) [CRAN]  
#> 44: RcppEigen    (0.3.3.9.1    -> 0.3.4.0.2   ) [CRAN]  
#> 45: nloptr       (1.2.2.2      -> 2.1.1       ) [CRAN]  
#> 46: minqa        (1.2.4        -> 1.2.8       ) [CRAN]  
#> 47: withr        (2.5.2        -> 3.0.1       ) [CRAN]  
#> 48: cpp11        (0.2.7        -> 0.5.0       ) [CRAN]  
#> 49: fansi        (0.5.0        -> 1.0.6       ) [CRAN]  
#> 50: pillar       (1.6.1        -> 1.9.0       ) [CRAN]  
#> 51: lme4         (1.1-27       -> 1.1-35.5    ) [CRAN]  
#> 52: quantreg     (5.85         -> 5.98        ) [CRAN]  
#> 53: carData      (3.0-4        -> 3.0-5       ) [CRAN]  
#> 54: backports    (1.2.1        -> 1.5.0       ) [CRAN]  
#> 55: car          (3.0-10       -> 3.1-2       ) [CRAN]  
#> 56: tidyselect   (1.1.1        -> 1.2.1       ) [CRAN]  
#> 57: corrplot     (0.88         -> 0.94        ) [CRAN]  
#> 58: dplyr        (1.0.6        -> 1.1.4       ) [CRAN]  
#> 59: tibble       (3.1.2        -> 3.2.1       ) [CRAN]  
#> 60: broom        (0.7.6        -> 1.0.6       ) [CRAN]  
#> 61: tidyr        (1.1.3        -> 1.3.1       ) [CRAN]  
#> 62: gtable       (0.3.0        -> 0.3.5       ) [CRAN]  
#> 63: isoband      (0.2.4        -> 0.2.7       ) [CRAN]  
#> 64: ggplot2      (3.4.4        -> 3.5.1       ) [CRAN]  
#> 65: crosstalk    (1.1.1        -> 1.2.1       ) [CRAN]  
#> 66: httpuv       (1.6.1        -> 1.6.15      ) [CRAN]  
#> 67: htmlwidgets  (1.5.3        -> 1.6.4       ) [CRAN]  
#> 68: commonmark   (1.7          -> 1.9.1       ) [CRAN]  
#> 69: crayon       (1.4.1        -> 1.5.3       ) [CRAN]  
#> 70: sourcetools  (0.1.7        -> 0.1.7-1     ) [CRAN]  
#> 71: data.table   (1.14.0       -> 1.16.0      ) [CRAN]  
#> 72: httr         (1.4.2        -> 1.4.7       ) [CRAN]  
#> 73: rstatix      (0.7.0        -> 0.7.2       ) [CRAN]  
#> 74: polynom      (1.4-0        -> 1.4-1       ) [CRAN]  
#> 75: ggsignif     (0.6.1        -> 0.6.4       ) [CRAN]  
#> 76: cowplot      (1.1.1        -> 1.1.3       ) [CRAN]  
#> 77: ggsci        (2.9          -> 3.2.0       ) [CRAN]  
#> 78: ggrepel      (0.9.1        -> 0.9.5       ) [CRAN]  
#> 79: DT           (0.19         -> 0.33        ) [CRAN]  
#> 80: shiny        (1.8.0        -> 1.9.1       ) [CRAN]  
#> 81: units        (0.8-0        -> 0.8-5       ) [CRAN]  
#> 82: remotes      (2.4.2        -> 2.5.0       ) [CRAN]  
#> 83: plotly       (16261c331... -> 3cf17c004...) [GitHub]
#> 84: ggpubr       (0.4.0        -> 0.6.0       ) [CRAN]  
#> 
#>   
   checking for file ‘/tmp/Rtmp3jyDG9/remotes18147fd45ae/moritzlindner-ERGtools2-f1617f5/DESCRIPTION’ ...
  
✔  checking for file ‘/tmp/Rtmp3jyDG9/remotes18147fd45ae/moritzlindner-ERGtools2-f1617f5/DESCRIPTION’
#> 
  
─  preparing ‘ERGtools2’:
#>    checking DESCRIPTION meta-information ...
  
✔  checking DESCRIPTION meta-information
#> 
  
─  excluding invalid files
#> 
  
   Subdirectory 'R' contains invalid file names:
#>      ‘ERGtools2-package.html’
#> ─  checking for LF line-endings in source and make files and shell scripts
#> ─  checking for empty or unneeded directories
#> 
  
─  building ‘ERGtools2_0.7.0.tar.gz’
#> 
  
   
#>
```

Note that `ERGtools2` depends on the github-deposited R
Packages `EPhysData` and `EPhysMethods`.
Installation usually works automatically. Updating, however may fail. If
this is the case, update manually using the following line of code:

```
remotes::install_github("moritzlindner/EPhysData")
#> These packages have more recent versions available.
#> It is recommended to update all of them.
#> Which would you like to update?
#> 
#>  1: All                                                 
#>  2: CRAN packages only                                  
#>  3: None                                                
#>  4: withr        (2.5.2        -> 3.0.1       ) [CRAN]  
#>  5: rlang        (1.1.2        -> 1.1.4       ) [CRAN]  
#>  6: glue         (1.6.2        -> 1.7.0       ) [CRAN]  
#>  7: cli          (3.6.2        -> 3.6.3       ) [CRAN]  
#>  8: stringi      (1.7.8        -> 1.8.4       ) [CRAN]  
#>  9: bit          (4.0.4        -> 4.0.5       ) [CRAN]  
#> 10: colorspace   (2.0-1        -> 2.1-1       ) [CRAN]  
#> 11: viridisLite  (0.4.0        -> 0.4.2       ) [CRAN]  
#> 12: RColorBrewer (1.1-2        -> 1.1-3       ) [CRAN]  
#> 13: munsell      (0.5.0        -> 0.5.1       ) [CRAN]  
#> 14: labeling     (0.4.2        -> 0.4.3       ) [CRAN]  
#> 15: farver       (2.1.0        -> 2.1.2       ) [CRAN]  
#> 16: MatrixModels (0.5-0        -> 0.5-3       ) [CRAN]  
#> 17: SparseM      (1.81         -> 1.84-2      ) [CRAN]  
#> 18: utf8         (1.2.1        -> 1.2.4       ) [CRAN]  
#> 19: RcppEigen    (0.3.3.9.1    -> 0.3.4.0.2   ) [CRAN]  
#> 20: Rcpp         (1.0.9        -> 1.0.13      ) [CRAN]  
#> 21: nloptr       (1.2.2.2      -> 2.1.1       ) [CRAN]  
#> 22: minqa        (1.2.4        -> 1.2.8       ) [CRAN]  
#> 23: cpp11        (0.2.7        -> 0.5.0       ) [CRAN]  
#> 24: fansi        (0.5.0        -> 1.0.6       ) [CRAN]  
#> 25: R6           (2.5.0        -> 2.5.1       ) [CRAN]  
#> 26: pillar       (1.6.1        -> 1.9.0       ) [CRAN]  
#> 27: lme4         (1.1-27       -> 1.1-35.5    ) [CRAN]  
#> 28: quantreg     (5.85         -> 5.98        ) [CRAN]  
#> 29: carData      (3.0-4        -> 3.0-5       ) [CRAN]  
#> 30: backports    (1.2.1        -> 1.5.0       ) [CRAN]  
#> 31: car          (3.0-10       -> 3.1-2       ) [CRAN]  
#> 32: tidyselect   (1.1.1        -> 1.2.1       ) [CRAN]  
#> 33: corrplot     (0.88         -> 0.94        ) [CRAN]  
#> 34: dplyr        (1.0.6        -> 1.1.4       ) [CRAN]  
#> 35: tibble       (3.1.2        -> 3.2.1       ) [CRAN]  
#> 36: broom        (0.7.6        -> 1.0.6       ) [CRAN]  
#> 37: tidyr        (1.1.3        -> 1.3.1       ) [CRAN]  
#> 38: gtable       (0.3.0        -> 0.3.5       ) [CRAN]  
#> 39: isoband      (0.2.4        -> 0.2.7       ) [CRAN]  
#> 40: ggplot2      (3.4.4        -> 3.5.1       ) [CRAN]  
#> 41: rstatix      (0.7.0        -> 0.7.2       ) [CRAN]  
#> 42: polynom      (1.4-0        -> 1.4-1       ) [CRAN]  
#> 43: ggsignif     (0.6.1        -> 0.6.4       ) [CRAN]  
#> 44: cowplot      (1.1.1        -> 1.1.3       ) [CRAN]  
#> 45: ggsci        (2.9          -> 3.2.0       ) [CRAN]  
#> 46: ggrepel      (0.9.1        -> 0.9.5       ) [CRAN]  
#> 47: remotes      (2.4.2        -> 2.5.0       ) [CRAN]  
#> 48: hdf5r        (097caa53f... -> dc4774c03...) [GitHub]
#> 49: ggpubr       (0.4.0        -> 0.6.0       ) [CRAN]  
#> 50: units        (0.8-0        -> 0.8-5       ) [CRAN]  
#> 
#>   
   checking for file ‘/tmp/Rtmp3jyDG9/remotes1814669659b7/moritzlindner-EPhysData-f88d9cc/DESCRIPTION’ ...
  
✔  checking for file ‘/tmp/Rtmp3jyDG9/remotes1814669659b7/moritzlindner-EPhysData-f88d9cc/DESCRIPTION’
#> 
  
─  preparing ‘EPhysData’:
#>    checking DESCRIPTION meta-information ...
  
✔  checking DESCRIPTION meta-information
#> 
  
─  checking for LF line-endings in source and make files and shell scripts
#> 
  
─  checking for empty or unneeded directories
#>    Omitted ‘LazyData’ from DESCRIPTION
#> ─  building ‘EPhysData_0.9.5.tar.gz’
#> 
  
   
#> 
remotes::install_github("moritzlindner/EPhysMethods")
#> These packages have more recent versions available.
#> It is recommended to update all of them.
#> Which would you like to update?
#> 
#> 1: All                            
#> 2: CRAN packages only             
#> 3: None                           
#> 4: Rcpp   (1.0.9 -> 1.0.13) [CRAN]
#> 5: units  (0.8-0 -> 0.8-5 ) [CRAN]
#> 6: signal (0.7-7 -> 1.8-1 ) [CRAN]
#> 7: pracma (2.3.3 -> 2.4.4 ) [CRAN]
#> 
#>   
   checking for file ‘/tmp/Rtmp3jyDG9/remotes18147d7e0f4/moritzlindner-EPhysMethods-d8b7a8c/DESCRIPTION’ ...
  
✔  checking for file ‘/tmp/Rtmp3jyDG9/remotes18147d7e0f4/moritzlindner-EPhysMethods-d8b7a8c/DESCRIPTION’
#> 
  
─  preparing ‘EPhysMethods’:
#>    checking DESCRIPTION meta-information ...
  
✔  checking DESCRIPTION meta-information
#> 
  
─  checking for LF line-endings in source and make files and shell scripts
#> 
  
─  checking for empty or unneeded directories
#>    Omitted ‘LazyData’ from DESCRIPTION
#> 
  
─  building ‘EPhysMethods_0.3.1.tar.gz’
#> 
  
   
#>
```

#### Example

This is a basic example which shows you how to solve a common
problem:

```
library(ERGtools2)

## a typical workflow
data(ERG) # load example data, to import own data, see the examples provided for newERGExam and ImportEspion
StimulusTable(ERG) # have a look whats inside
#>   Step     Description Intensity Background  Type
#> 1    1 DA 0 01 cd s m       0.01         DA Flash
#> 2    2    DA 1 cd s m       1.00         DA Flash
#> 3    3    DA 3 cd s m       3.00         DA Flash
Metadata(ERG)
#>    Step Channel Repeat Eye Channel_Name Recording
#> 1     1     ERG      1  RE          ERG         1
#> 2     1     ERG      1  LE          ERG         2
#> 3     1      OP      1  RE           OP         3
#> 4     1      OP      1  LE           OP         4
#> 5     2     ERG      1  RE          ERG         5
#> 6     2     ERG      1  LE          ERG         6
#> 7     2      OP      1  RE           OP         7
#> 8     2      OP      1  LE           OP         8
#> 9     3     ERG      1  RE          ERG         9
#> 10    3     ERG      1  LE          ERG        10
#> 11    3      OP      1  RE           OP        11
#> 12    3      OP      1  LE           OP        12
ERG<-ClearMeasurements(ERG) # Clear Measuremnts that migth be alredy in object
ERG<-SetStandardFunctions(ERG)
ggERGTrace(ERG, where = list( Step = as.integer(3), Eye = "RE", Channel ="ERG", Repeat = as.integer(1))) # pick one and have a look at the traces as imported
```

```
ERG <- AutoPlaceMarkers(ERG, Channel.names = pairlist(ERG = "ERG")) # automatically place markers
#> ========================================================================================================================================================================================================================
Measurements(ERG)
#> ========================================================================================================================================================================================================================
#>    Recording Step     Description Channel Repeat Eye Name Relative       Time         Voltage Channel_Name Recording.y
#> 1          1    1 DA 0 01 cd s m      ERG      1  RE    a     <NA> 0.0315 [s]  -53.17071 [uV]          ERG           1
#> 2          1    1 DA 0 01 cd s m      ERG      1  RE    B        a 0.0515 [s]  346.34079 [uV]          ERG           1
#> 3          2    1 DA 0 01 cd s m      ERG      1  LE    a     <NA> 0.0310 [s]  -85.88321 [uV]          ERG           2
#> 4          2    1 DA 0 01 cd s m      ERG      1  LE    B        a 0.0505 [s]  310.38093 [uV]          ERG           2
#> 5          5    2    DA 1 cd s m      ERG      1  RE    a     <NA> 0.0125 [s] -206.37706 [uV]          ERG           5
#> 6          5    2    DA 1 cd s m      ERG      1  RE    B        a 0.0355 [s]  547.86889 [uV]          ERG           5
#> 7          6    2    DA 1 cd s m      ERG      1  LE    a     <NA> 0.0130 [s] -209.66536 [uV]          ERG           6
#> 8          6    2    DA 1 cd s m      ERG      1  LE    B        a 0.0355 [s]  491.97428 [uV]          ERG           6
#> 9          9    3    DA 3 cd s m      ERG      1  RE    a     <NA> 0.0090 [s] -233.89249 [uV]          ERG           9
#> 10         9    3    DA 3 cd s m      ERG      1  RE    B        a 0.0345 [s]  587.60032 [uV]          ERG           9
#> 11        10    3    DA 3 cd s m      ERG      1  LE    a     <NA> 0.0095 [s] -240.61577 [uV]          ERG          10
#> 12        10    3    DA 3 cd s m      ERG      1  LE    B        a 0.0345 [s]  568.70641 [uV]          ERG          10
ggERGTrace(ERG, where = list( Step = as.integer(3), Eye = "RE", Channel ="ERG", Repeat = as.integer(1))) # pick one and have a look at the traces as imported
#> ========================================================================================================================================================================================================================
#> ========================================================================================================================================================================================================================
```
