## Supplemental Material 1 for "The ERGtools2 package: A Toolset for Processing and Analysing Visual Electrophysiology Data": README.Rmd

---
output: github_document
---

<!-- README.md is generated from README.Rmd. Please edit that file -->

```{r, include = FALSE}
knitr::opts_chunk$set(
  collapse = TRUE,
  comment = "#>",
  fig.path = "man/figures/README-",
  out.width = "100%",
  message = FALSE
)
```

# ERGtools2

<!-- badges: start -->
<!-- badges: end -->

The ERGtools2 package is an environment for working with electroretinogram data. It contains an import method for Diagnosys Espion data, but allows reading in of data coming in other formats with limited coding effort. Standard procedures like averaging, sub-setting an visualization of individual exams are supported.


## Installation

You can install the development version of ERGtools2 from [GitHub](https://github.com/) with:

``` {r installation}
if (!requireNamespace("remotes", quietly = TRUE)){
  install.packages("remotes")
}
remotes::install_github("moritzlindner/ERGtools2")
```

Note that  `ERGtools2` depends on the github-deposited R Packages  `EPhysData` and `EPhysMethods`. Installation usually works automatically. Updating, however may fail. If this is the case, update manually using the following line of code:

``` {r Update EPhysData and EPhysMethods}
remotes::install_github("moritzlindner/EPhysData")
remotes::install_github("moritzlindner/EPhysMethods")
```

## Example

This is a basic example which shows you how to solve a common problem:

```{r example}
library(ERGtools2)

## a typical workflow
data(ERG) # load example data, to import own data, see the examples provided for newERGExam and ImportEspion
StimulusTable(ERG) # have a look whats inside
Metadata(ERG)
ERG<-ClearMeasurements(ERG) # Clear Measuremnts that migth be alredy in object
ERG<-SetStandardFunctions(ERG)
ggERGTrace(ERG, where = list( Step = as.integer(3), Eye = "RE", Channel ="ERG", Repeat = as.integer(1))) # pick one and have a look at the traces as imported

ERG <- AutoPlaceMarkers(ERG, Channel.names = pairlist(ERG = "ERG")) # automatically place markers
Measurements(ERG)
ggERGTrace(ERG, where = list( Step = as.integer(3), Eye = "RE", Channel ="ERG", Repeat = as.integer(1))) # pick one and have a look at the traces as imported

```
