## Supplemental Material 1 for "The ERGtools2 package: A Toolset for Processing and Analysing Visual Electrophysiology Data": Rplots.pdf

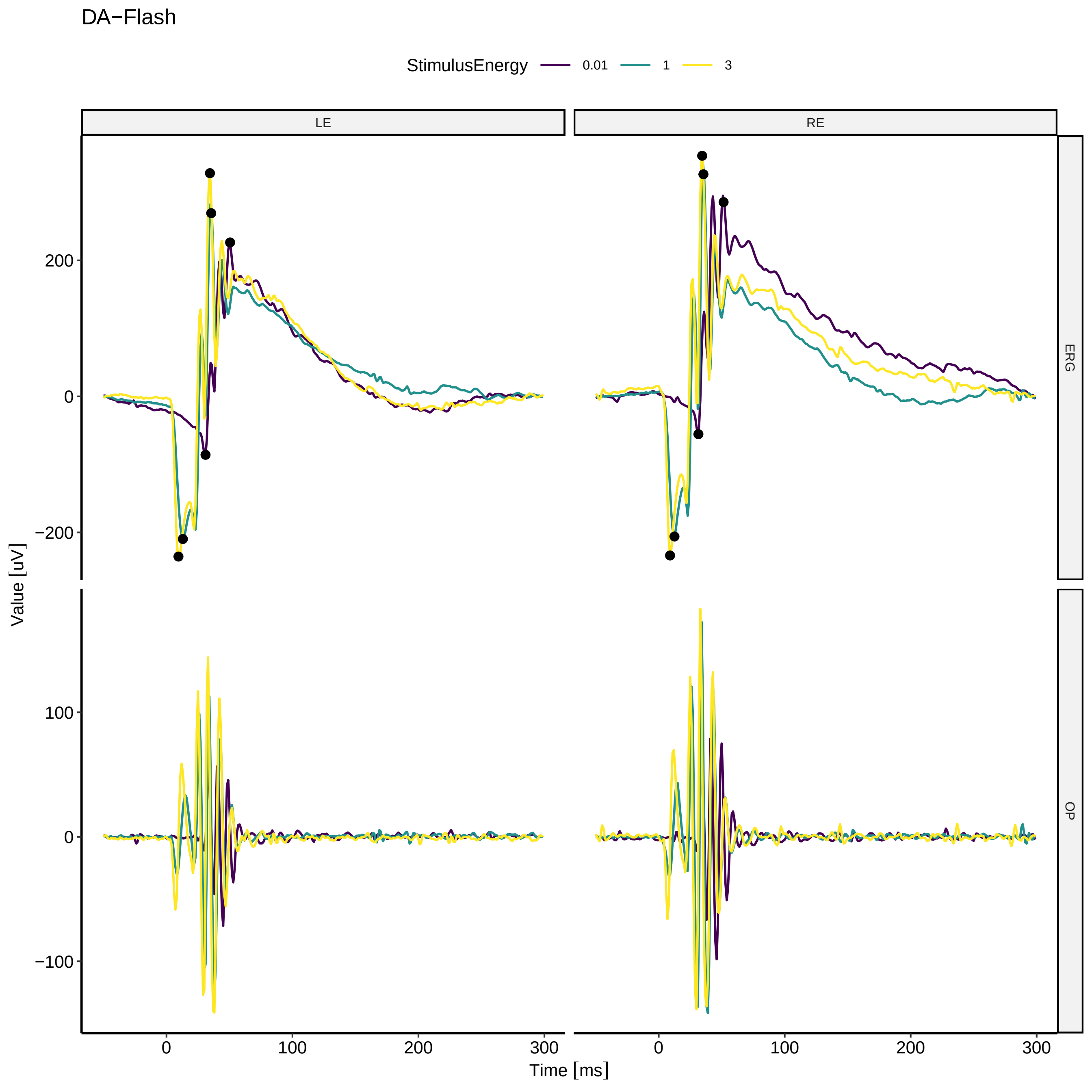
DA−Flash
StimulusEnergy
0.01
1
3
LE
RE
ERG
OP
200
0
−200
100
0
−100
Value [uV]
0
100
200
300
0
100
200
300
Time [ms]

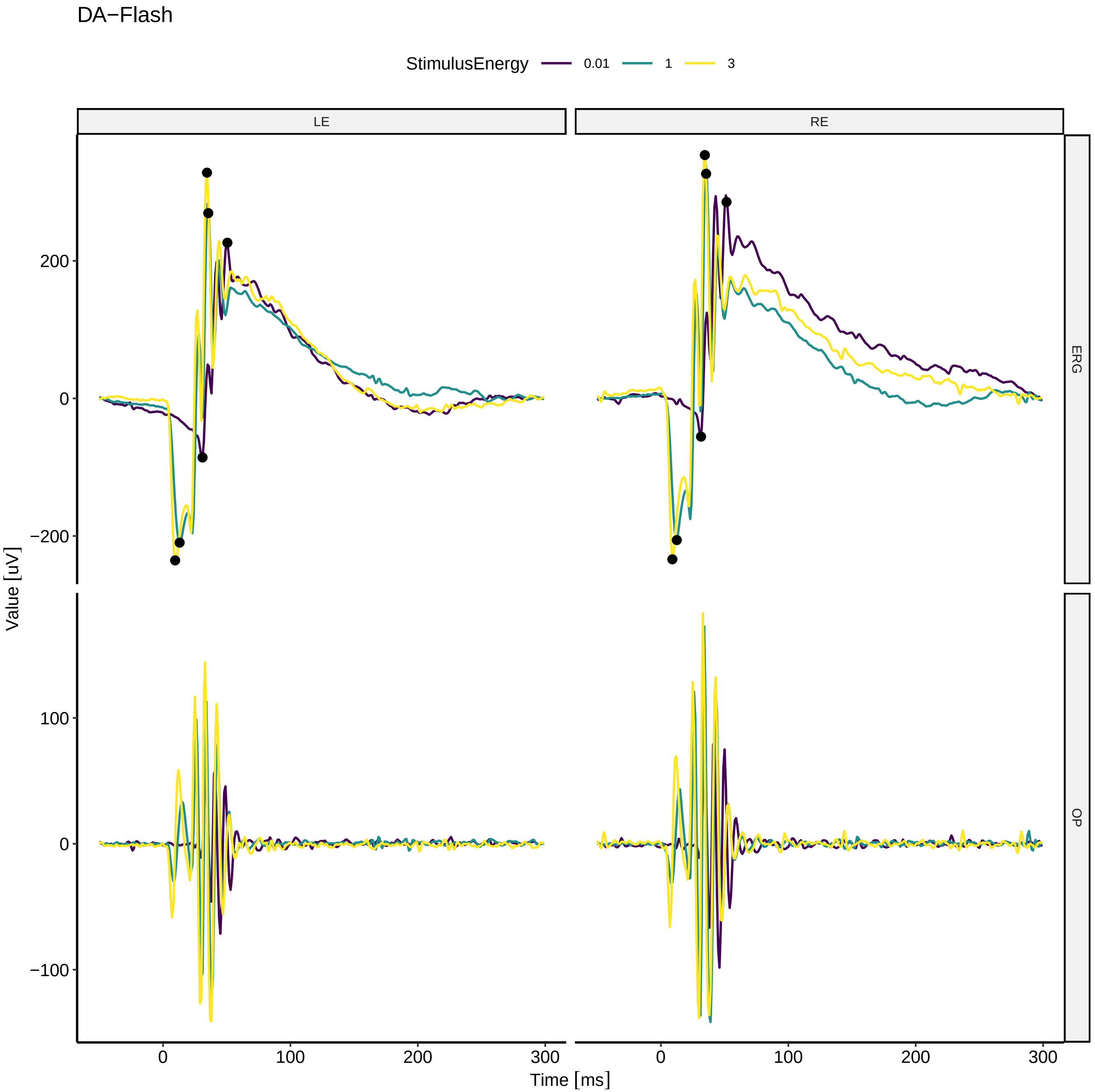
DA−Flash
StimulusEnergy
0.01
1
3
LE
RE
ERG
OP
Value [uV]
Time [ms]
200
0
−200
100
0
−100
0
100
200
300
0
100
200
300

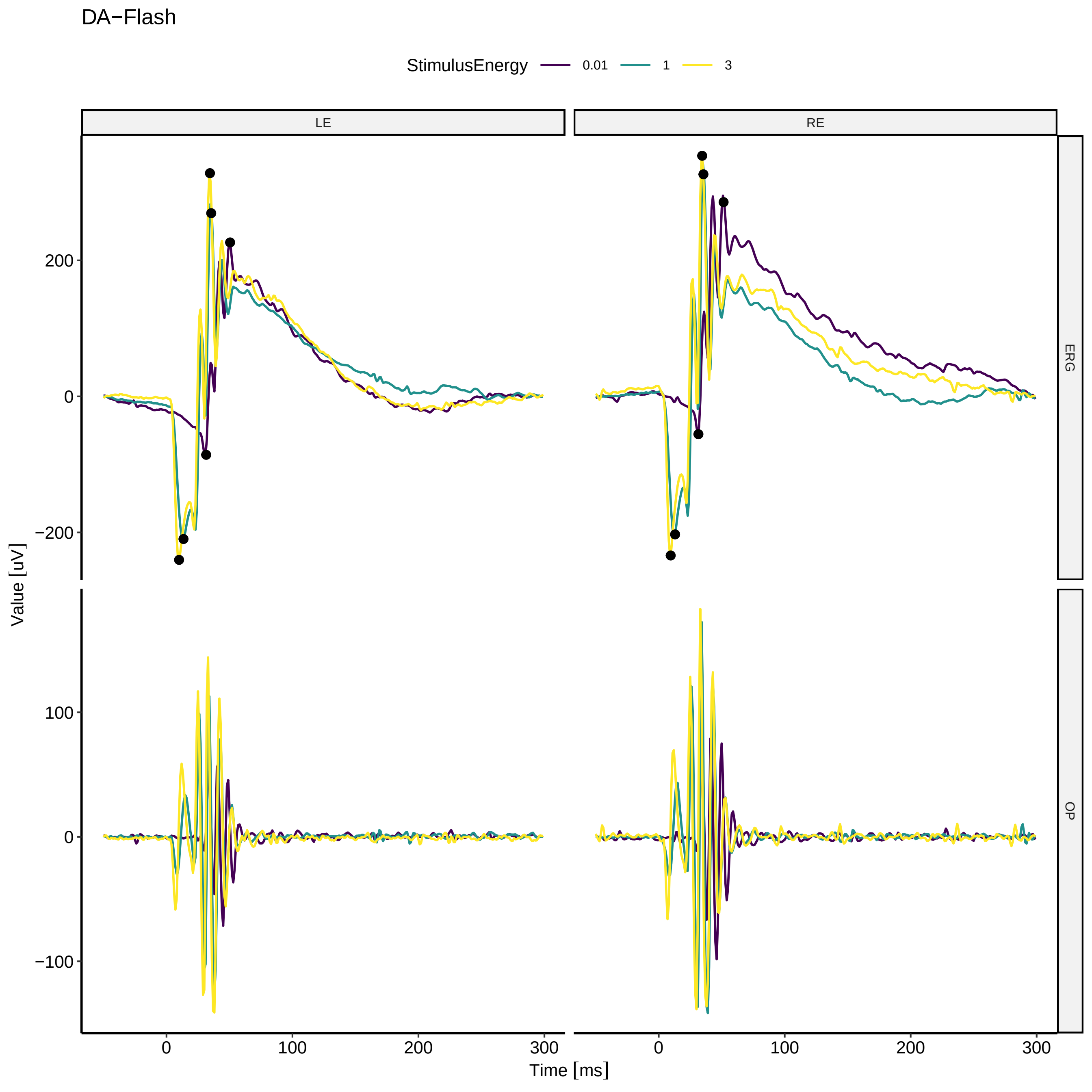

DA−Flash
StimulusEnergy
0.01
1
3
LE
RE
ERG
OP
Value [uV]
Time [ms]
200
0
−200
100
0
−100
0
100
200
300
0
100
200
300

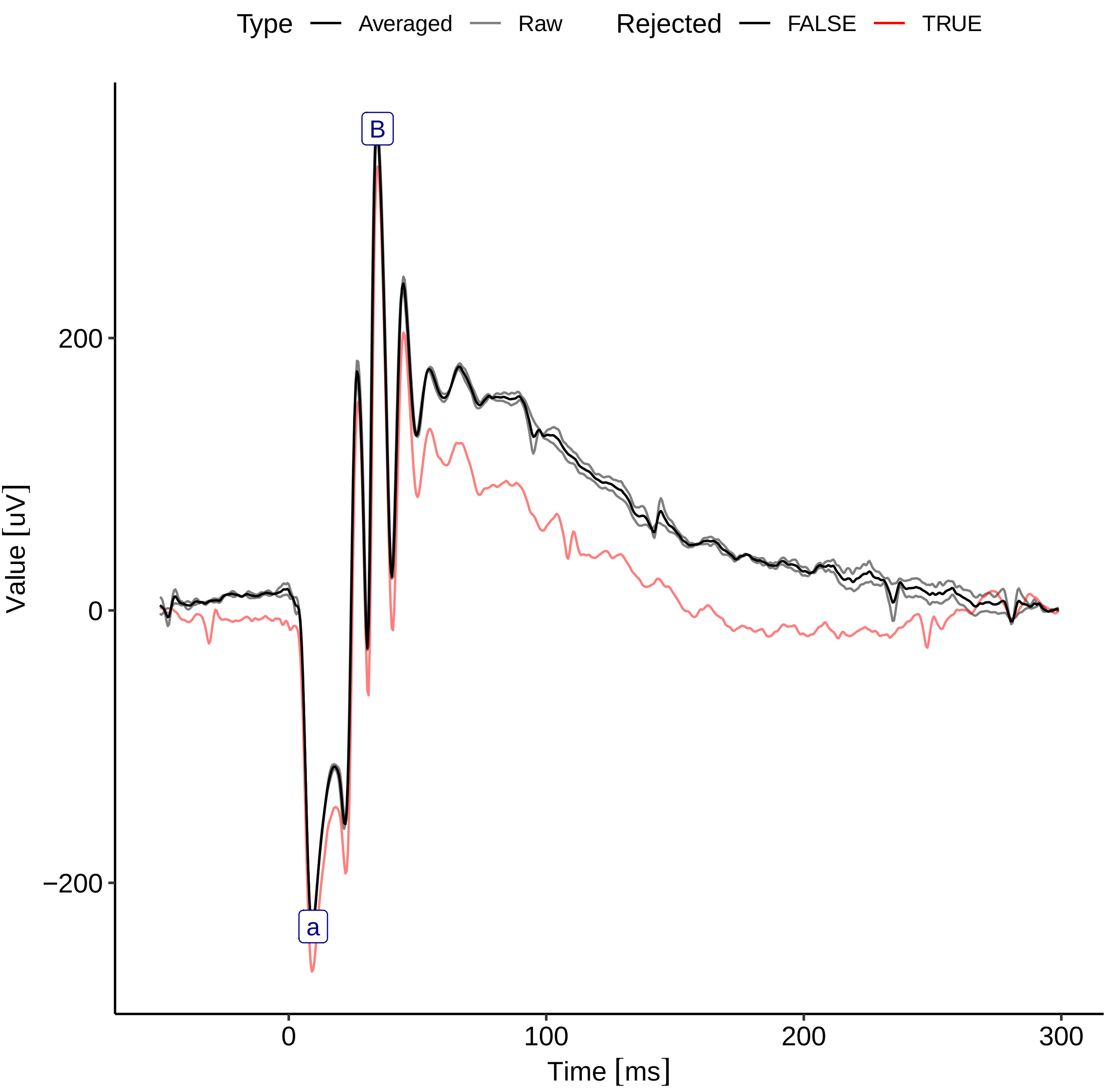

Type
Averaged
Raw
Rejected
FALSE
TRUE
B
a
200
0
−200
Value [uV]
0
100
200
300
Time [ms]

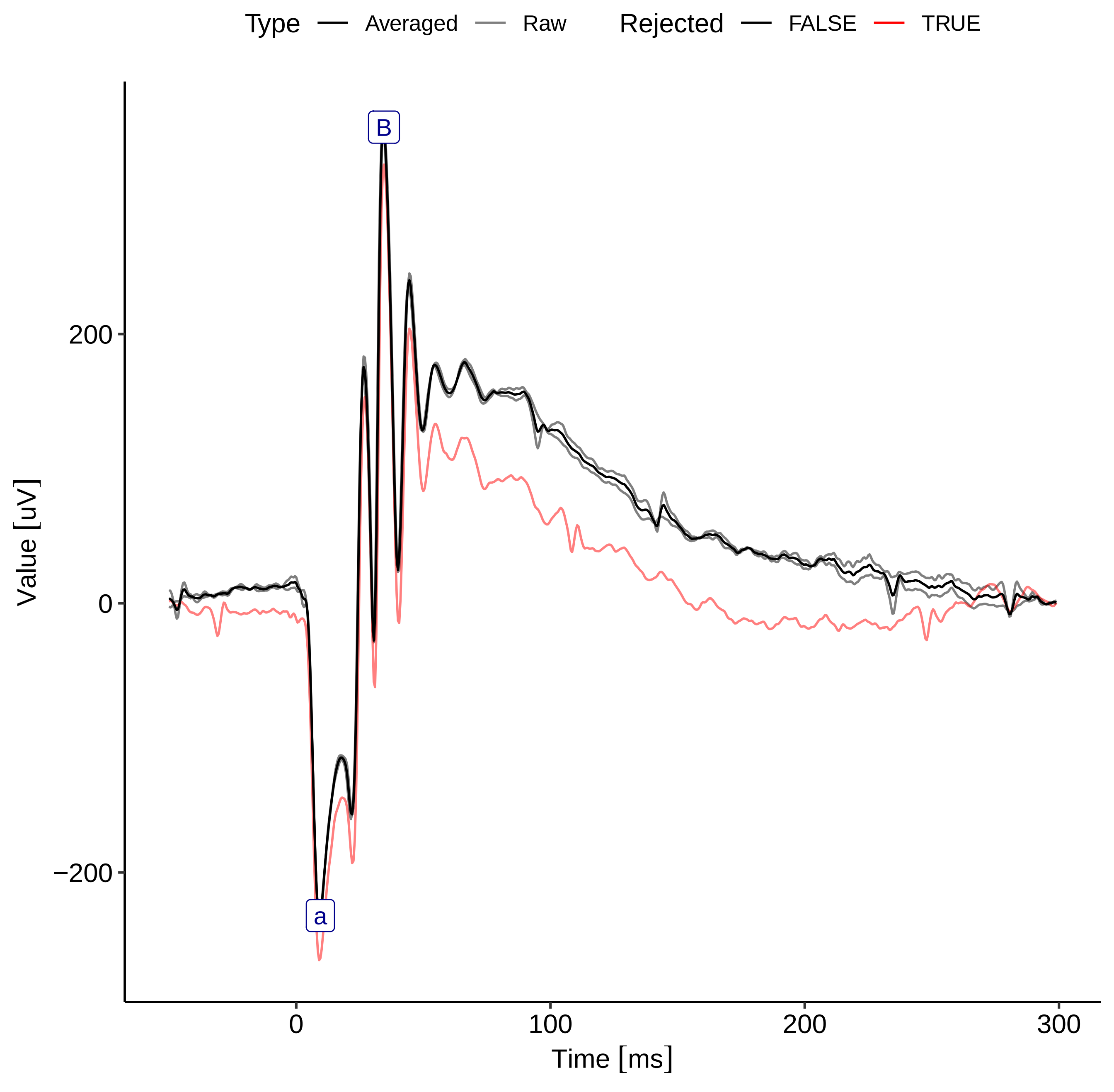

Type
Averaged
Raw
Rejected
FALSE
TRUE
B
a
200
0
−200
0
100
200
300
Value [uV]
Time [ms]

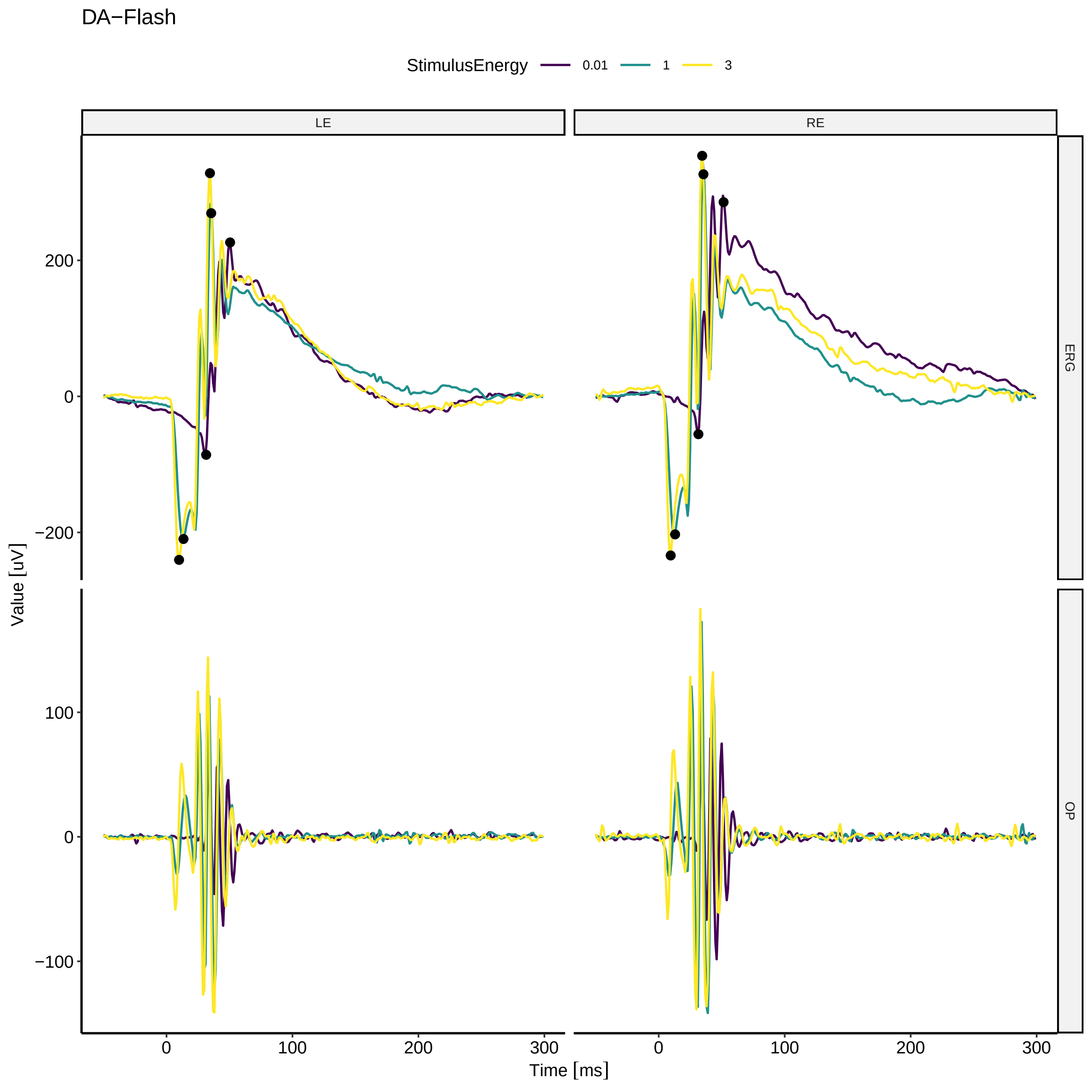
DA−Flash
StimulusEnergy
0.01
1
3
LE
RE
ERG
OP
Value [uV]
Time [ms]
200
0
−200
100
−100
300

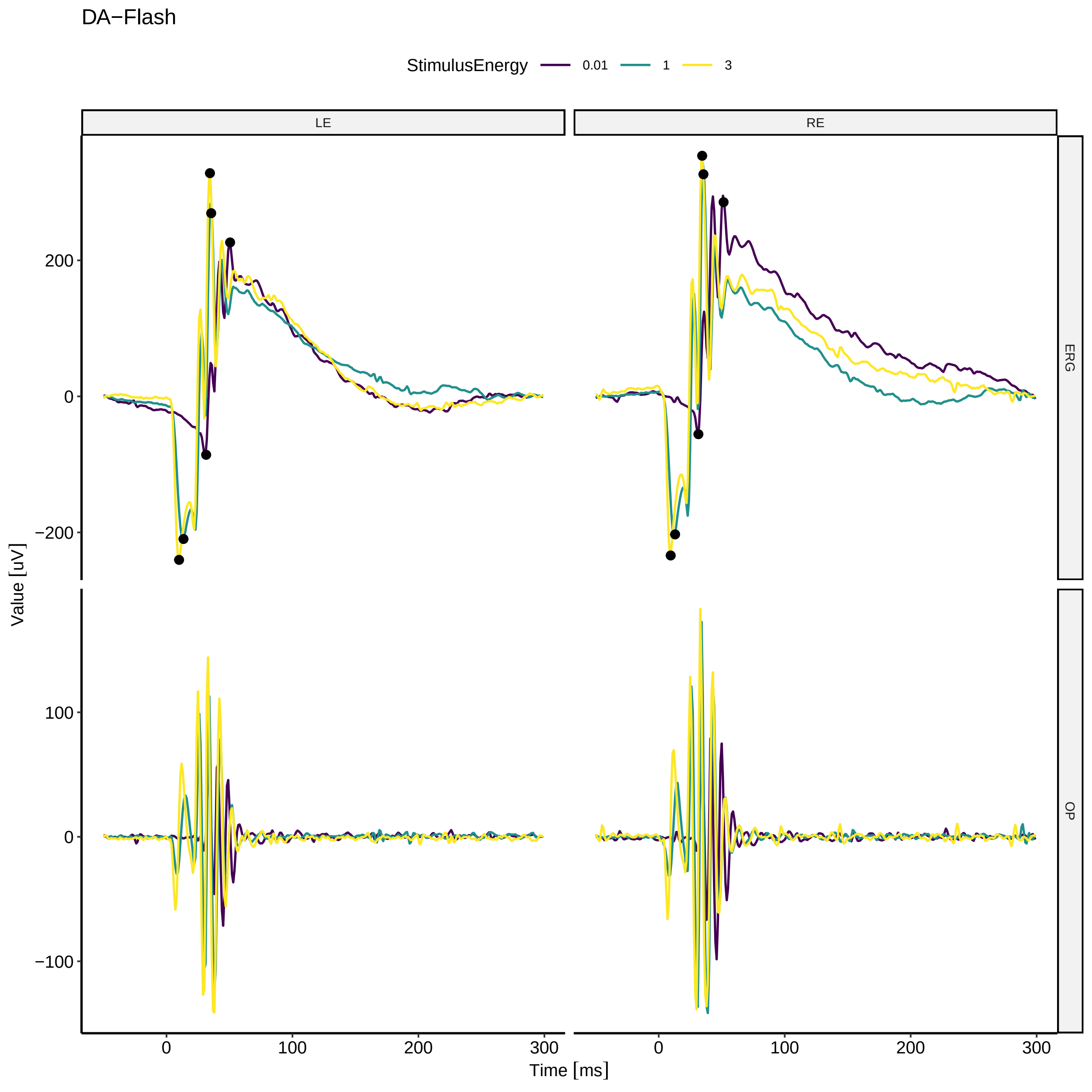

DA−Flash
StimulusEnergy
0.01
1
3
LE
RE
ERG
OP
Value [uV]
Time [ms]
200
0
−200
100
−100
300

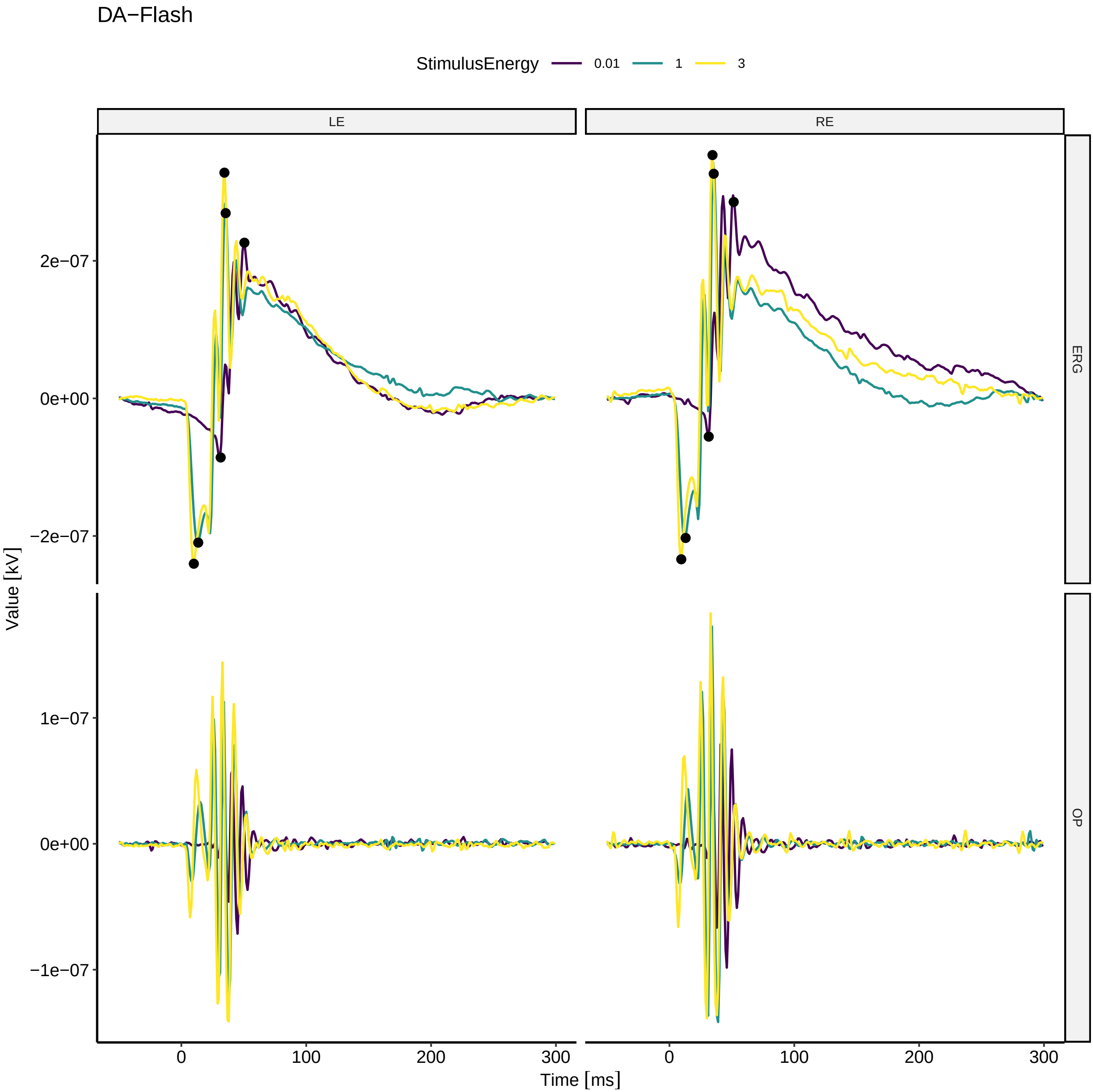
DA−Flash
StimulusEnergy
0.01
1
3
LE
RE
ERG
OP
2e−07
0e+00
−2e−07
1e−07
−1e−07
Value [kV]
0
100
200
300
Time [ms]

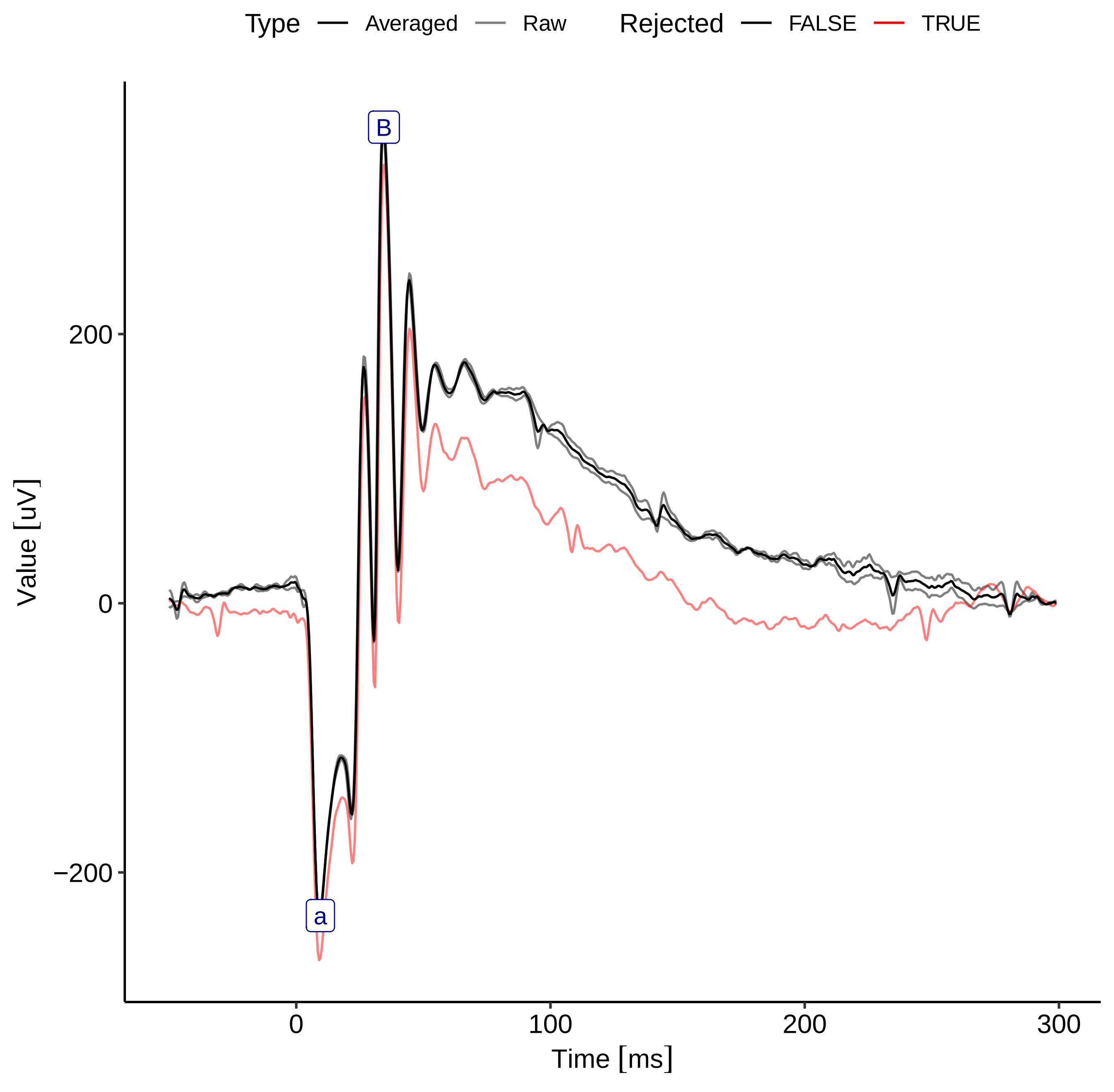

Type
Averaged
Raw
Rejected
FALSE
TRUE
B
a
200
0
−200
0
100
200
300
Value [uV]
Time [ms]

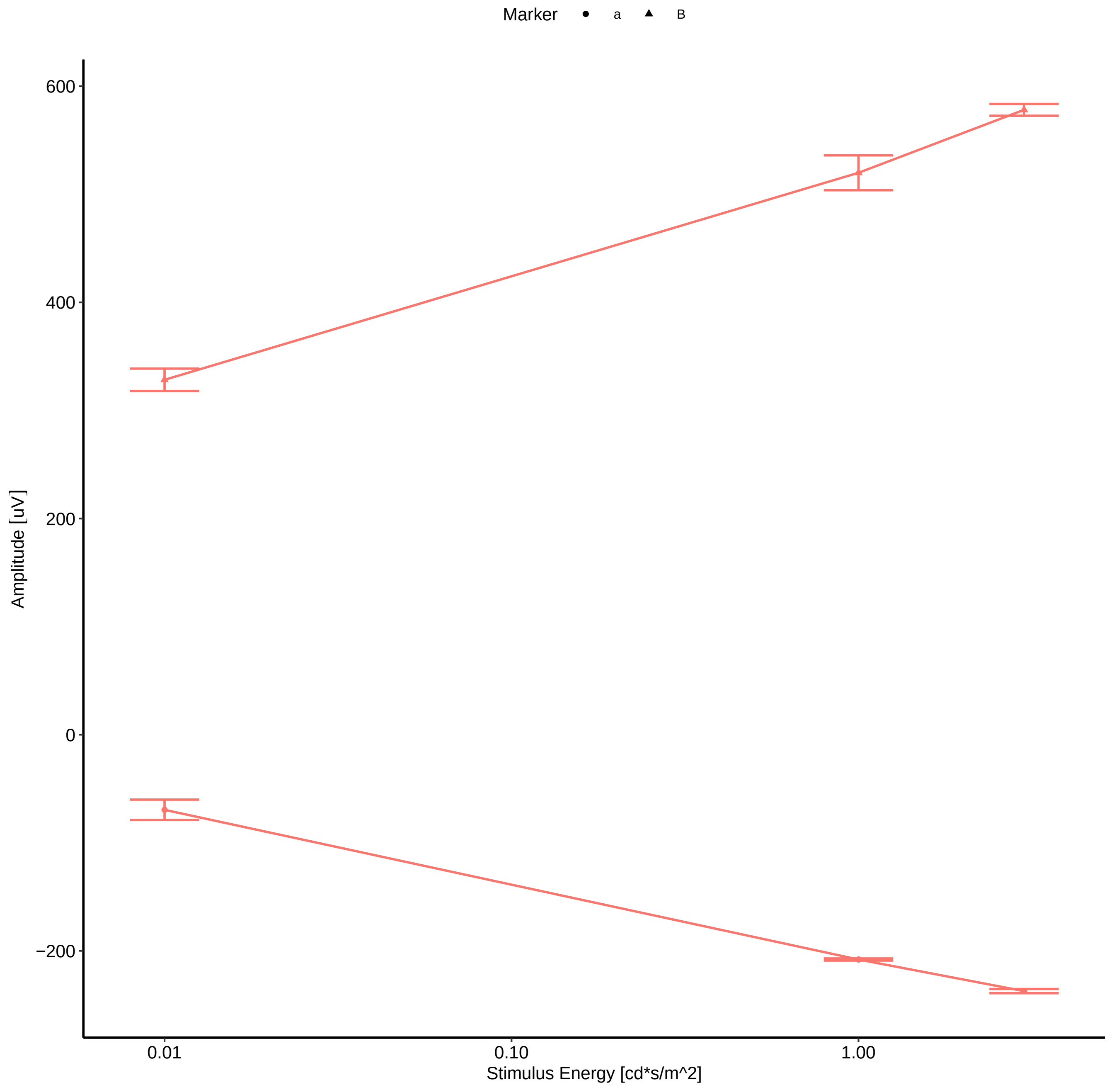
Marker
a
B
600
400
200
0
−200
Amplitude [uV]
0.01
0.10
1.00
Stimulus Energy [cd*s/m^2]

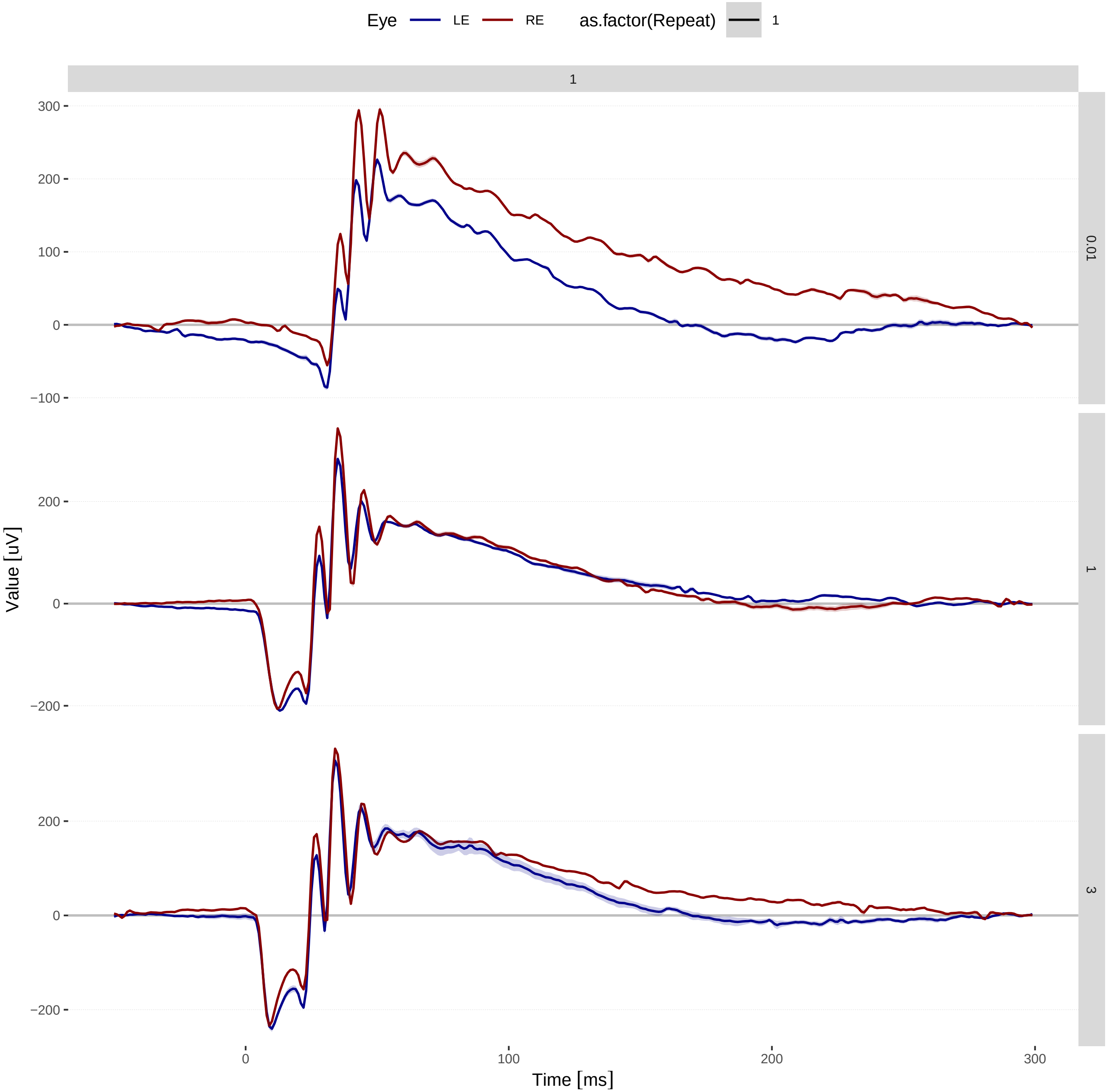

Eye
LE
RE
as.factor(Repeat)
1
1
0.01
1
3
300
200
100
0
−100
200
0
−200
200
0
−200
Value [uV]
0
100
200
300
Time [ms]

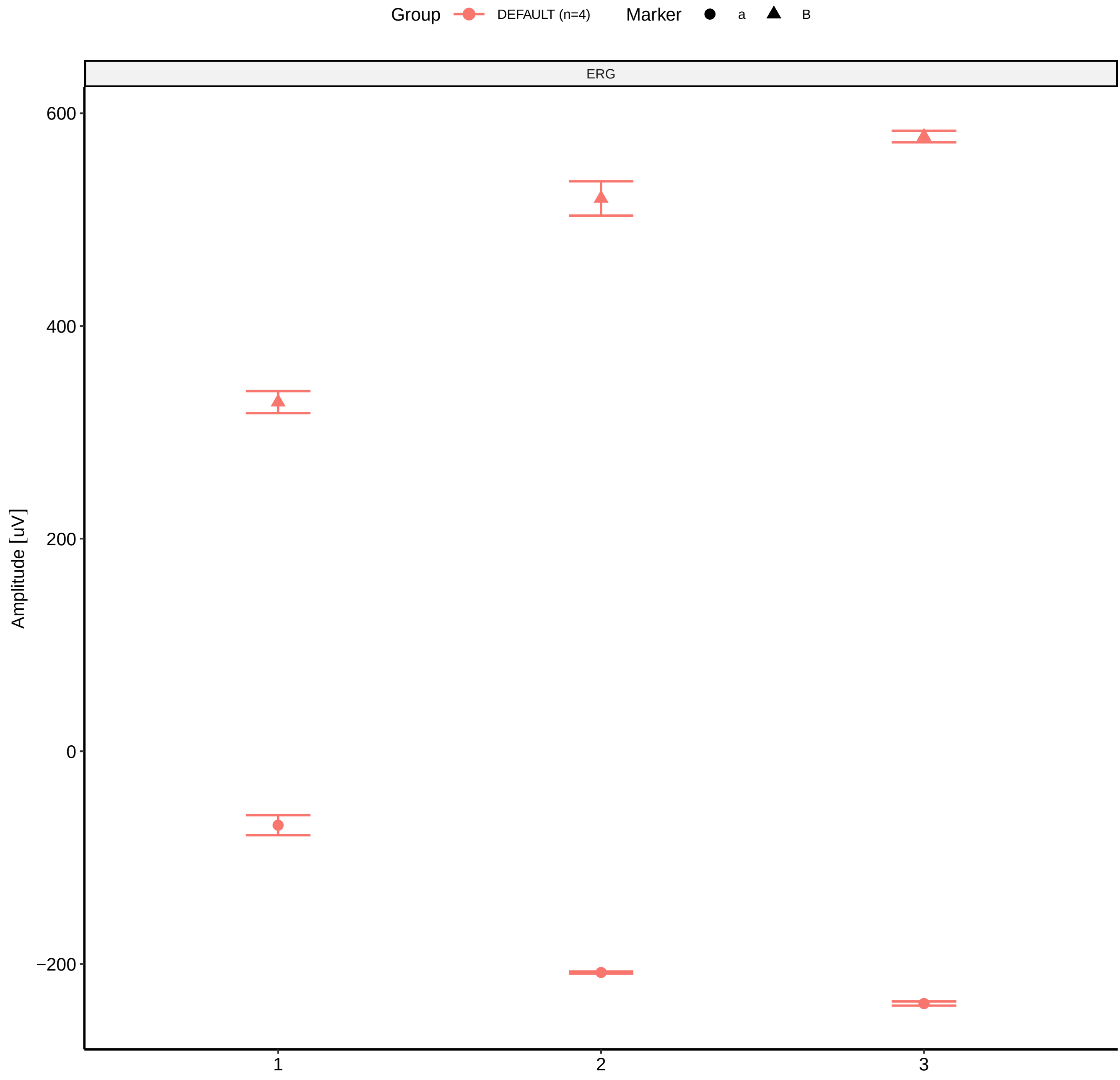

Group
DEFAULT (n=4)
Marker
a
B
ERG
600
400
200
0
−200
Amplitude [uV]
1
2
3

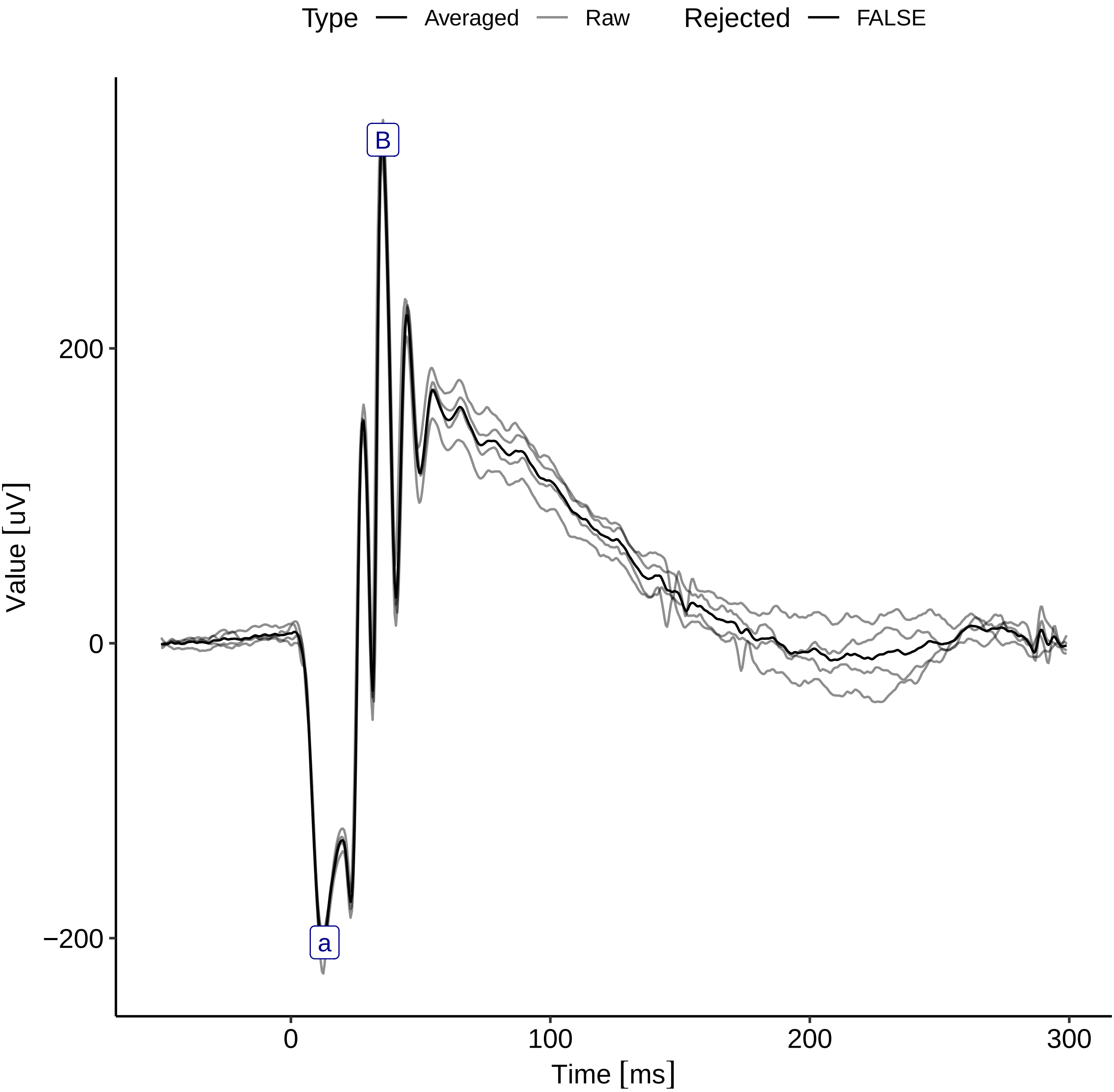
Type
Averaged
Raw
Rejected
FALSE
B
a
200
0
−200
Value [uV]
0
100
200
300
Time [ms]

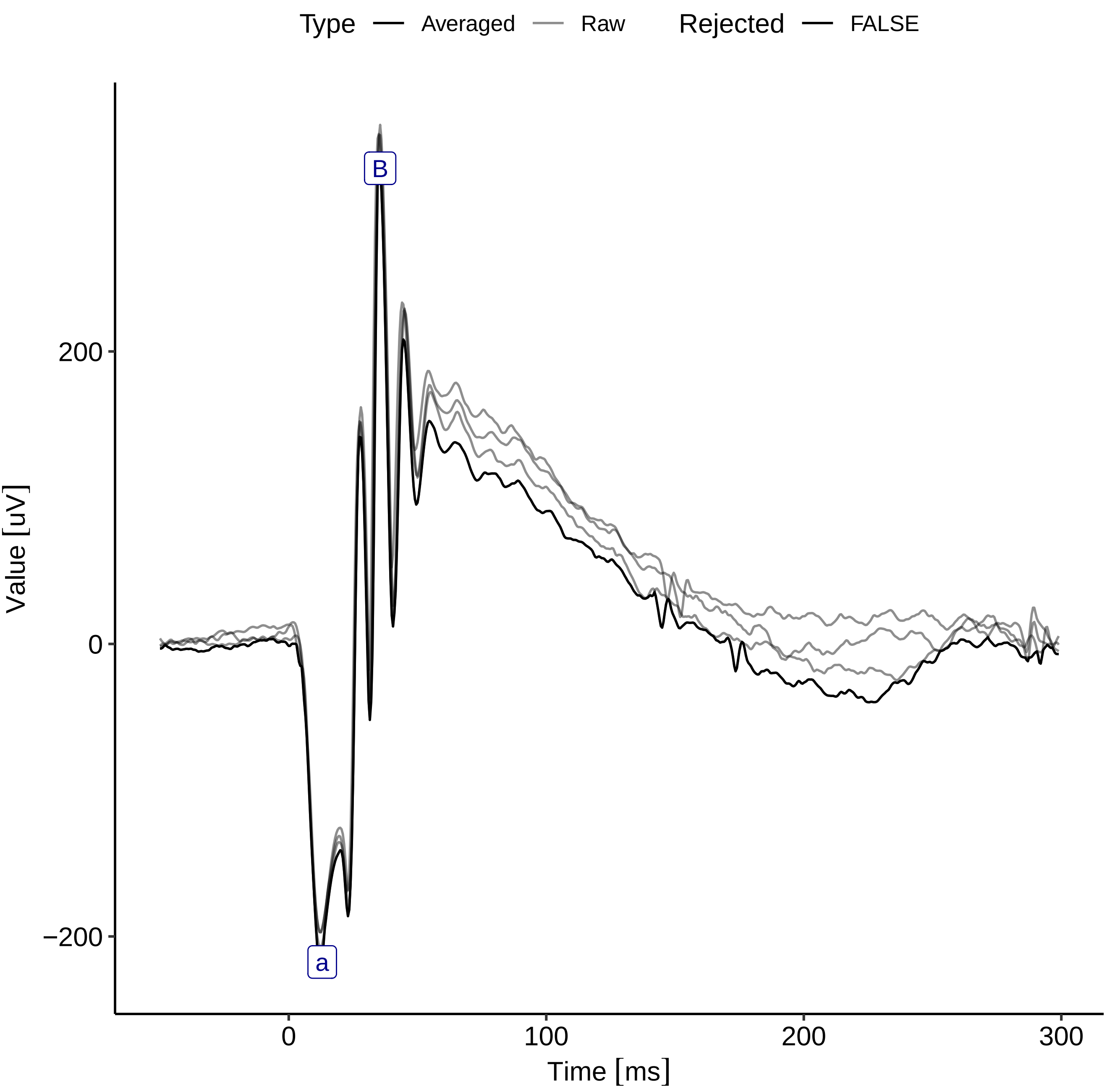
Type
Averaged
Raw
Rejected
FALSE
B
a
200
0
−200
0
100
200
300
Value [uV]
Time [ms]
