## Supplementary figures and images for "The ERGtools2 package: A Toolset for Processing and Analysing Visual Electrophysiology Data"

### dot-SampleERGExam-1.png

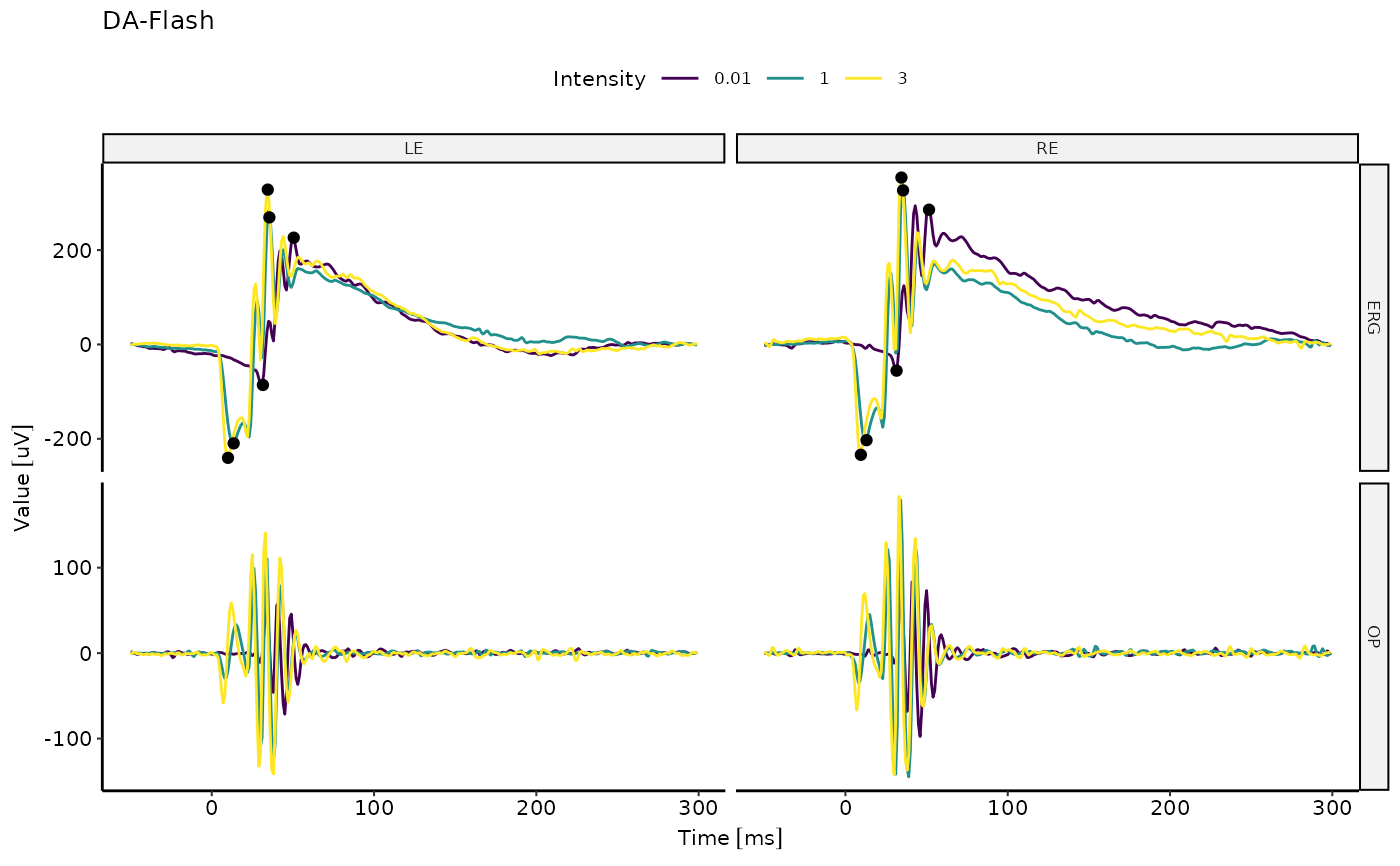

### ERGtools2-package-1.png

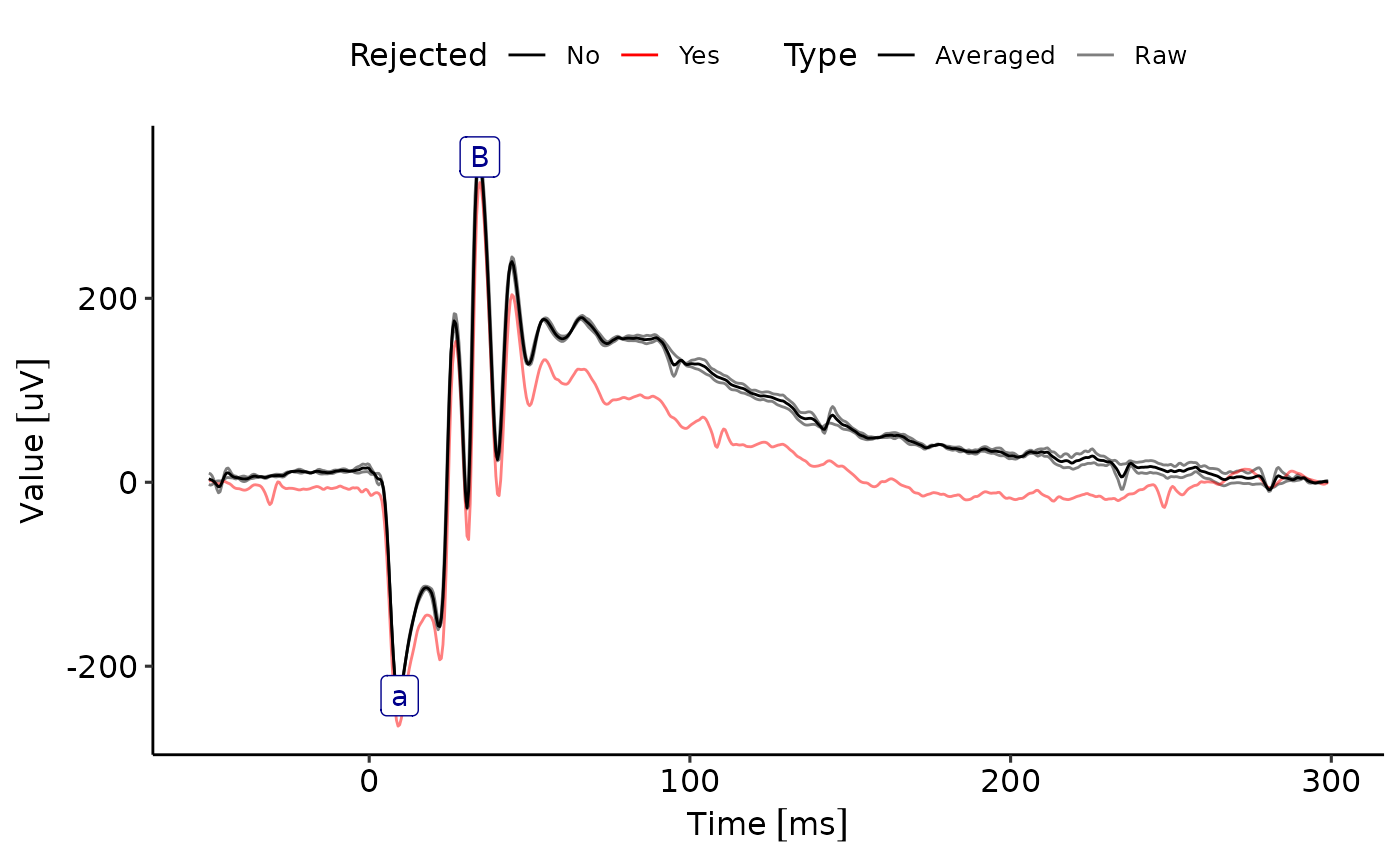

### ggERGExam-3.png

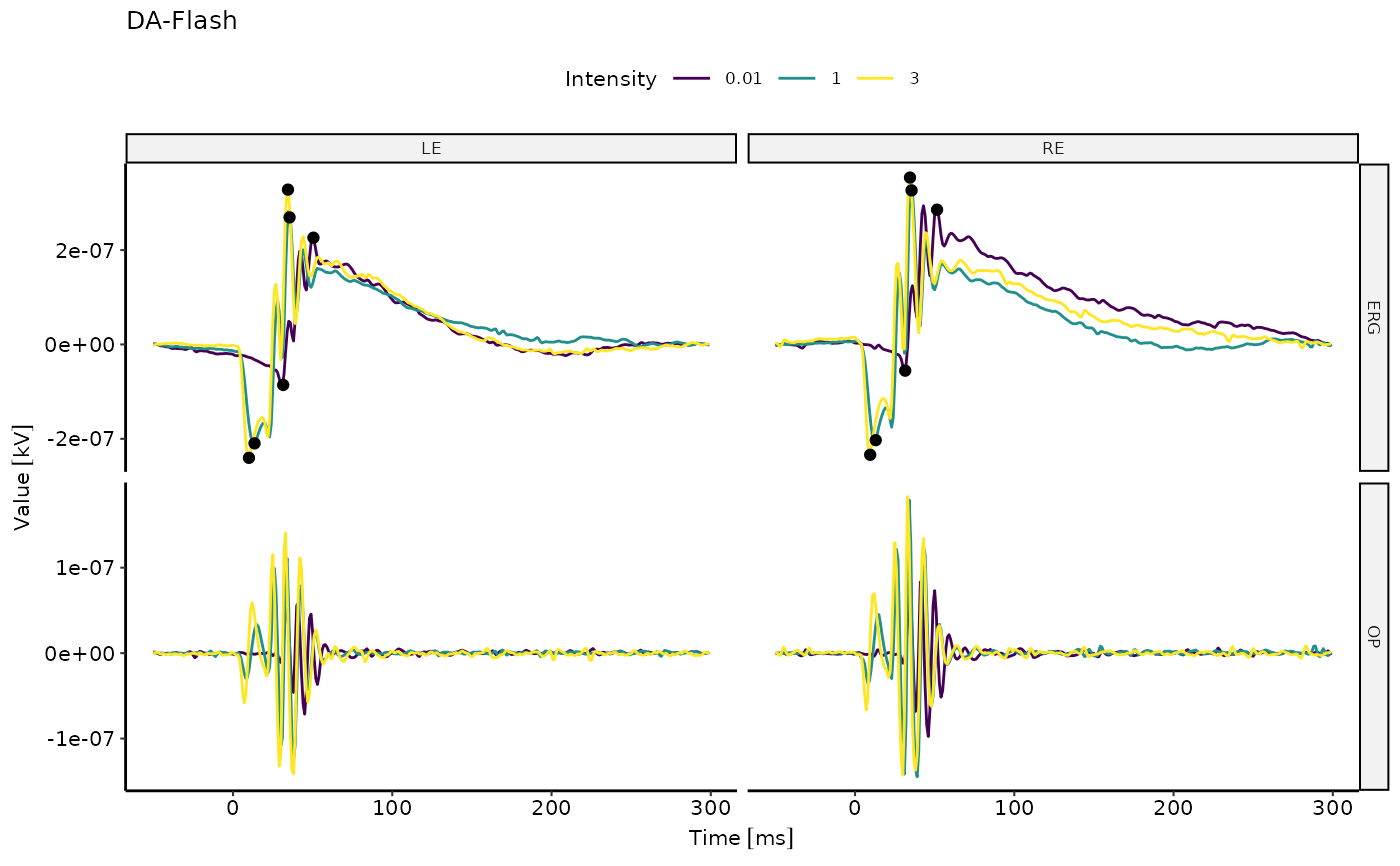

### ggIntensitySequence-1.png

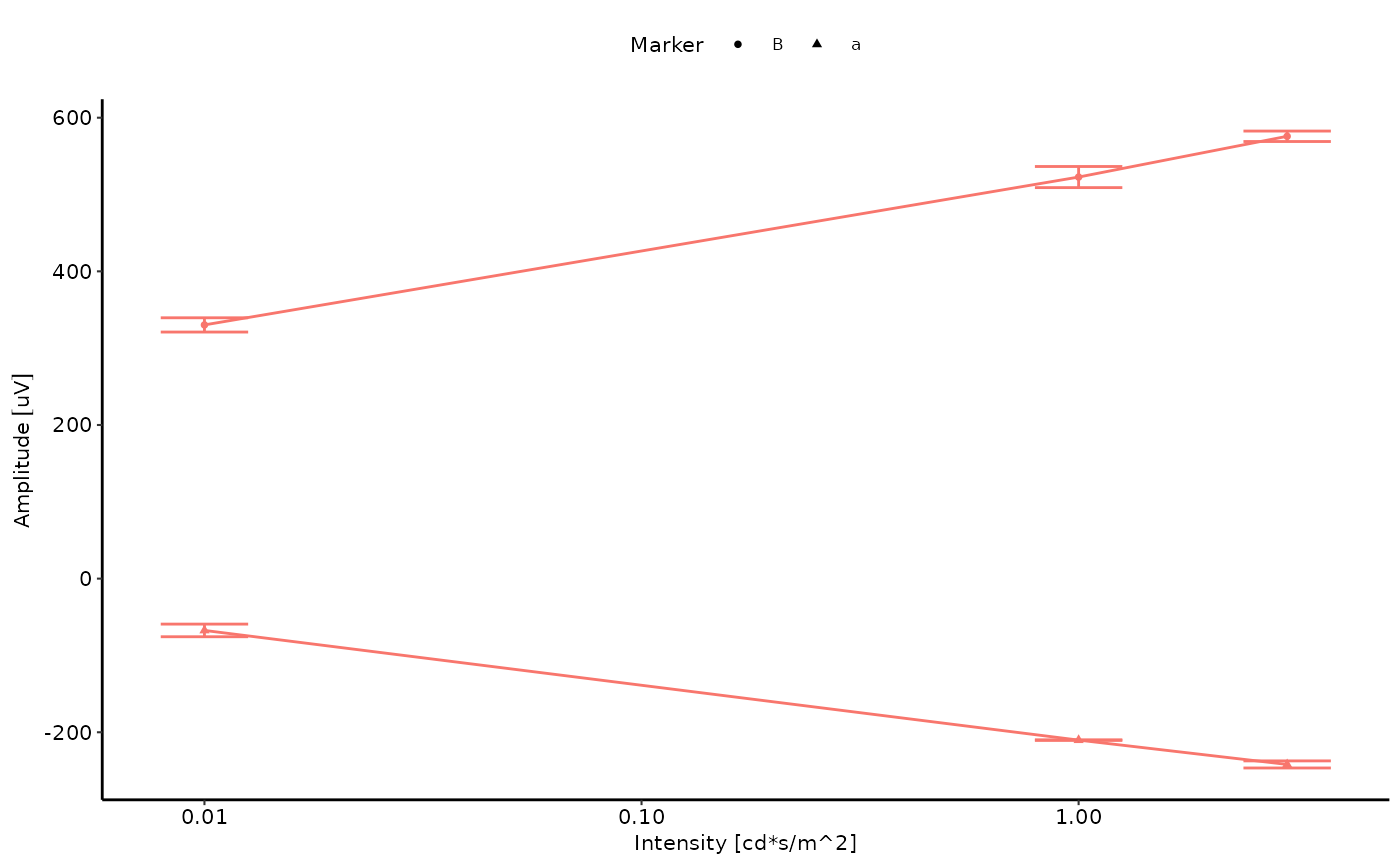

### ggStepSequence-1.png

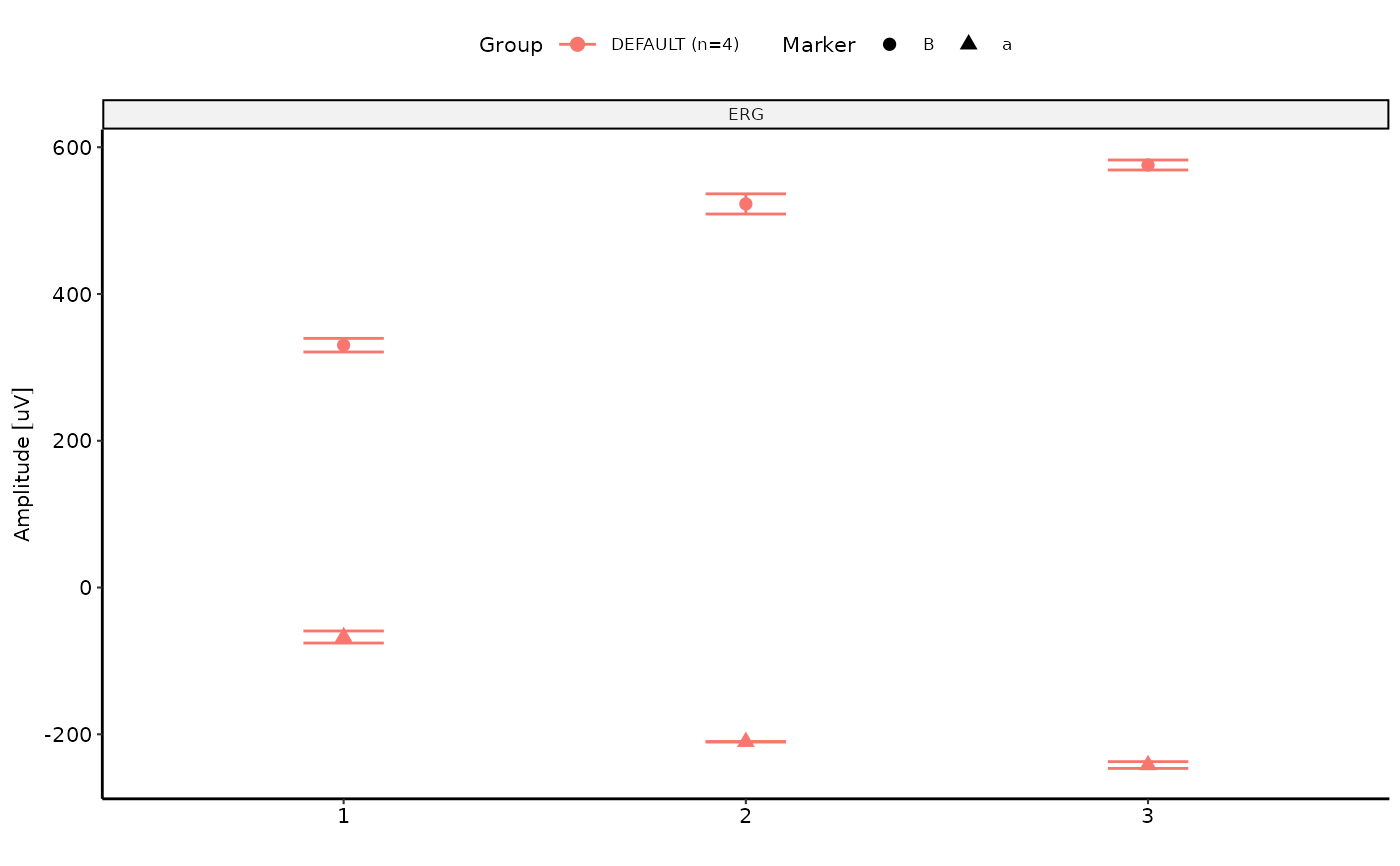

### README-example-1.png

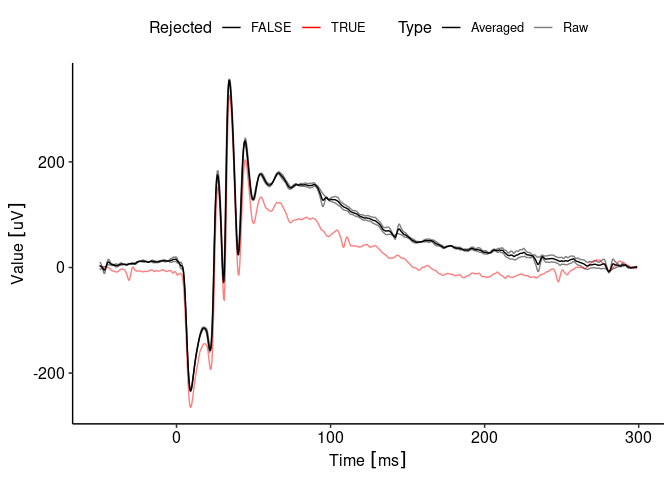

### README-example-2.png

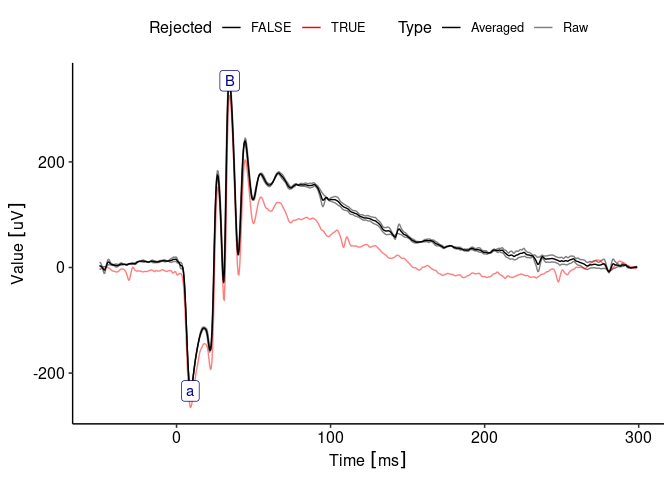

### README-pressure-1.png

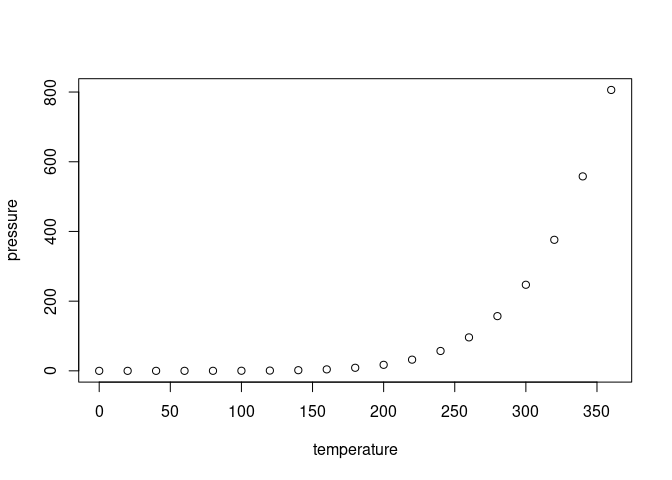

### Rplot001.png

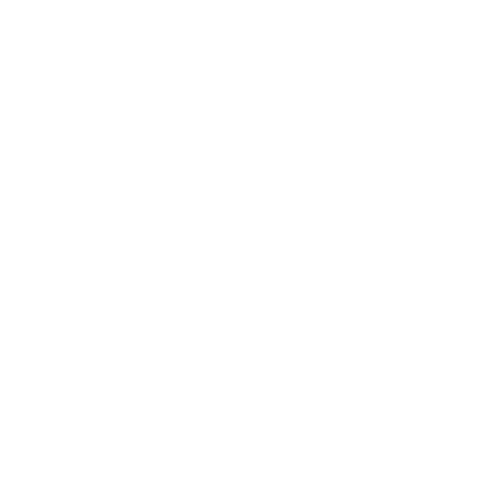

### Rplot002.png

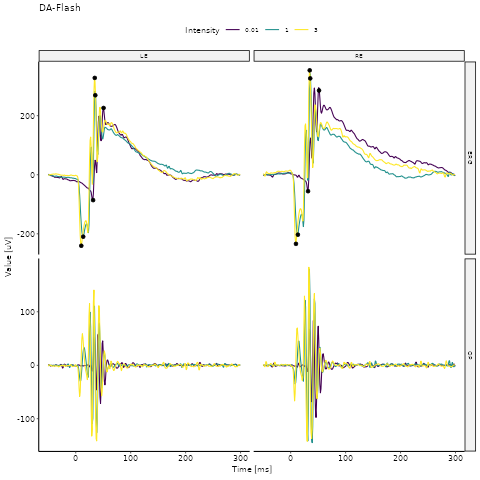

### Rplot003.png

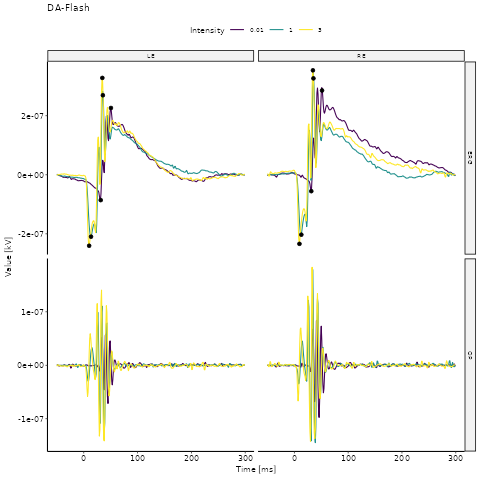

### UpdateProcessingMethods-1.png

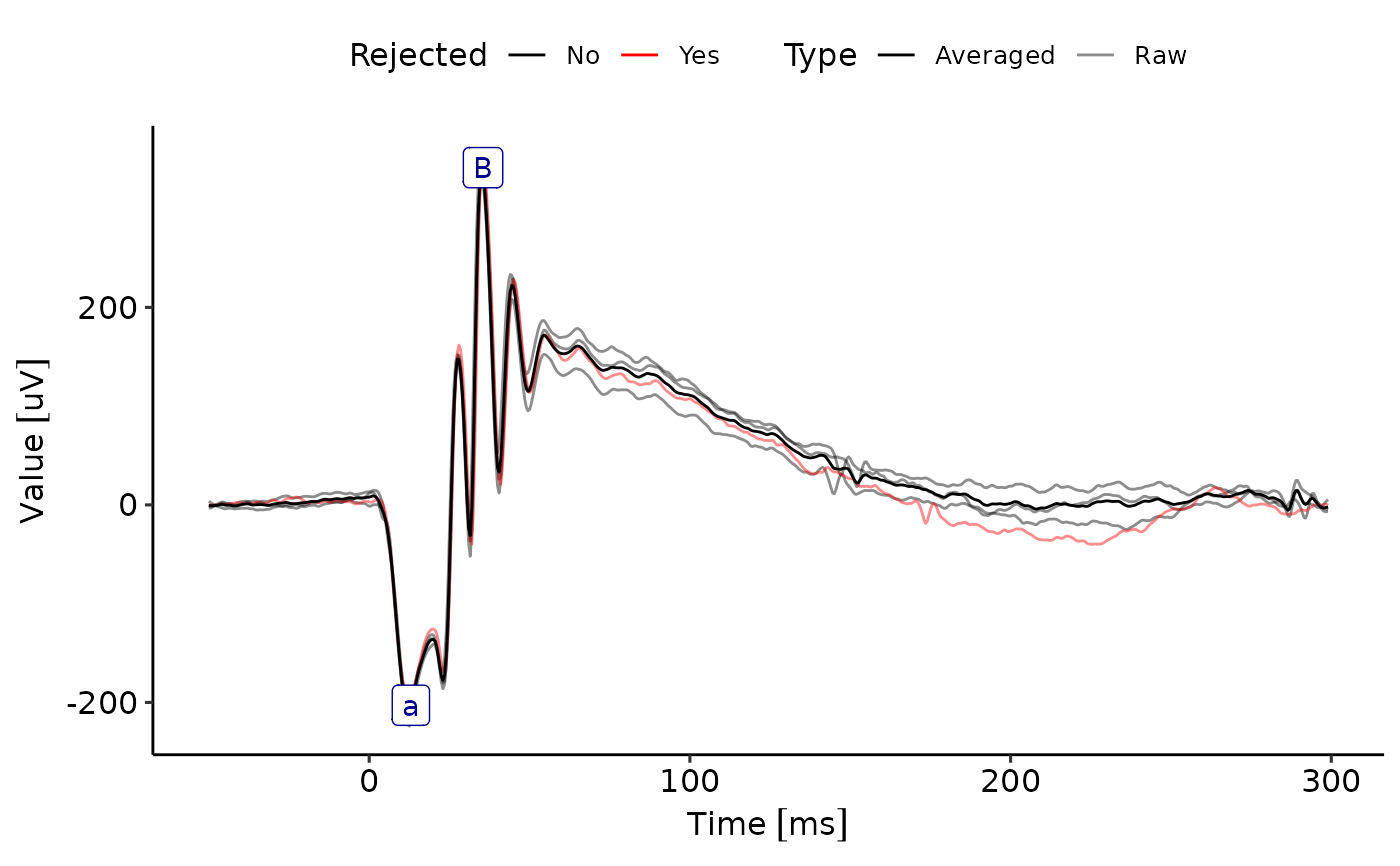

### UpdateProcessingMethods-2.png

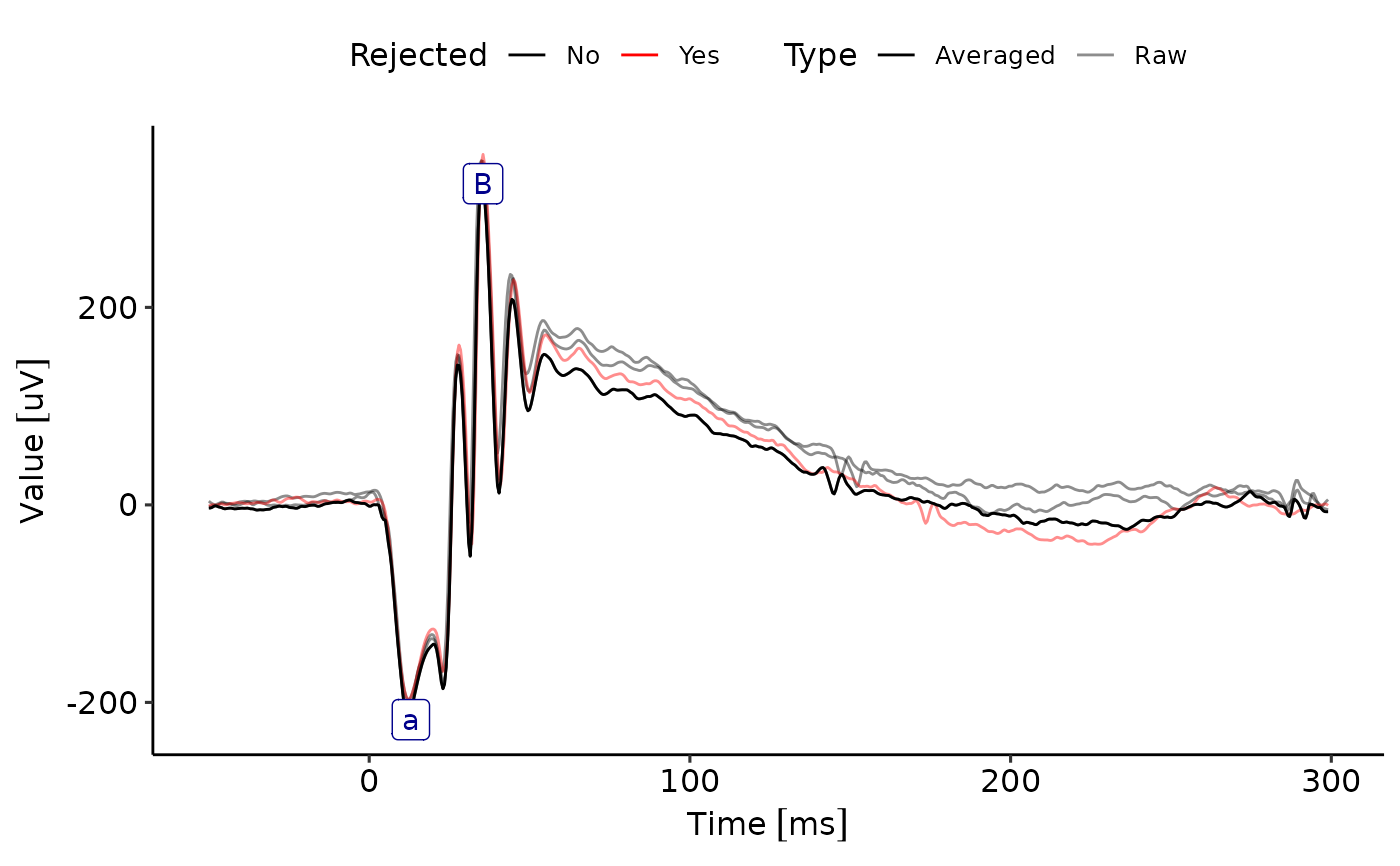
