## Supplemental Material 2 for "The ERGtools2 package: A Toolset for Processing and Analysing Visual Electrophysiology Data"

### ERGtools2 - Examples of how to create an ERGExam object from a set of CSV files

Moritz Lindner

15 Oktober, 2024

**Session info** **12**

*FilePath: "/media/moritz/DATADISK/02_Manuskripte/ERGtools2"*

```
require(ERGtools2)
```

```
## Lade nötiges Paket: ERGtools2

## Lade nötiges Paket: EPhysData

## Lade nötiges Paket: units

## udunits database from /usr/share/xml/udunits/udunits2.xml

## Registered S3 method overwritten by 'quantmod':
##   method            from
##   as.zoo.data.frame zoo
```

```
require(tidyr)
```

```
## Lade nötiges Paket: tidyr
```

```
require(stringr)
```

```
## Lade nötiges Paket: stringr
```

### Case 1: Raw data for each Recording are stored in a separate file

#### Make the example data: Extract data from an ERGexam object and store into CSV file

Metadata and stimulus information are stored in one file each, then each Recording of the exam is stored in a separate file, first column containing the time, and the sequential columns the individual trials

```
data(ERG)
MD <- Metadata(ERG)
Stim <- Stimulus(ERG)
dir.create("CVS_Example")
```

```
## Warning in dir.create("CVS_Example"): 'CVS_Example' existiert bereits
```

```
write.csv2(MD, paste0(getwd(), "/CVS_Example/Metadata.csv"), row.names =
             F) # may need adjustment depending on Operating System
write.csv2(Stim, paste0(getwd(), "/CVS_Example/Stimulus.csv"), row.names =
             F) # may need adjustment depending on Operating System
for (i in 1:nrow(MD)) {
  df <- as.data.frame(Subset(ERG,where=i),IncludeRejected = T)
  df <- df[,c("Trial","Time","Value")]
  df <- pivot_wider(
    df,
    names_from = "Trial",
    names_prefix = "Trial_",
    values_from = "Value"
  )
  fn <-
    paste0("ERG-Step-",
           MD$Step[i],
           "_Ch-",
           MD$Channel[i],
           "_Eye-",
           MD$Eye[i],
           ".csv")
  write.csv2(df, paste0(getwd(), "/CVS_Example/", fn), row.names = F, dec = ".") # may need adjustment
}
```

```
## Warning in write.csv2(df, paste0(getwd(), "/CVS_Example/", fn), row.names = F, :
## attempt to set 'dec' ignored
```

```
## Warning in write.csv2(df, paste0(getwd(), "/CVS_Example/", fn), row.names = F, :
## attempt to set 'dec' ignored
```

```
## Warning in write.csv2(df, paste0(getwd(), "/CVS_Example/", fn), row.names = F, :
## attempt to set 'dec' ignored

## Warning in write.csv2(df, paste0(getwd(), "/CVS_Example/", fn), row.names = F, :
## attempt to set 'dec' ignored

## Warning in write.csv2(df, paste0(getwd(), "/CVS_Example/", fn), row.names = F, :
## attempt to set 'dec' ignored

## Warning in write.csv2(df, paste0(getwd(), "/CVS_Example/", fn), row.names = F, :
## attempt to set 'dec' ignored

## Warning in write.csv2(df, paste0(getwd(), "/CVS_Example/", fn), row.names = F, :
## attempt to set 'dec' ignored

## Warning in write.csv2(df, paste0(getwd(), "/CVS_Example/", fn), row.names = F, :
## attempt to set 'dec' ignored

## Warning in write.csv2(df, paste0(getwd(), "/CVS_Example/", fn), row.names = F, :
## attempt to set 'dec' ignored

## Warning in write.csv2(df, paste0(getwd(), "/CVS_Example/", fn), row.names = F, :
## attempt to set 'dec' ignored

## Warning in write.csv2(df, paste0(getwd(), "/CVS_Example/", fn), row.names = F, :
## attempt to set 'dec' ignored

## Warning in write.csv2(df, paste0(getwd(), "/CVS_Example/", fn), row.names = F, :
## attempt to set 'dec' ignored
```

This is the content of the directory:

```
list.files(paste0(getwd(), "/CVS_Example/"))
```

```
##  [1] "ERG-Step-1_Ch-ERG_Eye-LE.csv" "ERG-Step-1_Ch-ERG_Eye-RE.csv"
##  [3] "ERG-Step-1_Ch-OP_Eye-LE.csv"  "ERG-Step-1_Ch-OP_Eye-RE.csv"
##  [5] "ERG-Step-1.csv"               "ERG-Step-2_Ch-ERG_Eye-LE.csv"
##  [7] "ERG-Step-2_Ch-ERG_Eye-RE.csv" "ERG-Step-2_Ch-OP_Eye-LE.csv"
##  [9] "ERG-Step-2_Ch-OP_Eye-RE.csv"  "ERG-Step-2.csv"
## [11] "ERG-Step-3_Ch-ERG_Eye-LE.csv" "ERG-Step-3_Ch-ERG_Eye-RE.csv"
## [13] "ERG-Step-3_Ch-OP_Eye-LE.csv"  "ERG-Step-3_Ch-OP_Eye-RE.csv"
## [15] "ERG-Step-3.csv"               "Metadata.csv"
## [17] "Stimulus.csv"
```

This is how the data in each file looks like:

```
head(read.csv2(paste0(getwd(), "/CVS_Example/", fn), dec = "."))
```

```
##    Time  Trial_1    Trial_2    Trial_3
## 1 -50.0 4502.994 -1815.1550   97.91454
## 2 -49.5 3901.460  -770.1742 1411.19934
## 3 -49.0 3787.964   437.4963 2480.57544
```

```
## 4 -48.5 3622.657  1076.1021 2860.36719
## 5 -48.0 2979.335  1277.6841 2616.53784
## 6 -47.5 1864.525  1173.5896 2426.05591
```

```
rm(list = ls())
```

##### Create an ERGExam from the data just stored

```
Metadata<-read.csv2(paste0(getwd(),"/CVS_Example/Metadata.csv"))
Stimulus<-read.csv2(paste0(getwd(),"/CVS_Example/Stimulus.csv"))

Recordings <- list() # make a list to store the recordings from each file (as EPhysData objects), this
for (i in 1:nrow(Metadata)) {
  fn <-
    paste0(
      "ERG-Step-",
      Metadata$Step[i],
      "_Ch-",
      Metadata$Channel[i],
      "_Eye-",
      Metadata$Eye[i],
      ".csv"
    )
  print(fn)
  # read the data for each recording, split into data and time trace and assign units
  df <- read.csv2(paste0(getwd(), "/CVS_Example/", fn), dec = ".")
  recording.data <- df[, str_detect(colnames(df), "Trial")]
  recording.data <- as_units(as.matrix(recording.data), "nV")
  time <- df[, "Time"]
  time <- as_units(time, "ms")
  # dreate the EPhysData object
  Recordings[[i]] <-
    newEPhysData(Data = recording.data, TimeTrace = time)
}
```

```
## [1] "ERG-Step-1_Ch-ERG_Eye-RE.csv"
## [1] "ERG-Step-1_Ch-ERG_Eye-LE.csv"
## [1] "ERG-Step-1_Ch-OP_Eye-RE.csv"
## [1] "ERG-Step-1_Ch-OP_Eye-LE.csv"
## [1] "ERG-Step-2_Ch-ERG_Eye-RE.csv"
## [1] "ERG-Step-2_Ch-ERG_Eye-LE.csv"
## [1] "ERG-Step-2_Ch-OP_Eye-RE.csv"
## [1] "ERG-Step-2_Ch-OP_Eye-LE.csv"
## [1] "ERG-Step-3_Ch-ERG_Eye-RE.csv"
## [1] "ERG-Step-3_Ch-ERG_Eye-LE.csv"
## [1] "ERG-Step-3_Ch-OP_Eye-RE.csv"
## [1] "ERG-Step-3_Ch-OP_Eye-LE.csv"
```

```
# An ERGExam object requires additional information on the subject (e.g. name, DOB) as well as on the e
SubjectInfo <- list(Subject = "Test", DOB = as.Date("2000-01-01"))
ExamInfo <- list(ProtocolName = "TestProtocol", ExamDate = as.POSIXct("2024-01-01"))
```

```r
# Now the ERGExam object is assembled
ImportedExam<-newERGExam(Data = Recordings,
                         Metadata = Metadata,
                         Stimulus = Stimulus,
                         Averaged = F,
                         ExamInfo = ExamInfo,
                         SubjectInfo = SubjectInfo)

validObject(ImportedExam)
```

```
## [1] TRUE
```

```r
# Compare (re)imported Exam to the original exam

data(ERG)
Original<-ERG

ImportedExam<-SetStandardFunctions(ImportedExam)
Original<-SetStandardFunctions(Original)

ggERGExam(ImportedExam)
```

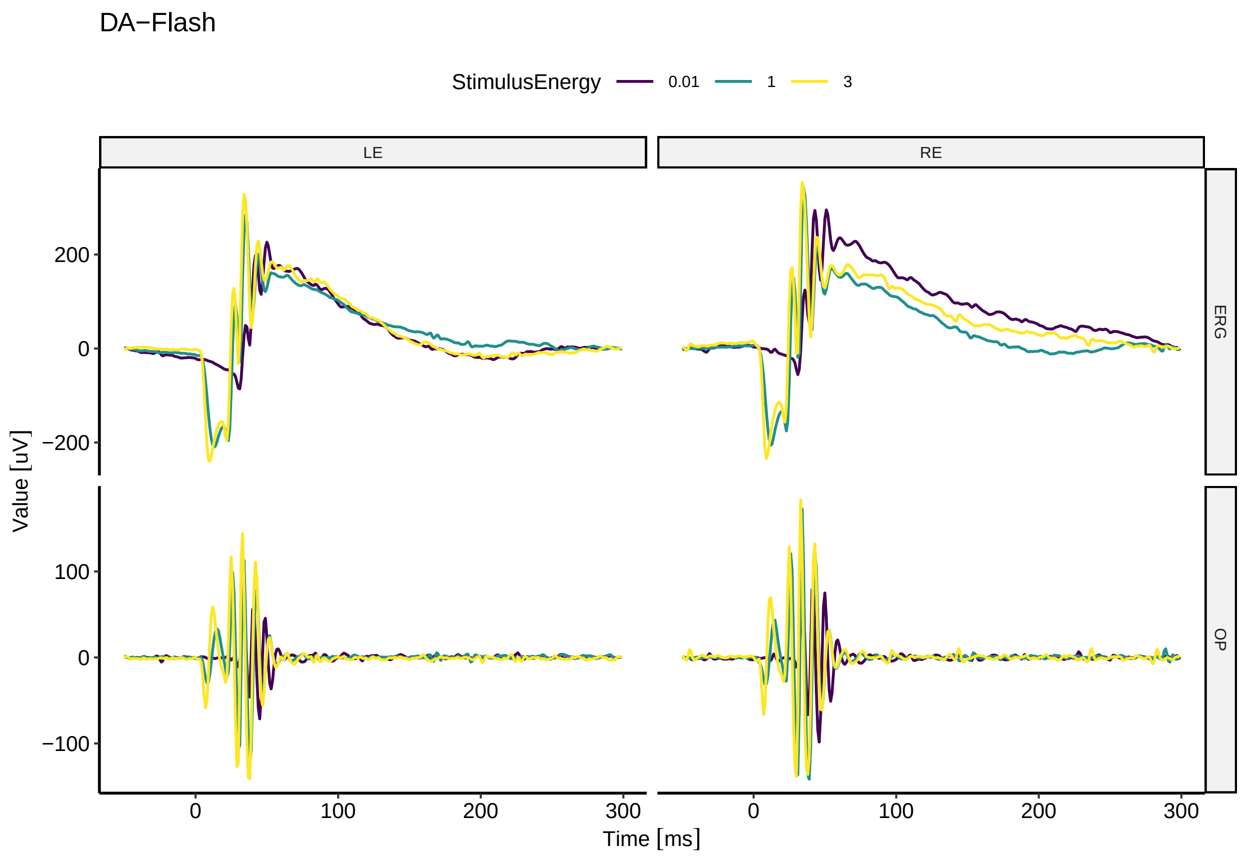

```
ggERGExam(Original)
```

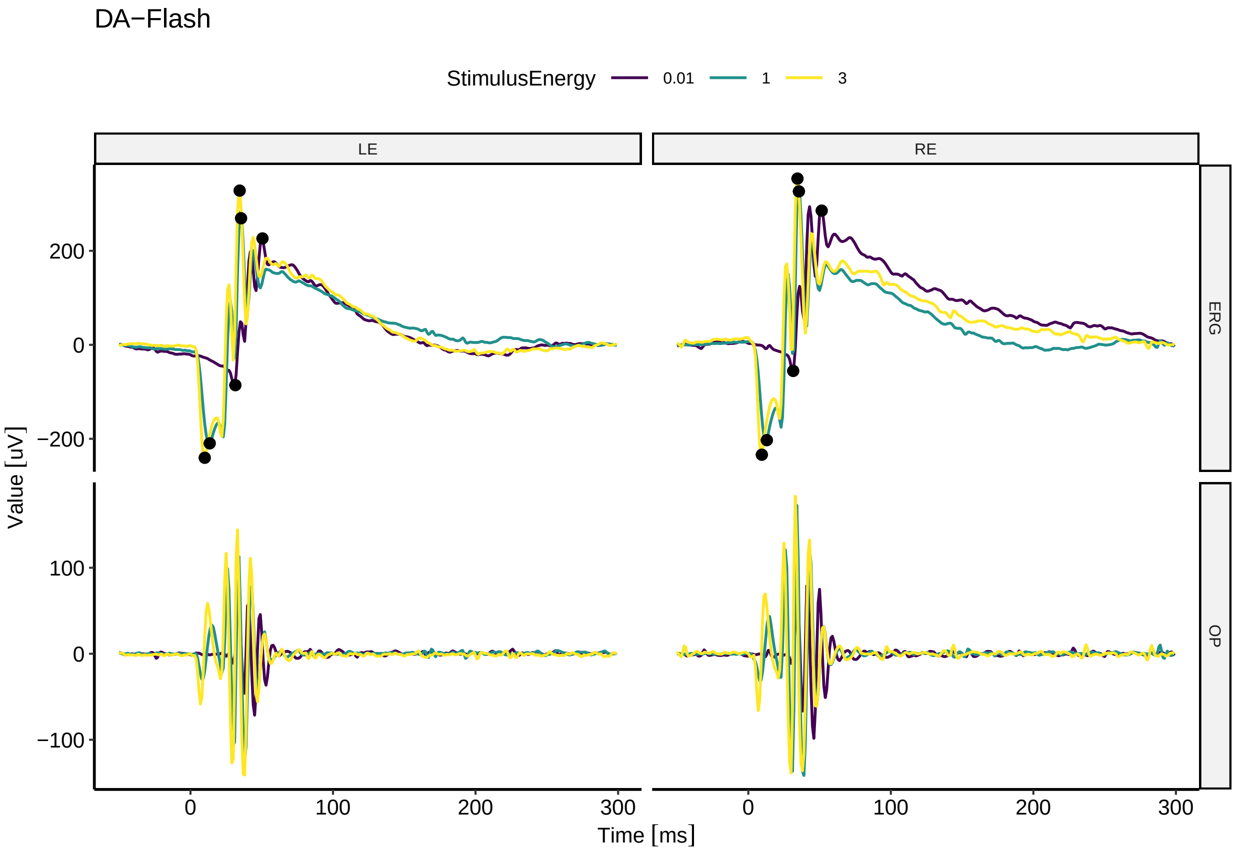

#### Case 2: Average traces from all Recordings per step/stimulus condition

##### Make the example data: Extract data from an ERGexam object and store into CSV file

Metadata and stimulus information are stored in one file each, then each Recording of the exam is stored in a separate file, first column containing the time, and the sequential columns the individual trials

```
unlink(paste0(getwd(), "/CVS_Example/"),recursive=T)
rm(list = ls())
data(ERG)
ERG<- Subset(ERG, Raw=F) # make object to store the averaged traces instead of raw data
MD <- Metadata(ERG)
Stim <- Stimulus(ERG)
dir.create("CVS_Example")
write.csv2(MD, paste0(getwd(), "/CVS_Example/Metadata.csv"), row.names =
             F) # may need adjustment depending on Operating System
write.csv2(Stim, paste0(getwd(), "/CVS_Example/Stimulus.csv"), row.names =
             F) # may need adjustment depending on Operating System
```

```r
for (i in 1:nrow(Stim)) {
  df <- as.data.frame(Subset(ERG,where=list(Step=Stim$Step[[i]]),IncludeRejected = T)
  df <- df[,c("Channel","Eye","Time","Value")]
  df <- pivot_wider(
    df,
    names_from = c("Channel","Eye"),
    names_prefix = "Value_",
    values_from = "Value"
  )
  fn <-
    paste0("ERG-Step-",
           Stim$Step[[i]],
           ".csv")
  write.csv2(df, paste0(getwd(), "/CVS_Example/", fn), row.names = F, dec = ".") # may need adjustment
}
```

```
## Warning in write.csv2(df, paste0(getwd(), "/CVS_Example/", fn), row.names = F, :
## attempt to set 'dec' ignored

## Warning in write.csv2(df, paste0(getwd(), "/CVS_Example/", fn), row.names = F, :
## attempt to set 'dec' ignored

## Warning in write.csv2(df, paste0(getwd(), "/CVS_Example/", fn), row.names = F, :
## attempt to set 'dec' ignored
```

This is the content of the directory:

```r
list.files(paste0(getwd(), "/CVS_Example/"))
```

```
## [1] "ERG-Step-1.csv" "ERG-Step-2.csv" "ERG-Step-3.csv" "Metadata.csv"
## [5] "Stimulus.csv"
```

This is how the data in each file looks like, note the column names contain the channel identifier (ERG/OP) and the eye identifier (RE/LE)

```r
head(read.csv2(paste0(getwd(), "/CVS_Example/", fn), dec = "."))
```

```
##    Time Value_ERG_RE Value_ERG_LE Value_OP_RE Value_OP_LE
## 1 -50.0    3797.9881   -2465.3628   2375.0339   -225.8795
## 2 -49.5    3014.5190   -1755.6906   1136.0682    360.1678
## 3 -49.0    2254.7980    -606.4151     32.0034   1081.8208
## 4 -48.5    1034.7684     172.9521  -1032.9668   1366.6543
## 5 -48.0    -827.4936     585.9339  -2169.7392   1138.6010
## 6 -47.5   -3086.9021     906.4600  -3207.4051    669.2753
```

```r
rm(list = ls())
```

#### Create an ERGExam from the data just stored

```r
Metadata<-read.csv2(paste0(getwd(),"/CVS_Example/Metadata.csv"))
Stimulus<-read.csv2(paste0(getwd(),"/CVS_Example/Stimulus.csv"))

Recordings <- list() # make a list to store the recordings from each file (as EPhysData objects), this
for (i in 1:nrow(Stimulus)) {
  fn <-
    paste0("ERG-Step-",
           Stimulus$Step[[i]],
           ".csv")
  print(fn)
  # read the data for each step, split into data and time trace and assign units
  df <- read.csv2(paste0(getwd(), "/CVS_Example/", fn), dec = ".")
  trace.data <- df[, str_detect(colnames(df), "Value")]
  trace.data <- as_units(as.matrix(trace.data), "nV")
  time <- df[, "Time"]
  time <- as_units(time, "ms")
  # dreate the EPhysData object
  current.metadata<-Metadata[Metadata$Step==Stimulus$Step[[i]],]
  for (j in 1:nrow(current.metadata)){ # for each Channel and Eye that has data from under the current
    current.column.name <-
      paste0("Value_",
             current.metadata$Channel[[j]],
             "_",
             current.metadata$Eye[[j]])
    current.trace<-trace.data[,current.column.name]
    Recordings<-append(Recordings, newEPhysData(Data = current.trace, TimeTrace = time))
  }
}
```

```
## [1] "ERG-Step-1.csv"
## [1] "ERG-Step-2.csv"
## [1] "ERG-Step-3.csv"
```

```r
# An ERGExam object requires additional information on the subject (e.g. name, DOB) as well as on the e
SubjectInfo <- list(Subject = "Test", DOB = as.Date("2000-01-01"))
ExamInfo <- list(ProtocolName = "TestProtocol", ExamDate = as.POSIXct("2024-01-01"))

# Now the ERGExam object is assembled
ImportedExam<-newERGExam(Data = Recordings,
                         Metadata = Metadata,
                         Stimulus = Stimulus,
                         Averaged = F,
                         ExamInfo = ExamInfo,
                         SubjectInfo = SubjectInfo)

validObject(ImportedExam)
```

```
## [1] TRUE
```

```r
# Compare (re)imported Exam to the original exam

data(ERG)
```

```
Original<-ERG

ImportedExam<-SetStandardFunctions(ImportedExam)
```

```
## Can't set a Rejected function for 'X', because 'X' contains only one trial. Keeping it.
## Can't set a Rejected function for 'X', because 'X' contains only one trial. Keeping it.
## Can't set a Rejected function for 'X', because 'X' contains only one trial. Keeping it.
## Can't set a Rejected function for 'X', because 'X' contains only one trial. Keeping it.
## Can't set a Rejected function for 'X', because 'X' contains only one trial. Keeping it.
## Can't set a Rejected function for 'X', because 'X' contains only one trial. Keeping it.
```

```
Original<-SetStandardFunctions(Original)

ggERGExam(ImportedExam)
```

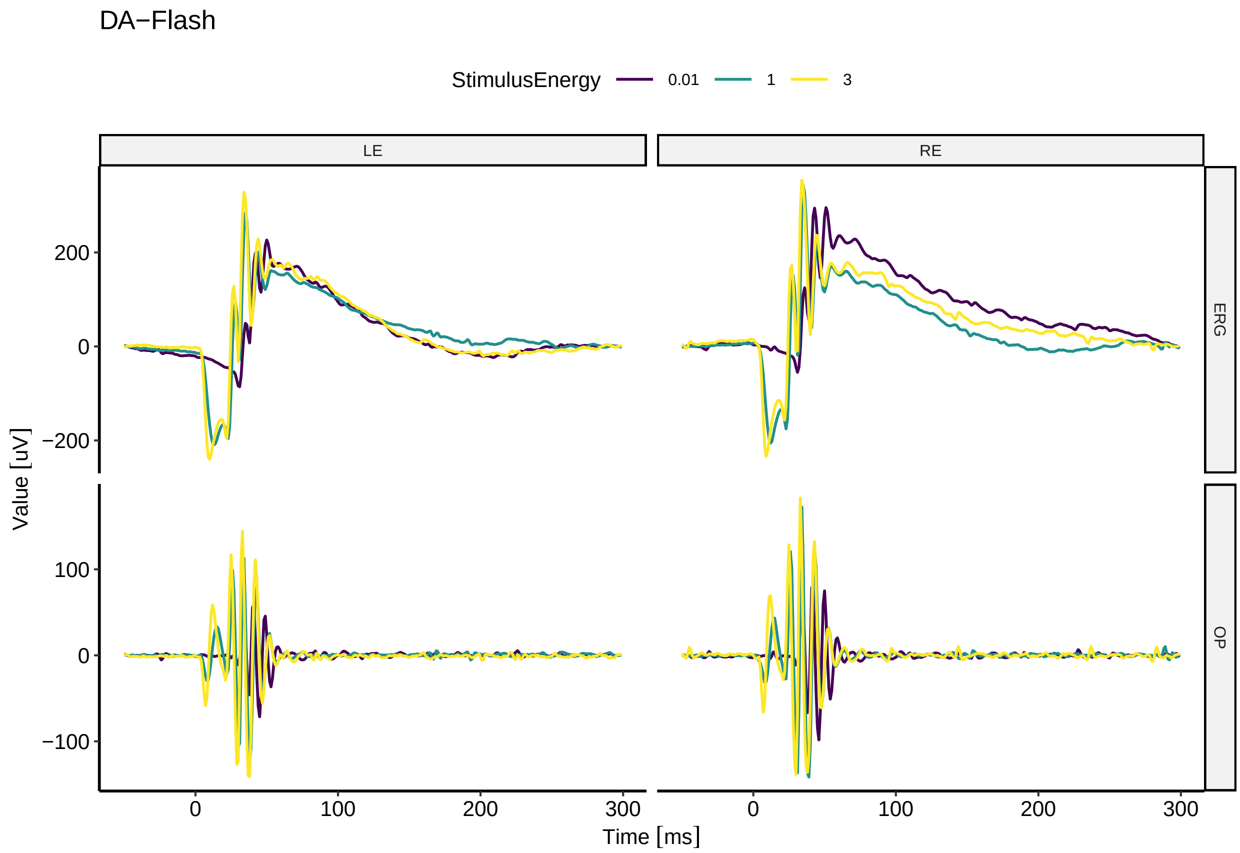

```
ggERGExam(Original)
```

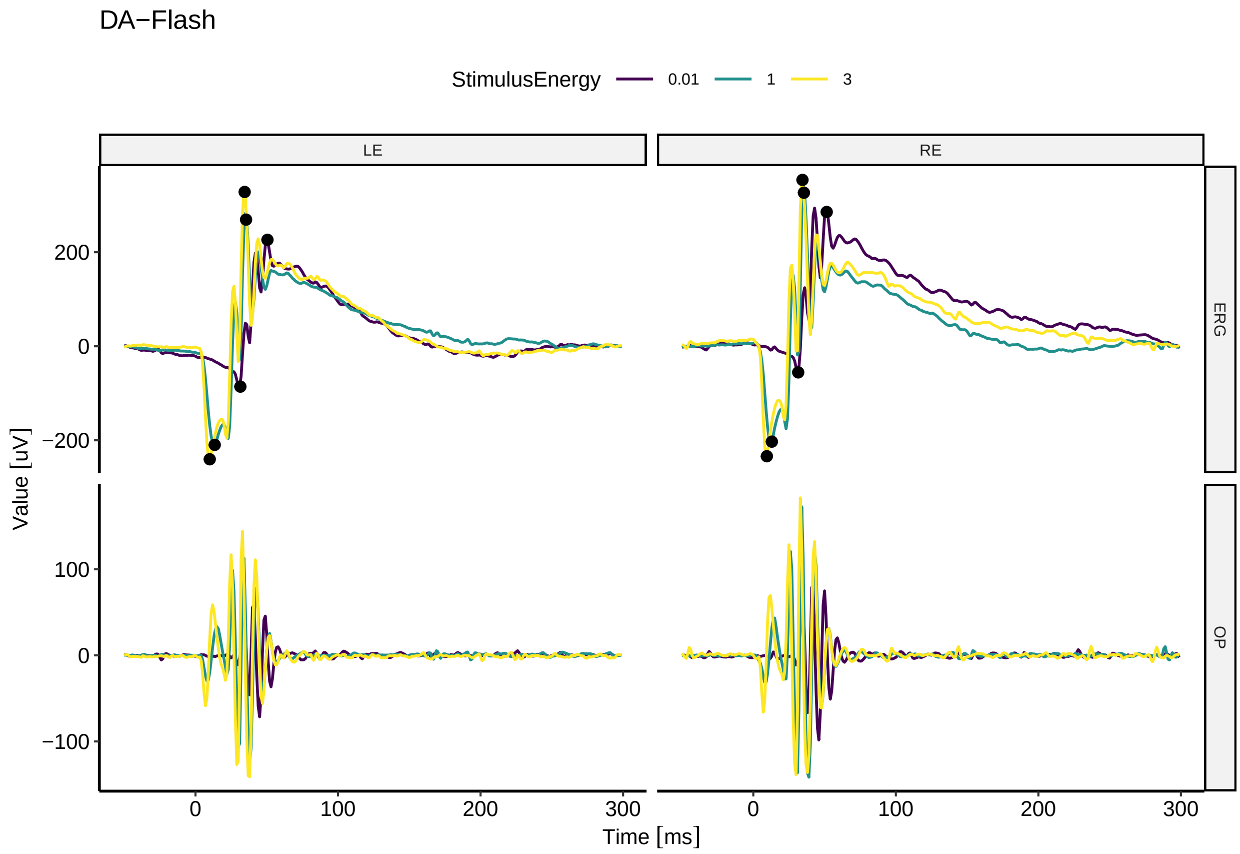

```
# Do some stuff with the imported exam (See other example script for more details)

ImportedExam_Subset<-DropRecordings(ImportedExam, where=list(Background="LA")) # drop all recordings ac
ggERGExam(ImportedExam_Subset)
```

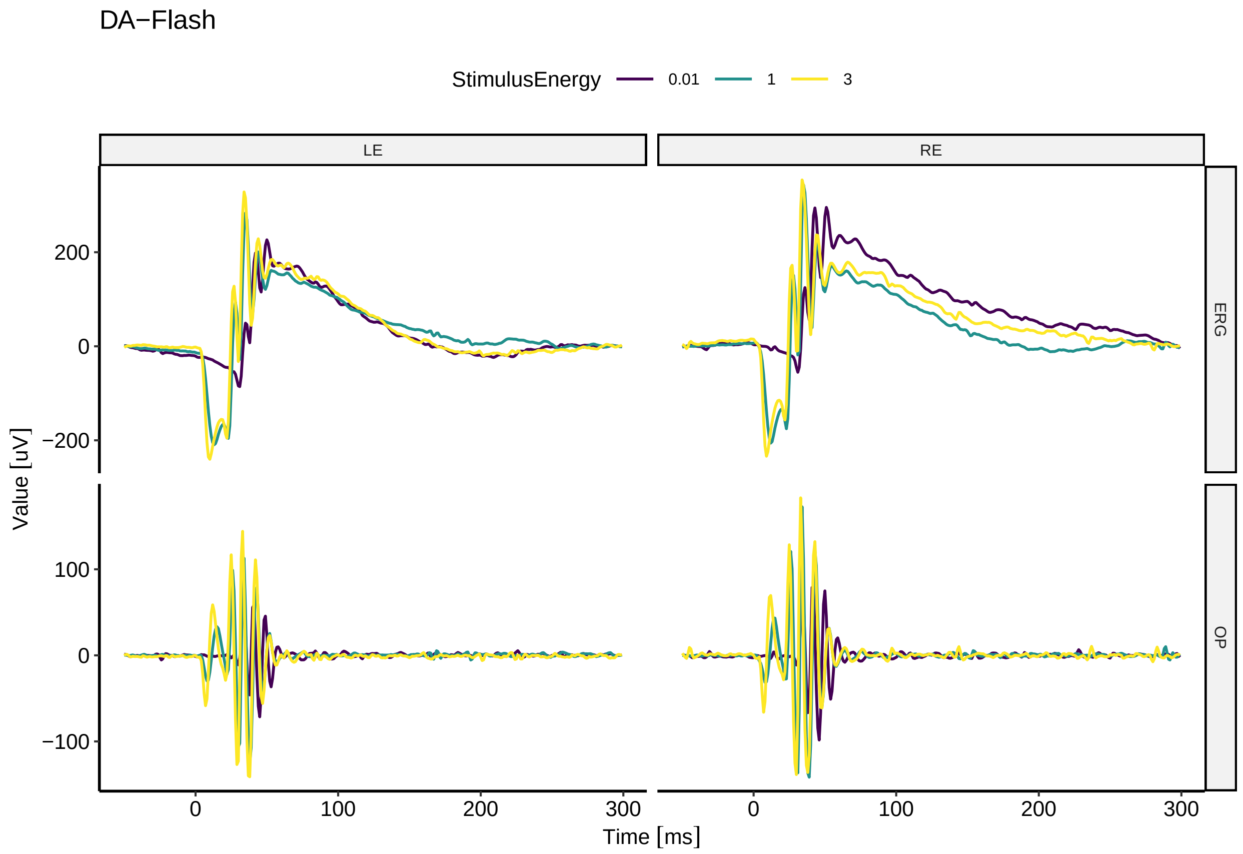

```r
ImportedExam<-AutoPlaceMarkers(ImportedExam)
ggERGExam(ImportedExam)
```

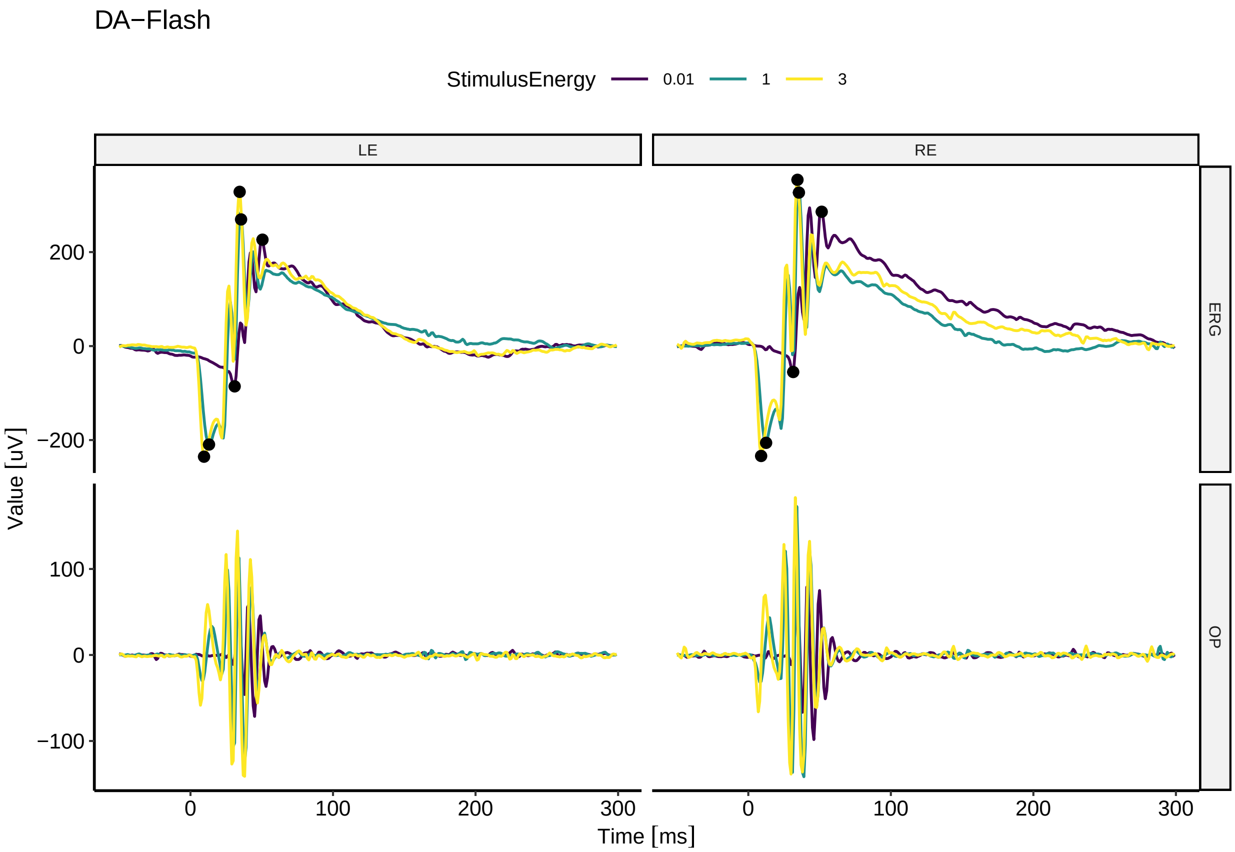

#### Session info

```
sessionInfo()
```

```
## R version 4.1.2 (2021-11-01)
## Platform: x86_64-pc-linux-gnu (64-bit)
## Running under: Ubuntu 22.04.5 LTS
##
## Matrix products: default
## BLAS:   /usr/lib/x86_64-linux-gnu/blas/libblas.so.3.10.0
## LAPACK: /usr/lib/x86_64-linux-gnu/lapack/liblapack.so.3.10.0
##
## locale:
##  [1] LC_CTYPE=de_DE.UTF-8       LC_NUMERIC=C
##  [3] LC_TIME=de_DE.UTF-8        LC_COLLATE=de_DE.UTF-8
##  [5] LC_MONETARY=de_DE.UTF-8    LC_MESSAGES=de_DE.UTF-8
##  [7] LC_PAPER=de_DE.UTF-8       LC_NAME=C
##  [9] LC_ADDRESS=C               LC_TELEPHONE=C
## [11] LC_MEASUREMENT=de_DE.UTF-8 LC_IDENTIFICATION=C
##
## attached base packages:
## [1] stats     graphics  grDevices utils     datasets  methods   base
##
```

```
## other attached packages:
## [1] stringr_1.5.1   tidyr_1.1.3     ERGtools2_0.8.0 EPhysData_0.9.7
## [5] units_0.8-0
##
## loaded via a namespace (and not attached):
##  [1] xts_0.12.1         bit64_4.0.5        httr_1.4.2         tools_4.1.2
##  [5] backports_1.2.1    utf8_1.2.1         R6_2.5.0           DT_0.19
##  [9] DBI_1.1.1          lazyeval_0.2.2     colorspace_2.0-1   withr_2.5.2
## [13] tidyselect_1.1.1   gridExtra_2.3      bit_4.0.4          curl_4.3.1
## [17] compiler_4.1.2     cli_3.6.2          hdf5r_1.3.4        shinyjs_2.1.0
## [21] plotly_4.10.4.9000 labeling_0.4.2     scales_1.3.0       digest_0.6.33
## [25] foreign_0.8-82     rmarkdown_2.18     rio_0.5.26         pkgconfig_2.0.3
## [29] htmltools_0.5.7    fastmap_1.1.1      highr_0.9          htmlwidgets_1.5.3
## [33] rlang_1.1.2        readxl_1.3.1       TTR_0.24.3         rstudioapi_0.13
## [37] quantmod_0.4.18    shiny_1.8.0        farver_2.1.0       generics_0.1.3
## [41] zoo_1.8-9          jsonlite_1.8.8     dplyr_1.0.6        zip_2.1.1
## [45] car_3.0-10         magrittr_2.0.3     Rcpp_1.0.9         munsell_0.5.0
## [49] fansi_0.5.0        abind_1.4-5        lifecycle_1.0.4    stringi_1.7.8
## [53] yaml_2.2.1         carData_3.0-4      MASS_7.3-55        grid_4.1.2
## [57] promises_1.2.0.1   forcats_0.5.1      crayon_1.4.1       EPhysMethods_0.3.1
## [61] lattice_0.20-45    haven_2.4.1        hms_1.1.0          knitr_1.33
## [65] pillar_1.6.1       ggpubr_0.4.0       ggsignif_0.6.1     glue_1.6.2
## [69] evaluate_0.23      data.table_1.14.0  vctrs_0.6.5        httpuv_1.6.1
## [73] cellranger_1.1.0   gtable_0.3.0       purrr_1.0.2        assertthat_0.2.1
## [77] ggplot2_3.5.1      xfun_0.34          openxlsx_4.2.3     mime_0.10
## [81] xtable_1.8-4       broom_0.7.6        pracma_2.3.3       rstatix_0.7.0
## [85] later_1.2.0        viridisLite_0.4.0  signal_0.7-7       tibble_3.1.2
## [89] ellipsis_0.3.2
```
