## Supplemental Material 3 for "The ERGtools2 package: A Toolset for Processing and Analysing Visual Electrophysiology Data"

### ERGtools2 - Case of use

Moritz Lindner

15 Oktober, 2024

**Session info** 22

*FilePath: "/media/moritz/DATADISK/02_Manuskripte/ERGtools2"*

#### Libraries

```
library(ERGtools2)
```

```
## Lade nötiges Paket: EPhysData

## Lade nötiges Paket: units

## udunits database from /usr/share/xml/udunits/udunits2.xml

## Registered S3 method overwritten by 'quantmod':
##   method            from
##   as.zoo.data.frame zoo
```

```
library(EPhysMethods) # containing filter functions, etc
library(cowplot)
library(ggplot2)
```

### A Standard Workflow

#### Importing data

```
Example_Exam <-
  ImportEspion(
    filename = paste0(getwd(),"/Data/ExampleSet/CR1377_22020812_MoLv1_DALAlong.TXT"),
    Import = c("Raw",
               "Measurements"),
    where = list(Channel = "ERG", Type= "Flash", Eye= "LE") # only import channels containing Flash ERG
  )
```

```
## Importing /media/moritz/DATADISK/02_Manuskripte/ERGtools2/Data/ExampleSet/CR1377_22020812_MoLv1_DALA

## Warning: Provision of the protocol is recommended and may be essential if the marker
## table is not provided or does not include markers for each step and channel of
## the recording.

## Warning: Provision of the protocol is recommended and may be essential if the marker
## table is not provided or does not include markers for each step and channel of
## the recording.
```

#### Configuring the data object for analysis

```
Example_Exam<-ClearMeasurements(Example_Exam) # to remove the Marker placement done in the manufacturer

Example_Exam<-SetStandardFunctions(Example_Exam) # set the standard functions for filtering, averaging
```

#### Visualizing and subsetting the data

```
FigStdWorkflow_A<-ggERGExam(Example_Exam)
```

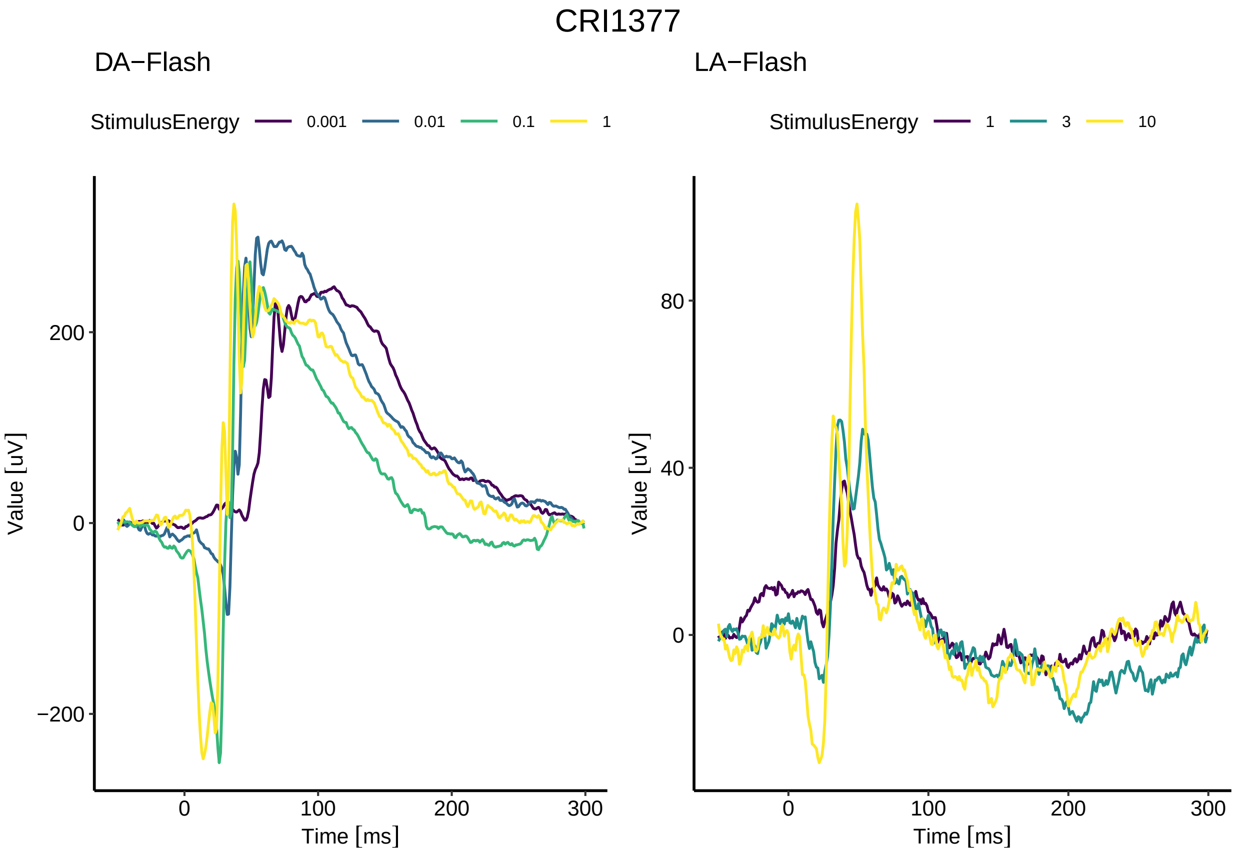

```
ggsave2("FigStdWorkflow_A.pdf", FigStdWorkflow_A, width = 7.5, height=6, units = "cm")
plot(FigStdWorkflow_A)
```

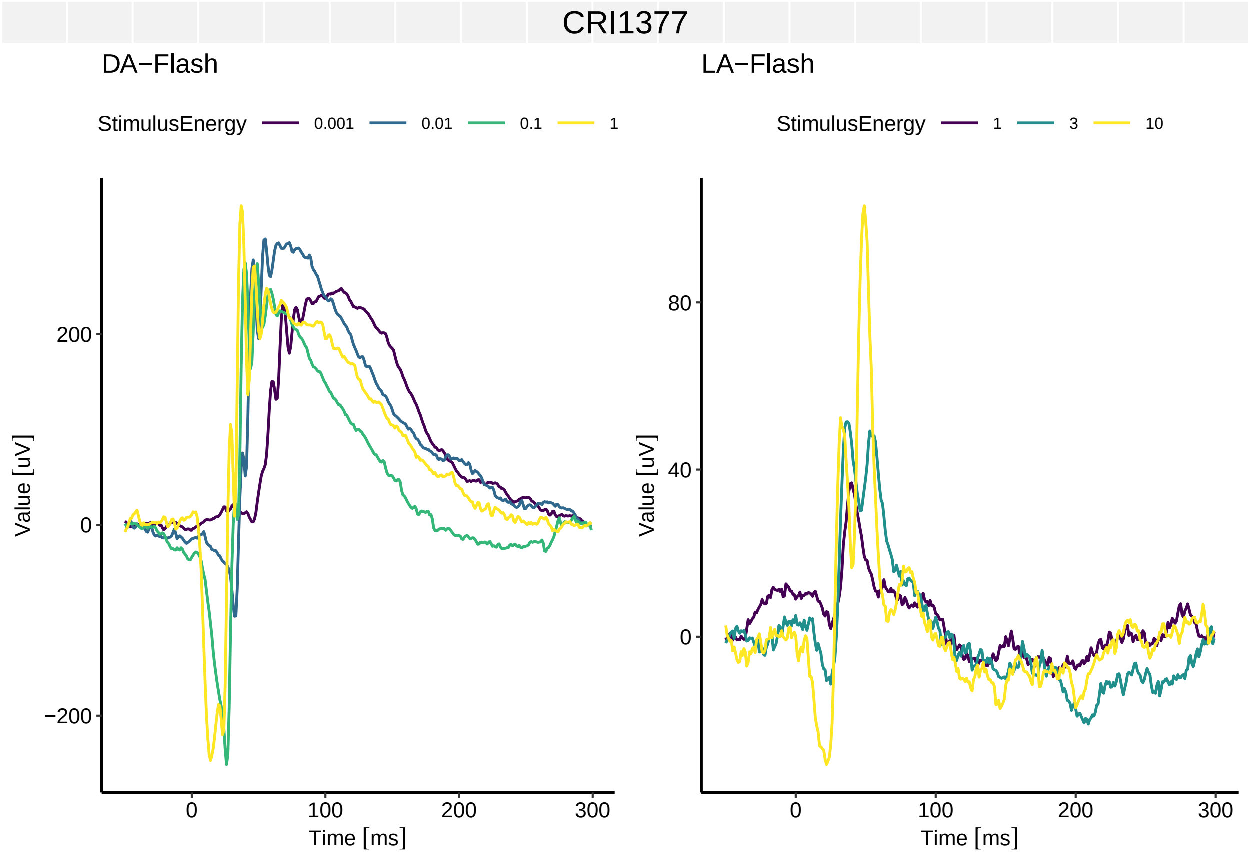

```
Example_Exam_Subset<-DropRecordings(Example_Exam, where=list(Background="LA")) # drop all recordings ac
ggERGExam(Example_Exam_Subset)
```

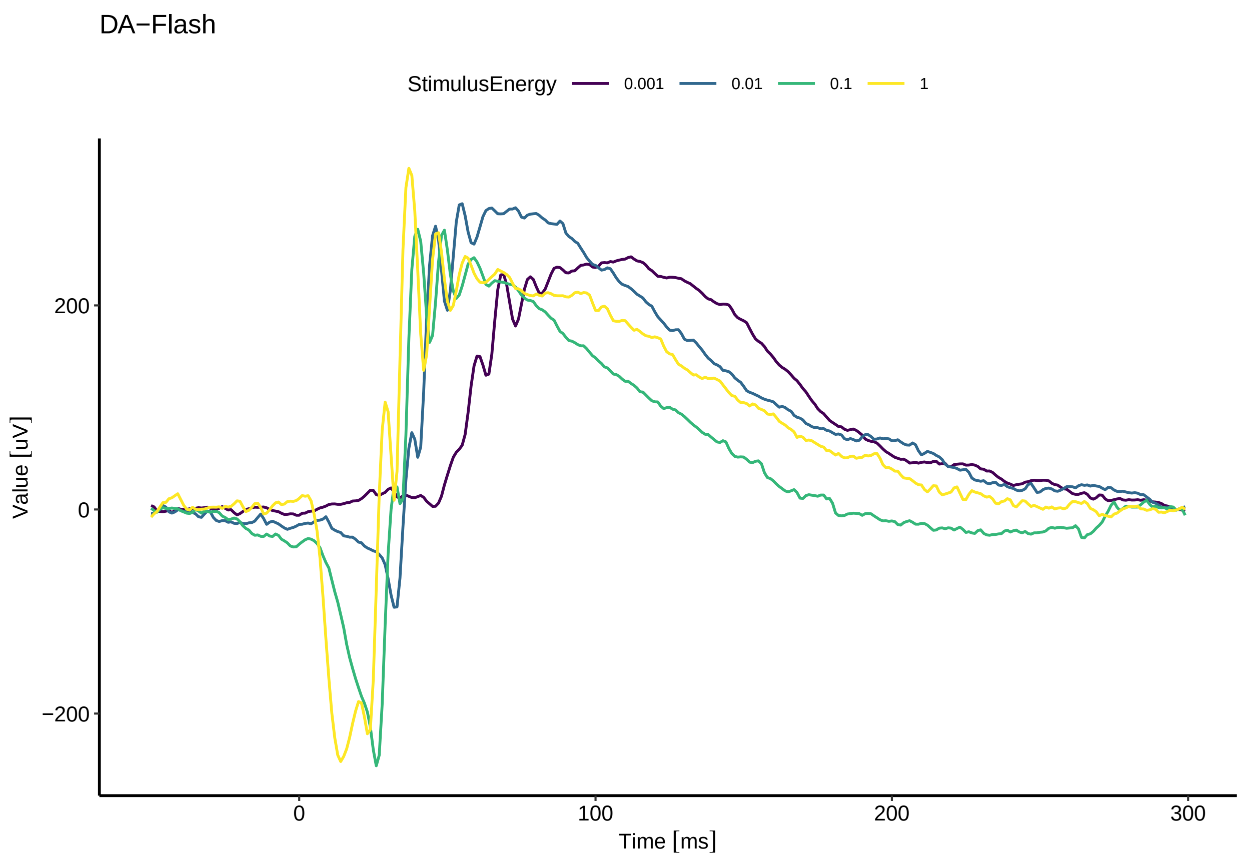

#### Placing Markers for Measurement

```
# Uncomment next line when running code manually
#Example_Exam<-exploreERGExam(Example_Exam)

Example_Exam<-AutoPlaceMarkers(Example_Exam)
ggERGExam(Example_Exam)
```

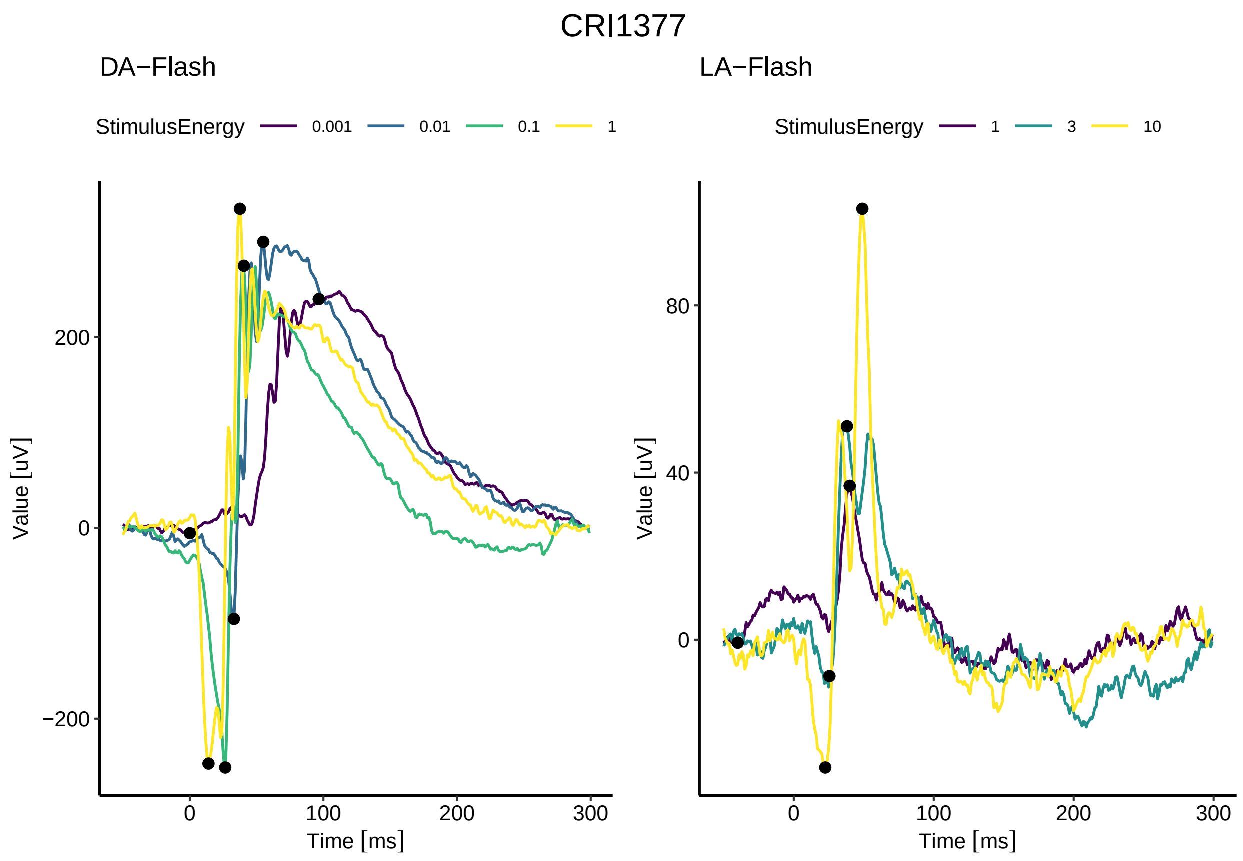

```
# Uncomment next line when running code manually
# Example_Exam<-interactiveMeasurements(Subset(Example_Exam,where=list(Background="LA")))
ggERGExam(Example_Exam)
```

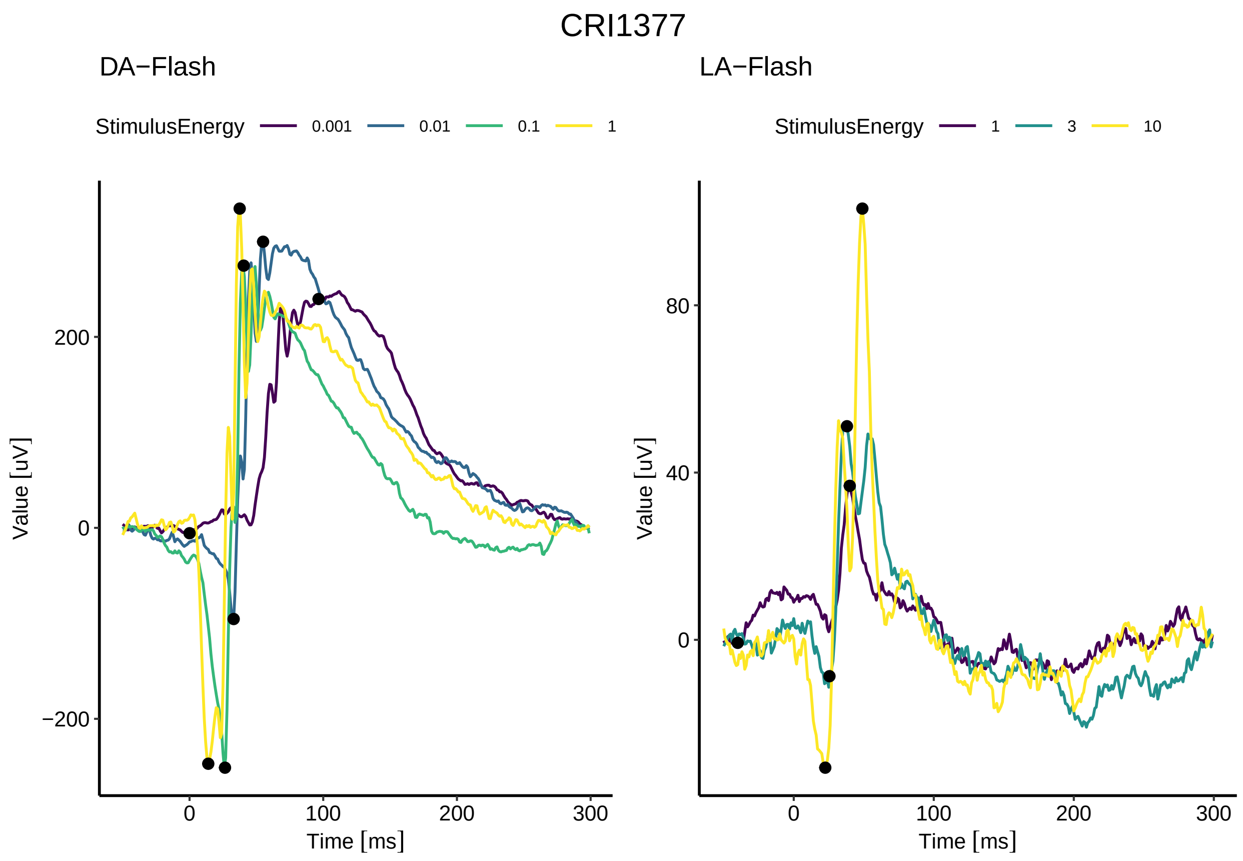

##### Summarizing and analysing a set of exams

```
dir<-paste0(getwd(),"/Data/ExampleSet/") # make a list of all files in a folder
recordings<-list.files(dir)
Exams<-list()
for (r in recordings){ # now loop through this list to import each file, the warnings we get can be ign
  Exam <-
    ImportEspion(
      filename = paste0(dir,r),
      Import = c("Raw",
                 "Measurements"),
      where = list(Channel = "ERG", Type= "Flash", Eye= "LE") # only import channels containing Flash E
    )
  Exam<-ClearMeasurements(Exam)
  Exam<-SetStandardFunctions(Exam)
  Exam<-AutoPlaceMarkers(Exam)
  Exams[[Subject(Exam)]]<-Exam
}
```

```
## Importing /media/moritz/DATADISK/02_Manuskripte/ERGtools2/Data/ExampleSet/CR1376_20220818_MoLv1_DALA
```

```
## Warning: Provision of the protocol is recommended and may be essential if the marker
## table is not provided or does not include markers for each step and channel of
```

```
## the recording.

## Warning: Provision of the protocol is recommended and may be essential if the marker
## table is not provided or does not include markers for each step and channel of
## the recording.

## Importing /media/moritz/DATADISK/02_Manuskripte/ERGtools2/Data/ExampleSet/CR1377_22020812_MoLv1_DALA[illegible]

## Warning: Provision of the protocol is recommended and may be essential if the marker
## table is not provided or does not include markers for each step and channel of
## the recording.

## Warning: Provision of the protocol is recommended and may be essential if the marker
## table is not provided or does not include markers for each step and channel of
## the recording.

## Importing /media/moritz/DATADISK/02_Manuskripte/ERGtools2/Data/ExampleSet/CR1379_20220818_MoLv1_DALA[illegible]

## Warning: Provision of the protocol is recommended and may be essential if the marker
## table is not provided or does not include markers for each step and channel of
## the recording.

## Warning: Provision of the protocol is recommended and may be essential if the marker
## table is not provided or does not include markers for each step and channel of
## the recording.

## Importing /media/moritz/DATADISK/02_Manuskripte/ERGtools2/Data/ExampleSet/CR1429_22020726_MoLv1_DALA[illegible]

## Warning: Provision of the protocol is recommended and may be essential if the marker
## table is not provided or does not include markers for each step and channel of
## the recording.

## Warning: Provision of the protocol is recommended and may be essential if the marker
## table is not provided or does not include markers for each step and channel of
## the recording.

## Warning in doTryCatch(return(expr), name, parentenv, handler): While importing
## TrialTrace for Step '1', Channel '2', Repeat '': It seems like Trials/Repeats
## have been rejected in the Espion software and were therefore not included into
## the export file. It is recommended to export all Trials and reject unwanted
## Trials inside ERGtools2.

## Warning in doTryCatch(return(expr), name, parentenv, handler): While importing
## TrialTrace for Step '2', Channel '2', Repeat '': It seems like Trials/Repeats
## have been rejected in the Espion software and were therefore not included into
## the export file. It is recommended to export all Trials and reject unwanted
## Trials inside ERGtools2.

## Importing /media/moritz/DATADISK/02_Manuskripte/ERGtools2/Data/ExampleSet/CR1459_22020926_MoLv1_DALA[illegible]

## Warning: Provision of the protocol is recommended and may be essential if the marker
## table is not provided or does not include markers for each step and channel of
## the recording.
```

```
## Warning: Provision of the protocol is recommended and may be essential if the marker
## table is not provided or does not include markers for each step and channel of
## the recording.

## Warning in doTryCatch(return(expr), name, parentenv, handler): While importing
## TrialTrace for Step '6', Channel '2', Repeat '': It seems like Trials/Repeats
## have been rejected in the Espion software and were therefore not included into
## the export file. It is recommended to export all Trials and reject unwanted
## Trials inside ERGtools2.
```

```r
ggPlotRecordings(Exams,
                 where = list(Background = "DA", Type = "Flash"))
```

```
## Running `ggPlotRecordings()`. This may take a while for long ERGExam lists.
```

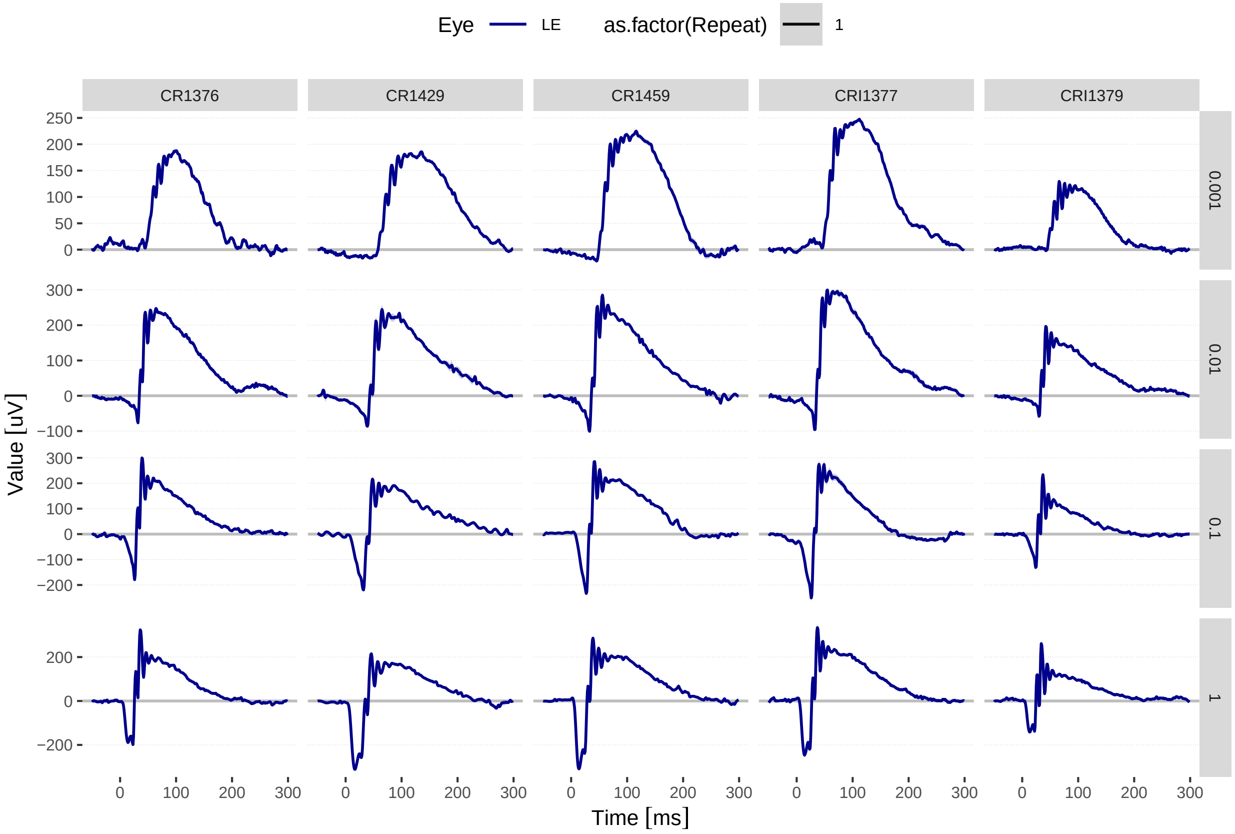

```r
ggsave2("FigStdWorkflow_2A.pdf", width = 9, height=9, units = "cm")

ggIntensitySequence(Exams,
                    where = list(Background = "DA", Type = "Flash"),
                    point.size = 1.5)
```

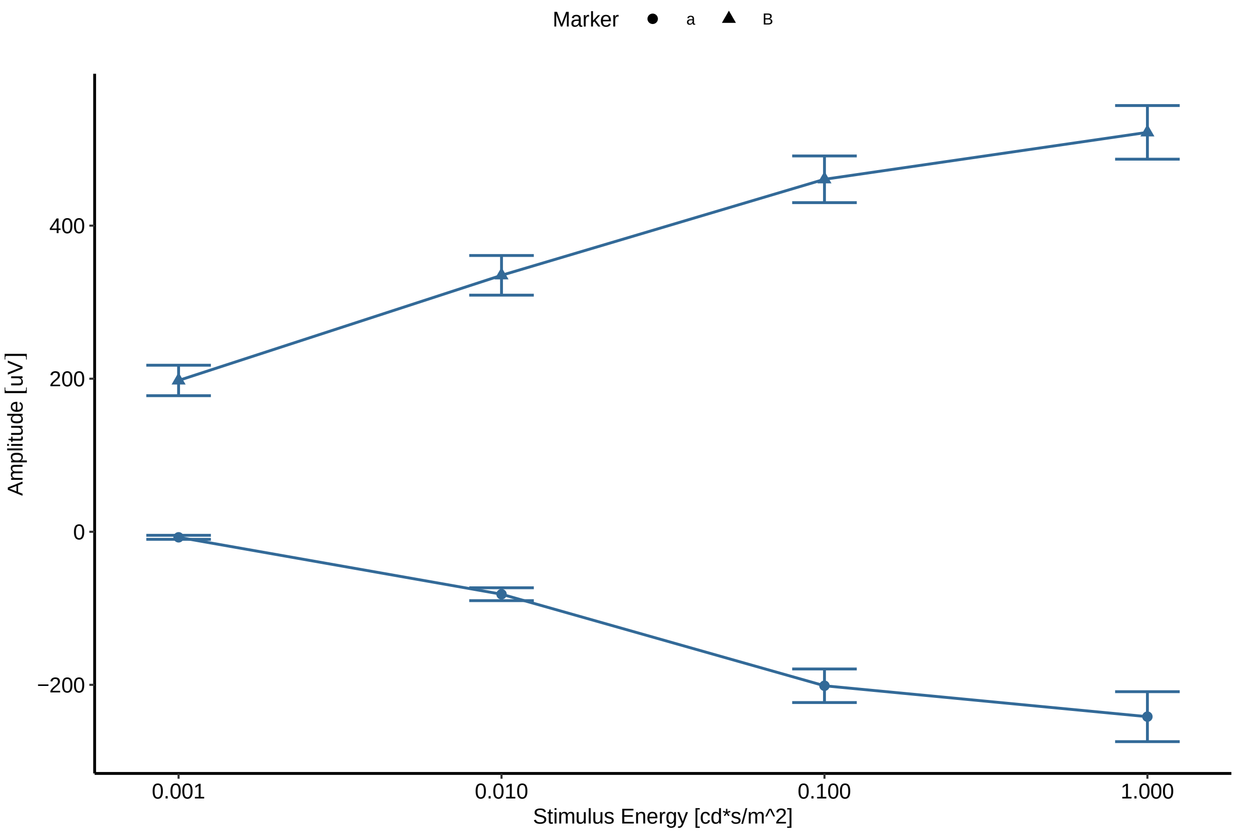

```r
ggsave2("FigStdWorkflow_2B.pdf", width = 7, height=6, units = "cm")

CollectMeasurements(Exams)
```

```
##    Step          Description Recording Channel Repeat Eye Name Relative
## 1     1     DA 0 001 cd s m         1     ERG      1  LE    a     <NA>
## 2     1     DA 0 001 cd s m         1     ERG      1  LE    B        a
## 3     2      DA 0 01 cd s m         2     ERG      1  LE    a     <NA>
## 4     2      DA 0 01 cd s m         2     ERG      1  LE    B        a
## 5     3       DA 0 1 cd s m         3     ERG      1  LE    a     <NA>
## 6     3       DA 0 1 cd s m         3     ERG      1  LE    B        a
## 7     4         DA 1 cd s m         4     ERG      1  LE    a     <NA>
## 8     4         DA 1 cd s m         4     ERG      1  LE    B        a
## 9     5   LA Single 1 0 Flash       5     ERG      1  LE    a     <NA>
## 10    5   LA Single 1 0 Flash       5     ERG      1  LE    B        a
## 11    6   LA Single 3 0 Flash       6     ERG      1  LE    a     <NA>
## 12    6   LA Single 3 0 Flash       6     ERG      1  LE    B        a
## 13    7 LA Single 10 0 Flash        7     ERG      1  LE    a     <NA>
## 14    7 LA Single 10 0 Flash        7     ERG      1  LE    B        a
## 15    1     DA 0 001 cd s m         1     ERG      1  LE    a     <NA>
## 16    1     DA 0 001 cd s m         1     ERG      1  LE    B        a
## 17    2      DA 0 01 cd s m         2     ERG      1  LE    a     <NA>
## 18    2      DA 0 01 cd s m         2     ERG      1  LE    B        a
## 19    3       DA 0 1 cd s m         3     ERG      1  LE    a     <NA>
## 20    3       DA 0 1 cd s m         3     ERG      1  LE    B        a
## 21    4         DA 1 cd s m         4     ERG      1  LE    a     <NA>
```

```
## 22    4         DA 1 cd s m          4     ERG      1  LE    B        a
## 23    5  LA Single 1 0 Flash         5     ERG      1  LE    a     <NA>
## 24    5  LA Single 1 0 Flash         5     ERG      1  LE    B        a
## 25    6  LA Single 3 0 Flash         6     ERG      1  LE    a     <NA>
## 26    6  LA Single 3 0 Flash         6     ERG      1  LE    B        a
## 27    7 LA Single 10 0 Flash         7     ERG      1  LE    a     <NA>
## 28    7 LA Single 10 0 Flash         7     ERG      1  LE    B        a
## 29    1     DA 0 001 cd s m          1     ERG      1  LE    a     <NA>
## 30    1     DA 0 001 cd s m          1     ERG      1  LE    B        a
## 31    2      DA 0 01 cd s m          2     ERG      1  LE    a     <NA>
## 32    2      DA 0 01 cd s m          2     ERG      1  LE    B        a
## 33    3       DA 0 1 cd s m          3     ERG      1  LE    a     <NA>
## 34    3       DA 0 1 cd s m          3     ERG      1  LE    B        a
## 35    4         DA 1 cd s m          4     ERG      1  LE    a     <NA>
## 36    4         DA 1 cd s m          4     ERG      1  LE    B        a
## 37    5  LA Single 1 0 Flash         5     ERG      1  LE    a     <NA>
## 38    5  LA Single 1 0 Flash         5     ERG      1  LE    B        a
## 39    6  LA Single 3 0 Flash         6     ERG      1  LE    a     <NA>
## 40    6  LA Single 3 0 Flash         6     ERG      1  LE    B        a
## 41    7 LA Single 10 0 Flash         7     ERG      1  LE    a     <NA>
## 42    7 LA Single 10 0 Flash         7     ERG      1  LE    B        a
## 43    1     DA 0 001 cd s m          1     ERG      1  LE    a     <NA>
## 44    1     DA 0 001 cd s m          1     ERG      1  LE    B        a
## 45    2      DA 0 01 cd s m          2     ERG      1  LE    a     <NA>
## 46    2      DA 0 01 cd s m          2     ERG      1  LE    B        a
## 47    3       DA 0 1 cd s m          3     ERG      1  LE    a     <NA>
## 48    3       DA 0 1 cd s m          3     ERG      1  LE    B        a
## 49    4         DA 1 cd s m          4     ERG      1  LE    a     <NA>
## 50    4         DA 1 cd s m          4     ERG      1  LE    B        a
## 51    5  LA Single 1 0 Flash         5     ERG      1  LE    a     <NA>
## 52    5  LA Single 1 0 Flash         5     ERG      1  LE    B        a
## 53    6  LA Single 3 0 Flash         6     ERG      1  LE    a     <NA>
## 54    6  LA Single 3 0 Flash         6     ERG      1  LE    B        a
## 55    7 LA Single 10 0 Flash         7     ERG      1  LE    a     <NA>
## 56    7 LA Single 10 0 Flash         7     ERG      1  LE    B        a
## 57    1     DA 0 001 cd s m          1     ERG      1  LE    a     <NA>
## 58    1     DA 0 001 cd s m          1     ERG      1  LE    B        a
## 59    2      DA 0 01 cd s m          2     ERG      1  LE    a     <NA>
## 60    2      DA 0 01 cd s m          2     ERG      1  LE    B        a
## 61    3       DA 0 1 cd s m          3     ERG      1  LE    a     <NA>
## 62    3       DA 0 1 cd s m          3     ERG      1  LE    B        a
## 63    4         DA 1 cd s m          4     ERG      1  LE    a     <NA>
## 64    4         DA 1 cd s m          4     ERG      1  LE    B        a
## 65    5  LA Single 1 0 Flash         5     ERG      1  LE    a     <NA>
## 66    5  LA Single 1 0 Flash         5     ERG      1  LE    B        a
## 67    6  LA Single 3 0 Flash         6     ERG      1  LE    a     <NA>
## 68    6  LA Single 3 0 Flash         6     ERG      1  LE    B        a
## 69    7 LA Single 10 0 Flash         7     ERG      1  LE    a     <NA>
## 70    7 LA Single 10 0 Flash         7     ERG      1  LE    B        a
##           Time           Voltage Subject Group            ExamDate
## 1   0.0310 [s]   -2.0383069 [uV]  CR1376     0 2022-08-16 10:53:41
## 2   0.0955 [s]  186.4152317 [uV]  CR1376     0 2022-08-16 10:53:41
## 3   0.0325 [s]  -70.7528330 [uV]  CR1376     0 2022-08-16 10:53:41
## 4   0.0645 [s]  318.0240514 [uV]  CR1376     0 2022-08-16 10:53:41
```

```
## 5   0.0265 [s] -175.4332551 [uV]  CR1376     0 2022-08-16 10:53:41
## 6   0.0400 [s]  472.5466852 [uV]  CR1376     0 2022-08-16 10:53:41
## 7   0.0230 [s] -198.6900201 [uV]  CR1376     0 2022-08-16 10:53:41
## 8   0.0370 [s]  516.8584293 [uV]  CR1376     0 2022-08-16 10:53:41
## 9   0.0295 [s]  -17.8737140 [uV]  CR1376     0 2022-08-16 10:53:41
## 10  0.0400 [s]   33.6568481 [uV]  CR1376     0 2022-08-16 10:53:41
## 11  0.0255 [s]   -5.5195789 [uV]  CR1376     0 2022-08-16 10:53:41
## 12  0.0530 [s]   58.5163887 [uV]  CR1376     0 2022-08-16 10:53:41
## 13  0.0235 [s]  -20.0560929 [uV]  CR1376     0 2022-08-16 10:53:41
## 14  0.0485 [s]  116.2060109 [uV]  CR1376     0 2022-08-16 10:53:41
## 15  0.0000 [s]   -5.6327413 [uV] CRI1377     0 2022-08-12 11:54:57
## 16  0.0965 [s]  245.5506086 [uV] CRI1377     0 2022-08-12 11:54:57
## 17  0.0330 [s]  -95.4677544 [uV] CRI1377     0 2022-08-12 11:54:57
## 18  0.0550 [s]  395.1253381 [uV] CRI1377     0 2022-08-12 11:54:57
## 19  0.0265 [s] -250.0754019 [uV] CRI1377     0 2022-08-12 11:54:57
## 20  0.0405 [s]  521.0791908 [uV] CRI1377     0 2022-08-12 11:54:57
## 21  0.0140 [s] -247.0355817 [uV] CRI1377     0 2022-08-12 11:54:57
## 22  0.0375 [s]  580.3660621 [uV] CRI1377     0 2022-08-12 11:54:57
## 23 -0.0400 [s]   -0.7195593 [uV] CRI1377     0 2022-08-12 11:54:57
## 24  0.0400 [s]   37.5468171 [uV] CRI1377     0 2022-08-12 11:54:57
## 25  0.0255 [s]  -10.4145864 [uV] CRI1377     0 2022-08-12 11:54:57
## 26  0.0380 [s]   61.4911811 [uV] CRI1377     0 2022-08-12 11:54:57
## 27  0.0225 [s]  -30.0617504 [uV] CRI1377     0 2022-08-12 11:54:57
## 28  0.0490 [s]  133.1896458 [uV] CRI1377     0 2022-08-12 11:54:57
## 29  0.0430 [s]   -1.6935731 [uV] CRI1379     0 2022-08-16 12:04:31
## 30  0.0665 [s]  130.9577513 [uV] CRI1379     0 2022-08-16 12:04:31
## 31  0.0310 [s]  -54.8687636 [uV] CRI1379     0 2022-08-16 12:04:31
## 32  0.0430 [s]  250.0899613 [uV] CRI1379     0 2022-08-16 12:04:31
## 33  0.0250 [s] -128.7135035 [uV] CRI1379     0 2022-08-16 12:04:31
## 34  0.0375 [s]  356.0739975 [uV] CRI1379     0 2022-08-16 12:04:31
## 35  0.0135 [s] -141.6833086 [uV] CRI1379     0 2022-08-16 12:04:31
## 36  0.0350 [s]  394.4244023 [uV] CRI1379     0 2022-08-16 12:04:31
## 37  0.0270 [s]   -9.4809895 [uV] CRI1379     0 2022-08-16 12:04:31
## 38  0.0390 [s]   15.9727114 [uV] CRI1379     0 2022-08-16 12:04:31
## 39  0.0195 [s]   -4.6232949 [uV] CRI1379     0 2022-08-16 12:04:31
## 40  0.0365 [s]   33.6790807 [uV] CRI1379     0 2022-08-16 12:04:31
## 41  0.0180 [s]  -11.3302291 [uV] CRI1379     0 2022-08-16 12:04:31
## 42  0.0495 [s]   68.7465564 [uV] CRI1379     0 2022-08-16 12:04:31
## 43  0.0025 [s]  -14.4315070 [uV]  CR1429     0 2022-09-14 11:01:00
## 44  0.1055 [s]  195.5306039 [uV]  CR1429     0 2022-09-14 11:01:00
## 45  0.0390 [s]  -86.2645861 [uV]  CR1429     0 2022-09-14 11:01:00
## 46  0.0655 [s]  329.6008365 [uV]  CR1429     0 2022-09-14 11:01:00
## 47  0.0320 [s] -219.5709215 [uV]  CR1429     0 2022-09-14 11:01:00
## 48  0.0485 [s]  435.4916330 [uV]  CR1429     0 2022-09-14 11:01:00
## 49  0.0170 [s] -311.2914575 [uV]  CR1429     0 2022-09-14 11:01:00
## 50  0.0460 [s]  526.3570672 [uV]  CR1429     0 2022-09-14 11:01:00
## 51  0.0320 [s]  -15.7517460 [uV]  CR1429     0 2022-09-14 11:01:00
## 52  0.0580 [s]   53.1868419 [uV]  CR1429     0 2022-09-14 11:01:00
## 53  0.0315 [s]  -20.1612616 [uV]  CR1429     0 2022-09-14 11:01:00
## 54  0.0555 [s]   78.3343671 [uV]  CR1429     0 2022-09-14 11:01:00
## 55  0.0265 [s]  -29.7262137 [uV]  CR1429     0 2022-09-14 11:01:00
## 56  0.0650 [s]  154.0166277 [uV]  CR1429     0 2022-09-14 11:01:00
## 57  0.0195 [s]  -12.4397352 [uV]  CR1459     0 2022-08-26 13:01:15
## 58  0.0970 [s]  230.1339388 [uV]  CR1459     0 2022-08-26 13:01:15
```

```
## 59  0.0330 [s] -100.7445542 [uV]  CR1459     0 2022-08-26 13:01:15
## 60  0.0565 [s]  382.6203862 [uV]  CR1459     0 2022-08-26 13:01:15
## 61  0.0275 [s] -232.3591804 [uV]  CR1459     0 2022-08-26 13:01:15
## 62  0.0420 [s]  517.6675889 [uV]  CR1459     0 2022-08-26 13:01:15
## 63  0.0140 [s] -309.3362752 [uV]  CR1459     0 2022-08-26 13:01:15
## 64  0.0395 [s]  591.4785377 [uV]  CR1459     0 2022-08-26 13:01:15
## 65  0.0280 [s]  -12.0952291 [uV]  CR1459     0 2022-08-26 13:01:15
## 66  0.0465 [s]   40.1823050 [uV]  CR1459     0 2022-08-26 13:01:15
## 67  0.0265 [s]  -17.6110215 [uV]  CR1459     0 2022-08-26 13:01:15
## 68  0.0410 [s]   66.2415868 [uV]  CR1459     0 2022-08-26 13:01:15
## 69  0.0215 [s]  -25.8541336 [uV]  CR1459     0 2022-08-26 13:01:15
## 70  0.0560 [s]  148.5445519 [uV]  CR1459     0 2022-08-26 13:01:15
##    StimulusEnergy Background  Type
## 1           1e-03         DA Flash
## 2           1e-03         DA Flash
## 3           1e-02         DA Flash
## 4           1e-02         DA Flash
## 5           1e-01         DA Flash
## 6           1e-01         DA Flash
## 7           1e+00         DA Flash
## 8           1e+00         DA Flash
## 9           1e+00         LA Flash
## 10          1e+00         LA Flash
## 11          3e+00         LA Flash
## 12          3e+00         LA Flash
## 13          1e+01         LA Flash
## 14          1e+01         LA Flash
## 15          1e-03         DA Flash
## 16          1e-03         DA Flash
## 17          1e-02         DA Flash
## 18          1e-02         DA Flash
## 19          1e-01         DA Flash
## 20          1e-01         DA Flash
## 21          1e+00         DA Flash
## 22          1e+00         DA Flash
## 23          1e+00         LA Flash
## 24          1e+00         LA Flash
## 25          3e+00         LA Flash
## 26          3e+00         LA Flash
## 27          1e+01         LA Flash
## 28          1e+01         LA Flash
## 29          1e-03         DA Flash
## 30          1e-03         DA Flash
## 31          1e-02         DA Flash
## 32          1e-02         DA Flash
## 33          1e-01         DA Flash
## 34          1e-01         DA Flash
## 35          1e+00         DA Flash
## 36          1e+00         DA Flash
## 37          1e+00         LA Flash
## 38          1e+00         LA Flash
## 39          3e+00         LA Flash
## 40          3e+00         LA Flash
## 41          1e+01         LA Flash
```

```
## 42          1e+01         LA Flash
## 43          1e-03         DA Flash
## 44          1e-03         DA Flash
## 45          1e-02         DA Flash
## 46          1e-02         DA Flash
## 47          1e-01         DA Flash
## 48          1e-01         DA Flash
## 49          1e+00         DA Flash
## 50          1e+00         DA Flash
## 51          1e+00         LA Flash
## 52          1e+00         LA Flash
## 53          3e+00         LA Flash
## 54          3e+00         LA Flash
## 55          1e+01         LA Flash
## 56          1e+01         LA Flash
## 57          1e-03         DA Flash
## 58          1e-03         DA Flash
## 59          1e-02         DA Flash
## 60          1e-02         DA Flash
## 61          1e-01         DA Flash
## 62          1e-01         DA Flash
## 63          1e+00         DA Flash
## 64          1e+00         DA Flash
## 65          1e+00         LA Flash
## 66          1e+00         LA Flash
## 67          3e+00         LA Flash
## 68          3e+00         LA Flash
## 69          1e+01         LA Flash
## 70          1e+01         LA Flash
```

### Special use cases

#### Automatic rejection of outlier recordings

```
Noisy_Exam<-Subset(Exams[[5]],where=list(StimulusEnergy=1,Background="LA"))
Noisy_Exam<-ClearMeasurements(Noisy_Exam)

# now replace the Rejection function for each Step/Recording in the object (in this case, there is only
Rejected(Noisy_Exam, where=list(Step=5))<-autoreject.by.distance
table(Rejected(Noisy_Exam[[1]]))
```

```
##
## FALSE  TRUE
##    14     6
```

```
Noise_dist_left <-
  ggERGTrace(Noisy_Exam, where = list(StimulusEnergy = 1, Background = "LA")) + theme(legend.position =
Noise_dist_left # Print all trials with the rejected highlighted
```

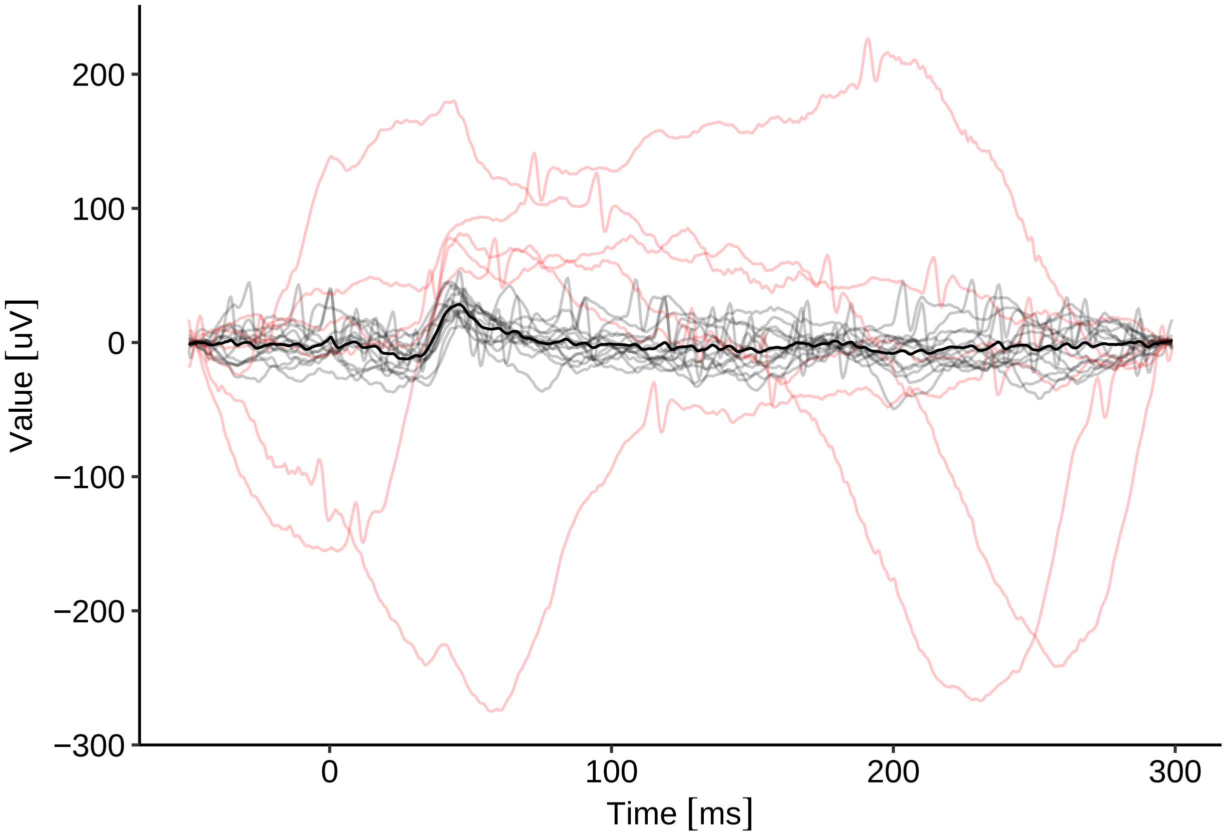

```
Noise_dist_right <-
  ggERGTrace(Noisy_Exam,
             where = list(StimulusEnergy = 1, Background = "LA"),
             Raw = F) + theme(legend.position = "none")
Noise_dist_right # Print averaged recording with the rejected highlighted
```

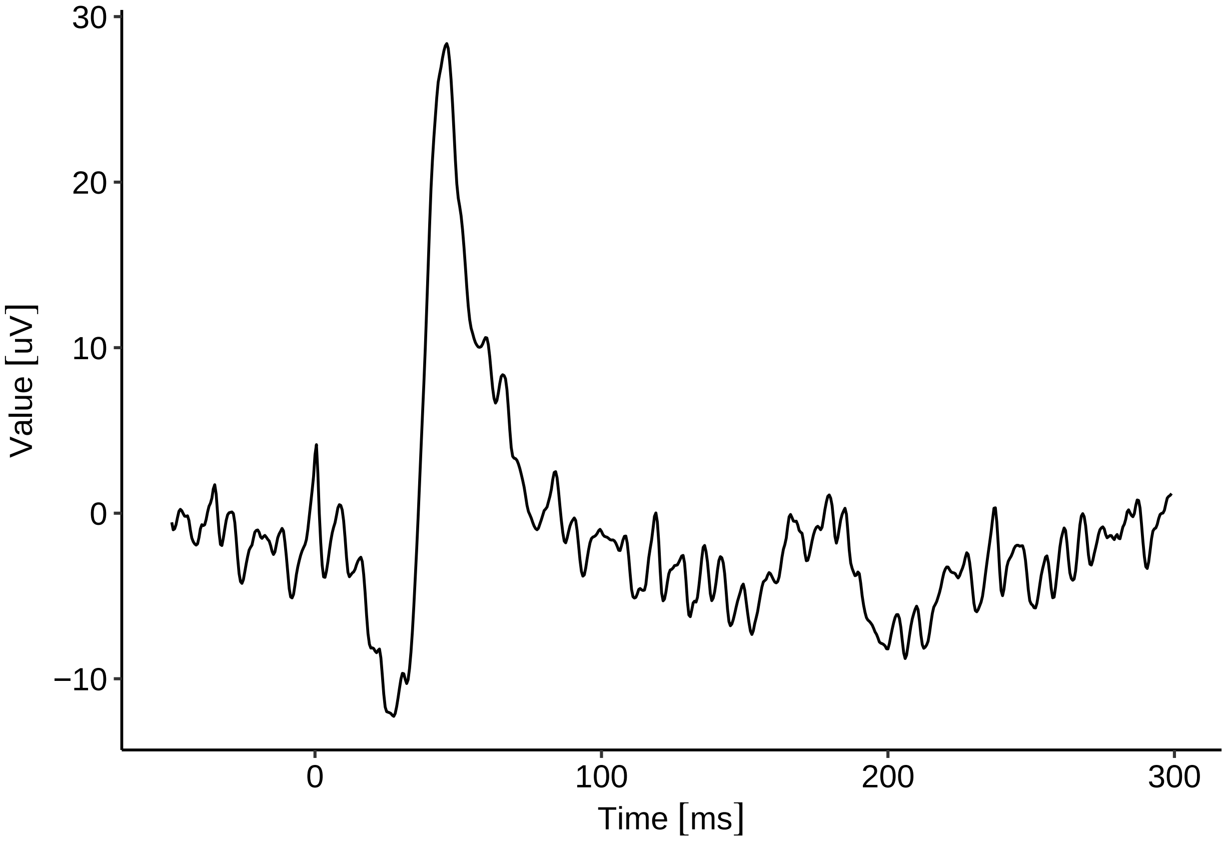

```
Rejected(Noisy_Exam, where = list(Step = 5)) <- function(x) {
  logical(dim(x)[2])
} # set all traces to be included, i.e. the function for rejection returns false for all trials.

table(Rejected(Noisy_Exam[[1]]))
```

```
##
## FALSE
##    20
```

```
Noise_thresh_middle <-
  ggERGTrace(Noisy_Exam,
             where = list(StimulusEnergy = 1, Background = "LA"),
             Raw = F) + theme(legend.position = "none")
Noise_thresh_middle
```

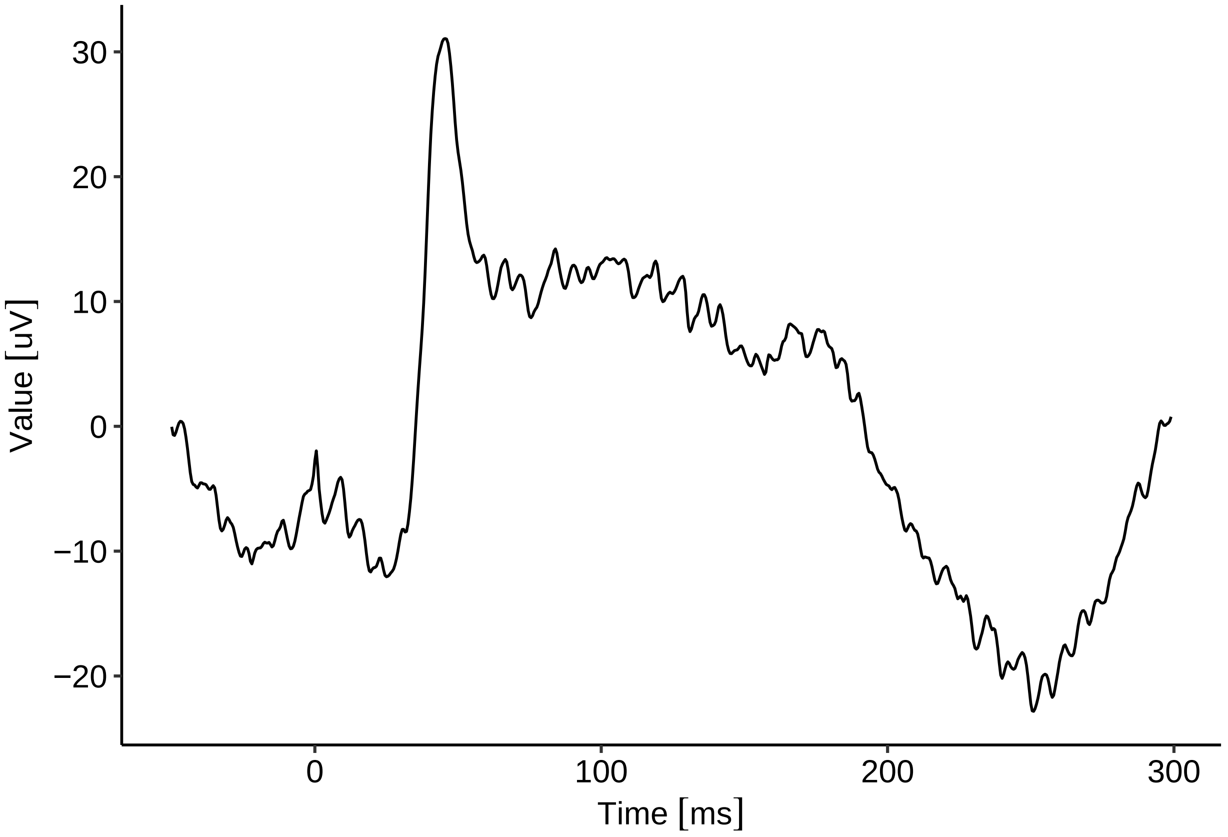

```
plot_grid(Noise_dist_left,Noise_thresh_middle,Noise_dist_right,nrow = 1)
```

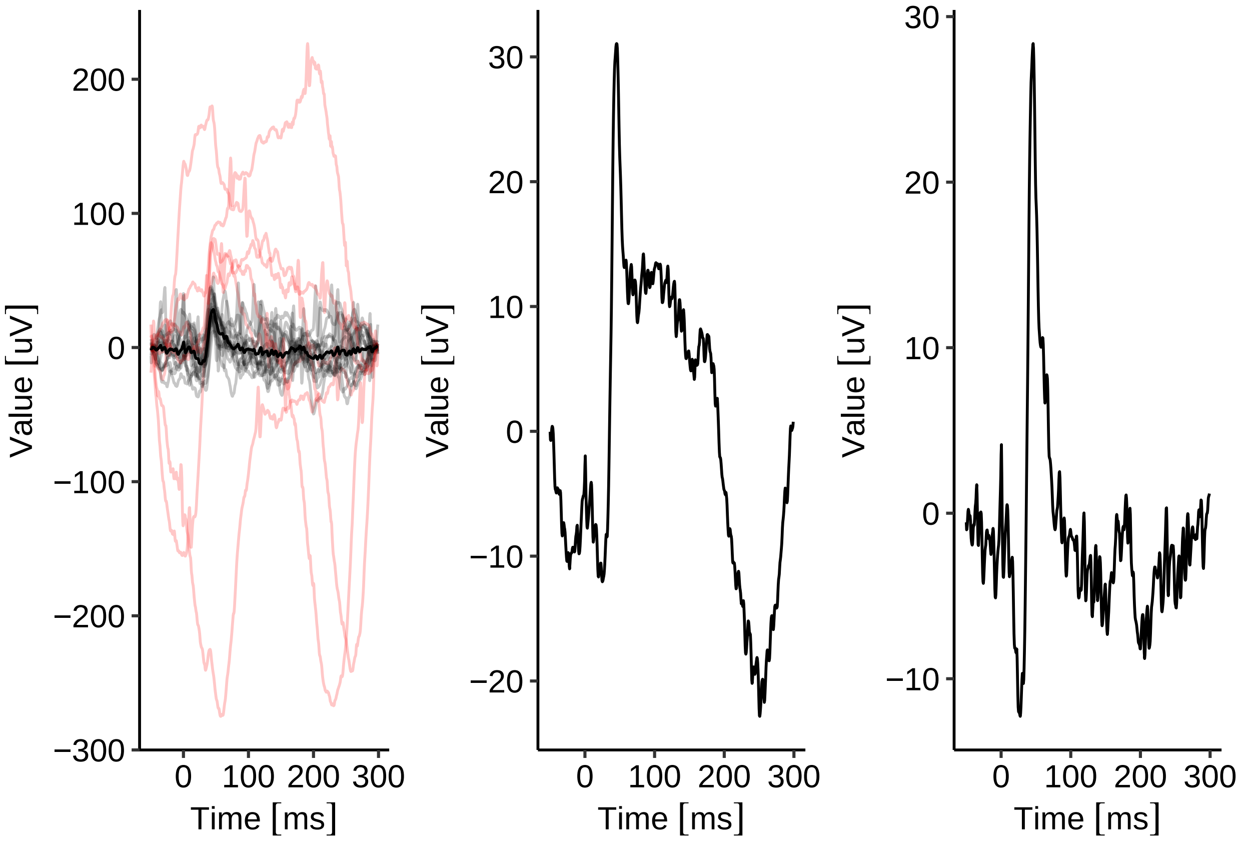

```
ggsave2("FigNOISE.PDF", width = 16, height=5, units = "cm")
```

##### Extracting and analysing oscillatory potentials

```
Exam_forOPs<-Subset(Exam,where=list(StimulusEnergy=0.1,Background="DA"))
Exam_forOPs<-ClearMeasurements(Exam_forOPs)
FigOP_A <-
  ggERGTrace(Exam_forOPs, where = list(StimulusEnergy = 0.1, Background = "DA")) + theme(legend.positio
FigOP_A
```

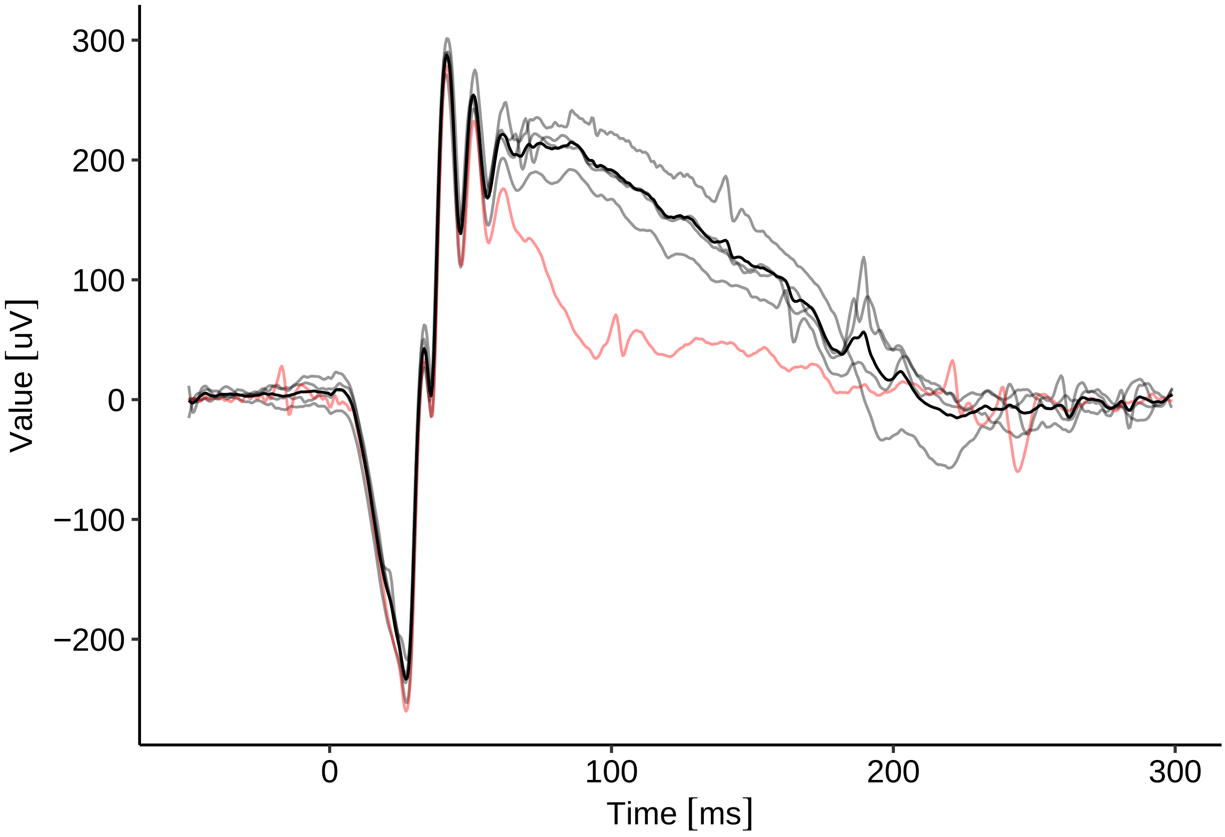

```
# filter and plot OPs
cutoff <-
  freq.to.w(c(75, 300), samp.freq = 2000) # convert to digital angular frequencies, we need to know the

FilterFunction(Exam_forOPs, where = list(Step = 3)) <-
  function(x) {
    # we set the new filter
    eval(substitute(
      filter.bandpass(x, low, high),
      list(low = cutoff[1], high = cutoff[2])
    ))
  }

FigOP_B <-
  ggERGTrace(Exam_forOPs, where = list(StimulusEnergy = 0.1, Background = "DA")) + theme(legend.positio
FigOP_B
```

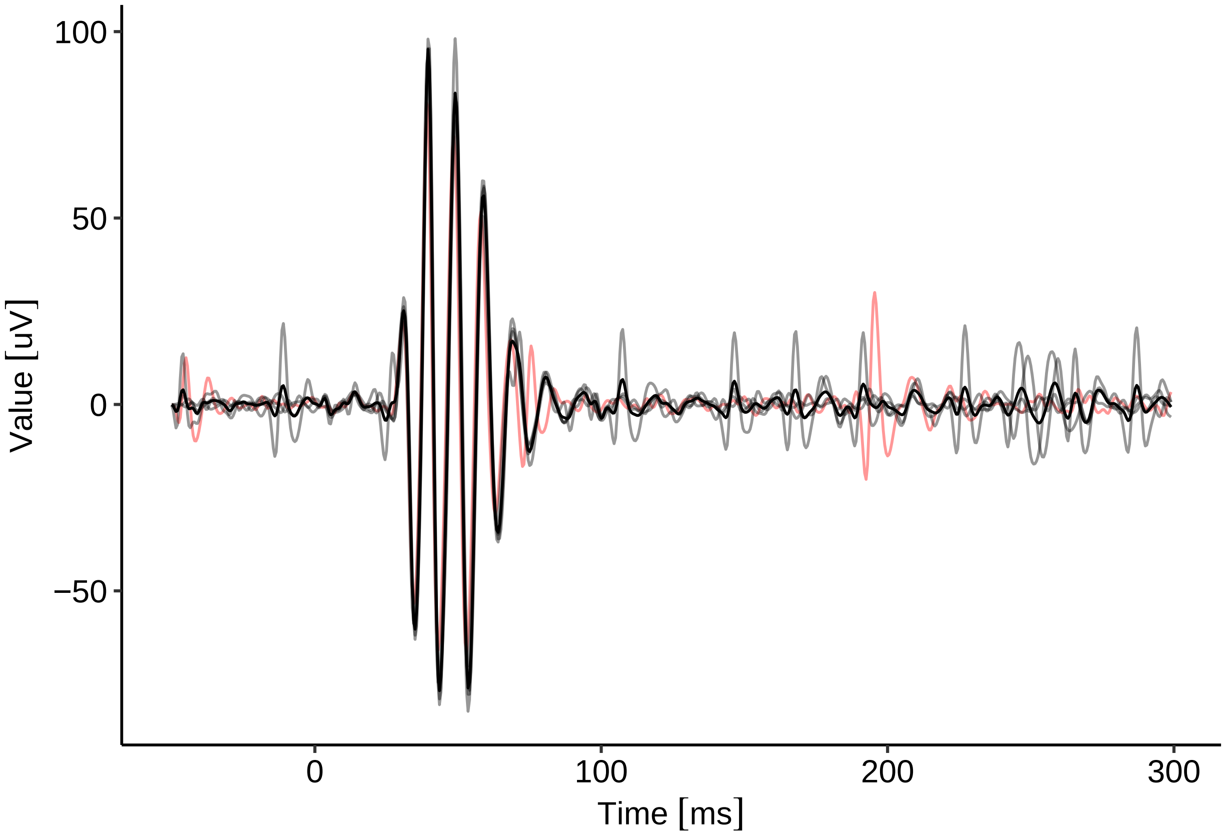

```
# calculate spectral power

cutoff <-
  freq.to.w(c(0.5, 300), samp.freq = 2000) # convert to digital angular frequencies, we need to know th

FilterFunction(Exam_forOPs, where = list(Step = 3)) <- function(x) { # we set the new filter
  filter.bandpass(x, cutoff[1], cutoff[2])
}
PSD <- lapply(Exam_forOPs, function(x) {
  psd<-PSD(Subset(x,Time = as_units(c(0,230),"ms")))
  psd<-Subset(psd,Time = as_units(c(55,250),"Hz"))
})
```

```
## Data is subsetted by time, thus resetting filter function.
## Data is subsetted by time, thus resetting filter function.
```

the PSD object now contains data of the unit nV2/Hz. The ggERGTrace method will expect a unit convertible to V, which is not preset here. So we have to use the lower-level function ggEPhysData.

```
FigOP_C <- ggEPhysData(PSD[[1]]) + theme(legend.position = "none")
FigOP_C
```

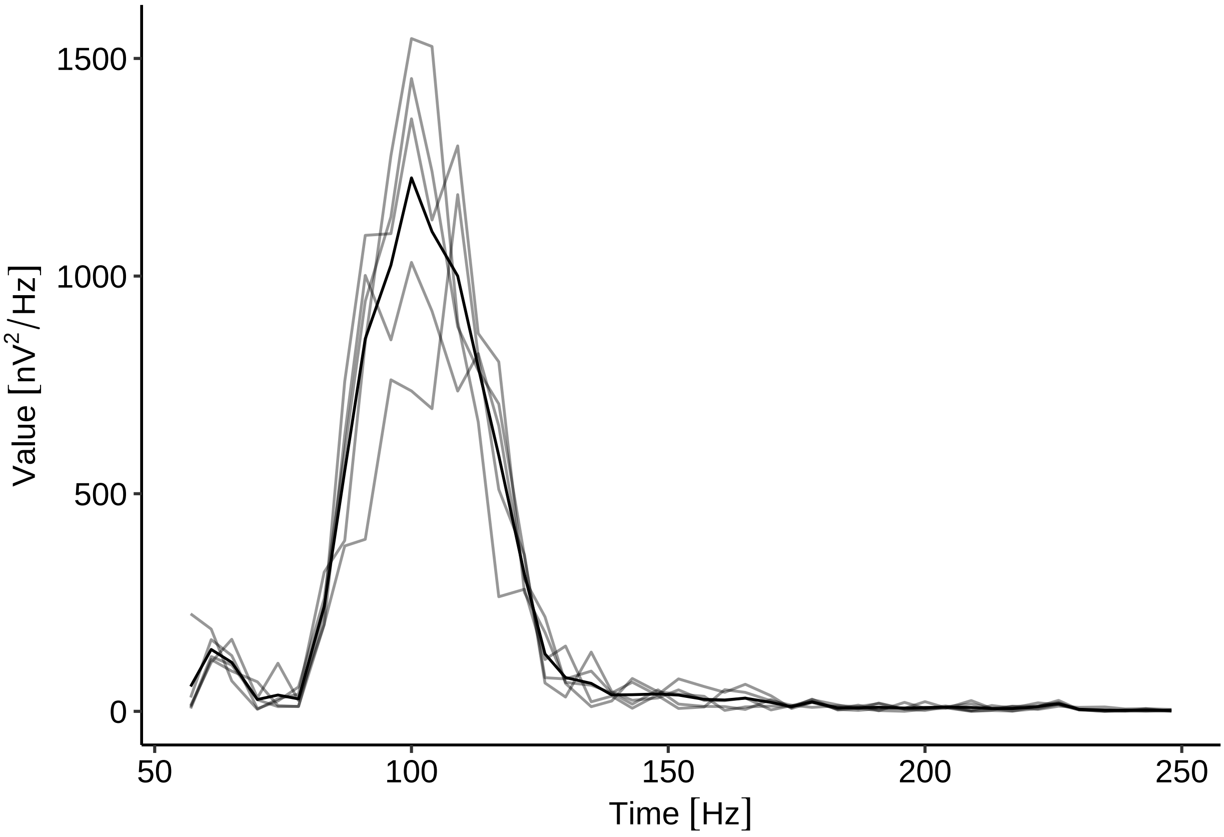

```r
# extract the values to perform further analyses

out <- lapply(PSD, function(y) {
  df_LA<-as.data.frame(y, Raw=F) # get averaged data
  df_LA
}, ReturnEPhysSet = F)

out<-out[[1]] # get the first (and in this case only) recording in the data set

# find the peak frequency
out$Time[which.max(out$Value)]
```

```
## 100 [Hz]
```

```r
# make composite figure

plot_grid(FigOP_A,FigOP_B,FigOP_C,labels=c("A","B","C"),nrow=1)
```

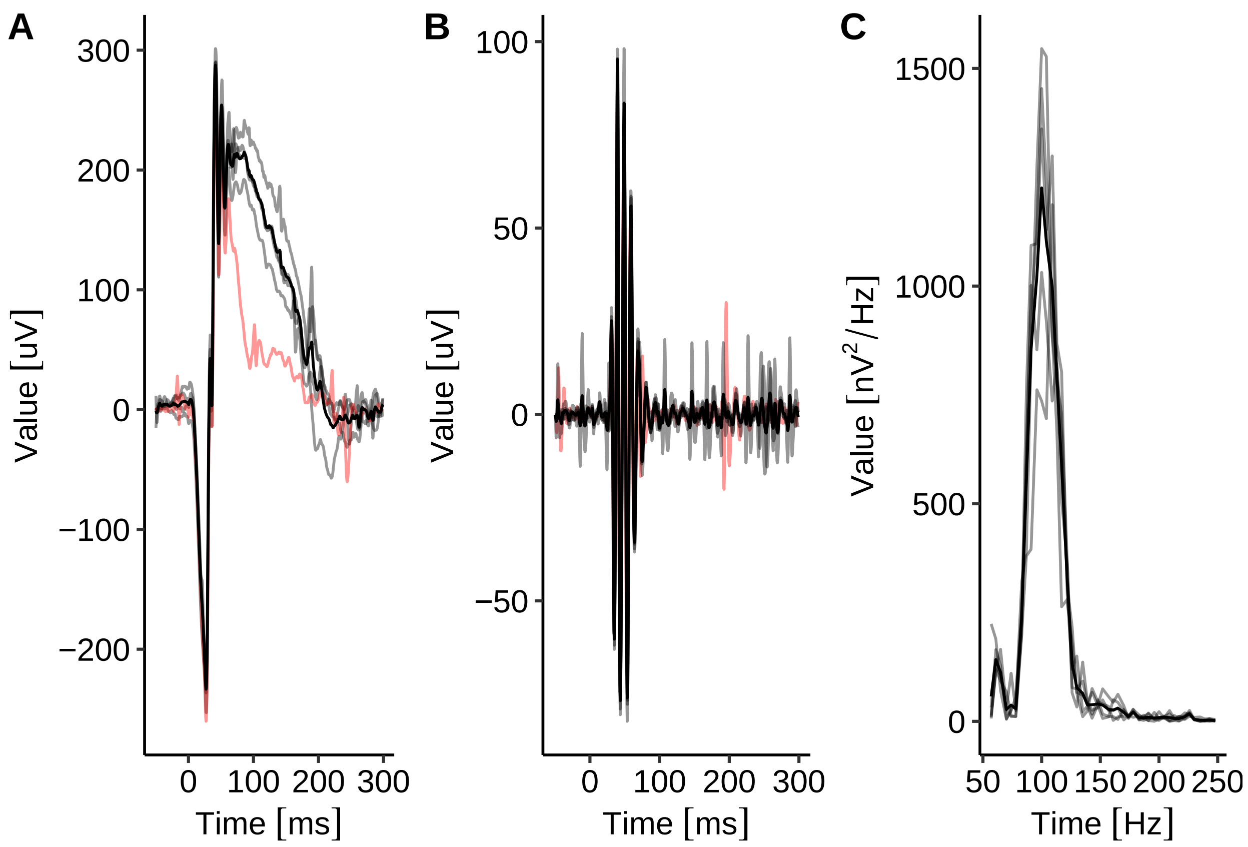

```
ggsave2("FigOP.PDF", width = 16, height=5, units = "cm")
```

#### Session info

```
sessionInfo()
```

```
## R version 4.1.2 (2021-11-01)
## Platform: x86_64-pc-linux-gnu (64-bit)
## Running under: Ubuntu 22.04.5 LTS
##
## Matrix products: default
## BLAS:   /usr/lib/x86_64-linux-gnu/blas/libblas.so.3.10.0
## LAPACK: /usr/lib/x86_64-linux-gnu/lapack/liblapack.so.3.10.0
##
## locale:
##  [1] LC_CTYPE=de_DE.UTF-8       LC_NUMERIC=C
##  [3] LC_TIME=de_DE.UTF-8        LC_COLLATE=de_DE.UTF-8
##  [5] LC_MONETARY=de_DE.UTF-8    LC_MESSAGES=de_DE.UTF-8
##  [7] LC_PAPER=de_DE.UTF-8       LC_NAME=C
##  [9] LC_ADDRESS=C               LC_TELEPHONE=C
## [11] LC_MEASUREMENT=de_DE.UTF-8 LC_IDENTIFICATION=C
##
```

```
## attached base packages:
## [1] stats     graphics  grDevices utils     datasets  methods   base
##
## other attached packages:
## [1] ggplot2_3.5.1      cowplot_1.1.1      EPhysMethods_0.3.1 ERGtools2_0.8.0
## [5] EPhysData_0.9.7    units_0.8-0
##
## loaded via a namespace (and not attached):
##  [1] xts_0.12.1         bit64_4.0.5        httr_1.4.2         tools_4.1.2
##  [5] backports_1.2.1    utf8_1.2.1         R6_2.5.0           DT_0.19
##  [9] DBI_1.1.1          lazyeval_0.2.2     colorspace_2.0-1   withr_2.5.2
## [13] tidyselect_1.1.1   gridExtra_2.3      bit_4.0.4          curl_4.3.1
## [17] compiler_4.1.2     textshaping_0.3.6  cli_3.6.2          hdf5r_1.3.4
## [21] shinyjs_2.1.0      plotly_4.10.4.9000 labeling_0.4.2     scales_1.3.0
## [25] systemfonts_1.0.4  stringr_1.5.1      digest_0.6.33      foreign_0.8-82
## [29] rmarkdown_2.18     rio_0.5.26         pkgconfig_2.0.3    htmltools_0.5.7
## [33] fastmap_1.1.1      highr_0.9          htmlwidgets_1.5.3  rlang_1.1.2
## [37] readxl_1.3.1       TTR_0.24.3         rstudioapi_0.13    quantmod_0.4.18
## [41] shiny_1.8.0        farver_2.1.0       generics_0.1.3     zoo_1.8-9
## [45] jsonlite_1.8.8     dplyr_1.0.6        zip_2.1.1          car_3.0-10
## [49] magrittr_2.0.3     Rcpp_1.0.9         munsell_0.5.0      fansi_0.5.0
## [53] abind_1.4-5        lifecycle_1.0.4    stringi_1.7.8      yaml_2.2.1
## [57] carData_3.0-4      MASS_7.3-55        grid_4.1.2         promises_1.2.0.1
## [61] forcats_0.5.1      crayon_1.4.1       lattice_0.20-45    haven_2.4.1
## [65] hms_1.1.0          knitr_1.33         pillar_1.6.1       ggpubr_0.4.0
## [69] ggsignif_0.6.1     glue_1.6.2         evaluate_0.23      data.table_1.14.0
## [73] vctrs_0.6.5        httpuv_1.6.1       cellranger_1.1.0   gtable_0.3.0
## [77] purrr_1.0.2        tidyr_1.1.3        assertthat_0.2.1   xfun_0.34
## [81] openxlsx_4.2.3     mime_0.10          xtable_1.8-4       broom_0.7.6
## [85] pracma_2.3.3       rstatix_0.7.0      later_1.2.0        ragg_1.2.1
## [89] viridisLite_0.4.0  signal_0.7-7       tibble_3.1.2       ellipsis_0.3.2
```
